# Spatio-temporal shifts driven by climate change threaten persistence and resilience of honey bee populations

**DOI:** 10.64898/2026.03.25.713998

**Authors:** Mert Kükrer

**Affiliations:** Biology Department, Middle East Technical University, Ankara, Turkey; Molecular Biology and Genetics Department, Kilis 7Aralik University, Kilis, Turkey

## Abstract

Understanding how climate shapes intraspecific genetic turnover is critical for predicting biodiversity responses to global change, yet such analyses remain limited for systems where natural adaptation and human-mediated dispersal jointly structure diversity. Here, we investigate the spatio-temporal dynamics of genetic composition in the western honey bee (*Apis mellifera*) across Anatolia and Thrace, a major historical refugium harboring five subspecies. Using a dataset of 672 individuals genotyped at 30 microsatellite loci, we characterize population structure and model ancestry compositions as a function of environmental and geographic variables. We integrate Gradient Forests and Generalized Dissimilarity Modelling to identify key climatic drivers of intra-specific turnover and project future changes under multiple CMIP6 climate scenarios.

We detect five major ancestral groups with widespread admixture structured by both spatial processes and environmental gradients. While geographic distance explains a substantial proportion of variation, climatic variables account for a large fraction of ancestry turnover. Spatial projections reveal distinct ecological regions corresponding to subspecies distributions, with high turnover zones aligned with major geographic and ecological barriers.

Climate projections indicate substantial restructuring of ancestry compositions over the 21^st^ century. Most ancestral groups show declines in persistence and resilience, whereas lineages associated with warmer and drier conditions expand under future scenarios. Regions of high uniqueness and refugia contract, while areas experiencing rapid turnover and novel ancestry compositions increase. Existing Genetic Conservation Areas provide incomplete representation of diversity and are projected to lose effectiveness under future climates.

Our results demonstrate that climate change is likely to disrupt spatial genetic structure, promote admixture, and threaten persistence and resilience of honey bee populations. By modeling ancestry composition as a multidimensional proxy for genetic variation, for the first time to our knowledge, this study provides a scalable framework for forecasting intraspecific biodiversity dynamics and informing conservation and management strategies under global change.

## INTRODUCTION

In the face of ongoing global change, biogeography has evolved into a dynamic field for understanding species’ responses to environmental transformations (Parmesan, 2006). As habitats shift, ranges fluctuate, and ecosystems rapidly change, spatially explicit genetic data enables the exploration of mechanisms shaping distributions (Joost et al., 2007; Gienapp et al., 2008; Eckert et al., 2008). However, identifying ecological patterns associated with genetic clines can be challenging (Jones et al., 2013). Nevertheless, approaches like Gradient Forests (GFs) and Generalized Dissimilarity Modeling (GDM) that account for non-linear interactions between environmental variables and ancestry compositions can be helpful (Ferrier et al., 2007; Ellis et al., 2012). GFs and GDM have been used in studying biodiversity at several layers, from ecosystems, communities, and species to populations, morphological traits, and genes (Bay et al., 2018; Mokany et al., 2019a, 2019b; Morgan et al., 2020; Gougherty et al., 2021; Fitzpatrick et al., 2021). Once biodiversity at any level is modeled as a function of multiple environmental gradients, the resulting models can be used to develop forecasts under global climate change scenarios (Fitzpatrick et al., 2011; Fitzpatrick & Keller, 2015), including invasion risk assessments (Chen et al., 2024).

The western honey bee (*Apis mellifera* L.) is a flagship species with a wide geographical range and significant ecological and economic roles (Franck et al., 2001; Meixner et al., 2010; Iwasaki & Hogendoorn, 2021). As generalist, widespread, and efficient pollinators, honey bees provide insights into ecosystems and help monitor community diversity and sustainability (Quigley et al., 2019; Cunningham et al., 2022). Moreover, spatio-temporal analyses on honey bee models can be applied to various species and biological questions, including invasive species adaptation (Dearden et al., 2009; Giordano et al., 2022). Despite human-mediated gene flow, environmental features and genotype-environment interactions often continue to shape the spatially structured genomic diversity of honey bees (Wallberg et al., 2014; Harpur et al., 2014; Cridland et al., 2017, 2018; Wragg et al., 2018; Parejo et al., 2020; Dogantzis et al., 2021; Gmel et al., 2023). Nevertheless, anthropogenic factors are also shown to influence gene flow, with queen/colony trade and migratory beekeeping creating mobile hybrid zones in space and time (Kükrer et al., 2021). While allelic divergence, selection candidates, and gene-environment associations are well-studied (Fuller et al., 2015; Chen et al., 2016, 2018; Wallberg et al., 2017; Henriques et al., 2018; Montero-Mendieta, 2019; Christmas et al., 2019; Ji et al., 2020; Cao et al., 2023), the role of climate in compositional turnover among honey bee populations remains understudied.

Climate regulates various processes on Earth, such as ecosystem productivity, and sustains life, including those of humans (Howden et al., 2007; Willis & Bhagwat, 2009; Bellard et al., 2012). As climate change disrupts these processes, understanding the interconnected relationships becomes essential to mitigate negative impacts, including local extinctions, changes in distribution ranges of species and ecosystems, community compositions, and ecosystem functioning (Thuiller et al., 2019; Babcock et al., 2019; Román-Palacios & Wiens, 2020; Pörtner et al., 2022). Additionally, climate change influences species invasions as invasive species exploit altered environments to spread, destabilizing native ecosystems (Dukes & Mooney, 1999; Bellard et al., 2016). High climate vulnerability often leads to economic damages and potential food insecurity, primarily affecting smallholder operations where environmental fluctuations amplify challenges (Cohn et al., 2017; Coronese et al., 2019). Mitigating economic losses and supporting food security can be enhanced by recognizing locally adapted geographic forms. In the case of honey bees, these forms may exhibit higher yield, improved colony development, enhanced performance, increased survival, lower pathogen levels, infrequent occurrence of diseases, and broader expression of desirable traits in swarming, defensiveness, or hygiene.

Quantitative studies on the impacts of global change on honey bees and beekeeping are disproportionately few relative to the severity of threats they face (Gordo & Sanz, 2006; Kovac & Stabentheiner, 2011; Delgado et al., 2012; Howlett et al., 2013; Wang et al., 2016; Langowska et al., 2017; Flores et al., 2019; Nürnberger et al., 2019; Rowland et al., 2021; Becsi et al., 2021). Furthermore, there is a notable absence of forecasts regarding the diverse impacts of environmental transformations on honey bees, including spatio-temporal analyses of intra-specific turnover (Kükrer & Bilgin, 2020). The growing interest in ecological forecasting stems from the urgency to provide vital information on future population, community, and ecosystem states to enhance conservation, management, and adaptation strategies (Petchey et al., 2015). A recent systematic review of climate change impacts on honey bees and beekeeping revealed significant negative effects on honey bee ecology and physiology, such as food reserves, plant-pollinator networks, mortality rates, gene expression, and metabolism (Zapata-Hernández et al., 2024). The assessment identified several knowledge gaps, including limited predictive studies and a lack of comprehensive climate analyses. Most studies focused on individual bee behavior rather than population dynamics and were conducted at short spatial (<10 km) and temporal (<5 years) scales, limiting their applicability to large-scale assessments. Additionally, environmental analyses have predominantly relied on short-term weather data rather than long-term climate trends, complicating efforts to forecast future impacts.

To assess honey bee diversity across temporal and spatial scales, we first characterize neutral genetic diversity and population structure across historical refugia in and around Anatolia, a natural laboratory with diverse bee habitats and vast environmental heterogeneity (Hewitt, 1999; Sönmez, 2022). Turkey hosts five native honey bee subspecies that meet, exchange genes, and adapt to local conditions, creating a unique experimental setting by blending genetic elements from Europe, Asia, and Africa (Kandemir et al., 2006; Kükrer et al., 2021). Deploying GFs, we identify key drivers of intra-specific turnover by modeling genetic composition as a function of climate and geography. Then, we apply GDM to site-pair dissimilarities in environmental variables selected by GFs to model ancestry estimates treated as relative abundances and serve as response variables. Finally, we conduct forecasts and spatio-temporal analyses to predict vulnerability and evaluate the persistence, resilience, and conservation efficacy of native populations to inform the management of honey bees.

## MATERIALS AND METHODS

### 2.1 Sampling and genotyping

We collected 460 honey bee samples from 392 localities in 75 provinces in Turkey between May 2016 and November 2019, covering the natural ranges of five subspecies: O lineage bees from *A. m. syriaca*, *A. m. caucasica*, *A. m. anatoliaca*, *A. m. meda*, and the Thracian ecotype of C lineage. Additional samples included 45 *A. m. carnica* from Austria and Germany, 12 *A. m. caucasica* from Georgia, and 174 samples from stationary beekeepers previously collected across Turkey (Kükrer, 2013; Oskay et al., 2019). In total, 691 samples were obtained (**Fig. 1a**). We isolated DNA from bee heads, grouped 30 microsatellite loci into four sets (see **Supplementary Table 1** for sample metadata, markers, and genotypes), amplified markers, determined fragment sizes, and binned the microsatellite alleles as specified in Kükrer et al. (2021). We excluded one locus due to inconsistent amplification, estimated genotyping errors by blind double-genotyping, and removed samples with sibling status or insufficient loci, leaving 672 samples for analysis. See **Supplementary Text** for details.

### 2.2 Exploring genetic diversity and population structure

We estimated null allele frequency, allelic richness, observed and expected heterozygosity, linkage disequilibrium, and other diversity metrics at the loci and population levels. We constructed phylogenies by UPGMA (Unweighted Pair Group Method with Arithmetic Mean) based on population genetic distances (Reynolds et al., 1983), visualized by the online tool Interactive Tree of Life v5 (Letunic and Bork, 2021). We examined population structure via genetic fixation index (F_st_), Analysis of Molecular Variance (AMOVA), Principal Component Analysis (PCA), and discriminant and spatially explicit versions of it. We estimated individual membership coefficients using Structure 2.3.4 (Pritchard et al., 2000). Based on these analyses, we identified five ancestral groups corresponding to the subspecies and removed 80 samples with obvious mismatches to spatially expected ancestry. After this filtering according to admixture patterns (i.e., the particular composition of ancestry estimates at a specific site), 592 samples were left for downstream analyses. Then, we interpolated ancestry estimates on geographic maps using a kriging model (Jay et al., 2012). See **Supplementary Text** for details.

### 2.3 Modeling intra-specific turnover and predicting ancestry compositions

The kriging model omits interactions between ancestry compositions and the environment besides geographic distance. Nevertheless, population structure and environmental factors interplay in more complex ways than isolation by distance alone. Therefore, we modeled intra-specific turnover to identify climatic and geographic drivers of admixture variation. We fitted Gradient Forests (GFs) and Generalized Dissimilarity Models (GDMs) to ancestry estimates of 554 individuals in and around Anatolia, excluding the 38 reference *carnica* samples from Austria and Germany to focus on the study area. We fitted the models using 19 bioclimatic variables from WorldClim 2.1 in 2.5-minute spatial resolution (Fick & Hijmans, 2017) and 18 climatic and topographic variables from the Envirem dataset (Title & Bemmels, 2018). These datasets describe annual and seasonal temperature and precipitation trends and extreme or limiting climatic factors. We retrieved altitude data from The Shuttle Radar Topography Mission (SRTM) at 90-meter resolution (Jarvis et al., 2008). We applied a 3 km buffer distance for each variable, aligning with the effective foraging radius of a worker bee (Haldane & Spurway, 1954; Visscher & Seeley, 1982). We summarize complete variable names, abbreviations, and units in **Table 1**.

GF is a machine learning approach that models compositional turnover by fitting regression trees to create cumulative importance functions of predictor variables and to assess how split values along a gradient explain biological variation. We fitted GFs as implemented in extendedForest 1.6.1 and gradientForest 0.1-32 (Ellis et al., 2012) to logit-transformed ancestry estimates to identify rapid changes and the most important environmental gradients. Since GFs can’t directly incorporate geographic distances, we included spatial effects using the first two Moran’s Eigenvector Maps (MEMs) with cumulative adjusted R^2^ values of 0.9, calculated from the geographic coordinates. We hypothesized that if local adaptation significantly influences genetic differentiation, the cumulative R^2^ of environmental predictors could exceed that of MEM variables. We expected *caucasica* ancestry to respond strongly to moisture-related predictors, reflecting adaptation to cool and moist environments during the flowering season, and *syriaca* ancestry to temperature-related predictors, reflecting adaptation to hot Mediterranean climates. Recognizing that environmental factors may vary between subspecies transition zones, we constructed additional regional GF models with relevant samples and ancestry components. We considered important variables with weighted R^2^ values larger than 0.01 in this basal GF analysis and included them in subsequent GDM processes as potential predictors.

GDMs explain biological variation as a function of climate and geography, identifying environmental gradients linked to compositional turnover. By relating biological distance to differences in environmental conditions or physical isolation, GDMs can infer where turnover is rapid along each gradient. We used gdm 1.5.0-9.1 (Fitzpatrick et al., 2022) to fit GDMs and infer admixture patterns across the area. We fitted additional regional models focusing on transition zones between ancestral group pairs. To address multicollinearity and select independent variables for our models, we applied an elimination procedure based on the variance inflation factor (VIF) by setting a correlation threshold of 0.75 and a VIF threshold of 5. We hypothesized those environmental predictors would explain more variance in compositional turnover than geographic distances alone if local adaptation were to play a major role in genetic differentiation. We spatially projected these GDMs, serving as surrogates for ancestry compositions across a densely sampled landscape. To visualize the transformed environmental variables, we used PCA to reduce outputs into synthetic variables, representing them in a red-green-blue color palette. We cross-validated our global model ten times, training with 90% of the data and testing on the remaining 10%. We grouped predictors into temperature, precipitation, or interaction (tXp) variables and evaluated variance partitioning.

### 2.4 Spatio-temporal analysis of biodiversity patterns and conservation implications

Once a spatial model based on transformed variables is established, it can predict ecological similarity between sites under current or future conditions. Using our dissimilarity model, we analyzed spatio-temporal variation in ecological distances to address questions about honey bee diversity patterns. Ecological similarity computations were derived and slightly modified from Mokany et al. (2022).

#### 2.4.1 Survey gaps, uniqueness, and turnover speed

First, we assessed potential survey gaps within our study area by recording the pairwise ecological similarity between each raster grid cell and its most similar sampling site, with lower scores indicating higher potential gaps. We evaluated each cell’s uniqueness by its mean ecological similarity to random reference cells covering 5% of the study area. We hypothesized that core regions where subspecies are found in unadmixed forms would show high uniqueness while transitional zones would show moderate values. We calculated the turnover speed at each site as the mean ecological similarity of the corresponding cell to its neighbors within a 0.5-degree radius. We predicted that geographic or ecological barriers to gene flow would exhibit fast turnover sites, indicating isolation by barriers (IBB) or environment (IBE). Regions like the Taurus Mountain Range, the Sea of Marmara and the Straits, the Anatolian Diagonal, and the East Anatolian Plateau might show high turnover rates. Conversely, we expect sites with low turnover speed to align with an isolation by distance (IBD) pattern.

#### 2.4.2 Ancestral group similarity and hierarchical classification

We used unadmixed sampling sites with ancestry estimates over 0.85 in one of the five ancestral groups to assess changes in ecological similarities across the study area. We calculated the mean ecological similarities to these unadmixed reference cells for each ancestral group and cell and constructed affinity maps. We used a supervised approach to classify the study area into five clusters corresponding to the five subspecies. First, we calculated pairwise ecological similarities between unadmixed reference cells and carried out a hierarchical clustering. Then, we assigned each cell to the most similar cluster. We repeated this at the six- and seven-cluster levels to explore potential ecotypes below the subspecies level. To evaluate the effectiveness of our method, we also used random reference cells and derived hit rates for both approaches in predicting the dominant ancestral group at sampling sites with ancestry estimates exceeding 0.625 (representing an F1 crossed to a backcrossed hybrid at a putative hybrid zone).

#### 2.4.3 Genetic Conservation Area resemblance and complementarity

Currently, eight Genetic Conservation Areas (GCA) in Turkey restrict migratory beekeeping and queen/colony sales, located in Adıyaman, Ardahan, Artvin, Düzce, Hatay, İzmir, Kırklareli, and Muğla, respectively. GCAs house breeding and conservation apiaries containing a representative sample of native colonies from each corresponding province. To evaluate representation, we calculated the ecological similarities of each cell in the study grid to those within the GCAs, recording the maximum value as the resemblance index. We iterated this procedure three times, adding hypothetical GCAs (Hakkari, Çankırı, and Muş) to enhance complementarity and representativeness. We compared resemblance indices under different conservation scenarios using a paired t-test and visualized stepwise resemblance gains. For each scenario, we compared the total protected surface area directly in GCAs or indirectly through a resemblance index above 0.7. Finally, we assessed the differential impact of additional GCAs on resemblance values within the area classified to each ancestral group using ANOVA followed by Tukey’s test.

#### 2.4.4 Temporal analyses

In the temporal analysis, we used six CMIP6 climate projections known for their short-term accuracy and varying long-term equilibrium climate sensitivity: CNRM-ESM2-1, INM-CM4-8, MPI-ESM1-2-HR, MIROC6, EC-Earth3-Veg, UKESM1-0-LL (Tokarska et al., 2020; Meehl et al., 2020). We computed composite means of WorldClim variables for Shared Socioeconomic Pathways (SSPs) 126, 245, 370, 585, and mid-years 2030, 2050, 2070, 2090. We then calculated Envirem variables from monthly average minimum/maximum temperatures and total precipitation.

We predicted 16 SSP-time period combinations across the study area using the dissimilarity model. We then assessed temporal changes in survey gaps, uniqueness, turnover speed, ancestral group similarities, and classification outcomes. We conducted paired t-tests for cell-level comparisons. We considered sites under-sampled if their gap values were below 0.45, unique if their uniqueness was below the 10^th^ percentile, fast turnover if within the 10^th^ percentile of turnover speed values, and high correspondence if their ecological similarity to any ancestral group exceeded 0.4. We hypothesized that climate change-induced impacts would manifest distinctly across the landscape. First, we anticipate increased survey gaps due to inadequate ecological gradient coverage, reflecting an increased mismatch between the characteristics of current sampling sites and future conditions prevailing in the study area. Second, we predict lowlands in cooler climates will face invasions by populations preadapted to hotter, arid conditions, leading to decreased uniqueness and dramatic shifts in ancestry compositions and classification outcomes within affected areas. Additionally, we expect an increase in fast turnover sites due to an expanded interface between ancestral groups occupying different altitudes.

#### 2.4.5 Persistence, resilience, disappearance, and emergence

To further evaluate climate change impacts on honey bee diversity, we employed four additional indices: persistence (inverse offset), resilience (refugia value), disappearance (forward offset), and emergence (reverse offset). For the persistence index, we calculated the future ecological similarity of each cell to itself to derive offset values. To avoid divisions by zero, we added 1 to the summed offset values before averaging across scenarios, measuring consistent durability in ancestry compositions. Since offsets inversely relate to persistence, we took the inverse of the final value. For the disappearance index, we calculated the ecological similarity of each cell under current conditions to all future cells in the random reference set, recorded the maximum value for each scenario, and averaged these to measure the continuous loss of site-specific ancestry compositions across SSPs and periods. For the emergence index, we calculated the ecological similarity of each future cell to the current random set, recorded the maximum value for each scenario, and averaged these as a measure of consistent novelty in ancestry compositions. For the resilience index, we calculated the ratio between the mean ecological similarity of each future cell to its 0.5-degree radius neighbors under current and future conditions. Deriving this resilience index, a ratio higher than 1 indicates higher ecological similarity to neighbors’ current conditions, suggesting a high value as a potential refugium. We averaged these ratios across SSP-period combinations to identify continuous refugia for ancestry compositions.

Sites with high resilience had refugia values greater than 1, low persistence if their average offset values were in the 4^th^ quartile, and high disappearance or emergence if their maximum similarity to reference cells was below 0.6. A decline in persistence and resilience indices, along with an uptick in disappearance and emergence indices over time, supports the assertion that climate change poses a significant threat to honey bee biodiversity. Climate change could impact different ancestral groups in various ways and show diverse effects across spatial scales or protection statuses. Disproportionate impacts within GCAs or unique sites may indicate conservation shortfalls. Thus, we analyzed how persistence, resilience, disappearance, emergence, and resemblance indices varied with spatial patterns. We checked if index values differed between densely/sparsely surveyed, unique/generic, fast/slow turnover, protected/unprotected sites, and those with high/low ecological similarity to any ancestral group or area classified to any of the clusters. We used a Bonferroni corrected t-test for each index and ANOVA followed by Tukey’s test for classification outcomes to infer interactions between indices and spatial patterns. We include detailed information regarding the R packages (R Core Team 2022 version 4.2.2) and the session information in **Supplementary Table 2**.

## RESULTS

### 3.1 Population structure points to distinct ancestral groups but also widespread admixture

Our exploration of population structure revealed multiple population profiles and substantial differentiation among honeybee populations. Using clustering methods and population genetic analyses, we identified distinct ancestral groups and widespread admixture across the landscape. In the phylogeny, the European population diverged first, followed by Thrace and East and South Marmara (**Fig. 1b**). Clustering analysis unveiled five main ancestral groups: Thracian, Anatolian, Caucasian, Levantine, and Zagrosian (**Fig. 1c**). At K = 2, individual membership coefficients differentiated samples from Europe and Asia, with Thracian samples appearing as a mixture of the two gene pools. At K = 5, Thracian samples formed their own cluster. By K = 7, all subspecies were observable, along with a spurious cluster within the *anatoliaca* range (see **Supplementary Text** for a more detailed analysis of genetic diversity and population structure). Populations with high Anatolian ancestry were mainly in western Turkey; still, those neighboring Thrace, Caucasus, Levant, and Zagros showed high ancestry estimates from these groups (**Fig. 1d**). Interpolation of ancestry estimates provided a detailed view of gradual admixture patterns, highlighting core regions where subspecies are found in unadmixed forms and adjacent transition zones. The kriging outcome also revealed areas where changes might be rapid (**Fig. 1e**). Notably, the transition between Anatolian and Zagrosian ancestral groups in East Anatolia deviated from gradualness, with Zagrosian ancestry tracking the Araxes valley and Anatolian ancestry following the Murat and Karasu rivers, the two main tributaries of the Euphrates.

### 3.2 Environmental gradients identified by GF models drive intra-specific turnover

Model performances for distinct ancestral groups in the global GF model averaged 0.64, ranging from 0.41 for the Zagrosian cluster to 0.74 for the Levantine cluster (**Supplementary Table 3**). The model identified the first two MEMs as the most important predictors of intra-specific turnover in ancestry compositions (**Fig. 2a** and **Supplementary Table 4**). The vital role of MEM1 and MEM2 (R^2^ values of 0.12 and 0.18) suggests the importance of spatial location or other unmeasured environmental predictors. Spatial MEMs had the highest relative contribution, but climatic variables still constituted 53% of the captured R^2^. When limited to selected variables with R^2^ over 0.01, the relative contribution of climatic factors was 40% (0.20 over 0.50). PETwettest, Pwarmest, Tdriest, and isothermality followed the MEMs with the highest R^2^-weighted importance. Other important variables included aridity, Tseasonality, TUR_alt, continentality, Pdriest, Pwettest, minTwarm, PETdriest, and Pcoldest. Thracian ancestry responded strongly to PETwettest (<30mm/month), TUR_alt (<500m), and Pcoldest. In contrast, Caucasian ancestry showed the highest sensitivity to Pwarmest (>150mm), Tdriest, aridity, Pwettest, and PETdriest (**Fig. 2b**). Zagrosian, Levantine, and Anatolian ancestral groups responded similarly to most predictors except isothermality and Tseasonality for Zagrosian cluster and minTwarm for Levantine cluster (>20°C).

### 3.3 GDMs disclose major climatic predictors of environment-ancestry composition relationships

The global GDM included geographic distance and seven environmental variables as predictors (Pdriest, minTwarm, Pwettest, PETdriest, continentality, PETwettest, and isothermality) after controlling for multicollinearity. These variables influenced predicted dissimilarities with summed coefficient values between 0.1 and 1.1. The mean dissimilarity between site pairs with identical predictor values was 0.17. Predictor variables, including geographic distance, explained 27.0% of the deviance with a mean absolute error of 0.19 in ten times cross-validation. The seven climatic variables combined accounted for 15.6% of the deviance, with 5.5% for temperature (isothermality, minTwarm, and continentality), 6.8% for precipitation (Pwettest and Pdriest), and 8.6% for tXp (PETdriest and PETwettest) related variables. Turnover was most sensitive to geographic distance and Pdriest, followed by minTwarm, Pwettest, and PETdriest (**Fig. 2c**). The least influential environmental variables—continentality, PETwettest, and isothermality—are associated more with transitions from Thracian to Anatolian or Anatolian to Zagrosian ancestral groups, which are more gradual than transitions to Caucasian or Levantine ancestral groups (**Fig. 3a**). While geographic distance was the most important variable in the global model, it dropped to second or third in regional models (see **Supplementary Text** for details about regional GFs and GDMs).

### 3.4 Spatial analyses unveil ecological patterns and turnover dynamics

Based on our GDM, we classified the study area via hierarchical clustering supervised by unadmixed reference cells and identified five primary bioregions representing ancestral groups within the study area (**Fig. 3b**). Our supervised classification approach proved accurate with a hit rate of 86% (n = 389), compared to the 59% accuracy of the approach based on random reference cells. The majority of the study area had low gap values, showing strong ecological consistency with the sampling sites (mean maximum similarity 0.78, SD = 0.07), enabling a robust analysis of the interplay between ancestry compositions and environmental conditions (**Fig. 4a**). Sites that deviated from the broader ecological context, identified as highly unique, corresponded mainly to core zones where subspecies were unadmixed (**Fig. 4b**), suggesting localized ecological conditions driving distinct adaptations. Spatial analysis revealed two regions with exceptionally high turnover speeds: the Taurus Mountains, which act as a physical barrier to gene flow, and the East Anatolian Plateau, indicating rapid ecological transitions due to dynamic environmental gradients or ecological boundaries (**Fig. 4c**).

### 3.5 Assessment of GCA resemblance prompts recognition of new conservation sites

Established GCAs encompassed all ancestral groups except Zagrosian (**Fig. 5a**). Adding further GCAs in Hakkari, Çankırı, and Muş significantly improved resemblance (**Fig. 5b, 5c, and 5d**), increasing protected area coverage from 59,711 km² to 81,949 km² (**Fig. 5e**). The indirectly protected area with high resemblance to GCAs expanded from 499,719 km² to 754,809 km², nearly covering Turkey’s entire surface area (**Supplementary Table 5**). The mean gain per cell in resemblance was 0.023 with Hakkari, 0.018 with Çankırı, and 0.007 with Muş (all p < 0.001), totaling nearly 0.05. Initial resemblance differences between ancestral groups were significant (all p < 0.001). Thracian and Caucasian groups are currently best represented, followed by the Anatolian group, with mean differences of 0.06 and 0.04 compared to the best-protected groups. The Zagrosian group is least represented, with mean differences of 0.12 and 0.11. Adding Hakkari reduced these to 0.05 and 0.04 (both p < 0.001). Including Çankırı reduced the Thracian-Anatolian difference to 0.02 (p < 0.001), equalizing Caucasian and Anatolian representation. After adding Muş, the resemblance of Zagrosian, Anatolian, and Caucasian groups equalized, each staying 0.02 below the Thracian group (p < 0.001). Across all scenarios, there was minimal improvement in the Levantine group’s resemblance (**Supplementary Table 6**).

### 3.6 Temporal analyses point to shifts in turnover patterns and the vulnerability of ancestral groups

Ecological boundaries remained relatively stable throughout the first half of the century. However, divergence emerged among the models in the latter half, especially under more pessimistic SSP scenarios (**Fig. 3c and 3d**). The impacts of climate change varied across the study area. Thrace experienced substantial early changes, followed by disruptions in the Caucasus. Later, divergence intensified between the highlands and lowlands within Anatolia. Coastal regions were initially more vulnerable to climate-induced changes, but inland areas were similarly affected later under intense SSP scenarios. Increases in minTwarm were a primary driver of these changes, along with significant impacts from PETdriest. SSP-period combinations showed variation in scenarios, with the Thracian group shrinking drastically in some, nearly disappearing within Turkey’s borders. High losses of ecological similarity were apparent in rasters associated with the Caucasian group, indicating retreat and fragmentation, particularly at lower altitudes. This pattern was also observable in the Anatolian and Zagrosian rasters, albeit to a lesser degree. In the late SSP585 scenario, the Levantine group gained excessive ground in central Anatolia and the central Black Sea. We compare spatial dynamics in SSP-period combinations in the **Supplementary Text**.

Ecological similarities to each ancestral group revealed a consistent decline in cells similar to any group, except for the Levantine group, which initially declined but later gained (**Fig. 6a**). By the end of the projection period, the number of cells showing high ecological similarity to unadmixed Thracian or Caucasian samples was halved. Although regions with high similarity to the Zagrosian group declined, the Anatolian group experienced the most drastic decrease in total surface area. Despite this decline, the Anatolian ancestral group maintained its classified area until the late stages of the projection period (**Fig. 6b**). The Thracian group steadily decreased from over 50,000 km² to 25,000 km², the Caucasian group from c. 100,000 km² to 75,000 km², and the Zagrosian group from c. 200,000 km² to 150,000 km². Conversely, the Levantine group nearly doubled its size from c. 175,000 km² to 300,000 km² by the end of the projection period, with total classification changes across the study area reaching 150,000 km².

Survey gaps notably increased, especially in areas of higher climate impact (**Fig. 4d**). There was a significant decrease in maximum ecological similarity to sampling sites, with similarity loss averaging around 0.06 per cell (p < 0.001). The proportion of the study area with survey gap values exceeding the threshold increased continuously under each scenario throughout the years, ranging from nearly none to 50,000 km² by the end of the century (**Fig. 6c**). Cell-level uniqueness increased slightly, averaging around 0.03 (p < 0.001), while the total area classified as highly unique consistently decreased from c. 100,000 km² to 75,000 km². Uniqueness declined consistently at Levant and Thrace, and the coherence of highly unique locations in the Caucasus was disrupted (**Fig. 4e**). Turnover speed did not exhibit substantial changes at the cell level. However, the proportion of sites with fast turnover increased across the study area, their surface area rising from c. 100,000 km² to 150,000 km². Notable changes in turnover patterns included a shift of the geographical barrier at the mid-portion of the Southeastern Taurus range some 125 kilometers north to the Munzur Mountains in East Anatolia (**Fig. 4f**). The Aegean mountainous areas and coastal regions along the Eastern Black Sea also displayed increased turnover speeds due to rising temperatures.

### 3.7 Climate change dynamics disrupts persistence and threatens resilience across spatial and temporal scales

Highly persistent sites and sites displaying high resilience declined over time and across scenarios, while sites with exceptionally high disappearance or emergence indices increased (**Fig. 6d**). Novel ancestry compositions emerged later due to intensifying environmental changes. Sites with high persistence decreased from 800,000 km² to 575,000 km², affecting nearly one-fourth of the study area. Low persistence was heterogeneous, with Thrace, Upper Euphrates, Levantine regions, Aegean, central and eastern Black Sea coasts experiencing the highest declines (**Fig. 7a**). East-Central Anatolia showed the highest persistence, followed by the highlands of West and East Anatolia. A Tukey’s test indicated that sites classified to various ancestral groups had significant mean differences (all p < 0.001, **Supplementary Table 7**), with the Thracian group having the lowest persistence scores. Sites with high ecological similarity to Anatolian or Zagrosian groups showed greater persistence (both p < 0.001, **Supplementary Table 8**). Persistence was lower in regions with fast turnover rates and high resemblance to GCAs, whereas highly unique regions had high persistence (all p < 0.001 with mean differences 0.70, 0.21, and 0.35). Under SSP126, persistence loss was limited, while SSP585 showed almost homogeneous low persistence across the study area, except for some restricted sites (see **Supplementary Text** for SSP scenario comparisons).

The area that could serve as refugia declined from 300,000 km² to 250,000 km². Refugium sites concentrated around mountainous areas, especially those surrounded by lowlands or adjacent to low-persistence sites (**Fig. 7b**). Regions classified to Thracian and Levantine ancestral groups suffered from an absence of such places, notably lacking effective refugia (all p < 0.001). In contrast, resilience was slightly higher at sites highly similar to the Zagrosian group and in areas with slow turnover (both p < 0.001 with mean differences 0.08 and 0.07). There were considerable differences across SSPs. In extreme cases, newly established refugia were sometimes overridden, such as in Thrace and Aegean, or even in the central Anatolian plateau and the mid-portion of the Southeastern Taurus range at later stages of the projection period.

Low persistence may lead to the replacement of local ancestry compositions with those from neighboring sites or, at other times, may end up in novel ancestry compositions. The disappearance of current ancestry compositions occurred relatively quickly, and their total extent stabilized at around 25,000 km². In contrast, the emergence of new ancestry compositions became more pronounced in the second half of the century, affecting an area of up to 100,000 km². Sites suffering the most from disappearance included the eastern Black Sea coast, a relatively restricted zone in South Aegean, Mount Uludağ in Marmara, northern Syria, and the southern lowlands of the mid-portion of the Southeastern Taurus range (**Fig. 7c**). Emergence was prevalent in central and eastern Black Sea coast, around the Bosporus, and the Araxes River valley north of Mount Ararat in the easternmost part of Anatolia (**Fig. 7d**). In the late stage pessimistic SSP scenarios, high disappearance was predicted throughout Thrace, entirely replaced by novel ancestry compositions extending to the western Black Sea region. Mean differences in disappearance and emergence indices were significant between sites classified to the Thracian group or others (all p < 0.001). The two indices were slightly larger in regions with fast turnover (both p < 0.001 with mean differences of 0.08).

## DISCUSSION

This study provides insights into the drivers of intra-specific turnover in honey bee ancestry compositions and the complex dynamics of population structure in response to environmental gradients. The findings highlight the importance of global and local factors in shaping genetic differentiation among subspecies, emphasizing the role of specific climatic variables beyond geographic distance in understanding ancestry turnover patterns. The spatio-temporal analyses of climate change impact indicate heightened vulnerability in distinct ancestral groups and underscore the need for new conservation sites to enhance the representation and resilience of honey bee populations. The study highlights potential implications of introgression and maladaptation in the absence of obvious reproductive barriers and has relevance to range shifts and predictions of invasibility and invasive spread.

### 4.1 Global and local drivers of intra-specific turnover

One of the key outcomes of this study was the identification of global and regional drivers of intra-specific turnover. The impact of localized climatic factors on the complexity of genetic differentiation highlights the importance of considering regional contexts and fine-scale ecological patterns (Kim et al., 2023). The global dissimilarity model relied on precipitation and potential evapotranspiration levels during the wettest and driest periods—essential predictors of soil moisture gradients, plant phenology, and community structure (Zhu et al., 2016; Liu et al., 2022; Dudenhöffer et al., 2022). Changes in phenology and composition affect periodic resource availability and diversity, require life-history adaptations, and can drive shifts in insect phenology involving polygenic effects (Alstad et al., 2016; Grünzweig et al., 2022). Among temperature-associated variables, the minimum temperature of the warmest period played a significant role. While extreme maxima define heat tolerance limits and can affect sperm viability and queen failure (Sinervo et al., 2010; McAfee et al., 2020), minimum temperature levels during the warmest period might act as a response threshold for epigenetic, physiological, or behavioral adaptations in thermoregulation (Stabentheiner et al., 2022; Zhang et al., 2022; Alghamdi & Alattal, 2023; Alattal & Alghamdi, 2023). Bees adapted to hotter environments exhibit extended foraging durations and ranges in their natural habitats compared to exotic subspecies (Alattal & Alghamdi, 2015).

Levantine ancestry responded strongly to minTwarm, and moisture-related variables were particularly predictive for Caucasian ancestry. *A. m. syriaca* bees exhibit adaptations to elevated temperatures, including smaller size, lighter coloration, and shorter hair (Ruttner, 1988), while *caucasica* bees are large, dark, and hairy propolis hoarders (Kekeçoğlu et al., 2020). Both subspecies display divergent behaviors adaptive to their native distributions (Brillet et al., 2002; Çakmak et al., 2010; Kence et al., 2013; Claudio et al., 2018; Yıldız & Karabağ, 2022). Additionally, Zagrosian ancestry within the native range of subspecies *meda* was most sensitive to continentality and isothermality. Differential neural, hormonal, and developmental responses across local populations shape the physiological and behavioral plasticity of bees in response to thermal fluctuations (Willmer & Stone, 2004; Grodzicki & Caputa, 2014; Abram et al., 2017; González-Tokman, 2020). Temperature oscillations distinctly affect populations and survival, causing up-regulated stress responses during cooling or heating (Fahrenholz et al., 1989; Torson et al., 2015; Mucci et al., 2021; Kaya-Zeeb et al., 2022). For instance, temperature increase rates influenced the critical thermal limits of *scutellata*-hybrids derived from African bees (Gonzalez et al., 2022). Besides direct impacts on individual and colony levels, thermal fluctuations are associated with plant productivity, pollen richness, land cover, and nutrition (Reitalu et al., 2019; Niemczyk et al., 2021), which can affect foraging and colony health.

Although climatic variables contributed significantly to the captured variance, spatial processes captured by MEMs in GFs and geographic distance in global GDM were the most critical predictors of intra-specific turnover. These results suggest that geographic distance and environmental factors not captured by other predictors have shaped genetic differentiation among honey bee populations, as observed in various species (Vanhove et al., 2021; Lima-Rezende et al., 2022). Physical barriers emerged as another geographic mechanism leading to isolation. Despite small ecological distances and gene flow across the Bosporus and Dardanelles straits, the Sea of Marmara formed a major boundary between western Anatolia and Thrace. Similarly, the transition from *syriaca* to *meda* and *anatoliaca* was primarily aligned with the Taurus Mountains. Consistent with other species, isolation by environment was evident along ecological boundaries and gradients, attributable to rapid ecological changes on the East Anatolian Plateau and the Anatolian Diagonal (Bilgin, 2011; Gür, 2016; Nielsen et al., 2021). When ecological barriers to gene flow are in place, only a limited number of alleles beneficial at both sides of the barrier may introgress (Akerman & Bürger, 2014). Without barriers or strong selection pressure, populations widely admix and homogenize.

### 4.2 Climate vulnerability in the form of declining persistence and resilience

Our temporal analyses of climate change impact raise concerns about the persistence and resilience of honey bee diversity. Persistence is uneven across the study area, indicating that local populations may experience changes in ancestry compositions and suggesting current geographically structured genetic diversity is vulnerable to climate change. Shrinking regions of exceptional uniqueness, dramatic declines in the proportion of sites with high similarity to any of the ancestral groups, and physical shifts at sites of fast turnover and along transition zones all point to wider admixture across the landscape. Climate change-induced hybridization in insect populations can result in introgression, genetic swamping, shifts in hybrid zone boundaries, species fusions, invasions, and local extinctions (Arce-Valdés & Sánchez-Guillén, 2022).

Except for *syriaca* adapted to elevated temperatures, local subspecies consistently shrink under each forecast, with the Thracian ancestral group being particularly threatened. Although physical barriers like the Sea of Marmara and the Taurus Mountains may buffer the spread of Levantine ancestry to Anatolia and Anatolian ancestry to Thrace, bees with Levantine ancestry appear to gain a competitive advantage over time, potentially initiating invasive dynamics. Interestingly, morphometric analyses of honey bee samples from the Jordan Valley, dating back 3000 years, suggest a different geographical distribution of subspecies in the past, with shifts from local *anatoliaca* ancestry to *syriaca* (Bloch et al., 2010). Intriguingly, *syriaca* bees are not favored by beekeepers due to their high defensiveness, tendency to swarm, and low honey yields. So, it is unlikely that this shift was human-mediated but rather was the outcome of past climate alterations. Our predictions about rapid shifts in ancestry compositions align with ecological forecasts for other insects, showing a mix of winners and losers (Neupane et al., 2024). Given their ectothermic nature, insect physiology and trophic or community interactions highly depend on ambient temperatures, rendering them vulnerable to warming and temperature extremes (Chen et al., 2011; Harvey et al., 2020). Furthermore, populations with enhanced survival responses to increased drought frequency and intensity have an evolutionary advantage (Exposito-Alonso et al., 2018).

Potential refugia for current ancestry compositions, concentrating around mountainous areas, decline across years and SSPs. Regions inhabited by Thracian and Levantine ancestral groups lack refugia, impairing resilience with increasing disappearance and emergence indices. Insect populations show diverse resilience through shifts and adaptations amid rapid anthropogenic change (Lancaster et al., 2016; Dudaniec et al., 2018; Halsch et al., 2021; McCulloch & Waters, 2023). Geographically restricted alpine species with limited dispersal face increased extinction risks, while surviving upland insect lineages may rapidly adapt (Kinzner et al., 2019; Shah et al., 2020). In our models, newly established mountainous refugia were sometimes overridden in later stages of pessimistic scenarios. In these instances, ecological similarity between the study area and sampling sites gradually declined. This increase in survey gaps underscores the importance of intensified monitoring and further sampling at those sites. Monitoring intraspecific genetic diversity is crucial for understanding species’ adaptation to changing environments and mitigating climate-induced risks, particularly in vulnerable regions (Pearman et al., 2024).

Climate responses can be asymmetric, often showing sharp declines beyond certain upper thresholds. Projected climate risks are significantly amplified under the SSP585 scenario compared to SSP126, underscoring the urgency for stringent emission controls. Considering the higher equilibrium climate sensitivity in the CMIP6 models, scenarios where warming exceeds 4 °C might not be unrealistic (Lee et al., 2023). Therefore, our results incorporating all four SSPs may serve as a baseline for understanding the climate vulnerability of honey bee populations. This vulnerability and the novel nature of human-mediated gene flow associated with climate change threaten populations alongside ongoing anthropogenic impacts from migratory beekeeping and trade (Kükrer et al., 2021). Although we focused on Anatolia and Thrace for our model system, our approach holds importance for monitoring and conserving managed and wild honey bee populations in neighboring countries such as Azerbaijan, Armenia, Bulgaria, Cyprus, Georgia, Greece, Iran, Iraq, and Syria. While our study addresses managed honey bees, feral populations and wild pollinators might face similar environmental challenges (Jaffe et al., 2010; Requier et al., 2019). Additionally, our modeling approach could benefit both domestic and wild animal and plant species, particularly non-native established and invasive ones in the Mediterranean, where similar pressures drive invasions.

### 4.3 Assessing and enhancing conservation strategies through resemblance analyses

This initial assessment provides essential first steps in genetic monitoring and systematic conservation planning of honey bee populations. It collates specimen and genetic data across the country, identifies conservation goals, evaluates existing conservation sites, and helps expansion designs (Kukkala & Moilanen, 2013). Our findings have significant implications for breeding and conservation management, emphasizing the potential benefits of incorporating genetic and environmental factors to evaluate complementarity and representativeness (Sarkar, 2006). Incorporating environmental variation in conservation decision-making is feasible when cross-taxon surrogates are unavailable (Rodrigues & Brooks, 2007). Hanson et al. (2017) confirmed that environmental and geographic variation could predict adaptive and neutral genetic variation in 27 plant species over the European Alps. Our assessment of GCA resemblance highlights the need to identify and incorporate new conservation sites to enhance ancestral group representation. According to our analysis, *A. m. meda* is currently unprotected, making it an urgent conservation priority.

Our analyses show that freely evolving GCAs may shift in ancestry compositions in response to environmental change. Alarmingly, sites with low persistence overlap with established GCAs. Thrace, the Aegean coast, and parts of the Caucasus range, each housing GCAs, suffer from low persistence. Moreover, sites resembling GCAs exhibit significantly reduced persistence. Breeding programs often depend on locally adapted geographic forms, yet climate change may significantly impact protected areas (Geldmann et al., 2019). Our study indicates that local genetic stocks in conservation sites might be maladapted to future conditions like many other animal and plant species (Hoffmann, 2010; Breed et al., 2013; Henry, 2016; Marsh et al., 2021). When selecting new GCAs, choosing sites with high persistence and resilience indices might be adaptive against invasibility. Assessing the climate efficacy of current GCAs by computing resemblance indices for future rasters could also be beneficial. Additionally, identifying current sites with a lowered resemblance to future GCAs can highlight those with eroding conservation statuses. See **Supplementary Text** for an extended discussion on conservation planning strategies.

### 4.4 Methodological considerations and enhancements

We assume past trends have been stable and current states reflect an optimally balanced situation. Complexity and difficulty in specifying ecological niches make predicting novel system states in response to change challenging. The unpredictability is further exacerbated by computational irreducibility, where evolution may be considered a chaotic process with low intrinsic predictability (Coreau et al., 2009; Doebeli & Ispolatov, 2010; Beckage et al., 2011). We employed six general circulation models across four SSPs and four periods to address challenges associated with future climate unpredictability. Additionally, we implemented a rigorous variable selection procedure, considering variance inflation factors and GF outcomes before modeling dissimilarities. Despite the robustness of our models, forecasting ancestry estimates is challenging due to unknown physiological limits, intragroup genetic diversity affecting adaptive capacities, developmental plasticity, and specific plant-pollinator interactions (Franks et al., 2014; Keeler et al., 2021). Still, our turnover predictions can be seen as an assessment of ongoing pressures and climatic stress on local populations that must cope with environmental transformations. The increased mismatch between existing gene combinations and the environment can undermine resilience.

To our knowledge, we employed GDM to model ancestry estimates and forecast intra-specific turnover in ancestry compositions for the first time. Previous research has mainly relied on genetic distances or differentiation indices, which may underestimate the true magnitude of local adaptation (DeMarche et al., 2019). Instead of reducing population differentiation to a single metric like F_st_, we used ancestry compositions as a multi-dimensional proxy to represent local-scale processes. Forecasting with multi-dimensional ancestry compositions is viable, as neighboring populations with similar environmental constraints to future site conditions may already exhibit preadaptations (Davis & Shaw, 2001). Elevated habitat suitability correlates positively with gene flow across landscapes, supported by both theoretical frameworks and empirical evidence from phylogeographical and landscape genetic studies (Auffret et al., 2017; Knowles & Massatti, 2017; Massatti & Winkler, 2022).

Space-for-time substitution is supported by common garden experiments and fossil data when spatial and temporal models capture comparable climate dissimilarities as in our models (Blois et al., 2013; Lovell et al., 2023). For honey bee subspecies, common garden experiments underline the role of competitive advantage provided by local adaptations (Costa et al., 2012; Hatjina et al., 2014; Büchler et al., 2014; Meixner et al., 2014; Uzunov et al., 2014). Under competitive advantage, *scutellata*-European hybrids derived from non-native African bees introduced to Brazil colonized the New World in less than 50 years, only to be halted at hybrid zones in California and Argentina. Besides, highland populations showed a notable decrease in *scutellata* ancestry compared to lowlands, with ancestry compositions correlating between populations in similar habitats despite a large geographical separation (Everitt et al., 2023). While Calfee et al. (2020) confirmed the genomic cohesion and polygenic basis of the rapid expansion of *scutellata* ancestry and related fitness costs in cooler climates, they faced challenges identifying the precise environmental variables driving the relationship in intra-specific turnover. Our findings bring honey bees to the list of species with comparative studies providing essential evidence on ecological factors and environmental gradients influencing variations in insect abundance and diversity (Blüthgen et al., 2022).

The four vulnerability indices (persistence, resilience, disappearance, and emergence) provide specific averages across scenario-period combinations to understand consistent and continuous impacts. Additionally, we introduce a highly sensitive classification approach, using sites with highly unadmixed samples to predict expected cluster distributions jointly from genetic and environmental variables. This supervised approach can be applied across different taxonomic levels, ecosystems, and communities while classifying biodiversity. Our simplified method for computing forward and reverse offsets is faster and less computationally demanding—a suitable tool for the developing world where computational resources might be limited (see **Supplementary Text** for an extended discussion on the methodology). Our streamlined approach and the provided **Supplementary Code** are instrumental for vulnerability assessment and enable straightforward interpretation.

## CONCLUSION

Our research sheds light on the drivers of intra-specific turnover in ancestry compositions across honey bee populations. Spatial analyses integrating GFs and GDMs provide insights into the diversity patterns and turnover dynamics. Our findings underscore spatial processes and specific climatic variables influential in genetic differentiation. Furthermore, declining persistence and resilience levels reveal the vulnerability of honey bee populations against climate change. These results highlight the urgent need to identify and incorporate new conservation sites to enhance the representation and resilience of ancestral groups. Overall, this study contributes essential knowledge to honey bee biogeography, facilitating informed conservation strategies to safeguard their unique diversity and persistence in the face of global change.

## ACKNOWLEDGEMENTS

We are indebted to the beekeepers and Robert Paxton, who generously provided samples for this study, as well as the various beekeeper associations, ministry personnel, and the Society for Conservation of Environment and Bees, along with all the individuals and entities in the field who supported and facilitated our research. This study was funded by the Middle East Technical University Revolving Funds. This manuscript has been released as a preprint (Kükrer et al., 2026) and incorporates elements from the corresponding author’s thesis (Kükrer, 2024).

---

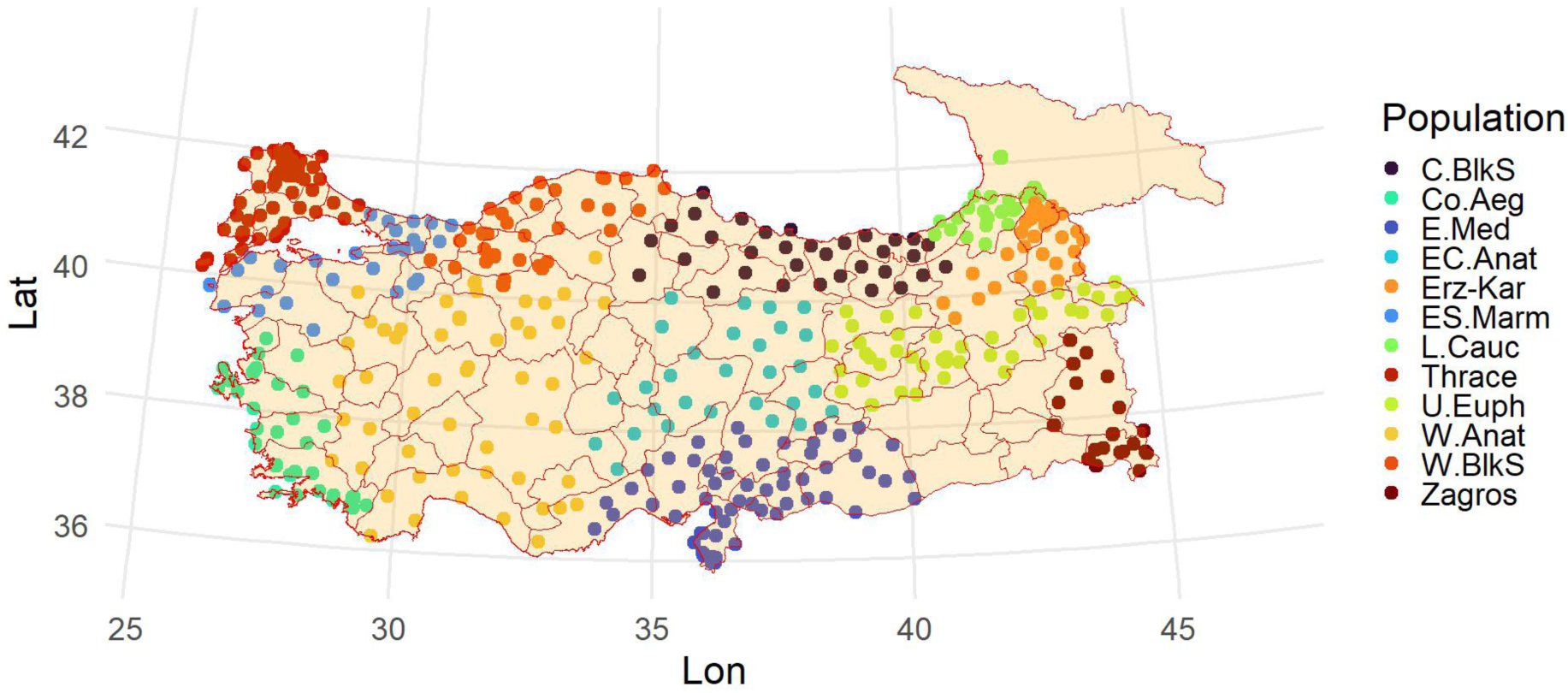

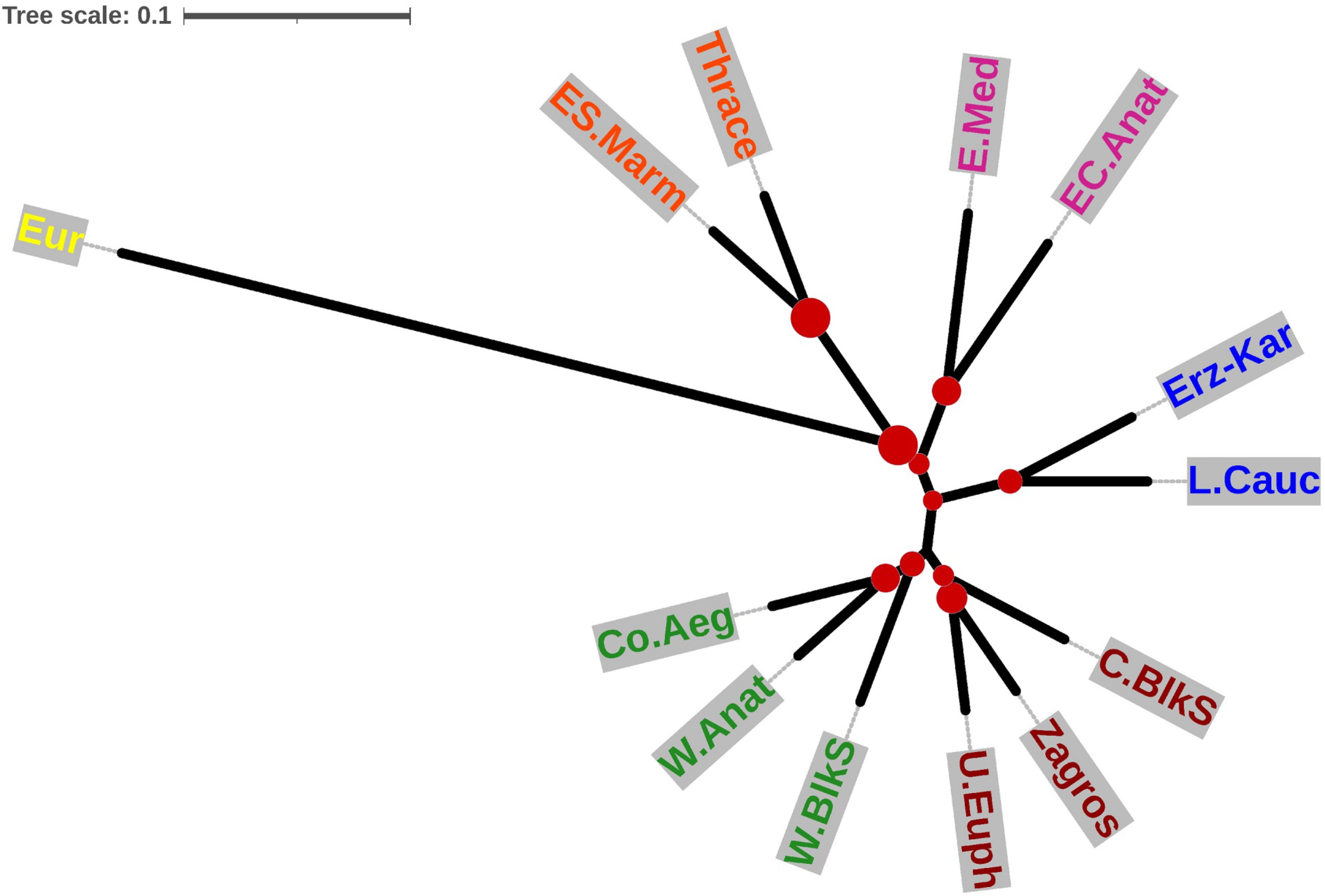

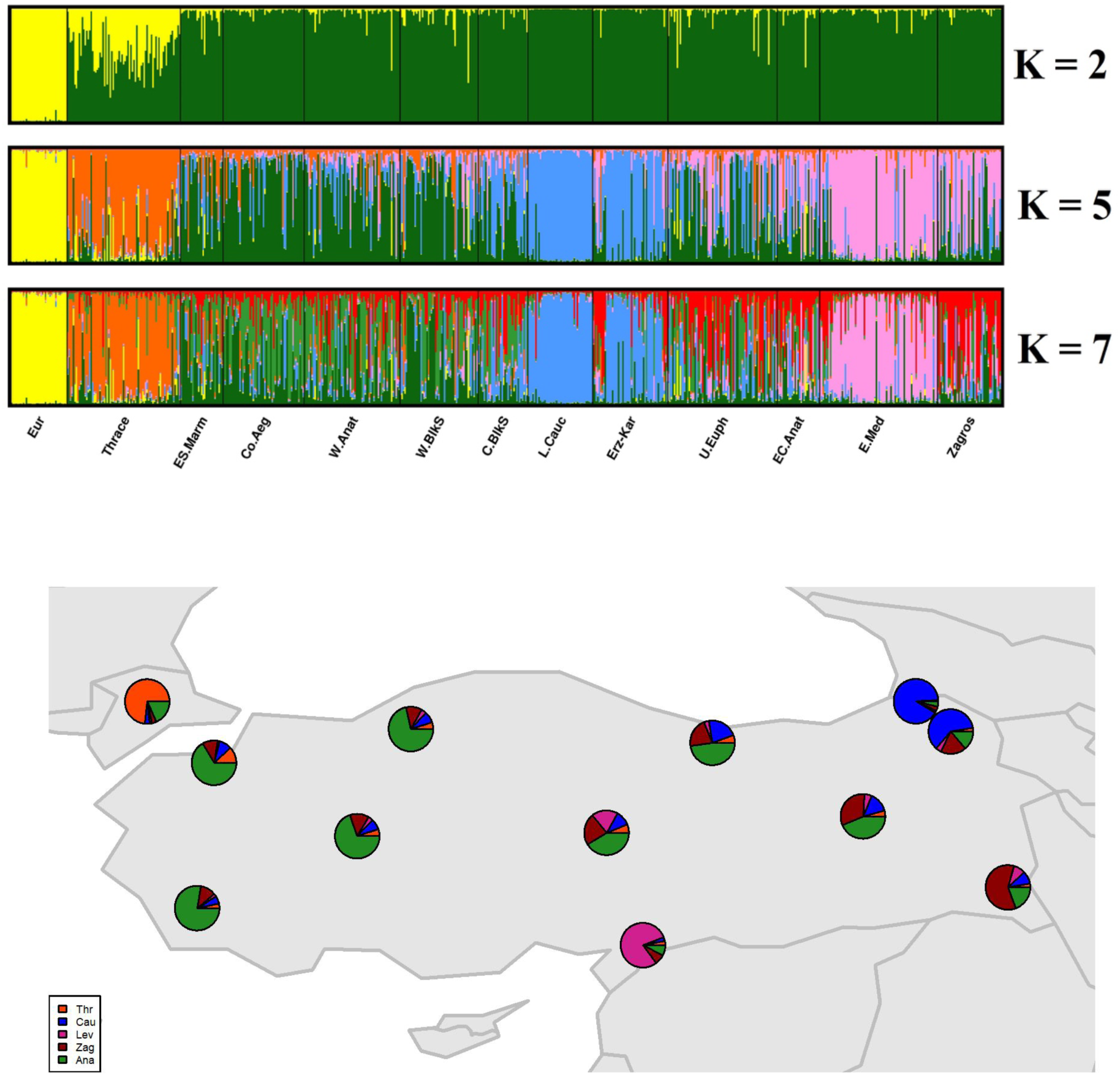

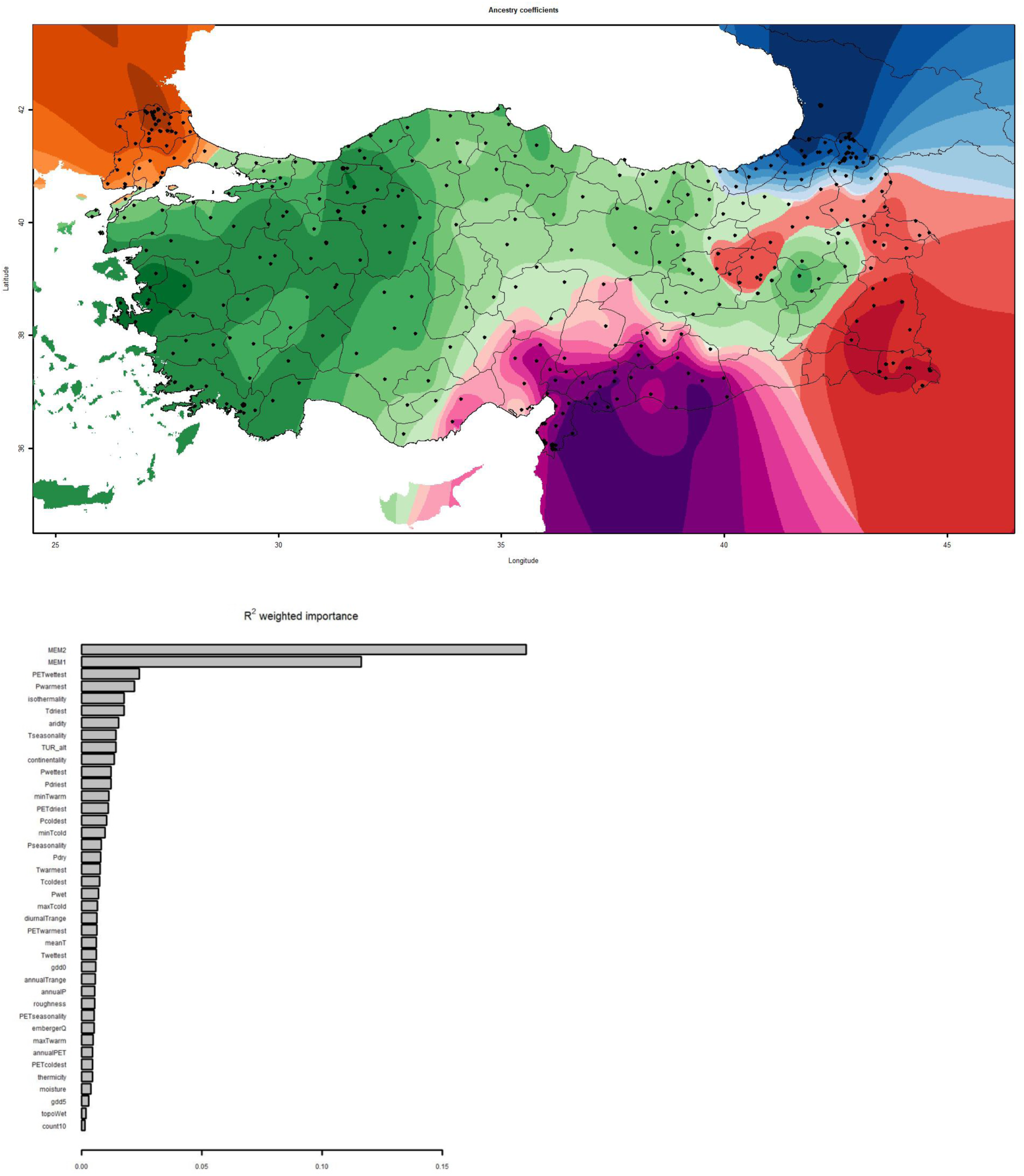

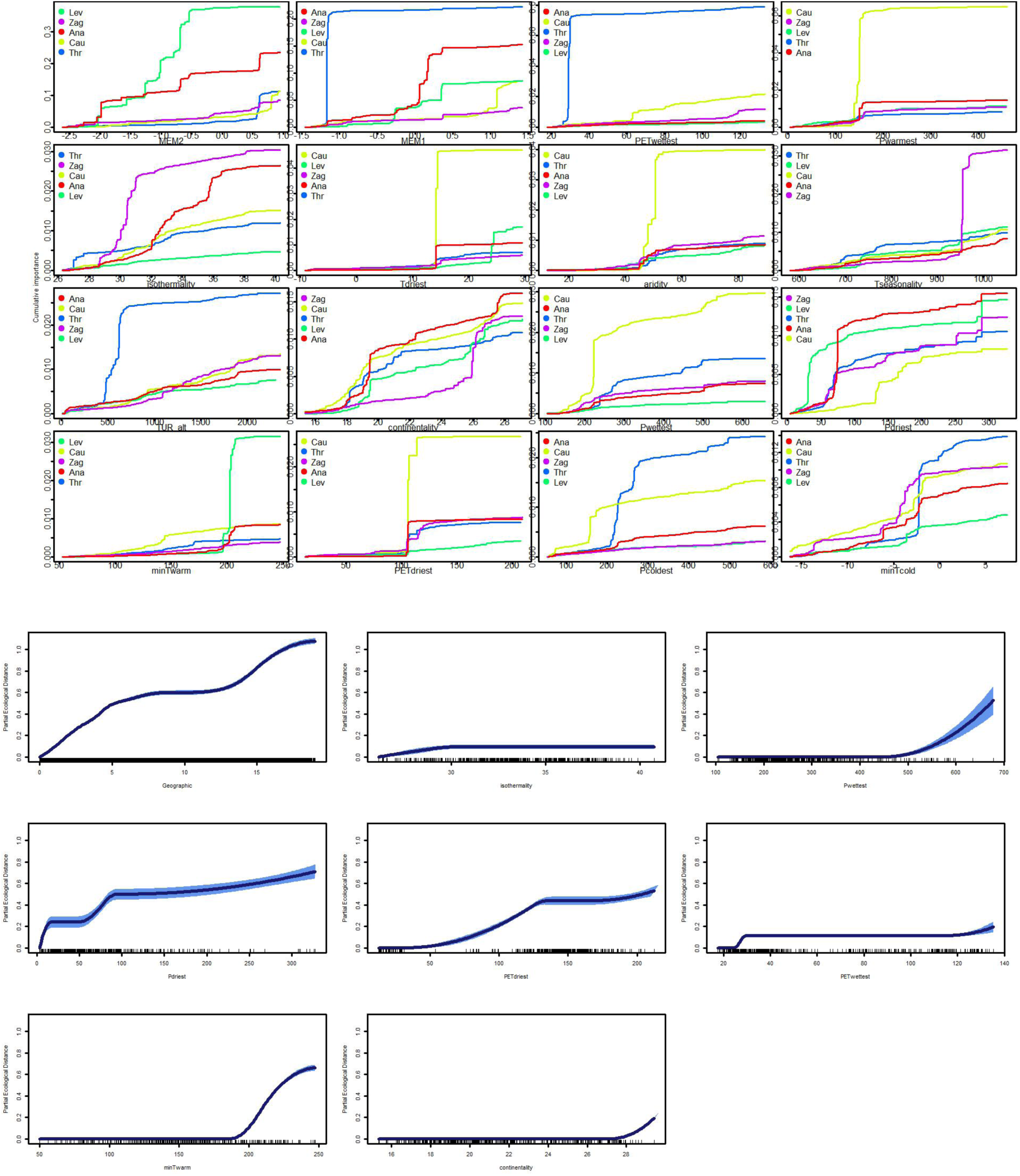

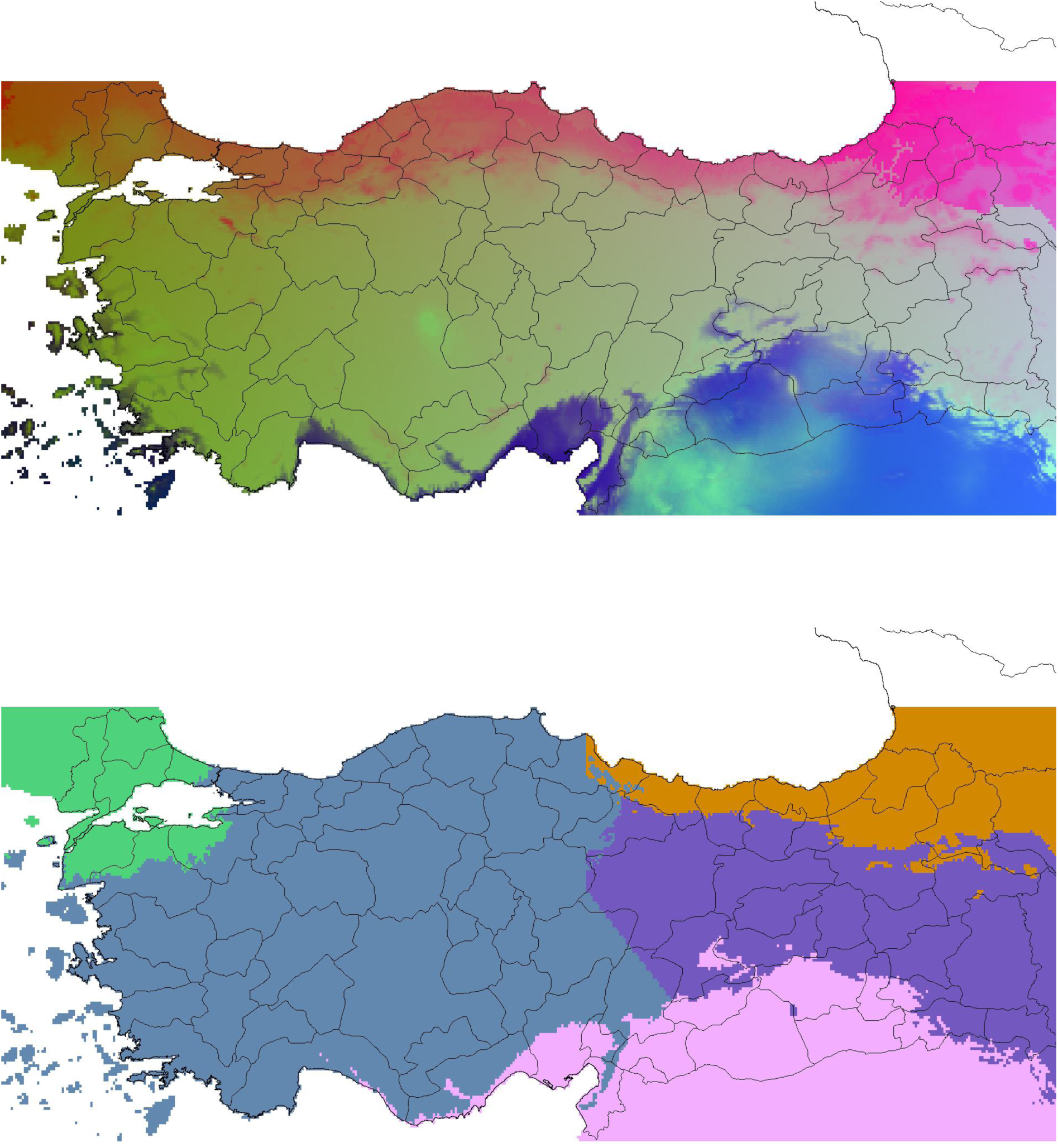

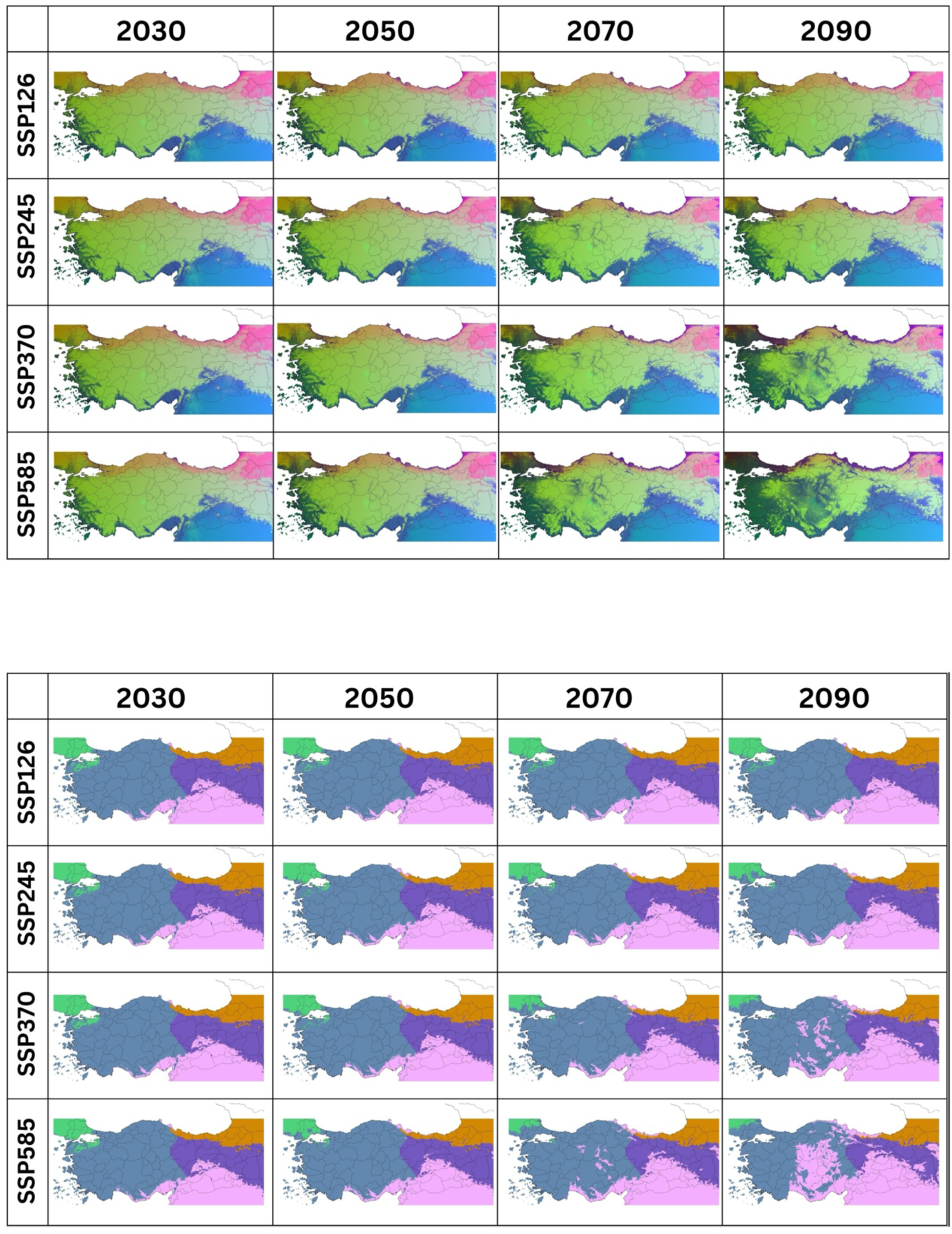

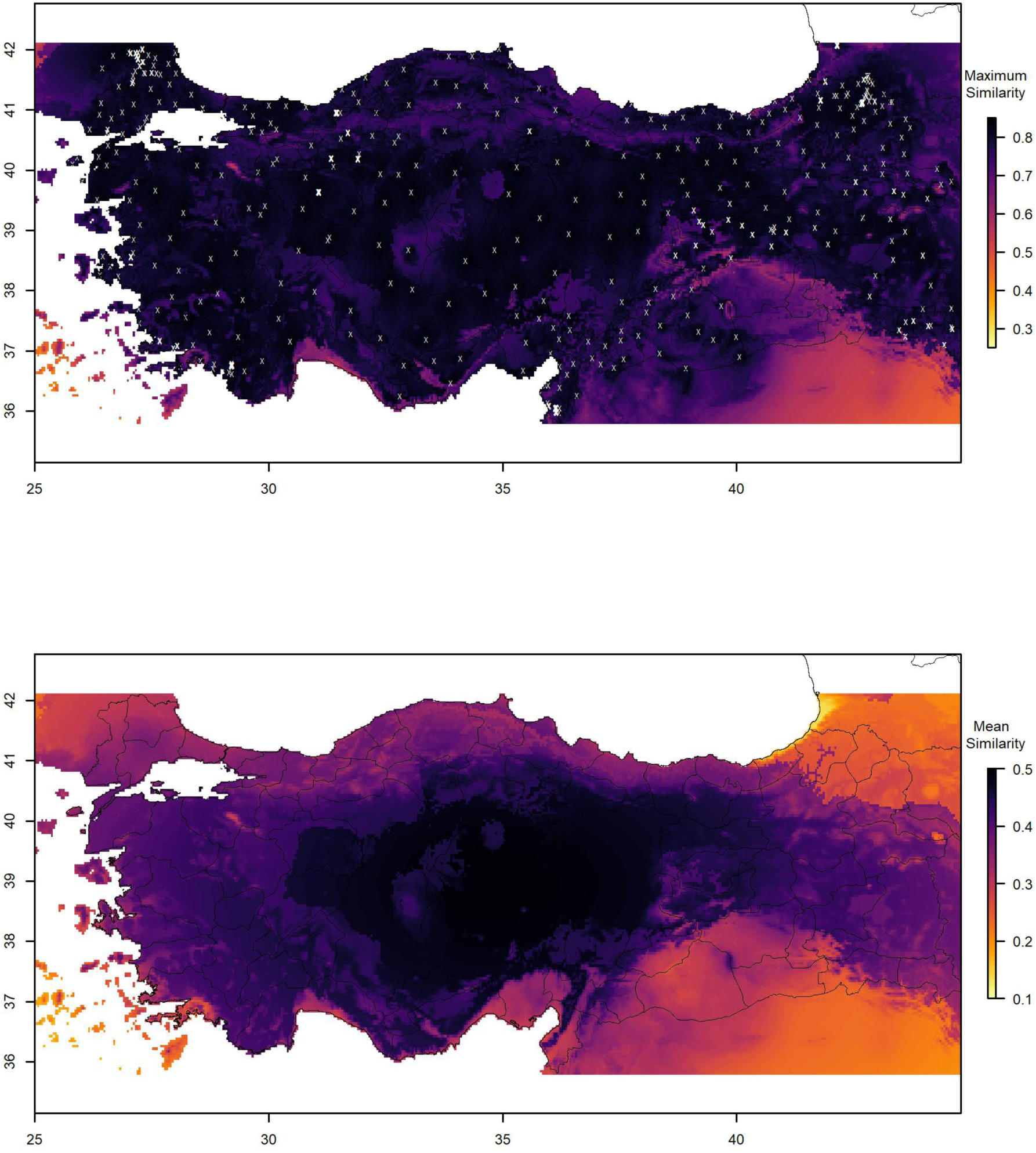

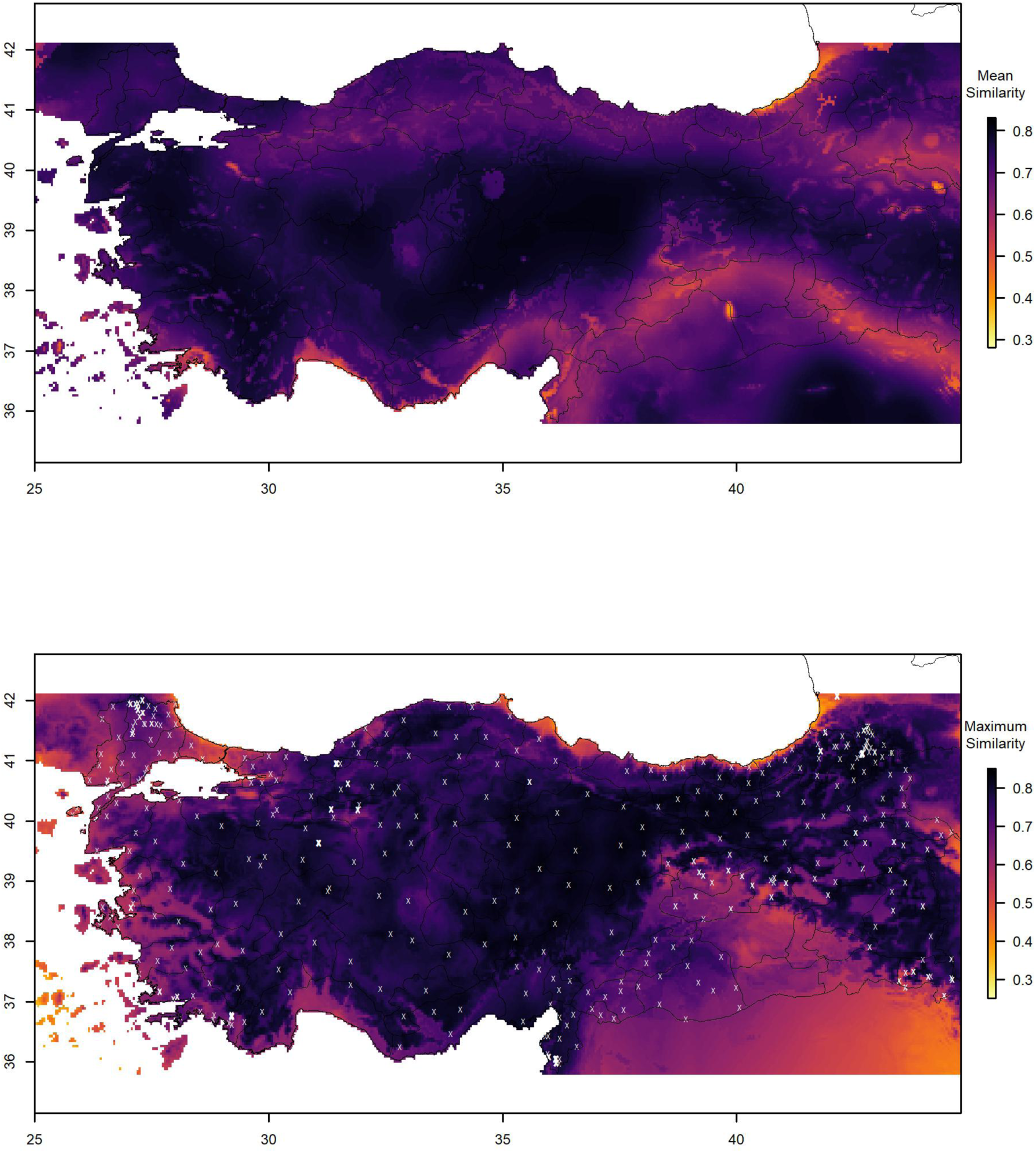

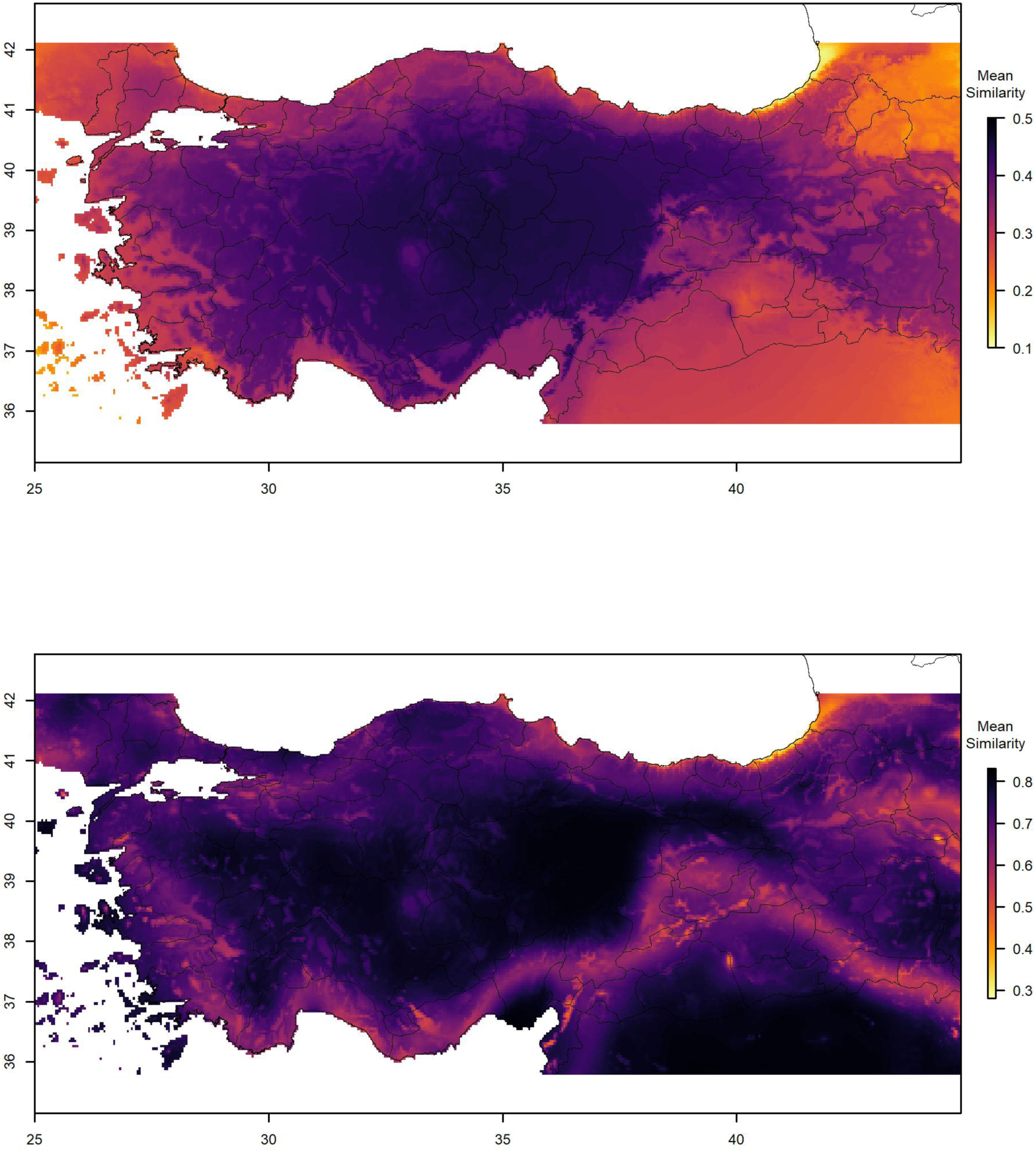

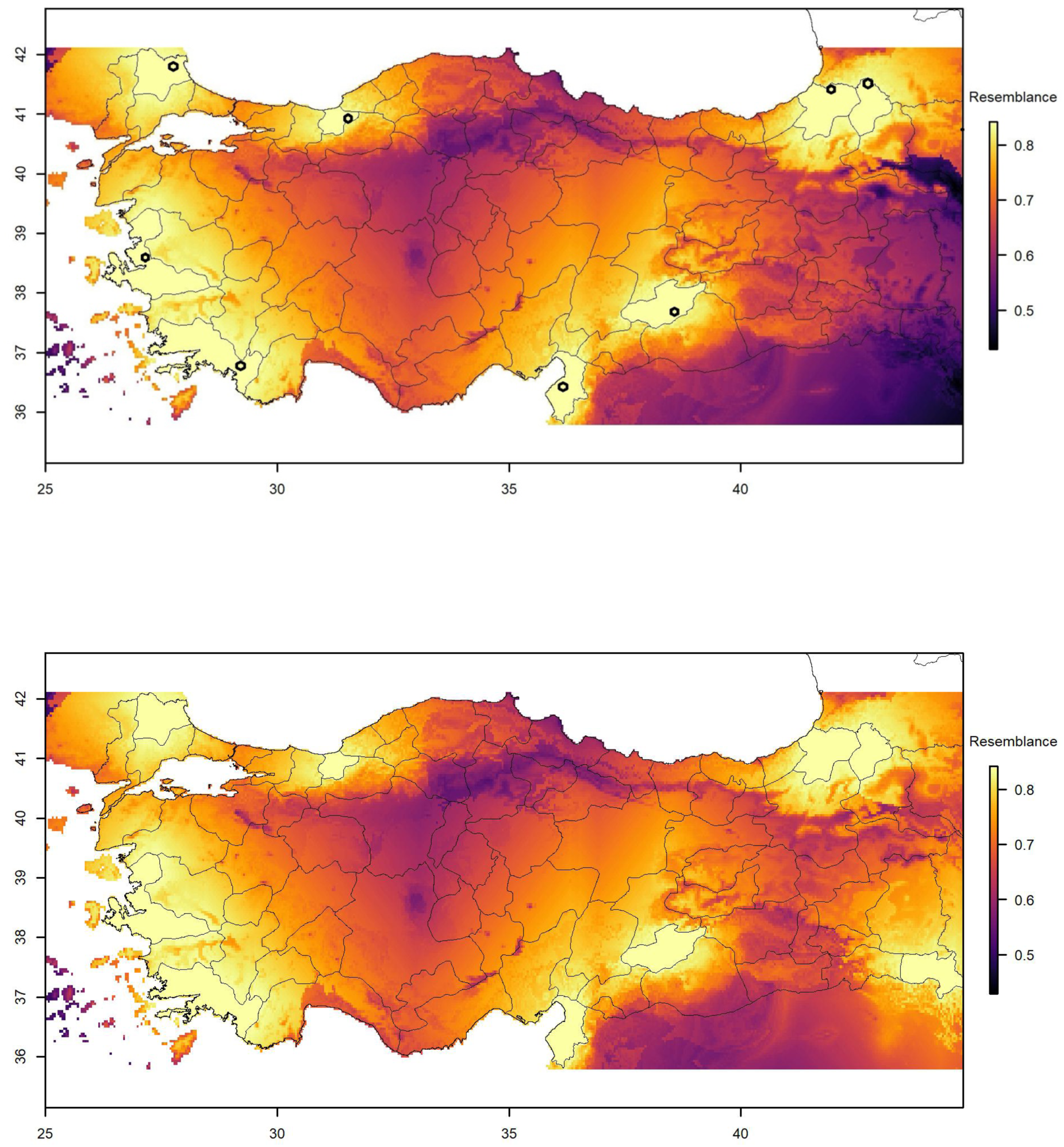

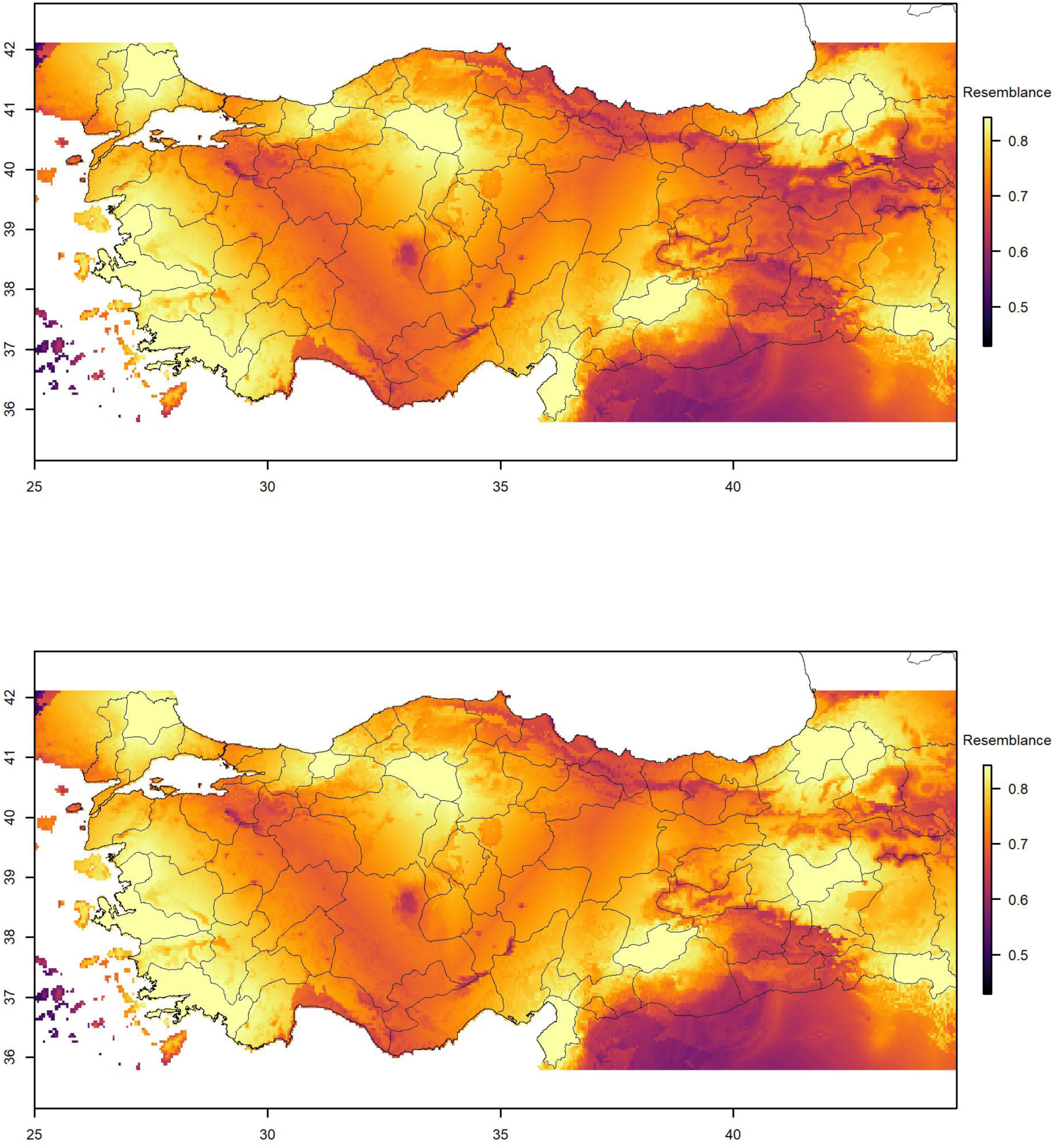

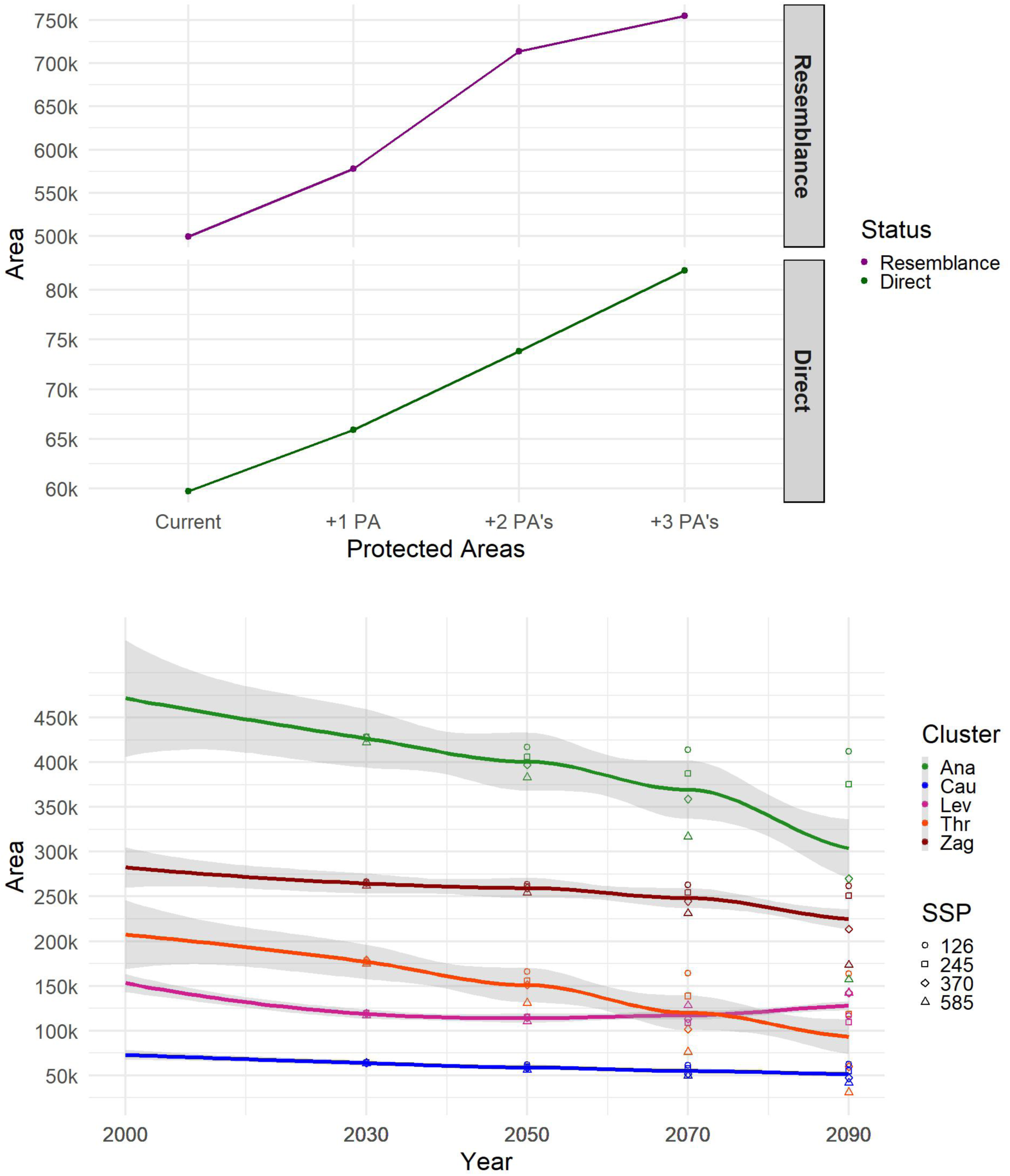

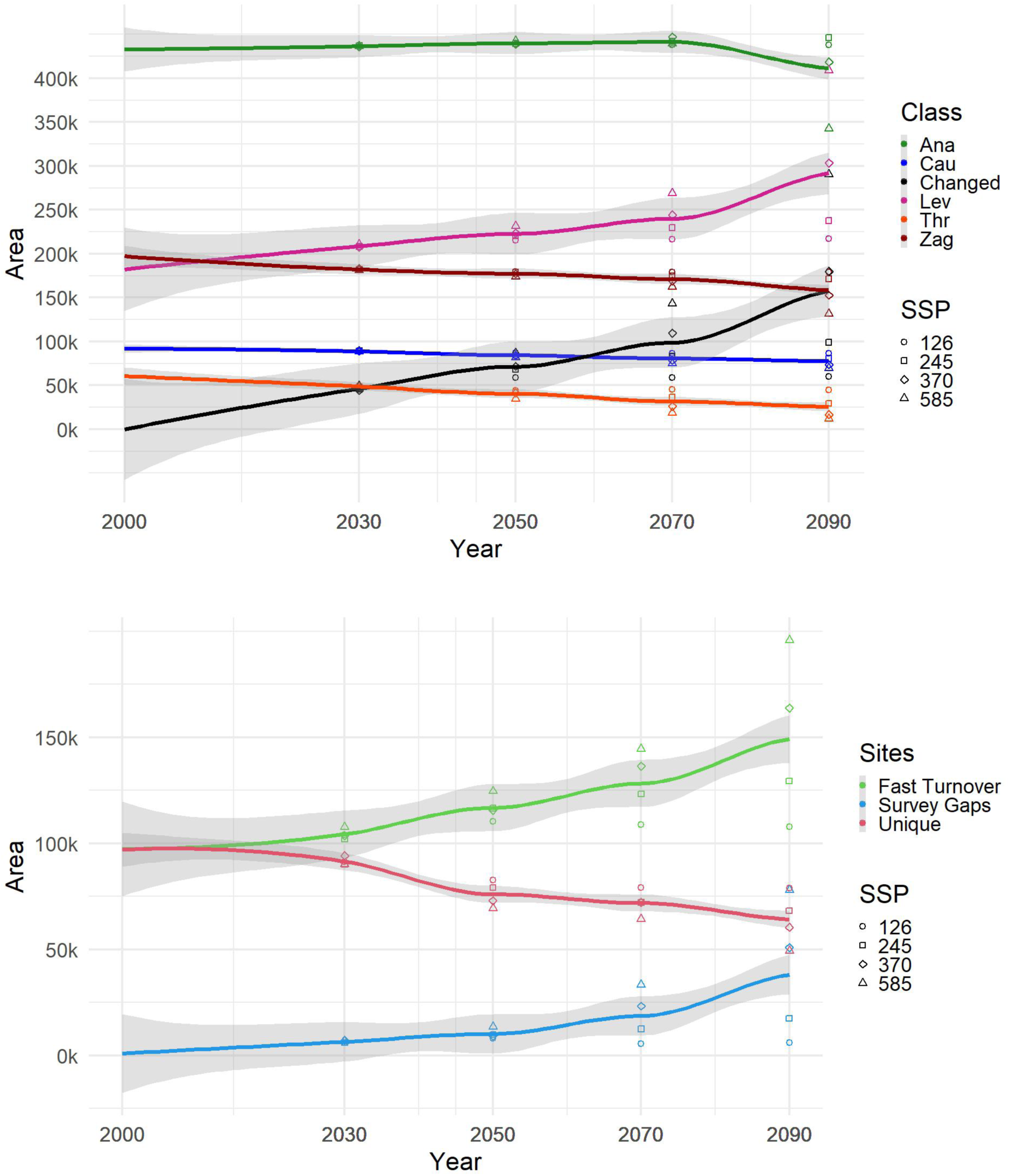

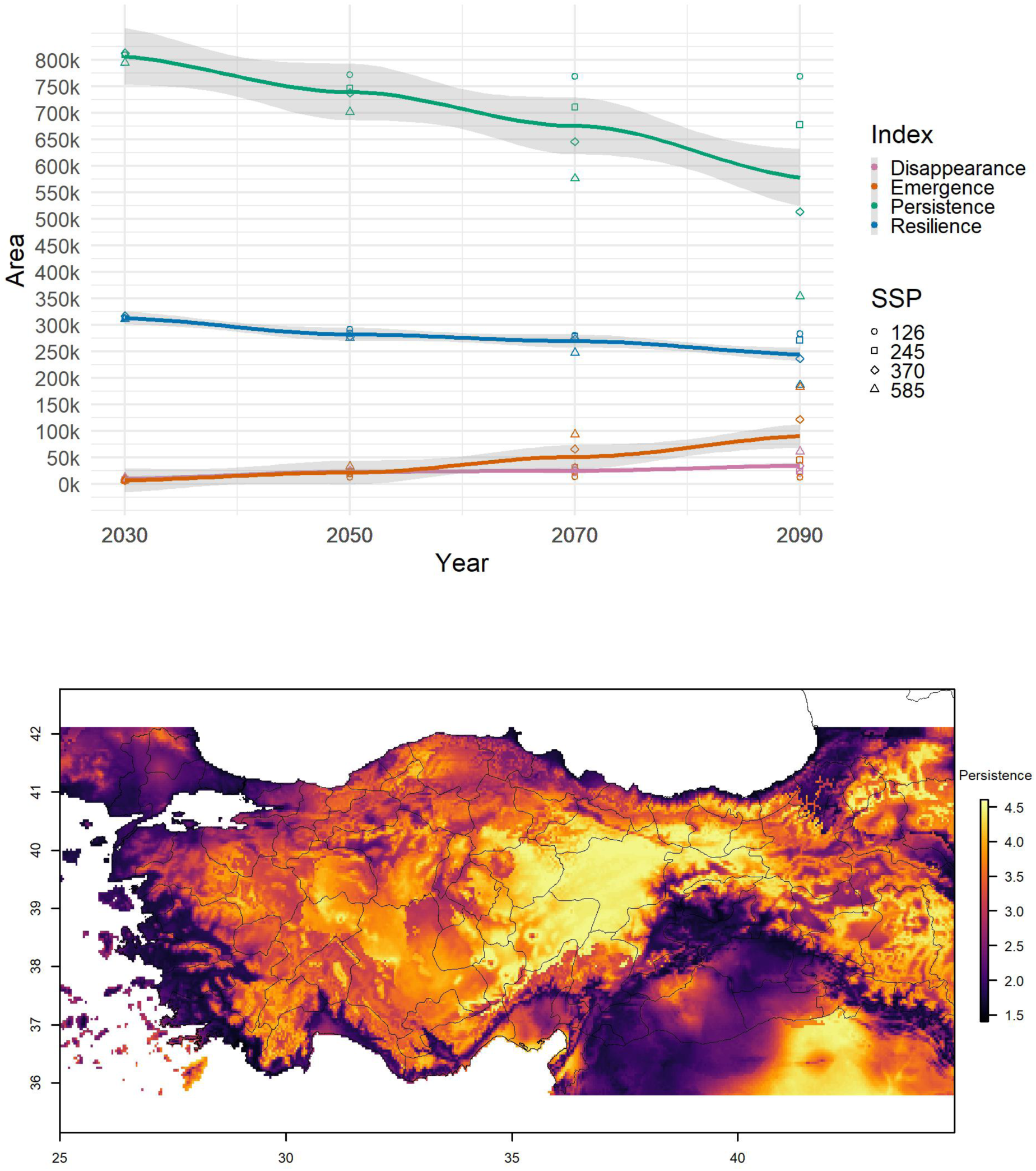

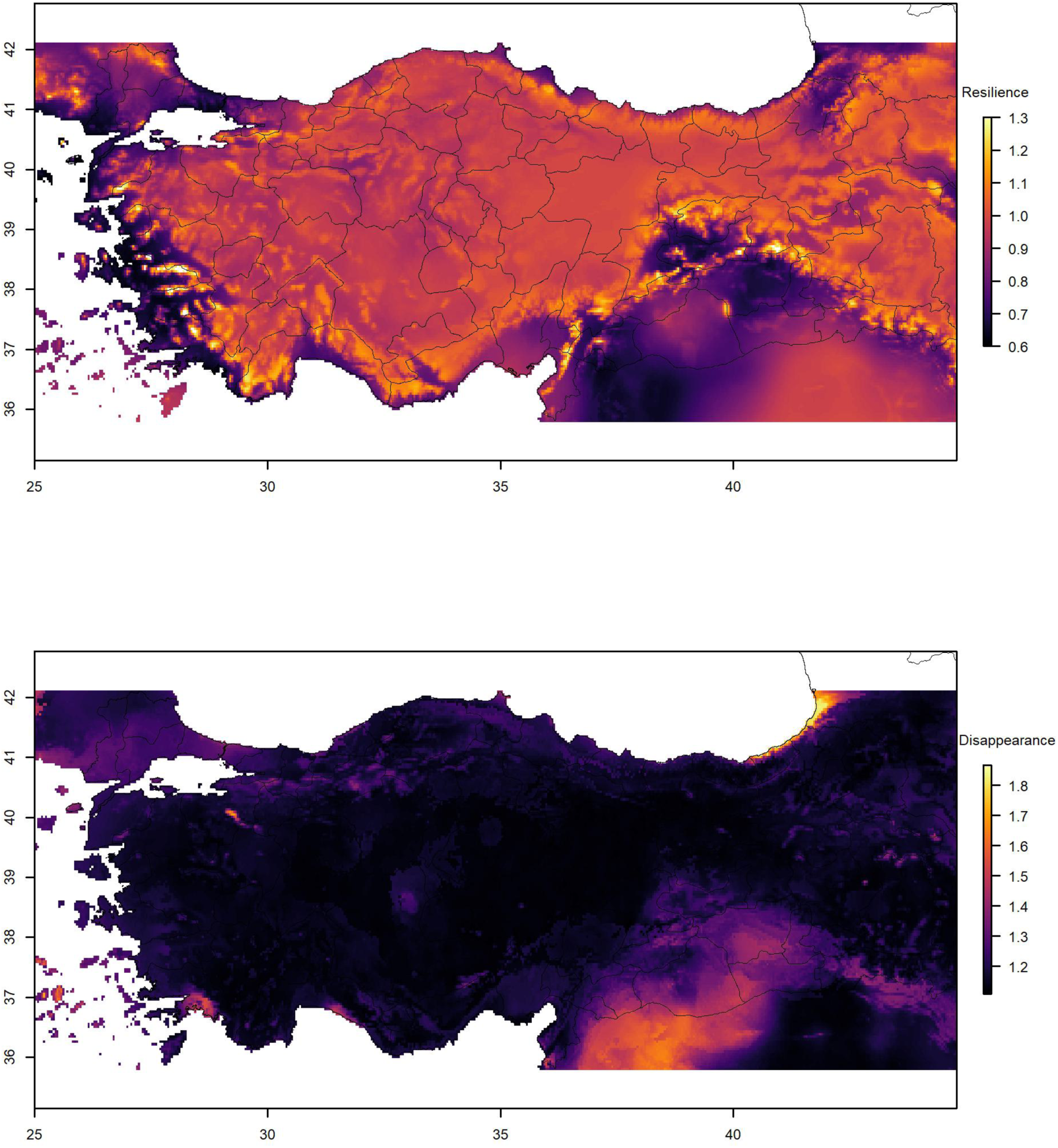

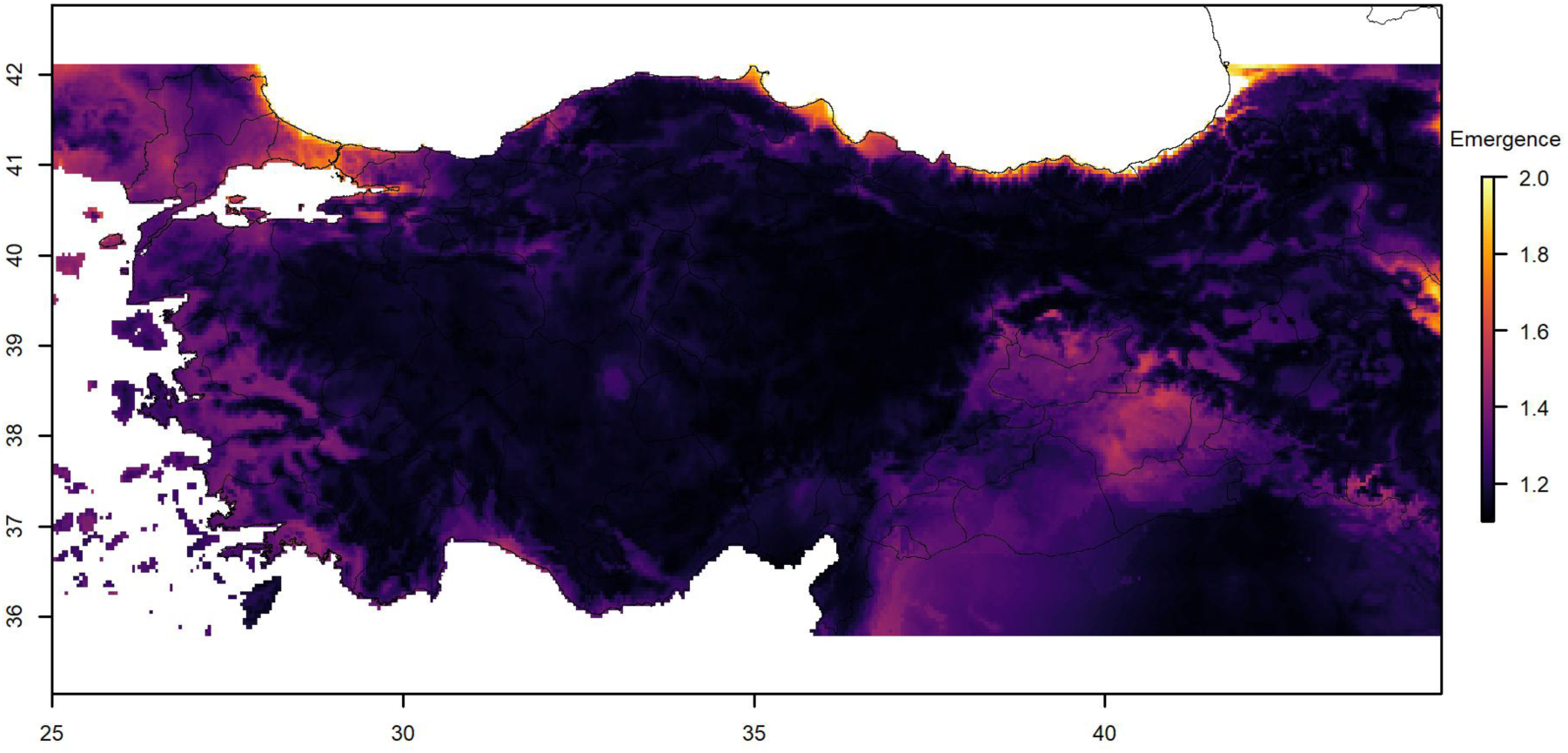

## REFERENCES

Abram, P. K., Boivin, G., Moiroux, J., & Brodeur, J. (2017). Behavioural effects of temperature on ectothermic animals: unifying thermal physiology and behavioural plasticity. Biological Reviews, 92(4), 1859–1876.

Akerman, A., & Bürger, R. (2014). The consequences of gene flow for local adaptation and differentiation: a two-locus two-deme model. Journal of mathematical biology, 68(5), 1135–1198.

Alattal, Y. Z., & Alghamdi, A. A. (2023). Linking histone methylation states and hsp transcriptional regulation in thermo-tolerant and thermo-susceptible A. mellifera L. subspecies in response to heat stress. Insects, 14(3), 225.

Alattal, Y., & AlGhamdi, A. (2015). Impact of temperature extremes on survival of indigenous and exotic honey bee subspecies, Apis mellifera, under desert and semiarid climates. Bulletin of Insectology, 68(2), 219–222.

Alghamdi, A. A., & Alattal, Y. Z. (2023). Expression levels of heat-shock proteins in Apis mellifera jemenetica and Apis mellifera carnica foragers in the desert climate of Saudi Arabia. Insects, 14(5), 432.

Alstad, A. O., Damschen, E. I., Givnish, T. J., Harrington, J. A., Leach, M. K., Rogers, D. A., & Waller, D. M. (2016). The pace of plant community change is accelerating in remnant prairies. Science Advances, 2(2), e1500975.

Arce-Valdés, L. R., & Sánchez-Guillén, R. A. (2022). The evolutionary outcomes of climate-change-induced hybridization in insect populations. Current Opinion in Insect Science, 54, 100966.

Auffret, A. G., Rico, Y., Bullock, J. M., Hooftman, D. A., Pakeman, R. J., Soons, M. B., … & Cousins, S. A. (2017). Plant functional connectivity–integrating landscape structure and effective dispersal. Journal of Ecology, 105(6), 1648–1656.

Babcock, R. C., Bustamante, R. H., Fulton, E. A., Fulton, D. J., Haywood, M. D., Hobday, A. J., … & Vanderklift, M. A. (2019). Severe continental-scale impacts of climate change are happening now: Extreme climate events impact marine habitat forming communities along 45% of Australia’s coast. Frontiers in Marine Science, 411.

Bay, R. A., Harrigan, R. J., Underwood, V. L., Gibbs, H. L., Smith, T. B., & Ruegg, K. (2018). Genomic signals of selection predict climate-driven population declines in a migratory bird. Science, 359(6371), 83–86.

Beckage, B., Gross, L. J., & Kauffman, S. (2011). The limits to prediction in ecological systems. Ecosphere, 2(11), 1–12.

Becsi, B., Formayer, H., & Brodschneider, R. (2021). A biophysical approach to assess weather impacts on honey bee colony winter mortality. Royal Society open science, 8(9), 210618.

Bellard, C., Bertelsmeier, C., Leadley, P., Thuiller, W., & Courchamp, F. (2012). Impacts of climate change on the future of biodiversity. Ecology letters, 15(4), 365–377.

Bellard, C., Cassey, P., & Blackburn, T. M. (2016). Alien species as a driver of recent extinctions. Biology letters, 12(2), 20150623.

Bilgin, R. (2011). Back to the suture: the distribution of intraspecific genetic diversity in and around Anatolia. International Journal of Molecular Sciences, 12(6), 4080–4103.

Bloch, G., Francoy, T. M., Wachtel, I., Panitz-Cohen, N., Fuchs, S., & Mazar, A. (2010). Industrial apiculture in the Jordan valley during Biblical times with Anatolian honeybees. Proceedings of the National Academy of Sciences, 107(25), 11240–11244.

Blois, J. L., Williams, J. W., Fitzpatrick, M. C., Jackson, S. T., & Ferrier, S. (2013). Space can substitute for time in predicting climate-change effects on biodiversity. Proceedings of the national academy of sciences, 110(23), 9374–9379.

Blüthgen, N., Staab, M., Achury, R., & Weisser, W. W. (2022). Unravelling insect declines: can space replace time?. Biology Letters, 18(4), 20210666.

Breed, M. F., Stead, M. G., Ottewell, K. M., Gardner, M. G., & Lowe, A. J. (2013). Which provenance and where? Seed sourcing strategies for revegetation in a changing environment. Conservation Genetics, 14, 1–10.

Brillet, C., Robinson, G. E., Bues, R., & Le Conte, Y. (2002). Racial differences in division of labor in colonies of the honey bee (Apis mellifera). Ethology, 108(2), 115–126.

Büchler, R., Costa, C., Hatjina, F., Andonov, S., Meixner, M. D., Conte, Y. L., … & Wilde, J. (2014). The influence of genetic origin and its interaction with environmental effects on the survival of Apis mellifera L. colonies in Europe. Journal of Apicultural Research, 53(2), 205–214.

Çakmak, I., Song, D. S., Mixson, T. A., Serrano, E., Clement, M. L., Savitski, A., … & Wells, H. (2010). Foraging response of Turkish honey bee subspecies to flower color choices and reward consistency. Journal of insect behavior, 23, 100–116.

Calfee, E., Agra, M. N., Palacio, M. A., Ramírez, S. R., & Coop, G. (2020). Selection and hybridization shaped the rapid spread of African honey bee ancestry in the Americas. PLoS genetics, 16(10), e1009038.

Cao, L., Dai, Z., Tan, H., Zheng, H., Wang, Y., Chen, J., … & Zhang, Z. (2023). Population Structure, Demographic History, and Adaptation of Giant Honeybees in China Revealed by Population Genomic Data. Genome Biology and Evolution, 15(3), evad025.

Chen, C., Liu, Z., Pan, Q., Chen, X., Wang, H., Guo, H., … & Shi, W. (2016). Genomic analyses reveal demographic history and temperate adaptation of the newly discovered honey bee subspecies Apis mellifera sinisxinyuan n. ssp. Molecular biology and evolution, 33(5), 1337–1348.

Chen, C., Wang, H., Liu, Z., Chen, X., Tang, J., Meng, F., & Shi, W. (2018). Population genomics provide insights into the evolution and adaptation of the eastern honey bee (Apis cerana). Molecular Biology and Evolution, 35(9), 2260–2271.

Chen, I. C., Hill, J. K., Shiu, H. J., Holloway, J. D., Benedick, S., Chey, V. K., … & Thomas, C. D. (2011). Asymmetric boundary shifts of tropical montane Lepidoptera over four decades of climate warming. Global Ecology and Biogeography, 20(1), 34–45.

Chen, Y., Gao, Y., Huang, X., Li, S., Zhang, Z., & Zhan, A. (2024). Incorporating adaptive genomic variation into predictive models for invasion risk assessment. Environmental Science and Ecotechnology, 18, 100299.

Christmas, M. J., Wallberg, A., Bunikis, I., Olsson, A., Wallerman, O., & Webster, M. T. (2019). Chromosomal inversions associated with environmental adaptation in honeybees. Molecular ecology, 28(6), 1358–1374.

Claudio, E. P., Rodriguez-Cruz, Y., Arslan, O. C., Giray, T., Rivera, J. L. A., Kence, M., … & Abramson, C. I. (2018). Appetitive reversal learning differences of two honey bee subspecies with different foraging behaviors. PeerJ, 6, e5918.

Cohn, A. S., Newton, P., Gil, J. D., Kuhl, L., Samberg, L., Ricciardi, V., … & Northrop, S. (2017). Smallholder agriculture and climate change. Annual Review of Environment and Resources, 42, 347–375.

Coreau, A., Pinay, G., Thompson, J. D., Cheptou, P. O., & Mermet, L. (2009). The rise of research on futures in ecology: rebalancing scenarios and predictions. Ecology letters, 12(12), 1277–1286.

Coronese, M., Lamperti, F., Keller, K., Chiaromonte, F., & Roventini, A. (2019). Evidence for sharp increase in the economic damages of extreme natural disasters. Proceedings of the National Academy of Sciences, 116(43), 21450–21455.

Costa, C., Lodesani, M., & Bienefeld, K. (2012). Differences in colony phenotypes across different origins and locations: evidence for genotype by environment interactions in the Italian honeybee (Apis mellifera ligustica)?. Apidologie, 43(6), 634–642.

Cridland, J. M., Ramirez, S. R., Dean, C. A., Sciligo, A., & Tsutsui, N. D. (2018). Genome sequencing of museum specimens reveals rapid changes in the genetic composition of honey bees in California. Genome biology and evolution, 10(2), 458–472.

Cridland, J. M., Tsutsui, N. D., & Ramírez, S. R. (2017). The complex demographic history and evolutionary origin of the western honey bee, Apis mellifera. Genome Biology and Evolution, 9(2), 457–472.

Cunningham, M. M., Tran, L., McKee, C. G., Polo, R. O., Newman, T., Lansing, L., … & Guarna, M. M. (2022). Honey bees as biomonitors of environmental contaminants, pathogens, and climate change. Ecological Indicators, 134, 108457.

Davis, M. B., & Shaw, R. G. (2001). Range shifts and adaptive responses to Quaternary climate change. Science, 292(5517), 673–679.

Dearden, P. K., Duncan, E. J., & Wilson, M. J. (2009). The honeybee Apis mellifera. Cold Spring Harbor Protocols, 2009(6), pdb-emo123.

Delgado, D. L., Pérez, M. E., Galindo-Cardona, A., Giray, T., & Restrepo, C. (2012). Forecasting the influence of climate change on agroecosystem services: potential impacts on honey yields in a small-island developing state. Psyche: A Journal of Entomology, 2012, 1–10.

DeMarche, M. L., Doak, D. F., & Morris, W. F. (2019). Incorporating local adaptation into forecasts of species’ distribution and abundance under climate change. Global Change Biology, 25(3), 775–793.

Doebeli, M., & Ispolatov, I. (2010). Complexity and diversity. Science, 328(5977), 494–497.

Dogantzis, Kathleen A., Tanushree Tiwari, Ida M. Conflitti, Alivia Dey, Harland M. Patch, Elliud M. Muli, Lionel Garnery et al. “Thrice out of Asia and the adaptive radiation of the western honey bee.” Science advances 7, no. 49 (2021): eabj2151.

Dudaniec, R. Y., Yong, C. J., Lancaster, L. T., Svensson, E. I., & Hansson, B. (2018). Signatures of local adaptation along environmental gradients in a range-expanding damselfly (Ischnura elegans). Molecular Ecology, 27(11), 2576–2593.

Dudenhöffer, J. H., Luecke, N. C., & Crawford, K. M. (2022). Changes in precipitation patterns can destabilize plant species coexistence via changes in plant–soil feedback. Nature Ecology & Evolution, 6(5), 546–554.

Dukes, J. S., & Mooney, H. A. (1999). Does global change increase the success of biological invaders?. Trends in ecology & evolution, 14(4), 135–139.

Eckert, C. G., Samis, K. E., & Lougheed, S. C. (2008). Genetic variation across species’ geographical ranges: the central–marginal hypothesis and beyond. Molecular ecology, 17(5), 1170–1188.

Ellis, N., Smith, S. J., & Pitcher, C. R. (2012). Gradient forests: calculating importance gradients on physical predictors. Ecology, 93(1), 156–168.

Everitt, T., Wallberg, A., Christmas, M. J., Olsson, A., Hoffmann, W., Neumann, P., & Webster, M. T. (2023). The genomic basis of adaptation to high elevations in Africanized honey bees. Genome biology and evolution, 15(9), evad157.

Exposito-Alonso, M., Vasseur, F., Ding, W., Wang, G., Burbano, H. A., & Weigel, D. (2018). Genomic basis and evolutionary potential for extreme drought adaptation in Arabidopsis thaliana. Nature ecology & evolution, 2(2), 352–358.

Fahrenholz, L., Lamprecht, I., & Schricker, B. (1989). Thermal investigations of a honey bee colony: thermoregulation of the hive during summer and winter and heat production of members of different bee castes. Journal of Comparative Physiology B, 159, 551–560.

Ferrier, S., Manion, G., Elith, J., & Richardson, K. (2007). Using generalized dissimilarity modelling to analyse and predict patterns of beta diversity in regional biodiversity assessment. Diversity and distributions, 13(3), 252–264.

Fick, S. E., & Hijmans, R. J. (2017). WorldClim 2: new 1-km spatial resolution climate surfaces for global land areas. International journal of climatology, 37(12), 4302–4315.

Fitzpatrick, M. C., & Keller, S. R. (2015). Ecological genomics meets community-level modelling of biodiversity: Mapping the genomic landscape of current and future environmental adaptation. Ecology letters, 18(1), 1–16.

Fitzpatrick, M. C., Chhatre, V. E., Soolanayakanahally, R. Y., & Keller, S. R. (2021). Experimental support for genomic prediction of climate maladaptation using the machine learning approach Gradient Forests. Molecular Ecology Resources, 21(8), 2749–2765.

Fitzpatrick, M. C., Sanders, N. J., Ferrier, S., Longino, J. T., Weiser, M. D., & Dunn, R. (2011). Forecasting the future of biodiversity: a test of single-and multi-species models for ants in North America. Ecography, 34(5), 836–847.

Fitzpatrick, M., Mokany, K., Manion, G., Nieto-Lugilde, D., Ferrier, S., Lisk, M., … & Fitzpatrick, M. M. (2022). Package ‘gdm’.

Flores, J. M., Gil-Lebrero, S., Gámiz, V., Rodríguez, M. I., Ortiz, M. A., & Quiles, F. J. (2019). Effect of the climate change on honey bee colonies in a temperate Mediterranean zone assessed through remote hive weight monitoring system in conjunction with exhaustive colonies assessment. Science of the Total Environment, 653, 1111–1119.

Franck, P., Garnery, L., Loiseau, A., Oldroyd, B. P., Hepburn, H. R., Solignac, M., & Cornuet, J. M. (2001). Genetic diversity of the honeybee in Africa: microsatellite and mitochondrial data. Heredity, 86(4), 420–430.

Franks, S. J., Weber, J. J., & Aitken, S. N. (2014). Evolutionary and plastic responses to climate change in terrestrial plant populations. Evolutionary applications, 7(1), 123–139.

Fuller, Z. L., Niño, E. L., Patch, H. M., Bedoya-Reina, O. C., Baumgarten, T., Muli, E., … & Miller, W. (2015). Genome-wide analysis of signatures of selection in populations of African honey bees (Apis mellifera) using new web-based tools. BMC genomics, 16(1), 1–18.

Geldmann, J., Manica, A., Burgess, N. D., Coad, L., & Balmford, A. (2019). A global-level assessment of the effectiveness of protected areas at resisting anthropogenic pressures. Proceedings of the National Academy of Sciences, 116(46), 23209–23215.

Gienapp, P., Teplitsky, C., Alho, J. S., Mills, J. A., & Merilä, J. (2008). Climate change and evolution: Disentangling environmental and genetic responses. Molecular Ecology, 17(1), 167–178.

Giordano, R., Galindo-Cardona, A., Melendez-Ackerman, E., Chen, S. C., & Giray, T. (2022). Adaptation of Invasive Species to Islands and the Puerto Rican Honey Bee. Frontiers in Ecology and Evolution, 10, 946737.

Gmel, A. I., Guichard, M., Dainat, B., Williams, G. R., Eynard, S., Vignal, A., … & Neuditschko, M. (2023). Identification of runs of homozygosity in Western honey bees (Apis mellifera) using whole-genome sequencing data. Ecology and Evolution, 13(1), e9723.

González-Tokman, D., Córdoba-Aguilar, A., Dáttilo, W., Lira-Noriega, A., Sánchez-Guillén, R. A., & Villalobos, F. (2020). Insect responses to heat: physiological mechanisms, evolution and ecological implications in a warming world. Biological Reviews, 95(3), 802–821.

Gonzalez, V. H., Oyen, K., Ávila, O., & Ospina, R. (2022). Thermal limits of Africanized honey bees are influenced by temperature ramping rate but not by other experimental conditions. Journal of Thermal Biology, 110, 103369.

Gordo, O., & Sanz, J. J. (2006). Temporal trends in phenology of the honey bee Apis mellifera (L.) and the small white Pieris rapae (L.) in the Iberian Peninsula (1952–2004). Ecological Entomology, 31(3), 261–268.

Gougherty, A. V., Keller, S. R., & Fitzpatrick, M. C. (2021). Maladaptation, migration and extirpation fuel climate change risk in a forest tree species. Nature Climate Change, 11(2), 166–171.

Grodzicki, P., & Caputa, M. (2014). Diurnal and seasonal changes in thermal preference of single, isolated bees and small groups of bees (Apis mellifera L.). Journal of insect behavior, 27, 701–711.

Grünzweig, J. M., De Boeck, H. J., Rey, A., Santos, M. J., Adam, O., Bahn, M., … & Yakir, D. (2022). Dryland mechanisms could widely control ecosystem functioning in a drier and warmer world. Nature ecology & evolution, 6(8), 1064–1076.

Gür, H. (2016). The Anatolian diagonal revisited: Testing the ecological basis of a biogeographic boundary. Zoology in the Middle East, 62(3), 189–199.

Haldane, J. B., & Spurway, H. (1954). A statistical analysis of communication in “Apis mellifera” and a comparison with communication in other animals. Insectes sociaux, 1, 247–283.

Halsch, C. A., Shapiro, A. M., Fordyce, J. A., Nice, C. C., Thorne, J. H., Waetjen, D. P., & Forister, M. L. (2021). Insects and recent climate change. Proceedings of the national academy of sciences, 118(2), e2002543117.

Hanson, J. O., Rhodes, J. R., Riginos, C., & Fuller, R. A. (2017). Environmental and geographic variables are effective surrogates for genetic variation in conservation planning. Proceedings of the National Academy of Sciences, 114(48), 12755–12760.

Harpur, B. A., Kent, C. F., Molodtsova, D., Lebon, J. M., Alqarni, A. S., Owayss, A. A., & Zayed, A. (2014). Population genomics of the honey bee reveals strong signatures of positive selection on worker traits. Proceedings of the National Academy of Sciences, 111(7), 2614–2619.

Harvey, J. A., Heinen, R., Gols, R., & Thakur, M. P. (2020). Climate change-mediated temperature extremes and insects: From outbreaks to breakdowns. Global change biology, 26(12), 6685–6701.

Hatjina, F., Costa, C., Büchler, R., Uzunov, A., Drazic, M., Filipi, J., … & Kezic, N. (2014). Population dynamics of European honey bee genotypes under different environmental conditions. Journal of Apicultural Research, 53(2), 233–247.

Henriques, D., Wallberg, A., Chávez-Galarza, J., Johnston, J. S., Webster, M. T., & Pinto, M. A. (2018). Whole genome SNP-associated signatures of local adaptation in honeybees of the Iberian Peninsula. Scientific Reports, 8(1), 11145.

Henry, R. J. (2016). Genomics strategies for germplasm characterization and the development of climate resilient crops. In Crop Breeding (pp. 25–34). Apple Academic Press.

Hewitt, G. M. (1999). Post-glacial re-colonization of European biota. Biological journal of the Linnean Society, 68(1-2), 87–112.

Hoffmann, I. (2010). Climate change and the characterization, breeding and conservation of animal genetic resources. Animal genetics, 41, 32–46.

Howden, S. M., Soussana, J. F., Tubiello, F. N., Chhetri, N., Dunlop, M., & Meinke, H. (2007). Adapting agriculture to climate change. Proceedings of the national academy of sciences, 104(50), 19691–19696.

Howlett, B. G., Butler, R. C., Nelson, W., & Donovan, B. J. (2013). Impact of climate change on crop pollinator in New Zealand. Wellington: Ministry for Primary Industries.

Iwasaki, J. M., & Hogendoorn, K. (2021). How protection of honey bees can help and hinder bee conservation. Current Opinion in Insect Science, 46, 112–118.

Jaffe, R., Dietemann, V., Allsopp, M. H., Costa, C., Crewe, R. M., Dall’Olio, R., … & Moritz, R. F. (2010). Estimating the density of honeybee colonies across their natural range to fill the gap in pollinator decline censuses. Conservation biology, 24(2), 583–593.

Jarvis A., H.I. Reuter, A. Nelson, E. Guevara, 2008, Hole-filled seamless SRTM data V4, International Centre for Tropical Agriculture (CIAT), available from https://srtm.csi.cgiar.org.

Jay, F., Manel, S., Alvarez, N., Durand, E. Y., Thuiller, W., Holderegger, R., … & François, O. (2012). Forecasting changes in population genetic structure of alpine plants in response to global warming. Molecular ecology, 21(10), 2354–2368.

Ji, Y., Li, X., Ji, T., Tang, J., Qiu, L., Hu, J., … & Zhou, X. (2020). Gene reuse facilitates rapid radiation and independent adaptation to diverse habitats in the Asian honeybee. Science Advances, 6(51), eabd3590.

Jones, M. R., Forester, B. R., Teufel, A. I., Adams, R. V., Anstett, D. N., Goodrich, B. A., … & Manel, S. (2013). Integrating landscape genomics and spatially explicit approaches to detect loci under selection in clinal populations. Evolution, 67(12), 3455–3468.

Joost, S., Bonin, A., Bruford, M. W., Després, L., Conord, C., Erhardt, G., … & Taberlet, P. (2007). A spatial analysis method (SAM) to detect candidate loci for selection: Towards a landscape genomics approach to adaptation. Molecular Ecology, 16(18), 3955–3969.

Kandemir, I., Kence, M., Sheppard, W. S., & Kence, A. (2006). Mitochondrial DNA variation in honey bee (Apis mellifera L.) populations from Turkey. Journal of apicultural research, 45(1), 33–38.

Kaya-Zeeb, S., Delac, S., Wolf, L., Marante, A. L., Scherf-Clavel, O., & Thamm, M. (2022). Robustness of the honeybee neuro-muscular octopaminergic system in the face of cold stress. Frontiers in physiology, 1979.

Keeler, A. M., Rose-Person, A., & Rafferty, N. E. (2021). From the ground up: Building predictions for how climate change will affect belowground mutualisms, floral traits, and bee behavior. Climate Change Ecology, 1, 100013.

Kekeçoğlu, M., Eroğlu, N., Kambur, M., & Münir, U. Ç. A. K. (2020). The relationships between propolis collecting capability and morphometric features of some honey bee races and ecotypes in Anatolia. Journal of Agricultural Sciences, 26(1), 71–77.

Kence, M., Oskay, D., Giray, T., & Kence, A. (2013). Honey bee colonies from different races show variation in defenses against the varroa mite in a ‘common garden’. Entomologia Experimentalis et Applicata, 149(1), 36–43.

Kim, S. W., Sommer, B., Beger, M., & Pandolfi, J. M. (2023). Regional and global climate risks for reef corals: Incorporating species-specific vulnerability and exposure to climate hazards. Global Change Biology.

Kinzner, M. C., Gamisch, A., Hoffmann, A. A., Seifert, B., Haider, M., Arthofer, W., … & Steiner, F. M. (2019). Major range loss predicted from lack of heat adaptability in an alpine Drosophila species. Science of the Total Environment, 695, 133753.

Knowles, L. L., & Massatti, R. (2017). Distributional shifts–not geographic isolation–as a probable driver of montane species divergence. Ecography, 40(12), 1475–1485.

Kovac, H., & Stabentheiner, A. (2011). Thermoregulation of foraging honeybees on flowering plants: seasonal variability and influence of radiative heat gain. Ecological entomology, 36(6), 686–699.

Kukkala, A. S., & Moilanen, A. (2013). Core concepts of spatial prioritisation in systematic conservation planning. Biological Reviews, 88(2), 443–464.

Kükrer, M. (2013). Genetic diversity of honey bee populations in Turkey based on microsatellite markers: A comparison between migratory versus stationary apiaries and isolated regions versus regions open to migratory beekeeping (Master’s thesis, Middle East Technical University).

Kükrer, M. (2024). From Diversity to Conservation: Insights for a National Genetic Monitoring Program for Honey Bees in the Face of Climate Change (Doctoral thesis, Middle East Technical University).

Kükrer, M., & Bilgin, C. C. (2020). Climate change prompts monitoring and systematic utilization of honey bee diversity in Turkey. Bee Studies, 12(1), 19–25.

Kükrer, M., Kence, M., & Kence, A. (2021). Honey bee diversity is swayed by migratory beekeeping and trade despite conservation practices: Genetic evidence for the impact of anthropogenic factors on population structure. Frontiers in Ecology and Evolution, 9, 556816.

Lancaster, L. T. (2016). Widespread range expansions shape latitudinal variation in insect thermal limits. Nature climate change, 6(6), 618–621.

Langowska, A., Zawilak, M., Sparks, T. H., Glazaczow, A., Tomkins, P. W., & Tryjanowski, P. (2017). Long-term effect of temperature on honey yield and honeybee phenology. International journal of biometeorology, 61, 1125–1132.

Lee, H., Calvin, K., Dasgupta, D., Krinner, G., Mukherji, A., Thorne, P., … & Park, Y. (2023). IPCC, 2023: Climate Change 2023: Synthesis Report, Summary for Policymakers. Contribution of Working Groups I, II and III to the Sixth Assessment Report of the Intergovernmental Panel on Climate Change [Core Writing Team, H. Lee and J. Romero (eds.)]. IPCC, Geneva, Switzerland.

Letunic, I., & Bork, P. (2021). Interactive Tree Of Life (iTOL) v5: an online tool for phylogenetic tree display and annotation. Nucleic acids research, 49(W1), W293–W296.

Lima-Rezende, C. A., Cabanne, G. S., Rocha, A. V., Carboni, M., Zink, R. M., & Caparroz, R. (2022). A comparative phylogenomic analysis of birds reveals heterogeneous differentiation processes among Neotropical savannas. Molecular Ecology, 31(12), 3451–3467.

Liu, Z., Fu, Y. H., Shi, X., Lock, T. R., Kallenbach, R. L., & Yuan, Z. (2022). Soil moisture determines the effects of climate warming on spring phenology in grasslands. Agricultural and Forest Meteorology, 323, 109039.

Lovell, R. S., Collins, S., Martin, S. H., Pigot, A. L., & Phillimore, A. B. (2023). Space-for-time substitutions in climate change ecology and evolution. Biological Reviews, 98(6), 2243–2270.

Marsh, J. I., Hu, H., Gill, M., Batley, J., & Edwards, D. (2021). Crop breeding for a changing climate: Integrating phenomics and genomics with bioinformatics. Theoretical and Applied Genetics, 134, 1677–1690.

Massatti, R., & Winkler, D. E. (2022). Spatially explicit management of genetic diversity using ancestry probability surfaces. Methods in Ecology and Evolution, 13(12), 2668–2681.

McAfee, A., Chapman, A., Higo, H., Underwood, R., Milone, J., Foster, L. J., … & Pettis, J. S. (2020). Vulnerability of honey bee queens to heat-induced loss of fertility. Nature Sustainability, 3(5), 367–376.

McCulloch, G. A., & Waters, J. M. (2023). Rapid adaptation in a fast-changing world: Emerging insights from insect genomics. Global Change Biology, 29(4), 943–954.

Meehl, G. A., Senior, C. A., Eyring, V., Flato, G., Lamarque, J. F., Stouffer, R. J., … & Schlund, M. (2020). Context for interpreting equilibrium climate sensitivity and transient climate response from the CMIP6 Earth system models. Science Advances, 6(26), eaba1981.

Meixner, M. D., Costa, C., Kryger, P., Hatjina, F., Bouga, M., Ivanova, E., & Büchler, R. (2010). Conserving diversity and vitality for honey bee breeding. Journal of Apicultural Research, 49(1), 85–92.

Meixner, M. D., Francis, R. M., Gajda, A., Kryger, P., Andonov, S., Uzunov, A., … & Wilde, J. (2014). Occurrence of parasites and pathogens in honey bee colonies used in a European genotype-environment interactions experiment. Journal of Apicultural Research, 53(2), 215–229.

Mokany, K., Bush, A., & Ferrier, S. (2019b). Community assembly processes restrict the capacity for genetic adaptation under climate change. Ecography, 42(6), 1164–1174.

Mokany, K., Harwood, T. D., Halse, S. A., & Ferrier, S. (2019a). Riddles in the dark: Assessing diversity patterns for cryptic subterranean fauna of the Pilbara. Diversity and Distributions, 25(2), 240–254.

Mokany, K., Ware, C., Woolley, S. N., Ferrier, S., & Fitzpatrick, M. C. (2022). A working guide to harnessing generalized dissimilarity modelling for biodiversity analysis and conservation assessment. Global Ecology and Biogeography, 31(4), 802–821.

Montero-Mendieta, S., Tan, K., Christmas, M. J., Olsson, A., Vilà, C., Wallberg, A., & Webster, M. T. (2019). The genomic basis of adaptation to high-altitude habitats in the eastern honey bee (Apis cerana). Molecular Ecology, 28(4), 746–760.

Morgan, K., Mboumba, J. F., Ntie, S., Mickala, P., Miller, C. A., Zhen, Y., … & Anthony, N. M. (2020). Precipitation and vegetation shape patterns of genomic and craniometric variation in the central African rodent Praomys misonnei. Proceedings of the Royal Society B, 287(1930), 20200449.

Mucci, C. A., Ramirez, L., Giffoni, R. S., & Lamattina, L. (2021). Cold stress induces specific antioxidant responses in honey bee brood. Apidologie, 52, 596–607.

Neupane, N., Larsen, E. A., & Ries, L. (2024). Ecological forecasts of insect range dynamics: a broad range of taxa include winners and losers under future climate. Current Opinion in Insect Science, 101159.

Nielsen, E. S., Henriques, R., Beger, M., & von der Heyden, S. (2021). Distinct interspecific and intraspecific vulnerability of coastal species to global change. Global Change Biology, 27(15), 3415–3431.

Niemczyk, M., Chmura, D. J., Socha, J., Wojda, T., Mroczek, P., Gil, W., & Thomas, B. R. (2021). How geographic and climatic factors affect the adaptation of Douglas-fir provenances to the temperate continental climate zone in Europe. European Journal of Forest Research, 140(6), 1341–1361.

Nürnberger, F., Härtel, S., & Steffan-Dewenter, I. (2019). Seasonal timing in honey bee colonies: phenology shifts affect honey stores and varroa infestation levels. Oecologia, 189, 1121–1131.

Oskay, D., Kükrer, M., & Kence, A. (2019). Muğla bal arısında (Apis mellifera anatoliaca) Amerikan yavru çürüklüğü hastalığına karşı direnç geliştirilmesi. Arıcılık Araştırma Dergisi, 11(1), 8–20.

Parejo, M., Wragg, D., Henriques, D., Charriere, J. D., & Estonba, A. (2020). Digging into the genomic past of Swiss honey bees by whole-genome sequencing museum specimens. Genome biology and evolution, 12(12), 2535–2551.

Parmesan, C. (2006). Ecological and evolutionary responses to recent climate change. Annual Review of Ecology, Evolution, and Systematics, 37, 637–669.

Pearman, P. B., Zachos, F. E., & Paz-Vinas, I. (2024). European monitoring of genetic diversity must expand to detect impacts of climate change. NATURE ECOLOGY & EVOLUTION.

Petchey, O. L., Pontarp, M., Massie, T. M., Kéfi, S., Ozgul, A., Weilenmann, M., … & Pearse, I. S. (2015). The ecological forecast horizon, and examples of its uses and determinants. Ecology letters, 18(7), 597–611.

Pörtner, H. O., Roberts, D. C., Poloczanska, E. S., Mintenbeck, K., Tignor, M., Alegría, A., … & Okem, A. (2022). IPCC, 2022: Summary for policymakers.

Pritchard, J. K., Stephens, M., & Donnelly, P. (2000). Inference of population structure using multilocus genotype data. Genetics, 155(2), 945–959.

Quigley, T. P., Amdam, G. V., & Harwood, G. H. (2019). Honey bees as bioindicators of changing global agricultural landscapes. Current opinion in insect science, 35, 132–137.

R Core Team (2022). R: A Language and Environment for Statistical Computing. R Foundation for Statistical Computing, Vienna, Austria. https://www.R-project.org/.

Reitalu, T., Bjune, A. E., Blaus, A., Giesecke, T., Helm, A., Matthias, I., … & Birks, H. J. B. (2019). Patterns of modern pollen and plant richness across northern Europe. Journal of Ecology, 107(4), 1662–1677.

Requier, F., Garnery, L., Kohl, P. L., Njovu, H. K., Pirk, C. W., Crewe, R. M., & Steffan-Dewenter, I. (2019). The conservation of native honey bees is crucial. Trends in ecology & evolution, 34(9), 789–798.

Reynolds, J., Weir, B. S., & Cockerham, C. C. (1983). Estimation of the coancestry coefficient: basis for a short-term genetic distance. Genetics, 105(3), 767–779.

Rodrigues, A. S., & Brooks, T. M. (2007). Shortcuts for biodiversity conservation planning: the effectiveness of surrogates. Annu. Rev. Ecol. Evol. Syst., 38, 713–737.

Román-Palacios, C., & Wiens, J. J. (2020). Recent responses to climate change reveal the drivers of species extinction and survival. Proceedings of the National Academy of Sciences, 117(8), 4211–4217.

Rowland, B. W., Rushton, S. P., Shirley, M. D., Brown, M. A., & Budge, G. E. (2021). Identifying the climatic drivers of honey bee disease in England and Wales. Scientific reports, 11(1), 21953.

Ruttner, F. (1988). Biogeography and Taxonomy of Honeybees. Springer-Verlag, Berlin Heidelberg.

Sarkar, S. (2006). Ecological diversity and biodiversity as concepts for conservation planning: comments on Ricotta. Acta Biotheoretica, 54, 133–140.

Shah, A. A., Dillon, M. E., Hotaling, S., & Woods, H. A. (2020). High elevation insect communities face shifting ecological and evolutionary landscapes. Current Opinion in Insect Science, 41, 1–6.

Sinervo, B., Mendez-De-La-Cruz, F., Miles, D. B., Heulin, B., Bastiaans, E., Villagrán-Santa Cruz, M., … & Sites Jr, J. W. (2010). Erosion of lizard diversity by climate change and altered thermal niches. Science, 328(5980), 894–899.

Sönmez, S. (2022). Assessing the potential of Anatolia as a climate refugium (Master’s thesis, Istanbul Technical University).

Stabentheiner, A., Nagy, J. M., Kovac, H., Käfer, H., Petrocelli, I., & Turillazzi, S. (2022). Effect of climate on strategies of nest and body temperature regulation in paper wasps, Polistes biglumis and Polistes gallicus. Scientific reports, 12(1), 3372.

Thuiller, W., Guéguen, M., Renaud, J., Karger, D. N., & Zimmermann, N. E. (2019). Uncertainty in ensembles of global biodiversity scenarios. Nature Communications, 10(1), 1446.

Title, P. O., & Bemmels, J. B. (2018). ENVIREM: an expanded set of bioclimatic and topographic variables increases flexibility and improves performance of ecological niche modeling. Ecography, 41(2), 291–307.

Tokarska, K. B., Stolpe, M. B., Sippel, S., Fischer, E. M., Smith, C. J., Lehner, F., & Knutti, R. (2020). Past warming trend constrains future warming in CMIP6 models. Science advances, 6(12), eaaz9549.

Torson, A. S., Yocum, G. D., Rinehart, J. P., Kemp, W. P., & Bowsher, J. H. (2015). Transcriptional responses to fluctuating thermal regimes underpinning differences in survival in the solitary bee Megachile rotundata. The Journal of Experimental Biology, 218(7), 1060–1068.

Uzunov, A., Costa, C., Panasiuk, B., Meixner, M., Kryger, P., Hatjina, F., … & Büchler, R. (2014). Swarming, defensive and hygienic behaviour in honey bee colonies of different genetic origin in a pan-European experiment. Journal of Apicultural Research, 53(2), 248–260.

Vanhove, M., Pina-Martins, F., Coelho, A. C., Branquinho, C., Costa, A., Batista, D., … & Paulo, O. S. (2021). Using gradient Forest to predict climate response and adaptation in Cork oak. Journal of Evolutionary Biology, 34(6), 910–923.

Visscher, P. K., & Seeley, T. D. (1982). Foraging strategy of honeybee colonies in a temperate deciduous forest. Ecology, 63(6), 1790–1801.

Wallberg, A., Han, F., Wellhagen, G., Dahle, B., Kawata, M., Haddad, N., … & Webster, M. T. (2014). A worldwide survey of genome sequence variation provides insight into the evolutionary history of the honeybee Apis mellifera. Nature Genetics, 46(10), 1081–1088.

Wallberg, A., Schoening, C., Webster, M. T., & Hasselmann, M. (2017). Two extended haplotype blocks are associated with adaptation to high altitude habitats in East African honey bees. PLoS Genetics, 13(5), e1006792.

Wang, Q., Xu, X., Zhu, X., Chen, L., Zhou, S., Huang, Z. Y., & Zhou, B. (2016). Low-temperature stress during capped brood stage increases pupal mortality, misorientation and adult mortality in honey bees. PloS one, 11(5), e0154547.

Willis, K. J., & Bhagwat, S. A. (2009). Biodiversity and climate change. Science, 326(5954), 806–807.

Willmer, P. G., & Stone, G. N. (2004). Behavioral, ecological, and physiological determinants of the activity patterns of bees. Advances in the Study of Behavior, 34(34), 347–466.

Wragg, D., Techer, M. A., Canale-Tabet, K., Basso, B., Bidanel, J. P., Labarthe, E., … & Vignal, A. (2018). Autosomal and mitochondrial adaptation following admixture: a case study on the honeybees of Reunion Island. Genome Biology and Evolution, 10(1), 220–238.

Yıldız, B. I., & Karabağ, K. (2022). Quantitation of neuroxin-1, ataxin-3 and atlastin genes related to grooming behavior in five races of honey bee, Apis mellifera L., 1758 (Hymenoptera: Apidae), in Turkey. Turkish Journal of Entomology, 46(1), 3–11.

Zapata-Hernández, G., Gajardo-Rojas, M., Calderón-Seguel, M., Muñoz, A. A., Yáñez, K. P., Requier, F., … & Arrieta, H. (2024). Advances and knowledge gaps on climate change impacts on honey bees and beekeeping: A systematic review. Global Change Biology, 30(3), e17219.

Zhang, M., Keenan, T. F., Luo, X., Serra-Diaz, J. M., Li, W., King, T., … & Gong, P. (2022). Elevated CO2 moderates the impact of climate change on future bamboo distribution in Madagascar. Science of The Total Environment, 810, 152235.

Zhu, J., Zhang, Y., & Wang, W. (2016). Interactions between warming and soil moisture increase overlap in reproductive phenology among species in an alpine meadow. Biology Letters, 12(7), 20150749.

