## Supplementary Text for "Spatio-temporal shifts driven by climate change threaten persistence and resilience of honey bee populations"

### INTRODUCTION

Divergent local geographic forms often arise from spatially heterogeneous selection acting on complex alternative phenotypes. These phenotypes are influenced by the interplay between the external environment, genes, and their immediate genomic environment (Chevin & Hospital, 2008; Slatkin, 2009; Schwander et al., 2014). These traits are subjects of polygenic local adaptation and highly interconnected regulatory networks (Pritchard et al., 2010; Jones et al., 2012; Boyle et al., 2017). Correspondingly, selection acts upon multiple end phenotypes of whole organisms rather than single traits, where harmoniously integrated gene complexes compromise between opposing selection pressures (Mayr, 1970). When outbreeding disrupts these coadapted gene complexes, locally adapted ecotypes can persist in parapatry despite gene flow (Lynch, 1991; Flaxman et al., 2013; Akerman & Bürger, 2014; Kulmuni & Westram, 2017). The geographic structure observed in such cases can be attributed to the interplay between the gene flow (migration) levels and selection pressure (local adaptation; Galindo et al., 2009; North et al., 2011; Kirk & Freeland, 2011).

Selective sweeps and background selection involved in local adaptation enhance differentiation signals even at distant neutral loci (Charlesworth et al., 1997). Moreover, pronounced differentiation at neutral loci in locally adapted populations is particularly evident under high levels of inbreeding (Eckert et al., 2010). These signals are further amplified when selection acts on many loci, making it difficult for alleles to disentangle from non-neutral loci (Barton & Bengtsson, 1986). Thus, in the presence of local adaptation, the overall genetic structure may serve as a proxy for the coupling between environment and organism, even when inferred from neutral markers. In six tree species, randomly selected SNPs outperformed candidate loci in predicting performance in common garden experiments (Fitzpatrick et al., 2021; Capblancq & Forester, 2021; Lachmuth et al., 2023; Lind et al.,

2023). In *Arabidopsis thaliana*, no individual SNP was significantly associated with highly heritable drought survival; instead, it was linked to genetic group membership (Exposito-Alonso et al., 2018). Similarly, Nielsen et al. (2021) found that both neutral and outlier loci followed biogeographical breaks in the Cape urchin, common shore crab, and granular limpet. These findings suggest that allele frequencies across the genome generally align with environmental gradients crucial for local adaptation, influencing both adaptive and neutral genomic backgrounds in parallel or proportionally. Although widely considered a confounding element in adaptive marker identification (Yu et al., 2006), population structure may have adaptive significance and inform about dynamic responses to environmental change.

#### MATERIALS AND METHODS

We grouped samples in and around Türkiye into 12 putative populations based on their proximity and similarities in climate, topography, and floral characteristics, alongside initial findings from prior research (Kandemir et al., 2006; Kükrer et al., 2021). Beekeepers in this study confirmed that they used honey bees from native stocks and had not purchased non-native queens or colonies in the last ten years. We isolated DNA from bee heads, grouped 30 microsatellite loci (Estoup et al., 1995; Solignac et al., 2003; Bodur et al., 2007; Shaibi et al., 2008; Tunca, 2009) into four sets (set 1: AP218, A113, AB024, AP249, A088, AP001, AP043; set 2: AP049, AP238, AC006, AP243, AP288, HBC1602, A107; set 3: A079, AC306, AP226, A007, HBC1601, AP068, A014, AP223; set 4: AP019, AB124, A043, A076, AP273, AP289, HBC1605, A028). We excluded locus A076 since it did not consistently amplify across samples. To address potential errors in genotyping, we blindly double-genotyped 18,310 alleles in 290 individuals to estimate errors associated with the genotyping process. We considered any mismatch between separate genotyping efforts as an error and corrected them by checking the raw electropherogram data. We removed eight samples genotyped at fewer than eight loci and a duplicated individual detected by the R-package poppr 2.9.4 (R Core Team 2022 version 4.2.2; Kamvar et al., 2014). To find full/half-sibs, we calculated relatedness by Colony 2.0 (Wang, 2012) and removed ten individuals assigned as siblings.

We estimated null allele frequencies by popgenreport 3.0.7 (Adamack & Gruber, 2014). We used the same package to calculate the total number of alleles per locus, observed and expected heterozygosity, deviations from Hardy-Weinberg equilibrium, linkage disequilibrium, the Shannon-Wiener diversity index, and the number of unique private alleles for each putative population. We estimated allelic richness and calculated the genetic fixation index ($F_{st}$), inbreeding coefficient ($F_{is}$), and allelic differentiation by hierfstat 0.5-11 (Goudet, 2005). We tested whether genetic differences within and between populations differ from random expectations by AMOVA implemented in poppr. Before running a model-based clustering algorithm, we examined population structure by analyzing principal components through Discriminant Analysis of Principal Components (DAPC; Jombart et al., 2010), regular Principal Component Analysis (PCA), and a spatially explicit version of it: sPCA (Jombart et al., 2008). The sPCA method produces independent synthetic variables that maximize genetic variance and spatial autocorrelation (implemented in adegenet 2.1.10; Jombart, 2008). We decided to conduct downstream analyses with five ancestral groups based on population structure revealed through Fst values, phylogenetic tree, AMOVA results, and sPCA. We estimated individual membership coefficients by Structure 2.3.4 (Pritchard et al., 2000), with the provided run parameters in **Supplementary Table 9**, analyzed distinct K-values (**Supplementary Figure 1**) and permuted ancestry estimates by Clumpak (Kopelman et al., 2015), and visualized them with dabestr 0.3.0 (Ho et al., 2019).

The Structure model assumes a hypothetical ancestral population by allowing each population to drift away from that ancestral population at a different rate and considers that allele frequencies tend to be similar in various populations (Falush et al., 2003). Correspondingly, the model allows the existence of multiple populations with closely matched allele frequencies to capture subtle genetic structure. However, this broad definition of what constitutes a population may lead to the inference of spurious populations not stemming from genetic discontinuity, especially for those that exhibit wide geographical distribution under a stepping-stone migration model (Falush et al., 2003; Guillot et al., 2005). Such populations might erroneously persist due to the failure of the MCMC algorithm to eliminate them effectively, indicating a potential convergence issue (Guillot et al., 2005). The minimum K-value where all the subspecies became identifiable was seven, showcasing a spurious cluster that cannot be clearly traced to any particular geography but still stays within the distribution of *anatoliaca*. The wide distribution of the subspecies is quite heterogeneous regarding environmental conditions and involves transitions to multiple other subspecies. Hence, we summed membership coefficients from these two *anatoliaca* clusters for further analysis. Also, we combined residual *carnica* and Thracian clusters' membership coefficients to account for the total C lineage ancestry found in the samples. This summation across clusters left us with five ancestry estimates for each individual, i.e., Levantine, Caucasian, Anatolian, Zagrosian, and Thracian ancestral groups.

We identified spatial outliers for each of the five ancestral groups with spdep 1.2-8 (Bivand & Wong, 2018) by plotting ancestry estimates of each individual against their spatially lagged values within a mean radius of ~80 km. We manually checked those outliers and removed 80 samples with obvious mismatches to spatially expected ancestry. Elimination included unexpected cases such as unadmixed (i.e., with an estimated ancestry larger than 0.75) Caucasian individuals on the Aegean coast or unadmixed Thracian individuals in East Anatolia (see **Supplementary Figures 2, 3, 4, 5, and 6**). We retrieved and analyzed spatial data using raster 3.6-23 (Hijmans, 2022), downloaded future bioclimatic variables with geodata 0.5-3 (Hijmans et al., 2023), and calculated Moran's Eigenvector Maps (MEMs) with adespatial 0.3-21 (Dray et al., 2023). To eliminate multicollinearity in predictor variables, we applied an elimination procedure based on the variance inflation factor (VIF) using usdm 2.1-6 (Naimi et al., 2014).

#### RESULTS

Double-genotyping efforts revealed a low overall error rate of 2.7%. We found 499 mismatches out of 18,310 microsatellite alleles genotyped and estimated initial per locus genotyping error rates between 0 and 0.07 (**Supplementary Table 10**). Since we rechecked the raw electropherograms of all the mismatches, the remaining genotyping error would be even lower. The number of alleles per locus ranged from 4 to 53, and the total number of alleles was 574, of which 140 were private alleles. The mean estimated frequency of null alleles was 0.05, with a maximum value of 0.12 (**Supplementary Table 10**). Four loci pairs showed significant linkage disequilibrium in three populations (**Supplementary Table 11**), and there were 46 population-loci pairs out of 377 with significant deviations from Hardy-Weinberg equilibrium (**Supplementary Figure 7**). No loci showed widespread linkage disequilibrium, deviation from Hardy-Weinberg equilibrium, and high null allele frequencies simultaneously, so we kept all the loci for the rest of the analysis. We present the several diversity indices calculated in **Supplementary Table 10**. The rest of these loci-based statistics include expected and observed heterozygosity levels, $F_{is}$, $F_{it}$, and $F_{st}$ estimations, evenness, $G'_{st}$, $G_{st}$, and Jost's D estimations.

At the population level, mean allelic richness per loci corrected for sample sizes ranged between 1.42 and 1.63, with a mean of 1.56, Thrace having the highest allelic richness observed. Europe, Thrace, and Eastern Mediterranean had the highest number of private alleles: 28, 25, and 23. Eastern Mediterranean and Thrace also showed the highest richness values at 4.38 and 4.34 in a Shannon-Wiener diversity index (**Supplementary Table 12**). Erzurum-Kars Volcanic Plateau and Lesser Caucasus populations had the highest inbreeding levels with $F_{is}$ estimations of 0.15 and 0.11, where overall $F_{is}$ was 0.04. We summarize expected and observed heterozygosity levels and $F_{st}$ values for each population in **Supplementary Table 12**. Pairwise $G'_{st}$, $G_{st}$, Jost's D, and $F_{st}$ estimations were highest among Europe and other populations (**Supplementary Table 13**). Notably, Thrace and East and South Marmara populations displayed a more pronounced differentiation level than the remaining populations in Anatolia, where the overall differentiation trend was comparatively weaker.

Samples from Europe and Anatolia formed poles at opposite ends of the PCA, whereas Thrace samples stayed in between (**Supplementary Figure 8**)—a pattern repeated in the phylogeny. On the other hand, in a spatially explicit sPCA with samples excluding Europe, each of the Thracian, Levantine, Caucasian, Anatolian, and Zagrosian ancestral groups formed distinct groups (**Supplementary Figures 9, 10, and 11**). Nevertheless, the transition between ancestral groups across the space was gradual—even demonstrating further substructure within Anatolian samples centered on Coastal Aegean and Western Black Sea populations (**Supplementary Figures 12 and 13**). Given the distinct clusters in the UPGMA tree and sPCA plots, where Thrace, Eastern Mediterranean, Lesser Caucasus, Zagros, and Coastal Aegean populations form the cores, we conducted an AMOVA with only these five populations. The results of the AMOVA proved significant differentiation between these potentially unadmixed populations, highlighting their genetic distinctiveness ($p = 0.01$). (**Supplementary Table 14** and **Supplementary Figures 14, 15, 16, 17, and 18**). Thrace and Coastal Aegean populations ($p = 0.01$) differed significantly from each other, just like Zagros differed from each of the Eastern Mediterranean ($p = 0.03$), Lesser Caucasus ($p = 0.04$), and Coastal Aegean ($p = 0.01$) populations. A DAPC plot also confirmed the presence of five genetic groups in honey bee populations (**Supplementary Figures 19 and 20**). Transition patterns between populations become evident when ancestry estimates are plotted separately for ancestral groups (**Supplementary Figures 21, 22, 23, 24, and 25**).

The ratio of $R^2$ captured in Gradient Forests (GF) by climatic variables showed substantial differences within subregions. The Thracian to Anatolian transition had the lowest figure (36%), while the Caucasian to Anatolian transition had the highest (72%; **Supplementary Table 15**). Regional GF models selected fewer variables with $R^2 > 0.01$ at transition zones between subspecies pairs (**Supplementary Table 16**). Nonetheless, the regional models highlighted specific environmental predictors that made minor contributions to the global model but played a significant role in local genetic differentiation, such as Pdriest at Thracian-Anatolian (**Supplementary Figures 26 and 27**), altitude and minTwarm at Anatolian-Levantine-Zagrosian (**Supplementary Figures 28 and 29**), or Pwettest at both Anatolian-Caucasian and Caucasian-Zagrosian transitions (**Supplementary Figures 30, 31, 32, and 33**). Similarly, regional Generalized Dissimilarity Models (GDMs) provided a nuanced understanding of fine-scale turnover patterns in ancestry compositions at a local level (**Supplementary Figures 34, 35, 36, and 37**). While geographic distance was the most important variable in the global model, it dropped to either second or third in regional models. For instance, at the Anatolian-Levantine-Zagrosian transition zone, minTwarm dominated geographic distance (**Supplementary Figure 38**), as did isothermality at Thracian-Anatolian and Caucasian-Zagrosian transitions (**Supplementary Figures 39 and 40**), and PETdriest at Anatolian-Caucasian and Caucasian-Zagrosian transitions (**Supplementary Figures 41 and 42**).

In the spatial analysis of our global GDM, the five ancestral groups showed strong associations with their respective regions. Survey sites displayed distinct patterns of ecological similarity to sites unadmixed in certain ancestral groups (**Supplementary Figures 43, 44, 45, 46, and 47**). Western Anatolia exhibited higher ecological similarity to the

Thracian group, while the Colchis Coast at the eastern Black Sea showed a greater affinity for the Caucasian. High similarity regions for the Levantine group extended towards East-Central Anatolia. In supervised hierarchical clustering, increasing the number of clusters to six and seven provided insights into potential ecotypes existing below the subspecies level, influenced by distinct ecological conditions (**Supplementary Figures 48 and 49**). The newly added clusters represented coastal and inland populations of *anatoliaca* and *caucasica* bees with distinct morphological and life-history traits. In 2000 randomly selected sites, successive additions of Hakkari, Çankırı, and Muş had notable impacts on the mean gains in resemblance per cell, as per the assessment of protected areas (**Supplementary Figure 50**). Notably, including Hakkari significantly enhanced the resemblance of the least protected sites.

The influence of GDM-transformed variables across the study space underscores the differential sensitivity of genetic turnover to various environmental variables (**Supplementary Figure 51**). Notably, increases in minTwarm and alterations in PETdriest were primary drivers of the observed effects due to climate change across several SSP-period combinations (**Supplementary Figure 52**). Our results incorporating all four SSPs provide a baseline for understanding the climate vulnerability of honey bee populations.

**Supplementary Figure 53** shows the mean differences in maximum ecological similarity to sampling sites (i.e., survey gaps), uniqueness, and turnover speed in 2000 randomly selected sites under climate change. **Supplementary Figures 54, 55, and 56** track the spatial impacts of climate change, which varied across SSP scenarios and periods for survey gaps, uniqueness, and turnover speed. **Supplementary Figures 57 (Thracian), 58 (Caucasian), 59 (Anatolian), 60 (Zagrosian), and 61 (Levantine)** illustrate the projected changes in ecological similarity to unadmixed sampling locations for different ancestral groups in response to climate change. **Supplementary Figures 62, 63, 64, and 65** detail the indices of persistence, resilience, disappearance, and emergence of ancestry compositions under SSP scenarios.

#### DISCUSSION

Both fitness and climate are multidimensional, and locally adapted lineages are expected to display a wide range of responses across various vital rates and environmental drivers (DeMarche et al., 2019). The diverse behavioral and ecological traits of honey bee subspecies reflect their adaptation to varying environmental conditions. Levantine bees swarm frequently with multiple queens, are highly defensive, show high hygienic and grooming behaviors linked to neurodevelopmental and behavioral genes, and sustain low mite infestations (Kence et al., 2013; Yıldız & Karabağ, 2022). Caucasian bees benefit from sequential resource availability throughout the flowering season, are less defensive, have low swarming tendencies, and exhibit reduced flower fidelity (Brillet et al., 2002; Çakmak et al., 2010). Forager *syriaca* shows high flower constancy with low learning flexibility, which is beneficial in Mediterranean climates with prolonged summer droughts and high predator densities. Conversely, *caucasica* bees exhibit plasticity to maximize honey stores for long winters (Claudio et al., 2018).

The space-for-time substitution method, widely used across eco-evolutionary subfields, including population phenotypes and genotypes, species distributions, and ecological communities (Thomas et al., 2004; Wilczek et al., 2014; Alexander et al., 2015; Gougherty et al., 2021), serves as an effective approach to study varied responses of locally adapted lineages. Complementing this, Jay et al. (2015) utilized a model-based approach that integrates genetic and geographic data to infer admixture coefficients based on correlations with environmental variables and make forecasts. However, our specific interest lies in modeling pairwise dissimilarities between sites based on ecological gradients, for which

GDM proves highly advantageous. GDM effectively accounts for nonlinear effects of interacting variable combinations on dissimilarity, including when pairs of assemblages are entirely different, and characterizes lower dissimilarity values more accurately when incorporating proximate sites with high and low environmental similarity (Mokany et al., 2022). Our dense sampling captures relationships between ancestry compositions and the environment (Anderson et al., 2010; Landguth & Schwartz, 2014) within our study area, focusing on a historical refugium with significant environmental heterogeneity spanning almost 1 million square kilometers, using microsatellite markers. While SNP markers are informative and can identify candidate regions for local adaptation, they often lack experimental validation for fitness consequences under different environmental conditions and genetic backgrounds (Barghi et al., 2020). Additionally, collating georeferenced SNP data with uniform and comparable methods across the entire range of *Apis mellifera* poses challenges.

We used pairwise ecological similarities, which incorporate climatic variation and the distribution of ancestry compositions across the landscape, to propose additional conservation sites. Including Muş in the conservation sites could be beneficial, as it appears to be the epicenter of Anatolian ancestry in the east, where *meda* and *anatoliaca* populations interpenetrate along distinct river beds of the Euphrates and the Araxes. Moreover, due to the ecological similarity of Muş to sites harboring high Zagrosian ancestry, such a conservation measure would also benefit that group. Adding Çankırı would introduce a previously neglected diversity component of Anatolian ancestry considering the distinct ecological conditions of the Black Sea hinterland. While our proposals could not increase the protected area resemblance for the Levantine ancestral group, an examination of the resemblance layers suggests that adding Mardin could enhance conservation through increased representativeness. However, due to the lack of samples from Mardin and its immediate neighbors to assess the genetic makeup of the region directly, we refrain from suggesting this province among the proposed protected areas at this time. In addition to proposing new conservation sites sequentially removing existing protected areas and monitoring changes in resemblance values can identify overlaps and inform efficient future resource allocations.

Resemblance analysis can significantly benefit and inform conservation efforts in the face of climate change. Resemblance between future sites and future protected areas can identify potential overlaps and reveal places that would be indirectly protected at a later stage, as well as those with low ecological similarity to protected areas, which would stay uncovered. In such cases, controlled mating, including artificial insemination, can benefit conservation herds. Mating control can effectively maintain or enhance protected area resemblance. Additionally, it can be strategically utilized to bolster adaptive capacity within protected areas, mainly by assisted gene flow of identified adaptive markers (Gaitán-Espitia & Hobday, 2021). In the future, our results can be incorporated into more sophisticated decision-making and advanced systematic conservation planning tools. These tools can integrate spatial, genetic, and ecological data to optimize conservation strategies and identify priority areas for honey bee populations (Zurell et al., 2022; Nielsen et al., 2022; Andrello et al., 2022).

To enhance our understanding of future genetic landscapes under climate change, we utilize disappearance and emergence indices. These indices allow us to determine (i) current ancestry compositions that may no longer be available in the future and (ii) novel ancestry compositions that are distinct from those currently observed. Our approach for index calculation dramatically reduces the computation time needed. A single run on a quad-core 2.40 GHz laptop with 12 GB of RAM, using a set of random reference cells representing 5% of the study space, took only ten hours. The analyses would take approximately ten days if all pairwise comparisons were used. While further testing is needed for broader efficacy and capturing extremely rare sites, our method remains informative by considering minimum ecological distances between sampling and random sites.
