## Supplementary Tables for "Spatio-temporal shifts driven by climate change threaten persistence and resilience of honey bee populations"

| ID | Thr | Cau | Lev | Zag | Ana | Region | Population | Lon | Lat | A007 | A014 | A028 |
|---|---|---|---|---|---|---|---|---|---|---|---|---|
| apis_UGB4 | 0,97 | 0,00 | 0,00 | 0,01 | 0,02 | 1 | Eur | 6,12 | 51,05 | 120_120 | 217_217 | 137_137 |
| apis_UGB5 | 0,99 | 0,00 | 0,00 | 0,00 | 0,00 | 1 | Eur | 6,12 | 51,06 | 120_120 | 217_217 | 131_137 |
| apis_UGA5 | 0,99 | 0,00 | 0,00 | 0,00 | 0,00 | 1 | Eur | 6,15 | 51,06 | 113_113 | 217_234 | 131_137 |
| apis_UGA1 | 0,99 | 0,00 | 0,00 | 0,00 | 0,01 | 1 | Eur | 7,46 | 53,42 | 113_120 | 217_234 | 137_137 |
| apis_UGA2 | 0,99 | 0,00 | 0,00 | 0,00 | 0,00 | 1 | Eur | 7,46 | 53,43 | 113_120 | 217_217 | 131_137 |
| apis_UGA3 | 0,99 | 0,00 | 0,00 | 0,00 | 0,00 | 1 | Eur | 7,47 | 53,42 | 113_113 | 217_234 | 127_131 |
| apis_UGB1 | 0,98 | 0,00 | 0,00 | 0,00 | 0,01 | 1 | Eur | 9,68 | 52,17 | 113_120 | 217_234 | NA |
| apis_UGB2 | 0,99 | 0,00 | 0,00 | 0,00 | 0,00 | 1 | Eur | 9,68 | 52,18 | 113_118 | 217_234 | 137_137 |
| apis_PaxC1 | 0,96 | 0,00 | 0,01 | 0,01 | 0,02 | 1 | Eur | 11,92 | 51,49 | 126_164 | 217_223 | 131_137 |
| apis_PaxC2 | 0,99 | 0,00 | 0,00 | 0,00 | 0,00 | 1 | Eur | 11,92 | 51,50 | 113_126 | 217_217 | 131_137 |
| apis_PaxC3 | 0,87 | 0,00 | 0,01 | 0,02 | 0,10 | 1 | Eur | 11,92 | 51,51 | 113_126 | 217_217 | 137_137 |
| apis_PaxC4 | 0,99 | 0,00 | 0,00 | 0,00 | 0,01 | 1 | Eur | 11,92 | 51,52 | 118_126 | 217_217 | 131_131 |
| apis_PaxC5 | 0,98 | 0,00 | 0,01 | 0,01 | 0,01 | 1 | Eur | 11,92 | 51,53 | 120_124 | 217_217 | 131_131 |
| apis_PaxB1 | 0,99 | 0,00 | 0,00 | 0,00 | 0,01 | 1 | Eur | 11,93 | 51,49 | 113_118 | 217_217 | 131_131 |
| apis_PaxB2 | 1,00 | 0,00 | 0,00 | 0,00 | 0,00 | 1 | Eur | 11,93 | 51,50 | 118_118 | 217_217 | 131_131 |
| apis_PaxB3 | 0,99 | 0,00 | 0,00 | 0,00 | 0,00 | 1 | Eur | 11,93 | 51,51 | 113_113 | 217_217 | 131_137 |
| apis_PaxB4 | 0,99 | 0,00 | 0,00 | 0,00 | 0,00 | 1 | Eur | 11,93 | 51,52 | 113_118 | 217_217 | 137_137 |
| apis_PaxB5 | 0,97 | 0,01 | 0,00 | 0,01 | 0,01 | 1 | Eur | 11,93 | 51,53 | 113_118 | 217_223 | 131_137 |
| apis_PaxA2 | 0,98 | 0,00 | 0,00 | 0,00 | 0,01 | 1 | Eur | 11,94 | 51,49 | 113_118 | 217_217 | 137_137 |
| apis_PaxA3 | 0,98 | 0,00 | 0,00 | 0,00 | 0,01 | 1 | Eur | 11,94 | 51,50 | 113_118 | 217_217 | 137_137 |
| apis_PaxA4 | 0,96 | 0,00 | 0,03 | 0,00 | 0,01 | 1 | Eur | 11,94 | 51,52 | 118_118 | 217_217 | 137_137 |
| apis_PaxA5 | 0,97 | 0,00 | 0,00 | 0,01 | 0,01 | 1 | Eur | 11,94 | 51,53 | 118_120 | 217_217 | 132_132 |
| apis_PaxA1 | 0,99 | 0,00 | 0,00 | 0,00 | 0,00 | 1 | Eur | 11,94 | 51,51 | 113_120 | 217_217 | 137_137 |
| apis_PaxD3 | 0,91 | 0,00 | 0,02 | 0,02 | 0,05 | 1 | Eur | 11,95 | 51,49 | 118_126 | 217_217 | 131_137 |
| apis_PaxD4 | 0,99 | 0,00 | 0,00 | 0,00 | 0,00 | 1 | Eur | 11,95 | 51,52 | 120_126 | 217_217 | 137_137 |
| apis_PaxD5 | 0,98 | 0,00 | 0,00 | 0,00 | 0,01 | 1 | Eur | 11,95 | 51,53 | 118_126 | 217_217 | 131_137 |
| apis_PaxE2 | 0,96 | 0,02 | 0,00 | 0,01 | 0,02 | 1 | Eur | 11,96 | 51,50 | 118_120 | 217_217 | 137_137 |
| apis_PaxE4 | 0,98 | 0,00 | 0,01 | 0,01 | 0,01 | 1 | Eur | 11,96 | 51,52 | 118_118 | 223_234 | 137_137 |
| apis_PaxE5 | 0,98 | 0,00 | 0,01 | 0,00 | 0,01 | 1 | Eur | 11,96 | 51,53 | 118_120 | 217_217 | 131_137 |
| apis_PaxF1 | 0,99 | 0,00 | 0,00 | 0,00 | 0,00 | 1 | Eur | 11,97 | 51,49 | 113_118 | 217_224 | 137_137 |
| apis_PaxF3 | 0,87 | 0,05 | 0,03 | 0,01 | 0,05 | 1 | Eur | 11,97 | 51,51 | 118_118 | 217_217 | 137_137 |
| apis_PaxF4 | 0,99 | 0,00 | 0,00 | 0,00 | 0,01 | 1 | Eur | 11,97 | 51,52 | 118_118 | 217_217 | 137_137 |
| apis_PaxF5 | 0,97 | 0,01 | 0,00 | 0,01 | 0,01 | 1 | Eur | 11,97 | 51,53 | 118_118 | 217_217 | 137_137 |
| apis_UAA1 | 0,96 | 0,00 | 0,00 | 0,01 | 0,02 | 1 | Eur | 14,75 | 47,16 | 113_118 | 217_234 | 133_137 |
| apis_UAA2 | 0,98 | 0,00 | 0,00 | 0,00 | 0,01 | 1 | Eur | 14,75 | 47,17 | 115_115 | 217_234 | 137_137 |
| apis_UAA3 | 0,99 | 0,00 | 0,00 | 0,00 | 0,00 | 1 | Eur | 14,76 | 47,16 | 115_118 | 217_217 | 132_137 |
| apis_UAA4 | 0,99 | 0,00 | 0,00 | 0,00 | 0,01 | 1 | Eur | 15,09 | 47,04 | 115_115 | 217_217 | 137_137 |
| apis_UAA5 | 0,98 | 0,00 | 0,01 | 0,00 | 0,01 | 1 | Eur | 15,09 | 47,05 | 113_118 | 217_234 | 137_137 |
| apis_091 | 0,23 | 0,05 | 0,10 | 0,05 | 0,57 | 2 | Thrace | 25,82 | 40,11 | 139_152 | 223_223 | 136_137 |
| apis_090 | 0,45 | 0,06 | 0,01 | 0,02 | 0,47 | 2 | Thrace | 25,91 | 40,22 | 118_130 | 223_223 | 132_135 |
| apis_084 | 0,89 | 0,01 | 0,00 | 0,01 | 0,09 | 2 | Thrace | 26,17 | 40,69 | NA | NA | 133_137 |
| apis_089 | 0,78 | 0,02 | 0,06 | 0,01 | 0,12 | 2 | Thrace | 26,37 | 40,23 | NA | NA | 131_131 |
| apis_R058 | 0,18 | 0,02 | 0,01 | 0,01 | 0,78 | 2 | Thrace | 26,39 | 40,93 | NA | NA | 130_130 |
| apis_083 | 0,95 | 0,01 | 0,00 | 0,01 | 0,02 | 2 | Thrace | 26,43 | 41,12 | 122_126 | 223_234 | 133_137 |
| apis_078 | 0,97 | 0,00 | 0,00 | 0,00 | 0,02 | 2 | Thrace | 26,44 | 41,70 | NA | NA | 137_137 |
| apis_R059 | 0,97 | 0,01 | 0,00 | 0,00 | 0,02 | 2 | Thrace | 26,55 | 40,63 | 126_162 | 223_234 | 136_143 |
| apis_R060 | 0,61 | 0,01 | 0,01 | 0,20 | 0,17 | 2 | Thrace | 26,56 | 40,63 | 141_166 | 223_223 | 137_137 |
| apis_085 | 0,76 | 0,01 | 0,01 | 0,03 | 0,19 | 2 | Thrace | 26,56 | 40,68 | NA | NA | 137_137 |

| | | | | | | | | | | | | |
|---|---|---|---|---|---|---|---|---|---|---|---|---|
| apis_088 | 0,17 | 0,08 | 0,08 | 0,01 | 0,67 | 2 | Thrace | 26,57 | 40,42 | 118_120 | 223_223 | 137_137 |
| apis_R061 | 0,50 | 0,01 | 0,01 | 0,01 | 0,48 | 2 | Thrace | 26,57 | 40,68 | 118_120 | 217_217 | 131_137 |
| apis_R057 | 0,74 | 0,01 | 0,01 | 0,03 | 0,21 | 2 | Thrace | 26,64 | 40,87 | NA | NA | 137_137 |
| apis_077 | 0,39 | 0,19 | 0,01 | 0,14 | 0,27 | 2 | Thrace | 26,67 | 41,91 | NA | NA | 137_137 |
| apis_082 | 0,66 | 0,02 | 0,02 | 0,04 | 0,27 | 2 | Thrace | 26,80 | 41,40 | 118_118 | 217_223 | 137_137 |
| apis_086 | 0,40 | 0,33 | 0,01 | 0,13 | 0,13 | 2 | Thrace | 26,88 | 40,97 | NA | NA | 137_137 |
| apis_087 | 0,01 | 0,01 | 0,00 | 0,01 | 0,97 | 2 | Thrace | 26,89 | 40,61 | 144_184 | 223_223 | 137_137 |
| apis_R105 | 0,78 | 0,17 | 0,00 | 0,01 | 0,03 | 2 | Thrace | 27,01 | 41,98 | 118_134 | 223_223 | 136_136 |
| apis_076 | 0,94 | 0,00 | 0,03 | 0,01 | 0,02 | 2 | Thrace | 27,05 | 41,95 | 118_125 | 217_219 | 137_137 |
| apis_R104 | 0,95 | 0,01 | 0,01 | 0,02 | 0,02 | 2 | Thrace | 27,05 | 41,96 | NA | NA | 133_137 |
| apis_R103 | 0,98 | 0,00 | 0,00 | 0,01 | 0,01 | 2 | Thrace | 27,06 | 41,95 | 118_118 | 217_223 | 137_137 |
| apis_R126 | 0,97 | 0,01 | 0,00 | 0,00 | 0,01 | 2 | Thrace | 27,08 | 41,45 | 135_154 | 219_223 | 137_137 |
| apis_R127 | 0,80 | 0,03 | 0,02 | 0,01 | 0,14 | 2 | Thrace | 27,09 | 41,45 | NA | NA | 137_137 |
| apis_R124 | 0,96 | 0,01 | 0,01 | 0,01 | 0,02 | 2 | Thrace | 27,10 | 41,49 | NA | NA | 133_137 |
| apis_R125 | 0,35 | 0,04 | 0,01 | 0,50 | 0,11 | 2 | Thrace | 27,10 | 41,58 | 130_164 | 222_223 | 133_137 |
| apis_079 | 0,97 | 0,01 | 0,00 | 0,01 | 0,01 | 2 | Thrace | 27,11 | 41,49 | 107_120 | 223_223 | 137_137 |
| apis_081 | 0,05 | 0,02 | 0,02 | 0,02 | 0,89 | 2 | Thrace | 27,12 | 41,10 | 142_184 | 219_223 | 133_133 |
| apis_R050 | 0,98 | 0,00 | 0,00 | 0,00 | 0,01 | 2 | Thrace | 27,16 | 41,65 | 120_140 | 217_223 | 133_137 |
| apis_R118 | 0,29 | 0,04 | 0,04 | 0,05 | 0,58 | 2 | Thrace | 27,16 | 41,95 | NA | NA | 133_137 |
| apis_R123 | 0,98 | 0,00 | 0,00 | 0,01 | 0,01 | 2 | Thrace | 27,16 | 41,96 | 122_152 | 217_223 | 132_136 |
| apis_R053 | 0,93 | 0,01 | 0,00 | 0,01 | 0,05 | 2 | Thrace | 27,17 | 41,95 | 118_122 | 217_223 | 137_137 |
| apis_R067 | 0,96 | 0,01 | 0,00 | 0,01 | 0,02 | 2 | Thrace | 27,18 | 41,90 | NA | NA | 137_137 |
| apis_R121 | 0,94 | 0,01 | 0,00 | 0,01 | 0,04 | 2 | Thrace | 27,18 | 41,91 | NA | NA | 133_141 |
| apis_R133 | 0,97 | 0,00 | 0,00 | 0,01 | 0,02 | 2 | Thrace | 27,19 | 41,90 | NA | NA | 137_137 |
| apis_R066 | 0,92 | 0,01 | 0,01 | 0,01 | 0,06 | 2 | Thrace | 27,20 | 41,85 | NA | NA | 133_137 |
| apis_R122 | 0,82 | 0,01 | 0,03 | 0,06 | 0,08 | 2 | Thrace | 27,20 | 41,86 | 118_126 | 217_219 | 137_137 |
| apis_R065 | 0,87 | 0,01 | 0,01 | 0,03 | 0,09 | 2 | Thrace | 27,21 | 41,84 | NA | NA | 137_137 |
| apis_R098 | 0,95 | 0,01 | 0,01 | 0,01 | 0,02 | 2 | Thrace | 27,21 | 41,85 | 118_118 | 223_223 | 137_137 |
| apis_R099 | 0,65 | 0,01 | 0,01 | 0,04 | 0,28 | 2 | Thrace | 27,21 | 41,86 | 142_150 | 219_223 | 133_137 |
| apis_R106 | 0,94 | 0,00 | 0,01 | 0,02 | 0,03 | 2 | Thrace | 27,21 | 41,96 | NA | NA | 135_137 |
| apis_R120 | 0,72 | 0,01 | 0,09 | 0,11 | 0,06 | 2 | Thrace | 27,21 | 41,97 | 118_138 | 217_223 | 137_137 |
| apis_R056 | 0,80 | 0,01 | 0,03 | 0,03 | 0,12 | 2 | Thrace | 27,22 | 40,67 | 122_140 | 223_223 | 137_137 |
| apis_R100 | 0,97 | 0,01 | 0,00 | 0,00 | 0,01 | 2 | Thrace | 27,22 | 41,85 | NA | NA | 137_137 |
| apis_R107 | 0,83 | 0,03 | 0,05 | 0,02 | 0,08 | 2 | Thrace | 27,22 | 41,86 | NA | NA | 132_137 |
| apis_R132 | 0,97 | 0,00 | 0,01 | 0,01 | 0,02 | 2 | Thrace | 27,22 | 41,96 | 126_132 | 219_223 | 131_137 |
| apis_R062 | 0,91 | 0,01 | 0,01 | 0,02 | 0,05 | 2 | Thrace | 27,24 | 41,74 | NA | NA | 133_137 |
| apis_R063 | 0,97 | 0,01 | 0,00 | 0,00 | 0,01 | 2 | Thrace | 27,25 | 41,74 | 118_118 | 217_223 | 137_137 |
| apis_092 | 0,36 | 0,05 | 0,02 | 0,06 | 0,51 | 2 | Thrace | 27,26 | 40,70 | 107_118 | 217_223 | 133_137 |
| apis_075 | 0,64 | 0,01 | 0,01 | 0,02 | 0,33 | 2 | Thrace | 27,30 | 42,01 | 118_130 | 219_223 | 137_137 |
| apis_R102 | 0,94 | 0,00 | 0,04 | 0,01 | 0,01 | 2 | Thrace | 27,30 | 42,02 | NA | NA | 133_137 |
| apis_074 | 0,90 | 0,01 | 0,01 | 0,05 | 0,03 | 2 | Thrace | 27,30 | 41,80 | 118_118 | 217_223 | 137_137 |
| apis_R052 | 0,94 | 0,01 | 0,01 | 0,01 | 0,04 | 2 | Thrace | 27,31 | 41,81 | 120_140 | 217_223 | 137_137 |
| apis_R101 | 0,98 | 0,00 | 0,00 | 0,01 | 0,01 | 2 | Thrace | 27,31 | 42,01 | 118_128 | 217_223 | 137_137 |
| apis_R116 | 0,77 | 0,01 | 0,14 | 0,04 | 0,05 | 2 | Thrace | 27,32 | 41,81 | 118_138 | 219_219 | 137_137 |
| apis_073 | 0,25 | 0,06 | 0,02 | 0,05 | 0,62 | 2 | Thrace | 27,34 | 41,63 | 107_118 | 217_222 | 133_137 |
| apis_R049 | 0,94 | 0,01 | 0,01 | 0,01 | 0,04 | 2 | Thrace | 27,36 | 40,80 | 118_135 | 219_234 | 137_137 |
| apis_R055 | 0,81 | 0,04 | 0,01 | 0,07 | 0,07 | 2 | Thrace | 27,36 | 41,62 | 120_166 | 223_223 | 137_137 |
| apis_093 | 0,83 | 0,04 | 0,03 | 0,04 | 0,06 | 2 | Thrace | 27,40 | 40,88 | 118_122 | 217_234 | 133_133 |

| | | | | | | | | | | | | |
|---|---|---|---|---|---|---|---|---|---|---|---|---|
| apis_R051 | 0,98 | 0,00 | 0,00 | 0,00 | 0,01 | 2 | Thrace | 27,44 | 41,92 | 118_122 | 217_217 | 137_137 |
| apis_080 | 0,92 | 0,01 | 0,00 | 0,01 | 0,05 | 2 | Thrace | 27,49 | 41,36 | 130_132 | 223_223 | 137_137 |
| apis_R108 | 0,66 | 0,02 | 0,01 | 0,03 | 0,29 | 2 | Thrace | 27,50 | 41,63 | 130_146 | 219_223 | 133_137 |
| apis_R109 | 0,98 | 0,00 | 0,00 | 0,00 | 0,01 | 2 | Thrace | 27,51 | 41,63 | 132_149 | 219_223 | 137_137 |
| apis_072 | 0,69 | 0,01 | 0,02 | 0,05 | 0,23 | 2 | Thrace | 27,55 | 41,75 | 118_142 | 217_223 | 137_137 |
| apis_071 | 0,91 | 0,00 | 0,00 | 0,01 | 0,07 | 2 | Thrace | 27,58 | 41,87 | 118_118 | 217_223 | 136_137 |
| apis_R068 | 0,95 | 0,00 | 0,01 | 0,01 | 0,03 | 2 | Thrace | 27,60 | 41,62 | 118_118 | 217_223 | 137_137 |
| apis_R119 | 0,91 | 0,02 | 0,01 | 0,01 | 0,05 | 2 | Thrace | 27,61 | 41,62 | NA | NA | 131_141 |
| apis_065 | 0,77 | 0,08 | 0,02 | 0,04 | 0,09 | 2 | Thrace | 27,67 | 41,14 | NA | NA | 131_137 |
| apis_068 | 0,97 | 0,01 | 0,00 | 0,01 | 0,02 | 2 | Thrace | 27,70 | 41,59 | 118_122 | 223_223 | 137_137 |
| apis_069 | 0,44 | 0,00 | 0,01 | 0,03 | 0,52 | 2 | Thrace | 27,86 | 41,78 | 118_132 | 217_223 | 137_137 |
| apis_066 | 0,73 | 0,06 | 0,01 | 0,03 | 0,16 | 2 | Thrace | 27,89 | 41,44 | NA | NA | 133_137 |
| apis_064 | 0,67 | 0,03 | 0,03 | 0,11 | 0,16 | 2 | Thrace | 28,01 | 41,10 | 118_142 | 219_223 | 137_137 |
| apis_070 | 0,27 | 0,01 | 0,01 | 0,02 | 0,70 | 2 | Thrace | 28,01 | 41,96 | 118_148 | 223_234 | 137_137 |
| apis_067 | 0,90 | 0,04 | 0,01 | 0,02 | 0,04 | 2 | Thrace | 28,02 | 41,61 | 118_126 | 217_223 | 137_137 |
| apis_063 | 0,45 | 0,05 | 0,02 | 0,07 | 0,42 | 2 | Thrace | 28,35 | 41,27 | 118_140 | 217_219 | 137_137 |
| apis_392A | 0,30 | 0,06 | 0,04 | 0,04 | 0,56 | 2 | Thrace | 28,60 | 41,03 | 120_144 | 223_223 | 131_137 |
| apis_392B | 0,18 | 0,47 | 0,01 | 0,02 | 0,32 | 2 | Thrace | 28,60 | 41,04 | 118_158 | 217_223 | 137_137 |
| apis_062 | 0,50 | 0,03 | 0,36 | 0,03 | 0,07 | 2 | Thrace | 28,87 | 41,27 | 138_152 | 223_223 | 137_137 |
| apis_283A | 0,03 | 0,10 | 0,01 | 0,12 | 0,73 | 3 | ES.Marm | 26,01 | 39,81 | 118_145 | 217_223 | 137_137 |
| apis_283C | 0,01 | 0,01 | 0,01 | 0,01 | 0,95 | 3 | ES.Marm | 26,01 | 39,82 | 118_118 | 217_223 | 137_137 |
| apis_283D | 0,02 | 0,01 | 0,01 | 0,01 | 0,95 | 3 | ES.Marm | 26,01 | 39,83 | 107_142 | 223_223 | 137_137 |
| apis_282 | 0,06 | 0,23 | 0,01 | 0,08 | 0,62 | 3 | ES.Marm | 26,34 | 39,52 | NA | NA | 133_137 |
| apis_281 | 0,02 | 0,24 | 0,01 | 0,03 | 0,70 | 3 | ES.Marm | 26,54 | 40,09 | 128_140 | 217_217 | 133_137 |
| apis_280 | 0,57 | 0,02 | 0,02 | 0,05 | 0,34 | 3 | ES.Marm | 26,76 | 40,31 | 118_118 | 223_234 | 137_137 |
| apis_120 | 0,01 | 0,02 | 0,01 | 0,02 | 0,95 | 3 | ES.Marm | 27,04 | 39,51 | 118_138 | 219_223 | 131_137 |
| apis_119 | 0,04 | 0,14 | 0,01 | 0,37 | 0,45 | 3 | ES.Marm | 27,17 | 39,81 | 118_180 | 223_234 | 133_137 |
| apis_279 | 0,01 | 0,03 | 0,01 | 0,01 | 0,93 | 3 | ES.Marm | 27,39 | 40,22 | 107_118 | 217_217 | 137_137 |
| apis_284 | 0,02 | 0,19 | 0,01 | 0,03 | 0,75 | 3 | ES.Marm | 27,59 | 39,67 | 130_152 | 223_223 | 133_137 |
| apis_278 | 0,02 | 0,05 | 0,02 | 0,06 | 0,86 | 3 | ES.Marm | 28,10 | 40,35 | NA | NA | 137_137 |
| apis_161 | 0,06 | 0,02 | 0,01 | 0,16 | 0,74 | 3 | ES.Marm | 28,18 | 39,30 | 120_160 | 217_223 | 133_137 |
| apis_277 | 0,16 | 0,17 | 0,03 | 0,38 | 0,27 | 3 | ES.Marm | 28,47 | 40,09 | NA | NA | 133_133 |
| apis_043 | 0,03 | 0,02 | 0,01 | 0,12 | 0,83 | 3 | ES.Marm | 28,89 | 40,52 | 118_118 | 217_217 | 137_137 |
| apis_095 | 0,26 | 0,01 | 0,01 | 0,56 | 0,15 | 3 | ES.Marm | 29,13 | 41,14 | 107_120 | 223_223 | 137_137 |
| apis_275 | 0,10 | 0,67 | 0,01 | 0,13 | 0,09 | 3 | ES.Marm | 29,26 | 40,32 | NA | NA | 137_137 |
| apis_094 | 0,53 | 0,23 | 0,01 | 0,07 | 0,15 | 3 | ES.Marm | 29,50 | 41,06 | 107_156 | 223_223 | 137_137 |
| apis_042 | 0,02 | 0,02 | 0,01 | 0,03 | 0,93 | 3 | ES.Marm | 29,64 | 40,64 | 118_140 | 219_223 | 133_133 |
| apis_044 | 0,18 | 0,19 | 0,03 | 0,06 | 0,53 | 3 | ES.Marm | 29,66 | 40,92 | NA | NA | 137_137 |
| apis_274 | 0,02 | 0,01 | 0,01 | 0,03 | 0,94 | 3 | ES.Marm | 29,77 | 39,98 | 118_158 | 219_223 | 137_137 |
| apis_041 | 0,09 | 0,05 | 0,01 | 0,37 | 0,47 | 3 | ES.Marm | 29,86 | 40,64 | 118_122 | 217_223 | 137_137 |
| apis_045 | 0,39 | 0,23 | 0,02 | 0,05 | 0,32 | 3 | ES.Marm | 30,02 | 41,08 | NA | NA | 137_137 |
| apis_040 | 0,02 | 0,01 | 0,01 | 0,10 | 0,86 | 3 | ES.Marm | 30,05 | 40,79 | 118_126 | 217_223 | 137_137 |
| apis_R069 | 0,05 | 0,01 | 0,00 | 0,02 | 0,92 | 3 | ES.Marm | 30,10 | 40,12 | 118_140 | 219_223 | 131_137 |
| apis_039 | 0,10 | 0,02 | 0,01 | 0,12 | 0,74 | 3 | ES.Marm | 30,16 | 40,69 | NA | NA | 137_137 |
| apis_273 | 0,22 | 0,03 | 0,01 | 0,04 | 0,70 | 3 | ES.Marm | 30,18 | 40,20 | NA | NA | 133_137 |
| apis_046 | 0,40 | 0,26 | 0,01 | 0,02 | 0,31 | 3 | ES.Marm | 30,38 | 41,08 | 134_152 | 219_223 | 133_137 |
| apis_048 | 0,06 | 0,03 | 0,01 | 0,06 | 0,84 | 3 | ES.Marm | 30,58 | 40,81 | 135_146 | 223_223 | 137_137 |
| apis_047 | 0,04 | 0,02 | 0,01 | 0,54 | 0,39 | 3 | ES.Marm | 30,80 | 41,06 | 128_144 | 219_223 | 137_137 |

| | | | | | | | | | | | | |
|---|---|---|---|---|---|---|---|---|---|---|---|---|
| apis_009 | 0,16 | 0,02 | 0,01 | 0,11 | 0,70 | 4 | Co.Aeg | 26,39 | 38,27 | 148_186 | 217_219 | 137_137 |
| apis_010 | 0,01 | 0,03 | 0,01 | 0,02 | 0,93 | 4 | Co.Aeg | 26,45 | 38,57 | 118_135 | 217_223 | 137_137 |
| apis_147 | 0,11 | 0,01 | 0,01 | 0,05 | 0,82 | 4 | Co.Aeg | 26,61 | 38,40 | 160_162 | 222_223 | 133_137 |
| apis_146 | 0,02 | 0,02 | 0,01 | 0,05 | 0,90 | 4 | Co.Aeg | 26,79 | 38,24 | 148_150 | 219_223 | 137_137 |
| apis_012E | 0,04 | 0,13 | 0,01 | 0,13 | 0,69 | 4 | Co.Aeg | 27,05 | 38,57 | 144_152 | 219_223 | 133_133 |
| apis_012A | 0,05 | 0,03 | 0,04 | 0,06 | 0,83 | 4 | Co.Aeg | 27,06 | 38,56 | NA | NA | 131_137 |
| apis_012C | 0,02 | 0,06 | 0,01 | 0,22 | 0,69 | 4 | Co.Aeg | 27,06 | 38,57 | NA | NA | 133_137 |
| apis_012D | 0,01 | 0,02 | 0,01 | 0,02 | 0,94 | 4 | Co.Aeg | 27,06 | 38,58 | NA | NA | 133_133 |
| apis_012B | 0,04 | 0,04 | 0,01 | 0,13 | 0,79 | 4 | Co.Aeg | 27,07 | 38,57 | NA | NA | 131_137 |
| apis_166 | 0,03 | 0,10 | 0,01 | 0,07 | 0,79 | 4 | Co.Aeg | 27,10 | 38,64 | NA | NA | 133_137 |
| apis_165 | 0,08 | 0,02 | 0,01 | 0,09 | 0,80 | 4 | Co.Aeg | 27,12 | 38,85 | 120_154 | 219_223 | 133_137 |
| apis_148 | 0,34 | 0,18 | 0,02 | 0,24 | 0,22 | 4 | Co.Aeg | 27,13 | 38,01 | NA | NA | 133_137 |
| apis_149 | 0,05 | 0,01 | 0,01 | 0,10 | 0,84 | 4 | Co.Aeg | 27,23 | 37,71 | NA | NA | 137_137 |
| apis_151 | 0,03 | 0,09 | 0,02 | 0,10 | 0,77 | 4 | Co.Aeg | 27,24 | 37,47 | NA | NA | 133_137 |
| apis_011 | 0,03 | 0,02 | 0,01 | 0,02 | 0,93 | 4 | Co.Aeg | 27,25 | 39,10 | NA | NA | 133_137 |
| apis_167 | 0,03 | 0,03 | 0,01 | 0,07 | 0,86 | 4 | Co.Aeg | 27,57 | 38,42 | NA | NA | 137_137 |
| apis_150 | 0,03 | 0,01 | 0,01 | 0,05 | 0,89 | 4 | Co.Aeg | 27,63 | 37,68 | NA | NA | 137_137 |
| apis_R081 | 0,07 | 0,03 | 0,01 | 0,05 | 0,83 | 4 | Co.Aeg | 27,67 | 37,17 | NA | NA | 133_137 |
| apis_152 | 0,11 | 0,15 | 0,02 | 0,26 | 0,46 | 4 | Co.Aeg | 27,68 | 37,17 | NA | NA | 133_137 |
| apis_R083 | 0,05 | 0,01 | 0,01 | 0,13 | 0,80 | 4 | Co.Aeg | 27,68 | 37,18 | 152_174 | 219_223 | 137_137 |
| apis_004 | 0,26 | 0,26 | 0,01 | 0,31 | 0,17 | 4 | Co.Aeg | 27,71 | 36,77 | NA | NA | 133_137 |
| apis_160 | 0,01 | 0,03 | 0,02 | 0,03 | 0,91 | 4 | Co.Aeg | 27,89 | 38,88 | 126_144 | 219_234 | 133_143 |
| apis_008 | 0,02 | 0,05 | 0,01 | 0,49 | 0,43 | 4 | Co.Aeg | 27,93 | 37,91 | NA | NA | 133_137 |
| apis_R089 | 0,03 | 0,02 | 0,03 | 0,05 | 0,88 | 4 | Co.Aeg | 27,96 | 37,05 | NA | NA | 133_137 |
| apis_R090 | 0,01 | 0,37 | 0,08 | 0,06 | 0,49 | 4 | Co.Aeg | 27,97 | 37,05 | NA | NA | 137_137 |
| apis_R080 | 0,01 | 0,01 | 0,02 | 0,39 | 0,57 | 4 | Co.Aeg | 28,05 | 37,09 | NA | NA | 133_137 |
| apis_R086 | 0,01 | 0,07 | 0,04 | 0,02 | 0,86 | 4 | Co.Aeg | 28,05 | 37,10 | NA | NA | 133_137 |
| apis_R084 | 0,01 | 0,41 | 0,01 | 0,12 | 0,46 | 4 | Co.Aeg | 28,06 | 37,09 | NA | NA | 137_137 |
| apis_159 | 0,05 | 0,12 | 0,02 | 0,07 | 0,75 | 4 | Co.Aeg | 28,08 | 38,35 | 107_122 | 219_219 | 137_137 |
| apis_005 | 0,02 | 0,01 | 0,03 | 0,08 | 0,86 | 4 | Co.Aeg | 28,12 | 36,74 | 122_158 | 219_223 | 133_137 |
| apis_006 | 0,04 | 0,01 | 0,02 | 0,23 | 0,71 | 4 | Co.Aeg | 28,23 | 37,56 | 118_138 | 223_223 | 133_133 |
| apis_003 | 0,03 | 0,03 | 0,01 | 0,17 | 0,76 | 4 | Co.Aeg | 28,38 | 37,10 | NA | NA | 137_137 |
| apis_007 | 0,03 | 0,01 | 0,01 | 0,05 | 0,90 | 4 | Co.Aeg | 28,54 | 37,83 | NA | NA | 135_137 |
| apis_154 | 0,02 | 0,01 | 0,01 | 0,02 | 0,94 | 4 | Co.Aeg | 28,55 | 36,84 | NA | NA | 137_137 |
| apis_R161 | 0,01 | 0,02 | 0,02 | 0,49 | 0,46 | 4 | Co.Aeg | 28,82 | 36,71 | NA | NA | 132_137 |
| apis_002 | 0,05 | 0,01 | 0,02 | 0,10 | 0,83 | 4 | Co.Aeg | 28,84 | 36,79 | 138_146 | 223_223 | 133_137 |
| apis_001 | 0,04 | 0,01 | 0,03 | 0,13 | 0,79 | 4 | Co.Aeg | 29,15 | 36,66 | 154_178 | 219_222 | 137_137 |
| apis_R026 | 0,01 | 0,03 | 0,03 | 0,11 | 0,83 | 4 | Co.Aeg | 29,19 | 36,77 | NA | NA | 133_137 |
| apis_R025 | 0,03 | 0,04 | 0,02 | 0,03 | 0,89 | 4 | Co.Aeg | 29,20 | 36,76 | 144_144 | 223_223 | 137_137 |
| apis_R029 | 0,01 | 0,02 | 0,01 | 0,08 | 0,88 | 4 | Co.Aeg | 29,20 | 36,78 | 118_118 | 223_225 | 137_140 |
| apis_155E | 0,14 | 0,02 | 0,01 | 0,03 | 0,81 | 4 | Co.Aeg | 29,20 | 36,77 | 138_146 | 219_219 | 137_137 |
| apis_155A | 0,02 | 0,63 | 0,01 | 0,12 | 0,22 | 4 | Co.Aeg | 29,21 | 36,76 | 135_154 | 223_223 | 133_137 |
| apis_155C | 0,01 | 0,58 | 0,01 | 0,14 | 0,26 | 4 | Co.Aeg | 29,21 | 36,77 | 122_172 | 223_223 | 137_137 |
| apis_155D | 0,04 | 0,08 | 0,01 | 0,07 | 0,80 | 4 | Co.Aeg | 29,21 | 36,78 | 135_194 | 223_223 | 137_137 |
| apis_R024 | 0,01 | 0,01 | 0,02 | 0,03 | 0,94 | 4 | Co.Aeg | 29,21 | 36,75 | 118_170 | 223_223 | 137_137 |
| apis_R027 | 0,01 | 0,01 | 0,01 | 0,01 | 0,97 | 4 | Co.Aeg | 29,21 | 36,79 | NA | NA | 137_137 |
| apis_155B | 0,02 | 0,02 | 0,01 | 0,10 | 0,85 | 4 | Co.Aeg | 29,22 | 36,77 | NA | NA | 137_137 |
| apis_R030 | 0,04 | 0,05 | 0,24 | 0,16 | 0,51 | 4 | Co.Aeg | 29,22 | 36,62 | NA | NA | 137_137 |

| | | | | | | | | | | | | |
|---|---|---|---|---|---|---|---|---|---|---|---|---|
| apis_R031 | 0,01 | 0,02 | 0,02 | 0,07 | 0,88 | 4 | Co.Aeg | 29,22 | 36,76 | NA | NA | 137_137 |
| apis_R032 | 0,02 | 0,25 | 0,02 | 0,27 | 0,44 | 4 | Co.Aeg | 29,22 | 36,78 | NA | NA | 128_133 |
| apis_R085 | 0,01 | 0,13 | 0,02 | 0,05 | 0,79 | 4 | Co.Aeg | 29,22 | 36,79 | 140_168 | 219_223 | 137_137 |
| apis_R028 | 0,01 | 0,01 | 0,00 | 0,03 | 0,94 | 4 | Co.Aeg | 29,23 | 36,76 | NA | NA | 131_137 |
| apis_R033 | 0,03 | 0,09 | 0,01 | 0,29 | 0,58 | 4 | Co.Aeg | 29,23 | 36,77 | 118_146 | 223_223 | 133_137 |
| apis_R034 | 0,01 | 0,01 | 0,00 | 0,09 | 0,88 | 4 | Co.Aeg | 29,23 | 36,78 | 118_118 | 222_223 | 137_137 |
| apis_R091 | 0,02 | 0,04 | 0,05 | 0,04 | 0,86 | 4 | Co.Aeg | 29,48 | 36,67 | NA | NA | 132_137 |
| apis_153 | 0,02 | 0,02 | 0,01 | 0,02 | 0,94 | 5 | W.Anat | 28,74 | 37,31 | NA | NA | 133_137 |
| apis_164 | 0,02 | 0,02 | 0,03 | 0,05 | 0,88 | 5 | W.Anat | 28,76 | 38,54 | NA | NA | 133_137 |
| apis_162 | 0,02 | 0,08 | 0,01 | 0,09 | 0,80 | 5 | W.Anat | 28,88 | 39,14 | NA | NA | 137_137 |
| apis_158 | 0,06 | 0,21 | 0,11 | 0,13 | 0,49 | 5 | W.Anat | 28,91 | 37,96 | NA | NA | 137_137 |
| apis_276 | 0,13 | 0,02 | 0,02 | 0,07 | 0,76 | 5 | W.Anat | 29,00 | 39,93 | NA | NA | 137_137 |
| apis_163 | 0,08 | 0,01 | 0,02 | 0,14 | 0,74 | 5 | W.Anat | 29,30 | 38,64 | NA | NA | 133_137 |
| apis_286 | 0,08 | 0,72 | 0,01 | 0,11 | 0,08 | 5 | W.Anat | 29,30 | 39,50 | NA | NA | 137_137 |
| apis_156 | 0,24 | 0,34 | 0,01 | 0,05 | 0,37 | 5 | W.Anat | 29,35 | 37,24 | NA | NA | 137_144 |
| apis_157 | 0,09 | 0,02 | 0,02 | 0,31 | 0,56 | 5 | W.Anat | 29,44 | 37,85 | NA | NA | 133_137 |
| apis_R070 | 0,02 | 0,01 | 0,01 | 0,01 | 0,96 | 5 | W.Anat | 29,58 | 39,38 | 118_180 | 223_223 | 133_137 |
| apis_R074 | 0,55 | 0,04 | 0,03 | 0,07 | 0,31 | 5 | W.Anat | 29,59 | 39,38 | NA | NA | 137_137 |
| apis_227 | 0,02 | 0,31 | 0,01 | 0,53 | 0,13 | 5 | W.Anat | 29,59 | 36,21 | NA | NA | 133_137 |
| apis_287 | 0,01 | 0,02 | 0,03 | 0,22 | 0,72 | 5 | W.Anat | 29,83 | 39,27 | NA | NA | 137_137 |
| apis_224 | 0,03 | 0,02 | 0,02 | 0,04 | 0,89 | 5 | W.Anat | 29,87 | 36,84 | NA | NA | 133_133 |
| apis_288 | 0,02 | 0,07 | 0,02 | 0,39 | 0,50 | 5 | W.Anat | 29,92 | 39,41 | NA | NA | 137_137 |
| apis_323 | 0,05 | 0,04 | 0,01 | 0,03 | 0,86 | 5 | W.Anat | 30,22 | 37,54 | NA | NA | 132_137 |
| apis_322 | 0,01 | 0,03 | 0,02 | 0,47 | 0,47 | 5 | W.Anat | 30,27 | 38,14 | NA | NA | 132_137 |
| apis_226 | 0,02 | 0,08 | 0,02 | 0,66 | 0,23 | 5 | W.Anat | 30,43 | 36,49 | NA | NA | 137_137 |
| apis_225 | 0,04 | 0,12 | 0,02 | 0,08 | 0,73 | 5 | W.Anat | 30,46 | 37,16 | NA | NA | 137_137 |
| apis_337 | 0,02 | 0,02 | 0,03 | 0,34 | 0,60 | 5 | W.Anat | 30,65 | 38,68 | NA | NA | 137_137 |
| apis_271 | 0,01 | 0,10 | 0,01 | 0,02 | 0,86 | 5 | W.Anat | 30,73 | 39,37 | NA | NA | 133_137 |
| apis_272 | 0,05 | 0,30 | 0,03 | 0,08 | 0,54 | 5 | W.Anat | 30,79 | 39,89 | NA | NA | 137_137 |
| apis_324 | 0,01 | 0,02 | 0,06 | 0,03 | 0,87 | 5 | W.Anat | 30,99 | 37,99 | NA | NA | 137_137 |
| apis_R072 | 0,02 | 0,36 | 0,01 | 0,25 | 0,36 | 5 | W.Anat | 31,06 | 39,64 | NA | NA | 137_137 |
| apis_R241 | 0,01 | 0,02 | 0,01 | 0,01 | 0,95 | 5 | W.Anat | 31,06 | 39,65 | NA | NA | 132_137 |
| apis_R071 | 0,03 | 0,47 | 0,08 | 0,04 | 0,38 | 5 | W.Anat | 31,07 | 39,63 | NA | NA | 137_137 |
| apis_R073 | 0,01 | 0,17 | 0,01 | 0,04 | 0,78 | 5 | W.Anat | 31,07 | 39,64 | 118_124 | 223_223 | 137_137 |
| apis_R239 | 0,04 | 0,02 | 0,00 | 0,01 | 0,93 | 5 | W.Anat | 31,07 | 39,65 | 132_138 | 223_223 | 137_137 |
| apis_R238 | 0,02 | 0,30 | 0,01 | 0,07 | 0,61 | 5 | W.Anat | 31,08 | 39,63 | 122_140 | 223_223 | 137_137 |
| apis_R240 | 0,01 | 0,05 | 0,14 | 0,03 | 0,77 | 5 | W.Anat | 31,08 | 39,64 | 118_154 | 219_219 | 137_137 |
| apis_R242 | 0,03 | 0,03 | 0,09 | 0,03 | 0,83 | 5 | W.Anat | 31,08 | 39,65 | 120_150 | 223_223 | 137_137 |
| apis_223 | 0,02 | 0,06 | 0,10 | 0,57 | 0,26 | 5 | W.Anat | 31,11 | 37,30 | 138_158 | 223_223 | 133_137 |
| apis_321 | 0,04 | 0,04 | 0,01 | 0,02 | 0,90 | 5 | W.Anat | 31,27 | 38,85 | 134_156 | 223_223 | 137_137 |
| apis_320 | 0,01 | 0,05 | 0,02 | 0,22 | 0,70 | 5 | W.Anat | 31,31 | 38,90 | 120_122 | 223_223 | 133_137 |
| apis_222 | 0,01 | 0,01 | 0,02 | 0,89 | 0,06 | 5 | W.Anat | 31,31 | 36,88 | NA | NA | 137_137 |
| apis_R019 | 0,01 | 0,05 | 0,08 | 0,24 | 0,62 | 5 | W.Anat | 31,33 | 40,20 | NA | NA | 137_137 |
| apis_R011 | 0,03 | 0,01 | 0,03 | 0,02 | 0,91 | 5 | W.Anat | 31,34 | 40,19 | NA | NA | 137_137 |
| apis_R021 | 0,02 | 0,01 | 0,01 | 0,01 | 0,96 | 5 | W.Anat | 31,34 | 40,20 | NA | NA | 133_133 |
| apis_R023 | 0,34 | 0,03 | 0,01 | 0,06 | 0,55 | 5 | W.Anat | 31,34 | 40,21 | NA | NA | 137_137 |
| apis_R022 | 0,01 | 0,03 | 0,01 | 0,10 | 0,86 | 5 | W.Anat | 31,35 | 40,20 | 140_156 | 219_223 | 137_137 |
| apis_034 | 0,01 | 0,24 | 0,01 | 0,05 | 0,69 | 5 | W.Anat | 31,39 | 40,07 | 132_134 | 219_223 | 133_137 |

| | | | | | | | | | | | | |
|---|---|---|---|---|---|---|---|---|---|---|---|---|
| apis_325 | 0,02 | 0,03 | 0,04 | 0,08 | 0,83 | 5 | W.Anat | 31,75 | 37,68 | 120_140 | 219_219 | 137_137 |
| apis_221 | 0,34 | 0,12 | 0,01 | 0,17 | 0,36 | 5 | W.Anat | 31,76 | 37,29 | 130_141 | 219_219 | 137_137 |
| apis_270 | 0,02 | 0,01 | 0,02 | 0,06 | 0,89 | 5 | W.Anat | 31,84 | 39,33 | 126_140 | 219_223 | 137_137 |
| apis_220 | 0,05 | 0,02 | 0,02 | 0,76 | 0,15 | 5 | W.Anat | 32,14 | 36,58 | NA | NA | 137_137 |
| apis_269 | 0,04 | 0,75 | 0,01 | 0,02 | 0,18 | 5 | W.Anat | 32,24 | 39,60 | 140_154 | 219_223 | 137_137 |
| apis_336 | 0,11 | 0,01 | 0,11 | 0,30 | 0,47 | 5 | W.Anat | 32,37 | 38,77 | 107_107 | 223_223 | 133_133 |
| apis_035 | 0,01 | 0,03 | 0,01 | 0,02 | 0,93 | 5 | W.Anat | 32,39 | 39,95 | 103_118 | 223_223 | 137_137 |
| apis_326 | 0,05 | 0,06 | 0,06 | 0,26 | 0,57 | 5 | W.Anat | 32,39 | 37,22 | 118_120 | 223_223 | 137_137 |
| apis_R005 | 0,01 | 0,02 | 0,01 | 0,14 | 0,83 | 5 | W.Anat | 32,49 | 39,47 | 142_156 | 219_223 | 137_137 |
| apis_335A | 0,16 | 0,01 | 0,34 | 0,10 | 0,39 | 5 | W.Anat | 32,61 | 38,13 | 142_160 | 223_223 | 137_137 |
| apis_335B | 0,41 | 0,08 | 0,05 | 0,07 | 0,39 | 5 | W.Anat | 32,62 | 38,13 | 135_142 | 219_223 | 137_137 |
| apis_R009 | 0,01 | 0,05 | 0,01 | 0,06 | 0,88 | 5 | W.Anat | 32,78 | 39,94 | 130_144 | 223_223 | 133_137 |
| apis_219 | 0,04 | 0,02 | 0,02 | 0,52 | 0,40 | 5 | W.Anat | 32,81 | 36,25 | 118_146 | 223_223 | 137_137 |
| apis_218 | 0,08 | 0,09 | 0,02 | 0,47 | 0,34 | 5 | W.Anat | 32,89 | 36,77 | 120_162 | 219_223 | 133_137 |
| apis_333 | 0,02 | 0,04 | 0,04 | 0,03 | 0,86 | 5 | W.Anat | 32,98 | 38,69 | NA | NA | 137_137 |
| apis_319 | 0,03 | 0,02 | 0,06 | 0,35 | 0,53 | 5 | W.Anat | 33,05 | 39,64 | NA | NA | 137_137 |
| apis_334 | 0,02 | 0,10 | 0,01 | 0,05 | 0,82 | 5 | W.Anat | 33,08 | 38,03 | NA | NA | 133_137 |
| apis_289 | 0,03 | 0,03 | 0,01 | 0,11 | 0,82 | 5 | W.Anat | 33,16 | 40,08 | NA | NA | 137_137 |
| apis_217 | 0,57 | 0,02 | 0,28 | 0,03 | 0,09 | 5 | W.Anat | 33,22 | 36,78 | 107_133 | 219_223 | 137_137 |
| apis_327 | 0,01 | 0,03 | 0,01 | 0,07 | 0,87 | 5 | W.Anat | 33,37 | 37,19 | 103_120 | 219_234 | 137_137 |
| apis_216 | 0,02 | 0,28 | 0,08 | 0,08 | 0,54 | 5 | W.Anat | 33,53 | 36,84 | 122_172 | 223_223 | 133_137 |
| apis_332 | 0,04 | 0,01 | 0,35 | 0,54 | 0,07 | 5 | W.Anat | 33,63 | 39,12 | 140_162 | 219_219 | 133_137 |
| apis_318 | 0,03 | 0,02 | 0,05 | 0,26 | 0,64 | 5 | W.Anat | 33,77 | 40,66 | NA | NA | 131_131 |
| apis_290 | 0,04 | 0,07 | 0,02 | 0,68 | 0,20 | 5 | W.Anat | 33,98 | 39,97 | NA | NA | 137_137 |
| apis_038 | 0,01 | 0,61 | 0,02 | 0,18 | 0,19 | 6 | W.BlkS | 30,42 | 40,51 | 130_140 | 223_223 | 137_137 |
| apis_037 | 0,08 | 0,38 | 0,01 | 0,02 | 0,51 | 6 | W.BlkS | 30,92 | 40,42 | NA | NA | 133_137 |
| apis_049 | 0,68 | 0,03 | 0,01 | 0,13 | 0,15 | 6 | W.BlkS | 30,98 | 40,83 | 148_160 | 219_223 | 137_137 |
| apis_051 | 0,21 | 0,21 | 0,01 | 0,40 | 0,16 | 6 | W.BlkS | 31,16 | 41,03 | 126_142 | NA | 133_137 |
| apis_R191 | 0,01 | 0,07 | 0,03 | 0,07 | 0,81 | 6 | W.BlkS | 31,44 | 40,95 | 154_162 | 219_219 | 137_137 |
| apis_R193 | 0,01 | 0,02 | 0,00 | 0,01 | 0,95 | 6 | W.BlkS | 31,44 | 40,96 | 144_152 | 219_219 | 133_137 |
| apis_R194 | 0,02 | 0,01 | 0,01 | 0,02 | 0,95 | 6 | W.BlkS | 31,44 | 40,97 | 135_154 | 219_223 | 137_137 |
| apis_R186 | 0,02 | 0,02 | 0,00 | 0,02 | 0,94 | 6 | W.BlkS | 31,45 | 40,95 | 130_134 | 222_223 | 133_137 |
| apis_R190 | 0,02 | 0,02 | 0,00 | 0,01 | 0,94 | 6 | W.BlkS | 31,45 | 40,96 | 107_138 | 223_223 | 133_137 |
| apis_R195 | 0,04 | 0,03 | 0,01 | 0,03 | 0,89 | 6 | W.BlkS | 31,45 | 40,97 | 140_144 | 223_223 | 137_137 |
| apis_R187 | 0,04 | 0,40 | 0,01 | 0,05 | 0,50 | 6 | W.BlkS | 31,46 | 40,95 | NA | NA | 137_137 |
| apis_R188 | 0,10 | 0,11 | 0,01 | 0,03 | 0,75 | 6 | W.BlkS | 31,46 | 40,96 | NA | NA | 137_137 |
| apis_R189 | 0,01 | 0,01 | 0,02 | 0,01 | 0,94 | 6 | W.BlkS | 31,46 | 40,97 | NA | NA | 133_137 |
| apis_R192 | 0,20 | 0,04 | 0,01 | 0,02 | 0,74 | 6 | W.BlkS | 31,47 | 40,96 | NA | NA | 137_137 |
| apis_050 | 0,01 | 0,80 | 0,01 | 0,02 | 0,16 | 6 | W.BlkS | 31,49 | 40,72 | NA | NA | 137_137 |
| apis_053 | 0,06 | 0,04 | 0,02 | 0,02 | 0,86 | 6 | W.BlkS | 31,54 | 40,96 | NA | NA | 137_137 |
| apis_036 | 0,02 | 0,01 | 0,04 | 0,06 | 0,87 | 6 | W.BlkS | 31,55 | 40,53 | NA | NA | 137_139 |
| apis_052 | 0,03 | 0,13 | 0,01 | 0,29 | 0,53 | 6 | W.BlkS | 31,57 | 41,35 | NA | NA | 137_137 |
| apis_R076 | 0,02 | 0,01 | 0,01 | 0,29 | 0,67 | 6 | W.BlkS | 31,70 | 40,62 | NA | NA | 137_137 |
| apis_R078 | 0,16 | 0,01 | 0,01 | 0,01 | 0,81 | 6 | W.BlkS | 31,70 | 40,63 | NA | NA | 137_137 |
| apis_R079 | 0,02 | 0,01 | 0,01 | 0,01 | 0,95 | 6 | W.BlkS | 31,70 | 40,64 | 152_176 | 219_223 | 133_137 |
| apis_R075 | 0,02 | 0,02 | 0,01 | 0,02 | 0,93 | 6 | W.BlkS | 31,71 | 40,63 | NA | NA | 133_137 |
| apis_R077 | 0,11 | 0,02 | 0,00 | 0,01 | 0,86 | 6 | W.BlkS | 31,82 | 41,28 | NA | NA | 133_137 |
| apis_R017 | 0,01 | 0,01 | 0,01 | 0,01 | 0,96 | 6 | W.BlkS | 31,90 | 40,19 | NA | NA | 137_137 |

| | | | | | | | | | | | | |
|---|---|---|---|---|---|---|---|---|---|---|---|---|
| apis_R047 | 0,02 | 0,03 | 0,03 | 0,54 | 0,38 | 6 | W.BlkS | 31,90 | 40,20 | NA | NA | 137_137 |
| apis_R016 | 0,01 | 0,05 | 0,03 | 0,35 | 0,56 | 6 | W.BlkS | 31,91 | 40,18 | 120_134 | 223_223 | 137_142 |
| apis_R018 | 0,01 | 0,04 | 0,01 | 0,02 | 0,92 | 6 | W.BlkS | 31,91 | 40,19 | 135_152 | 219_219 | 137_137 |
| apis_R045 | 0,01 | 0,02 | 0,14 | 0,16 | 0,67 | 6 | W.BlkS | 31,91 | 40,20 | 118_120 | 223_223 | 133_137 |
| apis_R020 | 0,00 | 0,06 | 0,00 | 0,17 | 0,76 | 6 | W.BlkS | 31,92 | 40,18 | 126_138 | 219_223 | 137_137 |
| apis_R046 | 0,01 | 0,01 | 0,01 | 0,06 | 0,91 | 6 | W.BlkS | 31,92 | 40,19 | NA | NA | 133_137 |
| apis_R048 | 0,01 | 0,92 | 0,01 | 0,03 | 0,03 | 6 | W.BlkS | 31,92 | 40,20 | NA | NA | 137_137 |
| apis_058 | 0,16 | 0,06 | 0,09 | 0,18 | 0,52 | 6 | W.BlkS | 31,93 | 41,15 | NA | NA | 137_140 |
| apis_033 | 0,06 | 0,02 | 0,04 | 0,42 | 0,47 | 6 | W.BlkS | 31,93 | 40,28 | NA | NA | 131_137 |
| apis_054 | 0,04 | 0,12 | 0,01 | 0,21 | 0,62 | 6 | W.BlkS | 32,07 | 41,55 | NA | NA | 137_137 |
| apis_060 | 0,02 | 0,02 | 0,04 | 0,19 | 0,73 | 6 | W.BlkS | 32,22 | 40,59 | NA | NA | 137_137 |
| apis_059 | 0,12 | 0,02 | 0,01 | 0,03 | 0,82 | 6 | W.BlkS | 32,30 | 40,96 | NA | NA | 137_137 |
| apis_055 | 0,22 | 0,03 | 0,01 | 0,03 | 0,72 | 6 | W.BlkS | 32,52 | 41,45 | NA | NA | 137_137 |
| apis_057 | 0,76 | 0,01 | 0,01 | 0,09 | 0,12 | 6 | W.BlkS | 32,57 | 41,79 | NA | NA | 137_137 |
| apis_R014 | 0,01 | 0,89 | 0,02 | 0,06 | 0,02 | 6 | W.BlkS | 32,62 | 40,50 | NA | NA | 137_137 |
| apis_R004 | 0,01 | 0,20 | 0,02 | 0,67 | 0,10 | 6 | W.BlkS | 32,68 | 40,47 | NA | NA | 133_133 |
| apis_R008 | 0,04 | 0,04 | 0,01 | 0,05 | 0,86 | 6 | W.BlkS | 32,69 | 40,47 | NA | NA | 131_137 |
| apis_061 | 0,01 | 0,05 | 0,02 | 0,09 | 0,82 | 6 | W.BlkS | 32,79 | 40,58 | NA | NA | 137_137 |
| apis_056 | 0,02 | 0,02 | 0,12 | 0,08 | 0,77 | 6 | W.BlkS | 32,89 | 41,68 | NA | NA | 137_137 |
| apis_317 | 0,09 | 0,02 | 0,27 | 0,33 | 0,29 | 6 | W.BlkS | 33,00 | 41,10 | NA | NA | 137_137 |
| apis_316 | 0,02 | 0,12 | 0,01 | 0,11 | 0,74 | 6 | W.BlkS | 33,58 | 41,47 | NA | NA | 137_137 |
| apis_313 | 0,08 | 0,01 | 0,11 | 0,24 | 0,57 | 6 | W.BlkS | 33,86 | 41,90 | NA | NA | 137_137 |
| apis_314 | 0,64 | 0,01 | 0,15 | 0,06 | 0,14 | 6 | W.BlkS | 33,92 | 41,89 | NA | NA | NA |
| apis_312 | 0,07 | 0,46 | 0,16 | 0,06 | 0,24 | 6 | W.BlkS | 34,01 | 41,41 | NA | NA | 137_137 |
| apis_311 | 0,01 | 0,50 | 0,01 | 0,04 | 0,44 | 6 | W.BlkS | 34,07 | 41,08 | NA | NA | 137_137 |
| apis_306 | 0,09 | 0,12 | 0,03 | 0,16 | 0,60 | 6 | W.BlkS | 34,36 | 41,90 | NA | NA | 137_137 |
| apis_307 | 0,04 | 0,14 | 0,21 | 0,31 | 0,31 | 6 | W.BlkS | 34,65 | 41,41 | NA | NA | 137_137 |
| apis_305 | 0,01 | 0,04 | 0,02 | 0,08 | 0,85 | 6 | W.BlkS | 34,93 | 42,02 | NA | NA | 132_137 |
| apis_304 | 0,01 | 0,19 | 0,18 | 0,07 | 0,54 | 6 | W.BlkS | 35,16 | 41,75 | NA | NA | 137_137 |
| apis_310 | 0,18 | 0,16 | 0,01 | 0,05 | 0,60 | 7 | C.BlkS | 34,66 | 40,41 | NA | NA | 137_137 |
| apis_309 | 0,01 | 0,08 | 0,01 | 0,22 | 0,67 | 7 | C.BlkS | 34,89 | 40,95 | NA | NA | 133_137 |
| apis_308 | 0,13 | 0,15 | 0,05 | 0,30 | 0,38 | 7 | C.BlkS | 35,30 | 41,18 | NA | NA | 133_137 |
| apis_299A | 0,04 | 0,49 | 0,02 | 0,17 | 0,27 | 7 | C.BlkS | 35,58 | 40,66 | NA | NA | 133_137 |
| apis_299B | 0,02 | 0,08 | 0,01 | 0,03 | 0,87 | 7 | C.BlkS | 35,59 | 40,66 | NA | NA | 133_137 |
| apis_303 | 0,01 | 0,03 | 0,03 | 0,08 | 0,85 | 7 | C.BlkS | 35,79 | 41,37 | NA | NA | 137_137 |
| apis_302 | 0,01 | 0,15 | 0,11 | 0,67 | 0,06 | 7 | C.BlkS | 35,96 | 41,69 | NA | NA | 137_137 |
| apis_300 | 0,08 | 0,05 | 0,01 | 0,17 | 0,69 | 7 | C.BlkS | 36,14 | 41,00 | NA | NA | 132_137 |
| apis_267 | 0,01 | 0,02 | 0,13 | 0,08 | 0,77 | 7 | C.BlkS | 36,17 | 40,15 | NA | NA | 137_137 |
| apis_301 | 0,01 | 0,82 | 0,02 | 0,05 | 0,10 | 7 | C.BlkS | 36,75 | 41,29 | NA | NA | 137_137 |
| apis_298 | 0,01 | 0,45 | 0,03 | 0,32 | 0,18 | 7 | C.BlkS | 36,82 | 40,46 | NA | NA | 133_137 |
| apis_297 | 0,01 | 0,89 | 0,01 | 0,03 | 0,06 | 7 | C.BlkS | 36,95 | 40,76 | NA | NA | 133_137 |
| apis_296 | 0,04 | 0,07 | 0,02 | 0,75 | 0,12 | 7 | C.BlkS | 37,24 | 41,07 | NA | NA | 137_137 |
| apis_292 | 0,01 | 0,17 | 0,02 | 0,05 | 0,75 | 7 | C.BlkS | 37,59 | 40,25 | NA | NA | 137_137 |
| apis_294 | 0,01 | 0,12 | 0,04 | 0,57 | 0,26 | 7 | C.BlkS | 37,66 | 40,84 | NA | NA | 137_137 |
| apis_295 | 0,03 | 0,32 | 0,07 | 0,24 | 0,34 | 7 | C.BlkS | 37,76 | 41,11 | NA | NA | 137_137 |
| apis_293 | 0,02 | 0,93 | 0,01 | 0,01 | 0,03 | 7 | C.BlkS | 37,88 | 40,56 | NA | NA | 137_137 |
| apis_168 | 0,18 | 0,01 | 0,13 | 0,48 | 0,19 | 7 | C.BlkS | 38,18 | 40,86 | NA | NA | 137_137 |
| apis_170 | 0,01 | 0,03 | 0,02 | 0,55 | 0,39 | 7 | C.BlkS | 38,34 | 40,25 | NA | NA | 137_137 |

| | | | | | | | | | | | | |
|---|---|---|---|---|---|---|---|---|---|---|---|---|
| apis_169 | 0,01 | 0,07 | 0,01 | 0,57 | 0,34 | 7 | C.BlkS | 38,47 | 40,73 | NA | NA | 137_137 |
| apis_171 | 0,01 | 0,28 | 0,02 | 0,03 | 0,66 | 7 | C.BlkS | 38,75 | 40,38 | NA | NA | 137_137 |
| apis_175 | 0,03 | 0,17 | 0,03 | 0,04 | 0,73 | 7 | C.BlkS | 38,86 | 40,88 | NA | NA | 137_137 |
| apis_174 | 0,14 | 0,04 | 0,01 | 0,22 | 0,60 | 7 | C.BlkS | 39,18 | 40,50 | NA | NA | 137_137 |
| apis_176 | 0,02 | 0,15 | 0,04 | 0,30 | 0,49 | 7 | C.BlkS | 39,29 | 40,99 | NA | NA | 137_137 |
| apis_172 | 0,13 | 0,07 | 0,01 | 0,07 | 0,72 | 7 | C.BlkS | 39,37 | 40,13 | NA | NA | 137_137 |
| apis_177 | 0,03 | 0,11 | 0,09 | 0,36 | 0,42 | 7 | C.BlkS | 39,64 | 40,73 | NA | NA | 136_136 |
| apis_173 | 0,01 | 0,28 | 0,02 | 0,21 | 0,47 | 7 | C.BlkS | 39,67 | 40,41 | NA | NA | 137_137 |
| apis_178 | 0,19 | 0,46 | 0,01 | 0,02 | 0,33 | 7 | C.BlkS | 39,90 | 40,90 | NA | NA | 137_137 |
| apis_181 | 0,01 | 0,50 | 0,01 | 0,29 | 0,18 | 7 | C.BlkS | 39,97 | 40,16 | NA | NA | 136_136 |
| apis_180 | 0,02 | 0,36 | 0,02 | 0,35 | 0,25 | 7 | C.BlkS | 40,27 | 40,64 | NA | NA | 136_136 |
| apis_179 | 0,01 | 0,71 | 0,01 | 0,11 | 0,16 | 7 | C.BlkS | 40,27 | 40,91 | NA | NA | 136_136 |
| apis_182 | 0,20 | 0,06 | 0,14 | 0,15 | 0,45 | 7 | C.BlkS | 40,42 | 40,34 | NA | NA | 137_137 |
| apis_184 | 0,01 | 0,86 | 0,01 | 0,05 | 0,07 | 7 | C.BlkS | 40,55 | 40,81 | NA | NA | 136_136 |
| apis_183 | 0,14 | 0,02 | 0,01 | 0,04 | 0,79 | 7 | C.BlkS | 40,90 | 40,46 | 132_135 | 219_223 | 133_137 |
| apis_185 | 0,01 | 0,78 | 0,01 | 0,06 | 0,13 | 8 | L.Cauc | 40,70 | 40,96 | NA | NA | 136_136 |
| apis_186 | 0,01 | 0,86 | 0,02 | 0,04 | 0,06 | 8 | L.Cauc | 40,83 | 41,16 | 128_130 | 219_222 | 137_137 |
| apis_187 | 0,01 | 0,79 | 0,01 | 0,02 | 0,17 | 8 | L.Cauc | 41,05 | 41,02 | 142_142 | 223_223 | 137_137 |
| apis_188 | 0,03 | 0,69 | 0,01 | 0,06 | 0,21 | 8 | L.Cauc | 41,20 | 41,20 | 107_120 | 219_219 | 133_137 |
| apis_198 | 0,01 | 0,89 | 0,01 | 0,02 | 0,07 | 8 | L.Cauc | 41,36 | 40,88 | 130_142 | 219_219 | 137_137 |
| apis_190 | 0,01 | 0,23 | 0,00 | 0,60 | 0,16 | 8 | L.Cauc | 41,42 | 41,34 | NA | NA | 136_136 |
| apis_189 | 0,00 | 0,98 | 0,00 | 0,01 | 0,01 | 8 | L.Cauc | 41,56 | 41,51 | NA | NA | 136_136 |
| apis_197 | 0,02 | 0,69 | 0,01 | 0,10 | 0,19 | 8 | L.Cauc | 41,73 | 40,77 | NA | NA | 137_137 |
| apis_191 | 0,02 | 0,82 | 0,04 | 0,07 | 0,05 | 8 | L.Cauc | 41,74 | 41,28 | 128_150 | 219_219 | 137_137 |
| apis_196 | 0,01 | 0,91 | 0,01 | 0,04 | 0,03 | 8 | L.Cauc | 41,77 | 41,04 | 130_140 | 217_219 | 137_137 |
| apis_R168 | 0,01 | 0,94 | 0,00 | 0,01 | 0,03 | 8 | L.Cauc | 41,79 | 41,17 | 126_144 | 219_219 | 137_137 |
| apis_R167 | 0,01 | 0,92 | 0,01 | 0,01 | 0,04 | 8 | L.Cauc | 41,80 | 41,16 | NA | NA | 131_137 |
| apis_R169 | 0,01 | 0,95 | 0,01 | 0,01 | 0,02 | 8 | L.Cauc | 41,80 | 41,17 | 130_150 | 219_222 | 137_137 |
| apis_R171 | 0,00 | 0,97 | 0,00 | 0,01 | 0,01 | 8 | L.Cauc | 41,80 | 41,18 | 107_134 | 219_219 | 137_140 |
| apis_R170 | 0,01 | 0,96 | 0,00 | 0,01 | 0,01 | 8 | L.Cauc | 41,81 | 41,17 | 130_144 | 219_219 | 137_137 |
| apis_206 | 0,01 | 0,95 | 0,01 | 0,01 | 0,03 | 8 | L.Cauc | 41,90 | 41,48 | 132_132 | 219_222 | 137_137 |
| apis_R142 | 0,01 | 0,96 | 0,01 | 0,01 | 0,02 | 8 | L.Cauc | 41,90 | 41,47 | NA | NA | 136_136 |
| apis_R143 | 0,01 | 0,91 | 0,02 | 0,03 | 0,03 | 8 | L.Cauc | 41,90 | 41,49 | 107_122 | 223_223 | 132_137 |
| apis_R141 | 0,01 | 0,97 | 0,01 | 0,01 | 0,02 | 8 | L.Cauc | 41,91 | 41,48 | 138_168 | 222_223 | 137_140 |
| apis_192 | 0,00 | 0,92 | 0,01 | 0,03 | 0,04 | 8 | L.Cauc | 42,09 | 41,25 | 124_132 | 217_219 | 137_137 |
| apis_Geo07 | 0,00 | 0,98 | 0,00 | 0,00 | 0,01 | 8 | L.Cauc | 42,14 | 41,25 | 142_162 | 219_223 | 127_137 |
| apis_Geo08 | 0,00 | 0,97 | 0,01 | 0,01 | 0,02 | 8 | L.Cauc | 42,14 | 42,08 | 107_130 | 219_219 | 137_137 |
| apis_R096 | 0,01 | 0,89 | 0,02 | 0,03 | 0,04 | 8 | L.Cauc | 42,14 | 42,09 | NA | NA | 136_136 |
| apis_Geo02 | 0,00 | 0,98 | 0,00 | 0,00 | 0,01 | 8 | L.Cauc | 42,15 | 42,07 | 130_140 | 217_223 | 137_137 |
| apis_Geo03 | 0,00 | 0,98 | 0,00 | 0,00 | 0,01 | 8 | L.Cauc | 42,15 | 42,09 | 140_144 | 219_219 | 137_140 |
| apis_Geo05 | 0,00 | 0,99 | 0,00 | 0,00 | 0,01 | 8 | L.Cauc | 42,16 | 42,09 | 132_132 | 219_224 | 127_140 |
| apis_Geo06 | 0,01 | 0,96 | 0,00 | 0,01 | 0,02 | 8 | L.Cauc | 42,16 | 42,07 | NA | NA | 133_137 |
| apis_Geo04 | 0,00 | 0,97 | 0,00 | 0,01 | 0,02 | 8 | L.Cauc | 42,16 | 42,08 | NA | NA | 136_136 |
| apis_Geo11 | 0,01 | 0,98 | 0,00 | 0,00 | 0,01 | 8 | L.Cauc | 42,17 | 42,07 | 142_150 | 219_219 | 137_137 |
| apis_Geo12 | 0,01 | 0,97 | 0,00 | 0,01 | 0,01 | 8 | L.Cauc | 42,17 | 42,09 | NA | NA | 136_136 |
| apis_195 | 0,01 | 0,92 | 0,01 | 0,04 | 0,02 | 8 | L.Cauc | 42,22 | 41,04 | NA | NA | 136_136 |
| apis_193 | 0,01 | 0,67 | 0,01 | 0,24 | 0,07 | 8 | L.Cauc | 42,27 | 41,42 | NA | NA | 136_136 |
| apis_R097 | 0,01 | 0,96 | 0,01 | 0,01 | 0,01 | 8 | L.Cauc | 42,35 | 41,25 | NA | NA | 137_137 |

| | | | | | | | | | | | | |
|---|---|---|---|---|---|---|---|---|---|---|---|---|
| apis_194 | 0,01 | 0,63 | 0,01 | 0,31 | 0,03 | 8 | L.Cauc | 42,39 | 41,28 | NA | NA | 131_137 |
| apis_237 | 0,01 | 0,89 | 0,05 | 0,03 | 0,02 | 8 | L.Cauc | 42,61 | 41,44 | NA | NA | 136_136 |
| apis_R092 | 0,01 | 0,97 | 0,00 | 0,00 | 0,01 | 8 | L.Cauc | 42,72 | 41,51 | NA | NA | 136_136 |
| apis_R093 | 0,01 | 0,97 | 0,00 | 0,01 | 0,01 | 8 | L.Cauc | 42,73 | 41,51 | NA | NA | 133_137 |
| apis_R136 | 0,00 | 0,97 | 0,00 | 0,01 | 0,01 | 8 | L.Cauc | 42,79 | 41,52 | NA | NA | 136_136 |
| apis_236 | 0,00 | 0,97 | 0,00 | 0,01 | 0,02 | 8 | L.Cauc | 42,79 | 41,58 | NA | NA | 137_137 |
| apis_R135 | 0,01 | 0,88 | 0,01 | 0,04 | 0,06 | 8 | L.Cauc | 42,81 | 41,48 | NA | NA | 136_136 |
| apis_R094 | 0,01 | 0,92 | 0,01 | 0,02 | 0,05 | 8 | L.Cauc | 42,82 | 41,58 | NA | NA | 137_137 |
| apis_R095 | 0,01 | 0,88 | 0,05 | 0,02 | 0,05 | 8 | L.Cauc | 42,83 | 41,58 | NA | NA | 137_137 |
| apis_R199 | 0,00 | 0,90 | 0,07 | 0,01 | 0,02 | 8 | L.Cauc | 42,87 | 41,48 | NA | NA | 132_137 |
| apis_238 | 0,01 | 0,92 | 0,01 | 0,01 | 0,05 | 8 | L.Cauc | 42,91 | 41,45 | NA | NA | 137_137 |
| apis_199 | 0,05 | 0,04 | 0,16 | 0,11 | 0,63 | 9 | Erz-Kar | 40,81 | 39,89 | NA | NA | 136_136 |
| apis_229 | 0,03 | 0,05 | 0,01 | 0,72 | 0,20 | 9 | Erz-Kar | 41,03 | 39,65 | NA | NA | 136_136 |
| apis_232 | 0,01 | 0,08 | 0,22 | 0,30 | 0,39 | 9 | Erz-Kar | 41,44 | 40,33 | NA | NA | 137_137 |
| apis_230 | 0,02 | 0,02 | 0,19 | 0,64 | 0,12 | 9 | Erz-Kar | 41,53 | 39,93 | NA | NA | 137_137 |
| apis_231 | 0,01 | 0,22 | 0,06 | 0,36 | 0,34 | 9 | Erz-Kar | 41,86 | 40,09 | NA | NA | 137_137 |
| apis_233 | 0,43 | 0,03 | 0,03 | 0,42 | 0,10 | 9 | Erz-Kar | 42,16 | 40,59 | NA | NA | 136_136 |
| apis_251 | 0,03 | 0,06 | 0,01 | 0,83 | 0,07 | 9 | Erz-Kar | 42,40 | 40,23 | NA | NA | 137_137 |
| apis_235 | 0,01 | 0,12 | 0,02 | 0,76 | 0,10 | 9 | Erz-Kar | 42,48 | 40,91 | NA | NA | 136_136 |
| apis_234 | 0,03 | 0,12 | 0,04 | 0,26 | 0,56 | 9 | Erz-Kar | 42,51 | 40,68 | NA | NA | 137_137 |
| apis_R198 | 0,00 | 0,95 | 0,02 | 0,01 | 0,02 | 9 | Erz-Kar | 42,58 | 41,06 | NA | NA | 137_137 |
| apis_R134 | 0,01 | 0,84 | 0,01 | 0,08 | 0,06 | 9 | Erz-Kar | 42,65 | 41,10 | NA | NA | 137_137 |
| apis_R177 | 0,00 | 0,96 | 0,01 | 0,01 | 0,01 | 9 | Erz-Kar | 42,67 | 41,12 | NA | NA | 137_137 |
| apis_R173 | 0,01 | 0,93 | 0,03 | 0,02 | 0,02 | 9 | Erz-Kar | 42,68 | 41,11 | NA | NA | 136_136 |
| apis_R174 | 0,01 | 0,69 | 0,04 | 0,07 | 0,19 | 9 | Erz-Kar | 42,68 | 41,12 | NA | NA | 137_137 |
| apis_R179 | 0,00 | 0,98 | 0,00 | 0,01 | 0,01 | 9 | Erz-Kar | 42,68 | 41,13 | NA | NA | 137_137 |
| apis_R175 | 0,01 | 0,96 | 0,00 | 0,01 | 0,02 | 9 | Erz-Kar | 42,69 | 41,10 | NA | NA | 136_136 |
| apis_R176 | 0,01 | 0,96 | 0,00 | 0,01 | 0,02 | 9 | Erz-Kar | 42,69 | 41,11 | NA | NA | 132_137 |
| apis_R178 | 0,01 | 0,40 | 0,01 | 0,03 | 0,56 | 9 | Erz-Kar | 42,69 | 41,12 | NA | NA | 137_137 |
| apis_R181 | 0,00 | 0,97 | 0,01 | 0,01 | 0,01 | 9 | Erz-Kar | 42,69 | 41,13 | NA | NA | 136_136 |
| apis_R183 | 0,01 | 0,92 | 0,01 | 0,01 | 0,05 | 9 | Erz-Kar | 42,69 | 41,14 | NA | NA | 137_137 |
| apis_R140 | 0,02 | 0,32 | 0,01 | 0,01 | 0,64 | 9 | Erz-Kar | 42,70 | 41,11 | NA | NA | 137_137 |
| apis_R180 | 0,01 | 0,91 | 0,01 | 0,02 | 0,06 | 9 | Erz-Kar | 42,70 | 41,12 | NA | NA | 137_137 |
| apis_R182 | 0,01 | 0,94 | 0,00 | 0,01 | 0,04 | 9 | Erz-Kar | 42,70 | 41,13 | NA | NA | 137_137 |
| apis_R196 | 0,01 | 0,82 | 0,01 | 0,07 | 0,10 | 9 | Erz-Kar | 42,70 | 41,14 | NA | NA | 137_137 |
| apis_R172 | 0,01 | 0,94 | 0,00 | 0,02 | 0,04 | 9 | Erz-Kar | 42,71 | 41,12 | NA | NA | 137_137 |
| apis_R184 | 0,01 | 0,97 | 0,00 | 0,00 | 0,01 | 9 | Erz-Kar | 42,71 | 41,13 | NA | NA | 137_137 |
| apis_246 | 0,03 | 0,02 | 0,05 | 0,48 | 0,42 | 9 | Erz-Kar | 42,73 | 40,83 | NA | NA | 137_137 |
| apis_R202 | 0,01 | 0,90 | 0,01 | 0,01 | 0,07 | 9 | Erz-Kar | 42,73 | 41,22 | NA | NA | 137_137 |
| apis_252 | 0,02 | 0,45 | 0,03 | 0,07 | 0,43 | 9 | Erz-Kar | 42,76 | 40,05 | NA | NA | 137_137 |
| apis_R137 | 0,07 | 0,89 | 0,01 | 0,01 | 0,03 | 9 | Erz-Kar | 42,77 | 41,25 | NA | NA | 136_136 |
| apis_R203 | 0,02 | 0,93 | 0,01 | 0,01 | 0,02 | 9 | Erz-Kar | 42,79 | 41,24 | NA | NA | 136_136 |
| apis_240 | 0,01 | 0,55 | 0,01 | 0,33 | 0,11 | 9 | Erz-Kar | 42,80 | 41,34 | NA | NA | 137_137 |
| apis_239 | 0,08 | 0,85 | 0,01 | 0,01 | 0,04 | 9 | Erz-Kar | 42,80 | 41,32 | NA | NA | 137_137 |
| apis_R197 | 0,01 | 0,51 | 0,03 | 0,10 | 0,35 | 9 | Erz-Kar | 42,84 | 41,15 | NA | NA | 136_136 |
| apis_R200 | 0,01 | 0,10 | 0,01 | 0,18 | 0,69 | 9 | Erz-Kar | 42,86 | 41,23 | NA | NA | 137_137 |
| apis_R201 | 0,01 | 0,64 | 0,28 | 0,05 | 0,02 | 9 | Erz-Kar | 42,87 | 41,23 | NA | NA | 137_137 |
| apis_250 | 0,02 | 0,02 | 0,01 | 0,08 | 0,86 | 9 | Erz-Kar | 42,88 | 40,55 | NA | NA | 137_137 |

| | | | | | | | | | | | | |
|---|---|---|---|---|---|---|---|---|---|---|---|---|
| apis_R138 | 0,01 | 0,91 | 0,01 | 0,02 | 0,06 | 9 | Erz-Kar | 42,96 | 41,14 | NA | NA | 137_137 |
| apis_R139 | 0,01 | 0,94 | 0,01 | 0,02 | 0,02 | 9 | Erz-Kar | 42,96 | 41,16 | NA | NA | 137_137 |
| apis_245 | 0,00 | 0,80 | 0,01 | 0,02 | 0,17 | 9 | Erz-Kar | 42,97 | 40,98 | NA | NA | 137_137 |
| apis_244 | 0,01 | 0,96 | 0,00 | 0,01 | 0,02 | 9 | Erz-Kar | 43,12 | 41,07 | NA | NA | 132_136 |
| apis_255 | 0,01 | 0,02 | 0,22 | 0,14 | 0,62 | 9 | Erz-Kar | 43,13 | 40,39 | NA | NA | 137_137 |
| apis_253 | 0,01 | 0,69 | 0,05 | 0,19 | 0,06 | 9 | Erz-Kar | 43,14 | 40,12 | 140_146 | 223_223 | 137_137 |
| apis_241 | 0,00 | 0,93 | 0,00 | 0,01 | 0,05 | 9 | Erz-Kar | 43,15 | 41,28 | 107_160 | 219_219 | 137_137 |
| apis_243 | 0,01 | 0,63 | 0,02 | 0,11 | 0,23 | 9 | Erz-Kar | 43,29 | 40,78 | NA | NA | 137_137 |
| apis_247 | 0,02 | 0,72 | 0,01 | 0,20 | 0,05 | 9 | Erz-Kar | 43,29 | 41,14 | NA | NA | 137_137 |
| apis_242 | 0,01 | 0,88 | 0,00 | 0,04 | 0,07 | 9 | Erz-Kar | 43,33 | 41,14 | NA | NA | 133_137 |
| apis_254 | 0,10 | 0,18 | 0,14 | 0,23 | 0,35 | 9 | Erz-Kar | 43,53 | 40,55 | NA | NA | 137_137 |
| apis_256 | 0,12 | 0,03 | 0,08 | 0,71 | 0,06 | 9 | Erz-Kar | 43,59 | 40,28 | NA | NA | 137_137 |
| apis_248 | 0,01 | 0,03 | 0,03 | 0,48 | 0,44 | 9 | Erz-Kar | 43,62 | 40,86 | NA | NA | 137_137 |
| apis_249 | 0,01 | 0,17 | 0,09 | 0,68 | 0,05 | 9 | Erz-Kar | 43,72 | 40,72 | NA | NA | 136_136 |
| apis_203 | 0,18 | 0,01 | 0,07 | 0,32 | 0,42 | 10 | U.Euph | 38,54 | 39,30 | 122_154 | 217_234 | 132_137 |
| apis_122 | 0,03 | 0,91 | 0,02 | 0,01 | 0,04 | 10 | U.Euph | 38,69 | 38,91 | 128_135 | 219_223 | 133_137 |
| apis_338A | 0,12 | 0,36 | 0,01 | 0,12 | 0,40 | 10 | U.Euph | 38,70 | 38,59 | 134_152 | 219_223 | 137_137 |
| apis_338B | 0,01 | 0,00 | 0,10 | 0,03 | 0,86 | 10 | U.Euph | 38,71 | 38,59 | 120_134 | 219_219 | 137_137 |
| apis_204 | 0,13 | 0,35 | 0,02 | 0,32 | 0,17 | 10 | U.Euph | 38,85 | 39,83 | 140_154 | 223_223 | 137_137 |
| apis_202 | 0,01 | 0,03 | 0,01 | 0,04 | 0,91 | 10 | U.Euph | 38,96 | 39,60 | NA | NA | 137_137 |
| apis_343A | 0,56 | 0,03 | 0,03 | 0,12 | 0,27 | 10 | U.Euph | 39,09 | 39,34 | NA | NA | 136_136 |
| apis_343C | 0,01 | 0,01 | 0,02 | 0,30 | 0,66 | 10 | U.Euph | 39,09 | 39,35 | 138_174 | 223_223 | 137_137 |
| apis_343B | 0,36 | 0,33 | 0,05 | 0,07 | 0,19 | 10 | U.Euph | 39,10 | 39,34 | 120_166 | 223_223 | 137_137 |
| apis_339A | 0,01 | 0,68 | 0,01 | 0,01 | 0,29 | 10 | U.Euph | 39,13 | 38,76 | 126_138 | 219_223 | 131_137 |
| apis_339B | 0,01 | 0,44 | 0,08 | 0,08 | 0,40 | 10 | U.Euph | 39,14 | 38,76 | NA | NA | 133_137 |
| apis_345A | 0,01 | 0,07 | 0,05 | 0,64 | 0,22 | 10 | U.Euph | 39,21 | 39,16 | 134_140 | 217_223 | 133_137 |
| apis_345B | 0,01 | 0,04 | 0,01 | 0,02 | 0,92 | 10 | U.Euph | 39,22 | 39,16 | 135_158 | 219_225 | NA |
| apis_344A | 0,03 | 0,01 | 0,01 | 0,05 | 0,90 | 10 | U.Euph | 39,29 | 39,10 | 111_126 | 219_223 | 133_137 |
| apis_344B | 0,19 | 0,24 | 0,01 | 0,39 | 0,18 | 10 | U.Euph | 39,30 | 39,10 | NA | NA | 131_137 |
| apis_346A | 0,04 | 0,01 | 0,01 | 0,77 | 0,16 | 10 | U.Euph | 39,30 | 38,37 | 126_132 | 223_223 | 137_137 |
| apis_346B | 0,02 | 0,01 | 0,06 | 0,29 | 0,63 | 10 | U.Euph | 39,30 | 38,38 | 107_135 | 219_223 | 137_137 |
| apis_340A | 0,01 | 0,01 | 0,06 | 0,55 | 0,37 | 10 | U.Euph | 39,48 | 38,99 | NA | NA | 137_137 |
| apis_340B | 0,01 | 0,04 | 0,09 | 0,15 | 0,72 | 10 | U.Euph | 39,49 | 38,99 | NA | NA | 137_137 |
| apis_201 | 0,01 | 0,66 | 0,11 | 0,14 | 0,08 | 10 | U.Euph | 39,66 | 39,72 | NA | NA | 137_137 |
| apis_341A | 0,01 | 0,02 | 0,01 | 0,61 | 0,36 | 10 | U.Euph | 39,83 | 39,18 | NA | NA | 133_137 |
| apis_341B | 0,02 | 0,02 | 0,03 | 0,43 | 0,50 | 10 | U.Euph | 39,83 | 39,19 | NA | NA | 137_137 |
| apis_342A | 0,01 | 0,09 | 0,35 | 0,35 | 0,20 | 10 | U.Euph | 39,86 | 39,45 | 135_166 | 223_223 | 137_137 |
| apis_342B | 0,02 | 0,02 | 0,01 | 0,90 | 0,05 | 10 | U.Euph | 39,87 | 39,45 | 135_160 | 217_219 | 133_137 |
| apis_347A | 0,01 | 0,02 | 0,04 | 0,27 | 0,67 | 10 | U.Euph | 39,89 | 38,54 | 122_130 | 219_219 | 133_137 |
| apis_347B | 0,12 | 0,01 | 0,03 | 0,04 | 0,79 | 10 | U.Euph | 39,89 | 38,55 | 135_135 | 219_223 | 132_137 |
| apis_349A | 0,02 | 0,26 | 0,01 | 0,23 | 0,48 | 10 | U.Euph | 40,12 | 39,09 | 126_132 | 219_219 | 137_137 |
| apis_349B | 0,03 | 0,03 | 0,17 | 0,73 | 0,04 | 10 | U.Euph | 40,13 | 39,09 | 124_132 | 219_223 | NA |
| apis_348A | 0,01 | 0,01 | 0,19 | 0,70 | 0,09 | 10 | U.Euph | 40,18 | 38,51 | NA | NA | 137_137 |
| apis_348B | 0,01 | 0,08 | 0,19 | 0,68 | 0,04 | 10 | U.Euph | 40,19 | 38,51 | NA | NA | 133_137 |
| apis_200 | 0,13 | 0,01 | 0,04 | 0,04 | 0,78 | 10 | U.Euph | 40,24 | 39,78 | NA | NA | 137_137 |
| apis_354A | 0,03 | 0,02 | 0,05 | 0,78 | 0,12 | 10 | U.Euph | 40,34 | 38,94 | NA | NA | 137_137 |
| apis_354B | 0,02 | 0,53 | 0,02 | 0,04 | 0,39 | 10 | U.Euph | 40,35 | 38,94 | 122_142 | 219_223 | 137_137 |
| apis_350B | 0,08 | 0,06 | 0,03 | 0,62 | 0,20 | 10 | U.Euph | 40,63 | 39,39 | 128_138 | 217_217 | 137_137 |

| | | | | | | | | | | | |
|---|---|---|---|---|---|---|---|---|---|---|---|
| apis_353A | 0,03 | 0,02 | 0,02 | 0,86 | 0,07 | 10 U.Euph | 40,73 | 39,03 | 120_135 | 219_223 | 137_137 |
| apis_353B | 0,01 | 0,01 | 0,01 | 0,05 | 0,92 | 10 U.Euph | 40,74 | 39,03 | 132_138 | 219_223 | 132_137 |
| apis_355A | 0,08 | 0,02 | 0,03 | 0,28 | 0,60 | 10 U.Euph | 40,75 | 38,74 | NA | NA | 137_137 |
| apis_355B | 0,04 | 0,03 | 0,23 | 0,51 | 0,18 | 10 U.Euph | 40,75 | 38,75 | NA | NA | 133_137 |
| apis_356A | 0,02 | 0,44 | 0,09 | 0,12 | 0,32 | 10 U.Euph | 40,80 | 38,99 | NA | NA | 137_137 |
| apis_356B | 0,05 | 0,02 | 0,10 | 0,63 | 0,19 | 10 U.Euph | 40,81 | 38,99 | NA | NA | 137_137 |
| apis_352A | 0,01 | 0,12 | 0,18 | 0,59 | 0,09 | 10 U.Euph | 40,81 | 39,07 | NA | NA | 137_137 |
| apis_352B | 0,01 | 0,02 | 0,12 | 0,81 | 0,04 | 10 U.Euph | 40,82 | 39,07 | NA | NA | 137_137 |
| apis_357A | 0,01 | 0,03 | 0,01 | 0,14 | 0,81 | 10 U.Euph | 41,06 | 38,97 | NA | NA | 137_137 |
| apis_357B | 0,01 | 0,01 | 0,01 | 0,08 | 0,90 | 10 U.Euph | 41,06 | 38,98 | NA | NA | 137_137 |
| apis_351A | 0,01 | 0,05 | 0,01 | 0,89 | 0,04 | 10 U.Euph | 41,13 | 39,20 | NA | NA | 133_137 |
| apis_351B | 0,01 | 0,85 | 0,01 | 0,03 | 0,09 | 10 U.Euph | 41,14 | 39,20 | NA | NA | 137_137 |
| apis_358A | 0,02 | 0,00 | 0,01 | 0,05 | 0,92 | 10 U.Euph | 41,68 | 39,05 | NA | NA | 137_137 |
| apis_358B | 0,02 | 0,01 | 0,01 | 0,03 | 0,94 | 10 U.Euph | 41,69 | 39,06 | NA | NA | 137_137 |
| apis_263 | 0,01 | 0,09 | 0,02 | 0,10 | 0,79 | 10 U.Euph | 41,74 | 39,31 | NA | NA | 137_137 |
| apis_361A | 0,02 | 0,01 | 0,03 | 0,06 | 0,88 | 10 U.Euph | 41,96 | 38,78 | NA | NA | 137_137 |
| apis_361B | 0,02 | 0,32 | 0,03 | 0,24 | 0,39 | 10 U.Euph | 41,97 | 38,78 | NA | NA | 131_137 |
| apis_360B | 0,01 | 0,25 | 0,01 | 0,20 | 0,52 | 10 U.Euph | 42,12 | 39,00 | NA | NA | 137_137 |
| apis_262 | 0,16 | 0,41 | 0,02 | 0,28 | 0,13 | 10 U.Euph | 42,33 | 39,64 | NA | NA | 137_137 |
| apis_362A | 0,01 | 0,03 | 0,04 | 0,58 | 0,34 | 10 U.Euph | 42,55 | 39,81 | NA | NA | 137_137 |
| apis_362B | 0,01 | 0,71 | 0,05 | 0,02 | 0,20 | 10 U.Euph | 42,56 | 39,81 | NA | NA | 137_137 |
| apis_365A | 0,00 | 0,16 | 0,05 | 0,20 | 0,59 | 10 U.Euph | 42,70 | 39,22 | NA | NA | 131_137 |
| apis_365B | 0,01 | 0,05 | 0,02 | 0,11 | 0,81 | 10 U.Euph | 42,71 | 39,22 | NA | NA | 137_137 |
| apis_363A | 0,01 | 0,82 | 0,02 | 0,04 | 0,11 | 10 U.Euph | 42,73 | 39,64 | NA | NA | 137_137 |
| apis_363B | 0,74 | 0,17 | 0,01 | 0,01 | 0,07 | 10 U.Euph | 42,74 | 39,64 | 122_128 | 223_223 | 137_137 |
| apis_364A | 0,16 | 0,10 | 0,00 | 0,33 | 0,40 | 10 U.Euph | 42,75 | 39,64 | 102_180 | 219_219 | 137_137 |
| apis_364B | 0,05 | 0,02 | 0,04 | 0,10 | 0,80 | 10 U.Euph | 42,76 | 39,64 | NA | NA | 137_137 |
| apis_366A | 0,87 | 0,01 | 0,03 | 0,04 | 0,05 | 10 U.Euph | 43,12 | 39,94 | NA | NA | 137_137 |
| apis_366B | 0,03 | 0,03 | 0,03 | 0,85 | 0,06 | 10 U.Euph | 43,13 | 39,94 | NA | NA | 133_137 |
| apis_367A | 0,02 | 0,01 | 0,01 | 0,61 | 0,35 | 10 U.Euph | 43,37 | 39,66 | 120_135 | 223_223 | 137_142 |
| apis_367B | 0,01 | 0,01 | 0,01 | 0,90 | 0,08 | 10 U.Euph | 43,38 | 39,66 | 130_146 | 223_223 | 137_137 |
| apis_368A | 0,01 | 0,29 | 0,01 | 0,10 | 0,59 | 10 U.Euph | 43,57 | 39,61 | NA | NA | 137_137 |
| apis_368B | 0,01 | 0,69 | 0,01 | 0,07 | 0,21 | 10 U.Euph | 43,58 | 39,61 | NA | NA | 137_137 |
| apis_257 | 0,02 | 0,02 | 0,03 | 0,54 | 0,40 | 10 U.Euph | 43,66 | 39,96 | 120_142 | 223_223 | 137_137 |
| apis_261 | 0,60 | 0,04 | 0,01 | 0,04 | 0,30 | 10 U.Euph | 43,93 | 39,82 | 118_148 | 219_223 | 137_137 |
| apis_369A | 0,00 | 0,06 | 0,02 | 0,85 | 0,07 | 10 U.Euph | 44,08 | 39,54 | 118_148 | 219_223 | 137_137 |
| apis_369B | 0,01 | 0,03 | 0,06 | 0,20 | 0,70 | 10 U.Euph | 44,09 | 39,54 | 135_142 | 223_223 | 137_137 |
| apis_260 | 0,04 | 0,50 | 0,04 | 0,39 | 0,03 | 10 U.Euph | 44,29 | 40,03 | 120_148 | 219_223 | 137_137 |
| apis_259 | 0,02 | 0,33 | 0,07 | 0,03 | 0,55 | 10 U.Euph | 44,38 | 39,78 | NA | NA | 137_137 |
| apis_258 | 0,02 | 0,08 | 0,41 | 0,41 | 0,07 | 10 U.Euph | 44,59 | 39,82 | 138_138 | 223_223 | 137_137 |
| apis_329 | 0,02 | 0,03 | 0,01 | 0,54 | 0,40 | 11 EC.Anat | 33,86 | 37,79 | 120_144 | 219_223 | 137_137 |
| apis_331 | 0,03 | 0,04 | 0,25 | 0,16 | 0,52 | 11 EC.Anat | 34,21 | 38,50 | 122_132 | 223_223 | 137_137 |
| apis_328 | 0,05 | 0,01 | 0,07 | 0,71 | 0,17 | 11 EC.Anat | 34,31 | 37,41 | 144_152 | 223_223 | 137_137 |
| apis_330 | 0,04 | 0,01 | 0,23 | 0,51 | 0,22 | 11 EC.Anat | 34,63 | 37,96 | 128_140 | 223_225 | 137_137 |
| apis_145 | 0,22 | 0,10 | 0,01 | 0,09 | 0,58 | 11 EC.Anat | 34,83 | 38,68 | 126_148 | 223_223 | 137_137 |
| apis_144 | 0,02 | 0,01 | 0,05 | 0,59 | 0,33 | 11 EC.Anat | 35,01 | 38,34 | NA | NA | 133_137 |
| apis_291 | 0,02 | 0,15 | 0,01 | 0,22 | 0,60 | 11 EC.Anat | 35,13 | 39,62 | 126_150 | 223_223 | 137_137 |
| apis_143 | 0,10 | 0,05 | 0,44 | 0,17 | 0,23 | 11 EC.Anat | 35,29 | 38,08 | 132_144 | 219_223 | 137_137 |

| | | | | | | | | | | | | |
|---|---|---|---|---|---|---|---|---|---|---|---|---|
| apis_268 | 0,03 | 0,67 | 0,02 | 0,18 | 0,10 | 11 | EC.Anat | 35,31 | 40,06 | 154_154 | 219_223 | 137_137 |
| apis_141 | 0,02 | 0,43 | 0,33 | 0,06 | 0,16 | 11 | EC.Anat | 35,32 | 38,86 | 130_156 | 223_223 | 137_137 |
| apis_142 | 0,01 | 0,13 | 0,03 | 0,66 | 0,16 | 11 | EC.Anat | 35,63 | 38,44 | NA | NA | 137_137 |
| apis_140 | 0,13 | 0,03 | 0,23 | 0,21 | 0,40 | 11 | EC.Anat | 35,79 | 39,21 | NA | NA | 133_137 |
| apis_137 | 0,03 | 0,02 | 0,03 | 0,19 | 0,74 | 11 | EC.Anat | 36,12 | 38,30 | 140_140 | 219_223 | 137_137 |
| apis_138 | 0,04 | 0,01 | 0,41 | 0,30 | 0,24 | 11 | EC.Anat | 36,42 | 38,95 | NA | NA | 137_137 |
| apis_139 | 0,02 | 0,23 | 0,01 | 0,31 | 0,43 | 11 | EC.Anat | 36,57 | 39,52 | 120_146 | 223_223 | 137_137 |
| apis_266 | 0,03 | 0,11 | 0,04 | 0,73 | 0,09 | 11 | EC.Anat | 36,79 | 39,99 | 142_162 | 223_225 | 137_137 |
| apis_127 | 0,02 | 0,02 | 0,86 | 0,08 | 0,03 | 11 | EC.Anat | 37,02 | 38,43 | 150_162 | 219_223 | 133_137 |
| apis_125 | 0,02 | 0,02 | 0,04 | 0,71 | 0,21 | 11 | EC.Anat | 37,10 | 39,34 | 124_135 | 223_223 | 137_137 |
| apis_126 | 0,02 | 0,07 | 0,48 | 0,34 | 0,09 | 11 | EC.Anat | 37,30 | 38,91 | 120_157 | 223_223 | 137_137 |
| apis_264 | 0,51 | 0,22 | 0,01 | 0,18 | 0,09 | 11 | EC.Anat | 37,31 | 39,92 | 135_142 | 223_225 | 137_137 |
| apis_128 | 0,08 | 0,01 | 0,36 | 0,14 | 0,41 | 11 | EC.Anat | 37,34 | 38,16 | 135_142 | 223_225 | 137_137 |
| apis_265 | 0,04 | 0,04 | 0,02 | 0,36 | 0,54 | 11 | EC.Anat | 37,53 | 39,61 | NA | NA | 137_137 |
| apis_129 | 0,03 | 0,01 | 0,15 | 0,73 | 0,08 | 11 | EC.Anat | 37,74 | 38,43 | 142_162 | 223_225 | 137_137 |
| apis_131 | 0,02 | 0,07 | 0,03 | 0,77 | 0,12 | 11 | EC.Anat | 37,87 | 38,10 | 134_150 | 223_223 | 137_137 |
| apis_124 | 0,09 | 0,06 | 0,52 | 0,12 | 0,22 | 11 | EC.Anat | 37,90 | 39,00 | 134_142 | 223_223 | 137_137 |
| apis_205 | 0,12 | 0,04 | 0,09 | 0,11 | 0,65 | 11 | EC.Anat | 38,00 | 39,91 | NA | NA | 137_137 |
| apis_121 | 0,02 | 0,02 | 0,01 | 0,07 | 0,88 | 11 | EC.Anat | 38,03 | 39,47 | 135_142 | 223_225 | 137_137 |
| apis_123 | 0,01 | 0,84 | 0,03 | 0,05 | 0,07 | 11 | EC.Anat | 38,18 | 38,60 | 132_162 | 223_223 | 137_137 |
| apis_130 | 0,01 | 0,04 | 0,12 | 0,69 | 0,15 | 11 | EC.Anat | 38,52 | 38,29 | 130_135 | 223_223 | 137_137 |
| apis_215 | 0,02 | 0,01 | 0,72 | 0,21 | 0,04 | 12 | E.Med | 33,90 | 36,47 | NA | NA | 137_137 |
| apis_214 | 0,02 | 0,03 | 0,62 | 0,26 | 0,07 | 12 | E.Med | 34,10 | 36,88 | NA | NA | 137_137 |
| apis_213 | 0,08 | 0,01 | 0,02 | 0,45 | 0,43 | 12 | E.Med | 34,25 | 36,70 | 144_154 | 219_223 | 137_137 |
| apis_211 | 0,03 | 0,07 | 0,03 | 0,65 | 0,22 | 12 | E.Med | 34,59 | 37,11 | 135_158 | 223_223 | 137_137 |
| apis_207 | 0,02 | 0,28 | 0,05 | 0,39 | 0,26 | 12 | E.Med | 34,89 | 37,40 | NA | NA | 137_137 |
| apis_212 | 0,01 | 0,12 | 0,01 | 0,49 | 0,37 | 12 | E.Med | 35,00 | 36,87 | 124_191 | 219_223 | 137_137 |
| apis_208 | 0,01 | 0,05 | 0,76 | 0,09 | 0,09 | 12 | E.Med | 35,30 | 37,59 | NA | NA | 137_137 |
| apis_210 | 0,03 | 0,02 | 0,11 | 0,29 | 0,55 | 12 | E.Med | 35,45 | 36,69 | 122_122 | 219_223 | NA |
| apis_209 | 0,01 | 0,09 | 0,52 | 0,28 | 0,10 | 12 | E.Med | 35,51 | 37,15 | 140_144 | 219_223 | NA |
| apis_032 | 0,06 | 0,02 | 0,87 | 0,02 | 0,04 | 12 | E.Med | 35,79 | 37,54 | 132_150 | 223_223 | 137_137 |
| apis_017 | 0,04 | 0,04 | 0,70 | 0,04 | 0,17 | 12 | E.Med | 35,80 | 36,28 | 120_134 | 223_223 | 137_137 |
| apis_031 | 0,06 | 0,03 | 0,81 | 0,03 | 0,08 | 12 | E.Med | 35,88 | 37,83 | 138_172 | 223_223 | 133_137 |
| apis_016 | 0,03 | 0,01 | 0,86 | 0,05 | 0,05 | 12 | E.Med | 35,92 | 36,43 | 130_138 | 223_223 | 137_142 |
| apis_018 | 0,01 | 0,01 | 0,87 | 0,01 | 0,09 | 12 | E.Med | 35,96 | 36,14 | 132_154 | 219_223 | 137_137 |
| apis_R205 | 0,01 | 0,04 | 0,46 | 0,44 | 0,05 | 12 | E.Med | 35,98 | 36,09 | 134_182 | 219_223 | 137_137 |
| apis_015 | 0,03 | 0,05 | 0,72 | 0,04 | 0,15 | 12 | E.Med | 35,98 | 36,43 | 120_158 | 223_223 | 137_142 |
| apis_R111 | 0,01 | 0,01 | 0,88 | 0,03 | 0,07 | 12 | E.Med | 35,99 | 36,08 | 142_146 | 223_223 | 137_144 |
| apis_R243 | 0,01 | 0,32 | 0,26 | 0,36 | 0,05 | 12 | E.Med | 35,99 | 36,09 | 120_154 | 223_223 | 137_137 |
| apis_R245 | 0,05 | 0,04 | 0,82 | 0,02 | 0,07 | 12 | E.Med | 35,99 | 36,10 | 130_135 | 223_223 | 137_137 |
| apis_R244 | 0,01 | 0,05 | 0,53 | 0,07 | 0,34 | 12 | E.Med | 36,00 | 36,09 | 142_150 | 223_223 | 137_137 |
| apis_014 | 0,02 | 0,01 | 0,92 | 0,01 | 0,04 | 12 | E.Med | 36,03 | 36,97 | 140_184 | 219_223 | 133_137 |
| apis_030 | 0,04 | 0,04 | 0,66 | 0,05 | 0,21 | 12 | E.Med | 36,10 | 37,40 | 120_140 | 219_223 | 137_137 |
| apis_R113 | 0,01 | 0,02 | 0,95 | 0,01 | 0,02 | 12 | E.Med | 36,13 | 36,04 | 120_122 | 223_223 | 132_137 |
| apis_R246 | 0,02 | 0,04 | 0,91 | 0,01 | 0,01 | 12 | E.Med | 36,13 | 36,05 | 128_146 | 223_223 | 137_137 |
| apis_R249 | 0,01 | 0,01 | 0,66 | 0,29 | 0,03 | 12 | E.Med | 36,13 | 36,06 | 118_140 | 223_223 | 133_137 |
| apis_R037 | 0,01 | 0,07 | 0,83 | 0,04 | 0,05 | 12 | E.Med | 36,14 | 35,99 | 138_148 | 219_223 | 137_137 |
| apis_R114 | 0,00 | 0,01 | 0,96 | 0,01 | 0,02 | 12 | E.Med | 36,14 | 36,00 | 103_138 | 219_219 | 137_137 |

| | | | | | | | | | | | | |
|---|---|---|---|---|---|---|---|---|---|---|---|---|
| apis_R144 | 0,00 | 0,01 | 0,95 | 0,02 | 0,02 | 12 | E.Med | 36,14 | 36,04 | 120_132 | 219_223 | 137_137 |
| apis_R247 | 0,01 | 0,00 | 0,97 | 0,01 | 0,01 | 12 | E.Med | 36,14 | 36,06 | 132_138 | 223_223 | 137_137 |
| apis_019 | 0,01 | 0,01 | 0,91 | 0,02 | 0,05 | 12 | E.Med | 36,14 | 36,05 | 128_140 | 223_223 | 137_137 |
| apis_R039 | 0,01 | 0,02 | 0,52 | 0,39 | 0,06 | 12 | E.Med | 36,15 | 35,98 | 135_140 | 219_223 | 137_137 |
| apis_R043 | 0,00 | 0,00 | 0,93 | 0,02 | 0,04 | 12 | E.Med | 36,15 | 36,00 | 135_152 | 222_223 | 137_137 |
| apis_R115 | 0,01 | 0,01 | 0,93 | 0,02 | 0,04 | 12 | E.Med | 36,15 | 36,04 | 120_138 | 223_223 | 133_137 |
| apis_R148 | 0,01 | 0,01 | 0,95 | 0,01 | 0,02 | 12 | E.Med | 36,15 | 36,05 | 135_162 | 223_223 | 137_137 |
| apis_R149 | 0,01 | 0,01 | 0,95 | 0,01 | 0,02 | 12 | E.Med | 36,15 | 36,06 | 118_134 | 217_223 | NA |
| apis_R250 | 0,01 | 0,01 | 0,93 | 0,03 | 0,02 | 12 | E.Med | 36,15 | 36,07 | 120_138 | 222_223 | 137_137 |
| apis_020 | 0,01 | 0,01 | 0,96 | 0,01 | 0,01 | 12 | E.Med | 36,15 | 35,99 | 140_182 | 219_219 | 133_137 |
| apis_R038 | 0,05 | 0,01 | 0,87 | 0,02 | 0,04 | 12 | E.Med | 36,16 | 35,98 | NA | NA | 137_137 |
| apis_R040 | 0,02 | 0,01 | 0,02 | 0,08 | 0,87 | 12 | E.Med | 36,16 | 35,99 | 118_130 | 223_223 | 137_137 |
| apis_R042 | 0,05 | 0,00 | 0,94 | 0,01 | 0,01 | 12 | E.Med | 36,16 | 36,00 | 122_132 | 223_223 | 137_137 |
| apis_R248 | 0,01 | 0,18 | 0,71 | 0,03 | 0,08 | 12 | E.Med | 36,16 | 36,06 | 138_142 | 221_223 | 137_137 |
| apis_029 | 0,03 | 0,01 | 0,94 | 0,01 | 0,02 | 12 | E.Med | 36,21 | 37,20 | 132_203 | 223_223 | 137_137 |
| apis_R041 | 0,00 | 0,01 | 0,96 | 0,02 | 0,01 | 12 | E.Med | 36,21 | 35,98 | 120_132 | 223_223 | 137_137 |
| apis_021 | 0,01 | 0,01 | 0,96 | 0,01 | 0,01 | 12 | E.Med | 36,22 | 36,05 | 140_142 | 217_223 | 137_144 |
| apis_022 | 0,02 | 0,01 | 0,88 | 0,07 | 0,03 | 12 | E.Med | 36,22 | 36,40 | 135_138 | 223_223 | 137_137 |
| apis_013 | 0,17 | 0,03 | 0,49 | 0,05 | 0,26 | 12 | E.Med | 36,23 | 36,75 | 140_140 | 223_223 | 133_137 |
| apis_023 | 0,01 | 0,01 | 0,95 | 0,01 | 0,02 | 12 | E.Med | 36,39 | 36,62 | 132_132 | 223_225 | 137_137 |
| apis_135 | 0,02 | 0,01 | 0,64 | 0,28 | 0,04 | 12 | E.Med | 36,42 | 37,60 | 138_152 | 219_219 | 137_137 |
| apis_028 | 0,07 | 0,01 | 0,72 | 0,02 | 0,18 | 12 | E.Med | 36,45 | 37,36 | 135_152 | 219_223 | 137_137 |
| apis_025 | 0,01 | 0,02 | 0,94 | 0,01 | 0,03 | 12 | E.Med | 36,51 | 36,79 | 111_122 | 222_223 | 137_137 |
| apis_024 | 0,01 | 0,01 | 0,96 | 0,01 | 0,01 | 12 | E.Med | 36,59 | 36,27 | 138_176 | 219_223 | 137_137 |
| apis_136 | 0,09 | 0,31 | 0,04 | 0,18 | 0,39 | 12 | E.Med | 36,65 | 38,05 | 134_138 | 219_223 | 137_137 |
| apis_026 | 0,39 | 0,02 | 0,45 | 0,05 | 0,08 | 12 | E.Med | 36,67 | 36,92 | 120_132 | 223_225 | 137_137 |
| apis_134 | 0,01 | 0,87 | 0,01 | 0,03 | 0,08 | 12 | E.Med | 36,81 | 37,85 | NA | NA | 137_137 |
| apis_027 | 0,01 | 0,04 | 0,86 | 0,02 | 0,07 | 12 | E.Med | 36,83 | 37,17 | 120_144 | 223_223 | 137_137 |
| apis_099 | 0,01 | 0,01 | 0,86 | 0,09 | 0,04 | 12 | E.Med | 36,92 | 36,89 | 140_158 | 223_223 | 137_137 |
| apis_098 | 0,01 | 0,01 | 0,95 | 0,01 | 0,02 | 12 | E.Med | 37,11 | 36,78 | 132_156 | 223_223 | 137_137 |
| apis_133 | 0,03 | 0,02 | 0,04 | 0,06 | 0,85 | 12 | E.Med | 37,13 | 37,48 | 130_140 | 217_225 | 133_137 |
| apis_100 | 0,01 | 0,01 | 0,91 | 0,01 | 0,05 | 12 | E.Med | 37,20 | 37,09 | 140_172 | 223_223 | 137_137 |
| apis_097 | 0,01 | 0,01 | 0,95 | 0,01 | 0,01 | 12 | E.Med | 37,39 | 36,72 | 111_120 | 219_223 | 137_137 |
| apis_118 | 0,02 | 0,01 | 0,91 | 0,03 | 0,03 | 12 | E.Med | 37,53 | 37,18 | 148_183 | 223_223 | 137_144 |
| apis_101 | 0,01 | 0,01 | 0,97 | 0,01 | 0,01 | 12 | E.Med | 37,54 | 37,35 | 120_124 | 223_228 | NA |
| apis_132 | 0,34 | 0,01 | 0,27 | 0,16 | 0,21 | 12 | E.Med | 37,55 | 37,82 | 122_132 | 223_223 | NA |
| apis_096 | 0,02 | 0,01 | 0,81 | 0,06 | 0,10 | 12 | E.Med | 37,60 | 36,87 | NA | NA | 133_137 |
| apis_116 | 0,01 | 0,01 | 0,94 | 0,02 | 0,02 | 12 | E.Med | 37,91 | 37,26 | NA | NA | 137_137 |
| apis_117 | 0,02 | 0,01 | 0,16 | 0,73 | 0,07 | 12 | E.Med | 37,99 | 36,95 | 140_178 | 219_223 | 137_137 |
| apis_102 | 0,01 | 0,05 | 0,89 | 0,03 | 0,02 | 12 | E.Med | 38,08 | 37,65 | 120_130 | 219_223 | 133_137 |
| apis_103 | 0,01 | 0,01 | 0,87 | 0,01 | 0,10 | 12 | E.Med | 38,13 | 37,81 | NA | NA | 137_137 |
| apis_104 | 0,02 | 0,02 | 0,60 | 0,02 | 0,34 | 12 | E.Med | 38,27 | 38,04 | 130_134 | 223_223 | 137_137 |
| apis_113 | 0,04 | 0,01 | 0,58 | 0,24 | 0,13 | 12 | E.Med | 38,36 | 36,96 | 122_134 | 223_223 | 137_137 |
| apis_114 | 0,01 | 0,01 | 0,89 | 0,03 | 0,06 | 12 | E.Med | 38,38 | 37,43 | 111_144 | 219_219 | 133_137 |
| apis_105 | 0,17 | 0,13 | 0,13 | 0,45 | 0,12 | 12 | E.Med | 38,65 | 37,91 | 111_142 | 219_219 | 131_131 |
| apis_112 | 0,03 | 0,01 | 0,85 | 0,04 | 0,06 | 12 | E.Med | 38,92 | 36,72 | 132_178 | 223_223 | 137_137 |
| apis_115 | 0,01 | 0,02 | 0,85 | 0,07 | 0,04 | 12 | E.Med | 38,95 | 37,60 | 113_130 | 217_219 | 137_137 |
| apis_106 | 0,01 | 0,03 | 0,83 | 0,09 | 0,03 | 12 | E.Med | 39,04 | 38,03 | 118_138 | 217_219 | 137_137 |

| | | | | | | | | | | | | |
|---|---|---|---|---|---|---|---|---|---|---|---|---|
| apis_111 | 0,05 | 0,00 | 0,92 | 0,01 | 0,02 | 12 | E.Med | 39,20 | 37,33 | 135_138 | 219_223 | NA |
| apis_110 | 0,01 | 0,01 | 0,93 | 0,02 | 0,03 | 12 | E.Med | 39,51 | 37,20 | 111_170 | 225_225 | NA |
| apis_107 | 0,03 | 0,44 | 0,02 | 0,13 | 0,38 | 12 | E.Med | 39,68 | 37,75 | 128_138 | 219_219 | 137_137 |
| apis_109 | 0,01 | 0,01 | 0,93 | 0,03 | 0,02 | 12 | E.Med | 39,99 | 37,24 | 124_144 | 222_223 | 137_137 |
| apis_108 | 0,02 | 0,01 | 0,83 | 0,10 | 0,05 | 12 | E.Med | 40,06 | 36,91 | NA | NA | 137_142 |
| apis_375A | 0,03 | 0,07 | 0,02 | 0,74 | 0,13 | 13 | Zagros | 42,85 | 37,91 | 144_166 | 223_223 | NA |
| apis_375B | 0,01 | 0,01 | 0,03 | 0,91 | 0,05 | 13 | Zagros | 42,86 | 37,92 | 135_158 | 223_223 | 137_137 |
| apis_374A | 0,03 | 0,04 | 0,11 | 0,10 | 0,73 | 13 | Zagros | 42,97 | 38,26 | NA | NA | 137_137 |
| apis_374B | 0,04 | 0,03 | 0,03 | 0,76 | 0,15 | 13 | Zagros | 42,98 | 38,26 | 120_120 | 223_223 | 133_137 |
| apis_370A | 0,01 | 0,18 | 0,01 | 0,21 | 0,60 | 13 | Zagros | 43,29 | 39,20 | 107_128 | 219_219 | NA |
| apis_370B | 0,03 | 0,60 | 0,05 | 0,10 | 0,21 | 13 | Zagros | 43,30 | 39,20 | 107_164 | 219_219 | 137_137 |
| apis_372A | 0,01 | 0,01 | 0,65 | 0,18 | 0,16 | 13 | Zagros | 43,32 | 38,83 | 146_158 | 223_223 | NA |
| apis_372B | 0,01 | 0,01 | 0,08 | 0,77 | 0,13 | 13 | Zagros | 43,33 | 38,83 | 111_138 | 219_219 | 137_137 |
| apis_373A | 0,04 | 0,01 | 0,04 | 0,77 | 0,14 | 13 | Zagros | 43,35 | 38,53 | NA | NA | 133_133 |
| apis_373B | 0,30 | 0,08 | 0,02 | 0,52 | 0,08 | 13 | Zagros | 43,36 | 38,53 | 140_142 | 219_219 | 137_137 |
| apis_383A | 0,01 | 0,01 | 0,05 | 0,91 | 0,02 | 13 | Zagros | 43,48 | 37,35 | NA | NA | 131_137 |
| apis_383B | 0,05 | 0,03 | 0,18 | 0,60 | 0,14 | 13 | Zagros | 43,49 | 37,35 | NA | NA | 137_137 |
| apis_371A | 0,01 | 0,71 | 0,01 | 0,13 | 0,15 | 13 | Zagros | 43,60 | 38,99 | 140_152 | 217_223 | 137_137 |
| apis_371B | 0,01 | 0,01 | 0,07 | 0,89 | 0,02 | 13 | Zagros | 43,61 | 38,99 | 132_144 | 219_223 | 137_137 |
| apis_382A | 0,02 | 0,04 | 0,15 | 0,21 | 0,58 | 13 | Zagros | 43,61 | 37,24 | NA | NA | 133_137 |
| apis_381A | 0,01 | 0,01 | 0,08 | 0,86 | 0,04 | 13 | Zagros | 43,62 | 37,25 | 135_154 | 219_223 | NA |
| apis_382B | 0,01 | 0,12 | 0,04 | 0,52 | 0,31 | 13 | Zagros | 43,62 | 37,49 | 107_135 | 219_223 | 137_137 |
| apis_381B | 0,01 | 0,02 | 0,17 | 0,76 | 0,04 | 13 | Zagros | 43,63 | 37,49 | 135_156 | 219_223 | NA |
| apis_380A | 0,01 | 0,23 | 0,12 | 0,57 | 0,06 | 13 | Zagros | 43,78 | 37,51 | 140_144 | 219_223 | 137_137 |
| apis_380B | 0,02 | 0,03 | 0,46 | 0,16 | 0,34 | 13 | Zagros | 43,79 | 37,51 | 146_146 | 223_223 | 133_137 |
| apis_376A | 0,05 | 0,02 | 0,14 | 0,66 | 0,13 | 13 | Zagros | 43,98 | 38,59 | 130_140 | 219_223 | 137_137 |
| apis_376B | 0,01 | 0,03 | 0,01 | 0,88 | 0,08 | 13 | Zagros | 43,99 | 38,59 | 132_166 | 219_219 | 137_137 |
| apis_378A | 0,04 | 0,16 | 0,58 | 0,15 | 0,07 | 13 | Zagros | 43,99 | 37,70 | 146_178 | 219_223 | 137_137 |
| apis_378B | 0,02 | 0,25 | 0,06 | 0,36 | 0,31 | 13 | Zagros | 43,99 | 37,71 | 135_166 | 219_219 | 137_137 |
| apis_379B | 0,01 | 0,01 | 0,00 | 0,01 | 0,96 | 13 | Zagros | 43,99 | 37,72 | 135_140 | 219_219 | 137_137 |
| apis_379A | 0,03 | 0,15 | 0,64 | 0,08 | 0,10 | 13 | Zagros | 44,00 | 37,71 | 138_146 | 219_223 | 133_137 |
| apis_387A | 0,01 | 0,00 | 0,01 | 0,95 | 0,02 | 13 | Zagros | 44,10 | 37,42 | 126_128 | 219_219 | NA |
| apis_387B | 0,01 | 0,03 | 0,08 | 0,77 | 0,12 | 13 | Zagros | 44,11 | 37,42 | 120_126 | 214_219 | 137_137 |
| apis_386 | 0,01 | 0,08 | 0,01 | 0,80 | 0,10 | 13 | Zagros | 44,16 | 37,43 | NA | NA | 137_137 |
| apis_377A | 0,02 | 0,04 | 0,30 | 0,44 | 0,20 | 13 | Zagros | 44,16 | 38,10 | 138_142 | 219_223 | 137_137 |
| apis_377B | 0,05 | 0,81 | 0,03 | 0,06 | 0,05 | 13 | Zagros | 44,17 | 38,10 | 118_130 | 223_223 | 131_137 |
| apis_384A | 0,01 | 0,01 | 0,19 | 0,15 | 0,64 | 13 | Zagros | 44,37 | 37,54 | NA | NA | 137_137 |
| apis_384B | 0,01 | 0,04 | 0,01 | 0,10 | 0,84 | 13 | Zagros | 44,38 | 37,54 | 142_142 | 219_223 | 137_137 |
| apis_391A | 0,02 | 0,01 | 0,05 | 0,21 | 0,71 | 13 | Zagros | 44,44 | 37,11 | 128_128 | 219_219 | 137_137 |
| apis_391B | 0,04 | 0,01 | 0,04 | 0,56 | 0,36 | 13 | Zagros | 44,45 | 37,11 | 128_144 | 222_223 | 137_137 |
| apis_388A | 0,01 | 0,08 | 0,04 | 0,85 | 0,02 | 13 | Zagros | 44,59 | 37,40 | NA | NA | 140_140 |
| apis_388B | 0,03 | 0,07 | 0,02 | 0,85 | 0,03 | 13 | Zagros | 44,60 | 37,40 | 132_135 | 223_223 | 133_137 |
| apis_385A | 0,01 | 0,21 | 0,18 | 0,32 | 0,28 | 13 | Zagros | 44,60 | 37,71 | NA | NA | 137_137 |
| apis_385B | 0,02 | 0,06 | 0,05 | 0,29 | 0,58 | 13 | Zagros | 44,60 | 37,72 | 120_148 | 217_219 | 137_137 |
| apis_389A | 0,01 | 0,02 | 0,53 | 0,08 | 0,36 | 13 | Zagros | 44,60 | 37,38 | 124_135 | 219_223 | 137_137 |
| apis_389B | 0,01 | 0,09 | 0,25 | 0,52 | 0,13 | 13 | Zagros | 44,61 | 37,38 | 132_138 | 219_219 | 137_137 |
| apis_390A | 0,00 | 0,01 | 0,06 | 0,84 | 0,08 | 13 | Zagros | 44,62 | 37,38 | 132_135 | 219_223 | 133_137 |
| apis_390B | 0,01 | 0,05 | 0,06 | 0,70 | 0,18 | 13 | Zagros | 44,63 | 37,38 | 116_142 | 221_223 | 132_132 |

| A043 | A079 | A088 | A107 | A113 | AB024 | AB124 | AC006 | AC306 | AP001 |
|---|---|---|---|---|---|---|---|---|---|
| 140_142 | 104_106 | 149_150 | 160_166 | 213_237 | 104_106 | 214_228 | 150_158 | 163_177 | 210_210 |
| 140_140 | 110_112 | 150_150 | 162_166 | 213_213 | 104_106 | 214_214 | 150_158 | 163_177 | 210_213 |
| 125_140 | 106_110 | 150_150 | 156_162 | 213_219 | 104_104 | 212_214 | 150_158 | 177_177 | 210_210 |
| 125_140 | 104_106 | 150_150 | 160_162 | 213_219 | 104_104 | 212_214 | 148_150 | 163_177 | 210_210 |
| 125_125 | 106_110 | 150_150 | 159_160 | 213_219 | 104_104 | 212_214 | 150_158 | 163_177 | 210_210 |
| 140_142 | 104_106 | 150_150 | 156_162 | 213_219 | 104_104 | 214_216 | 150_158 | 177_177 | 210_210 |
| 140_140 | 102_104 | 149_149 | 158_160 | 213_219 | 104_106 | 218_222 | 150_158 | 163_172 | 210_210 |
| 140_142 | 104_110 | 149_150 | 162_170 | 213_213 | 104_106 | 222_222 | 150_158 | 163_177 | 210_213 |
| 127_140 | 98_102 | 149_150 | 165_172 | 213_213 | 96_104 | 214_233 | 150_158 | 168_184 | 210_210 |
| 140_140 | 106_110 | 150_150 | 158_160 | 219_233 | 96_106 | 212_214 | 150_158 | 163_163 | 213_253 |
| 140_140 | 98_104 | 150_150 | 158_168 | 213_233 | 96_106 | 212_222 | 150_152 | 163_184 | 210_210 |
| 140_140 | 110_112 | 150_150 | 158_158 | 213_219 | 96_106 | 214_214 | 150_152 | 163_184 | 210_253 |
| 127_140 | 106_110 | 142_150 | 158_168 | 219_233 | 96_104 | 212_214 | 150_152 | 163_163 | 210_263 |
| 127_140 | 104_106 | 149_150 | 164_171 | 213_213 | 106_106 | 214_228 | 158_158 | 163_184 | 210_210 |
| 127_127 | 106_110 | 149_150 | 162_164 | 213_213 | 96_104 | 214_216 | 150_150 | 175_184 | 210_210 |
| 127_140 | 106_106 | 150_150 | 168_173 | 213_213 | 96_104 | 214_214 | 150_158 | 163_175 | 210_210 |
| 125_140 | 106_106 | 150_150 | 164_168 | 213_213 | 96_104 | 214_214 | 150_158 | 163_175 | 210_210 |
| 127_140 | 102_106 | 150_150 | 162_164 | 213_213 | 106_106 | 214_228 | 150_158 | 163_175 | 210_210 |
| 139_142 | 102_106 | 142_150 | 160_160 | 213_213 | 106_106 | 214_214 | 150_158 | 163_163 | 210_215 |
| 140_140 | 98_112 | 142_150 | 160_164 | 213_213 | 104_104 | 214_214 | 150_158 | 163_172 | 210_210 |
| 125_140 | 102_106 | 150_154 | 160_165 | 213_219 | 104_106 | 214_218 | 158_158 | 163_177 | 210_243 |
| 125_142 | 106_112 | 142_150 | 160_160 | 213_219 | 106_106 | 214_214 | 155_158 | 177_179 | 210_215 |
| 140_142 | 106_112 | 150_150 | 160_168 | 213_219 | 106_106 | 214_222 | 158_158 | 163_177 | 210_215 |
| 127_140 | 92_106 | 148_150 | 158_162 | 219_219 | 104_104 | 214_216 | 150_152 | 167_184 | 210_247 |
| 127_140 | 92_104 | 150_150 | 162_170 | 213_213 | 106_106 | 214_214 | 152_158 | 163_163 | 210_213 |
| 127_142 | 92_110 | 150_150 | 162_170 | 213_219 | 106_106 | 214_216 | 150_158 | 163_163 | 210_210 |
| 125_142 | 106_116 | 142_150 | 160_164 | 213_219 | 96_106 | 220_224 | 150_150 | 163_163 | 210_213 |
| 125_140 | 106_108 | 142_150 | 168_171 | 213_213 | 106_106 | 212_222 | 150_150 | 163_172 | 210_210 |
| 127_140 | 106_106 | 142_150 | 164_168 | 213_219 | 98_106 | 212_222 | 150_158 | 163_163 | 210_213 |
| 142_142 | 108_108 | 142_154 | 157_171 | 213_219 | 104_106 | 218_222 | 150_150 | 163_163 | 210_215 |
| 142_142 | 108_112 | 146_150 | 168_171 | 213_219 | 104_106 | 214_218 | 150_158 | 163_163 | 210_210 |
| 142_142 | 106_112 | 145_150 | 159_168 | 213_219 | 104_104 | 214_218 | 150_152 | 163_184 | 210_215 |
| 140_142 | 106_112 | 142_150 | 166_171 | 213_219 | 104_104 | 214_218 | 150_152 | 163_163 | 210_215 |
| 140_142 | 92_106 | 150_152 | 166_166 | 213_215 | 104_106 | 218_222 | 150_150 | 163_163 | 210_230 |
| 125_140 | 92_110 | 142_150 | 160_166 | 213_215 | 104_104 | 218_220 | 150_150 | 163_163 | 210_210 |
| 125_140 | 104_110 | 145_145 | 159_160 | 213_213 | 104_104 | 214_218 | 150_150 | 163_175 | 210_210 |
| 140_140 | 92_106 | 142_150 | 164_166 | 213_215 | 104_104 | 218_220 | 150_150 | 163_163 | 210_210 |
| 125_125 | 104_106 | 144_144 | 161_168 | 213_213 | 104_106 | 214_218 | 146_150 | 163_175 | 210_210 |
| 140_140 | 98_98 | 139_142 | 146_179 | 229_233 | 104_108 | 220_220 | 152_152 | 167_172 | 213_215 |
| 140_140 | 98_104 | 142_142 | 140_181 | 213_231 | 104_108 | 220_230 | 152_152 | 172_175 | 210_215 |
| 133_140 | NA | NA | 144_144 | NA | NA | 220_222 | 152_152 | NA | NA |
| 134_140 | NA | NA | 144_171 | NA | NA | 220_226 | 152_152 | NA | NA |
| 133_133 | NA | NA | 144_173 | NA | NA | NA | 152_152 | NA | NA |
| 140_140 | 104_104 | 139_142 | 156_162 | 229_233 | 104_106 | 220_222 | 152_152 | 163_172 | 215_215 |
| 140_140 | NA | NA | 144_177 | NA | NA | 210_230 | 152_152 | NA | NA |
| 125_140 | 98_104 | 142_150 | 140_181 | 225_231 | 104_106 | 220_220 | 150_152 | 163_167 | 210_215 |
| 140_140 | 98_104 | 139_139 | 164_175 | 213_227 | 104_104 | 224_243 | 152_158 | 167_175 | 210_210 |
| 140_140 | NA | NA | 145_171 | NA | NA | 222_228 | 152_152 | NA | NA |

| | | | | | | | | | |
|---|---|---|---|---|---|---|---|---|---|
| 140_140 | 98_108 | 149_150 | 160_171 | 213_227 | 104_106 | 210_220 | 150_152 | 163_163 | 210_210 |
| 119_125 | 104_106 | 139_150 | 164_173 | 213_213 | 104_106 | 214_222 | 150_152 | 163_170 | 210_213 |
| 140_142 | NA | NA | 144_144 | NA | NA | 220_222 | 152_152 | NA | NA |
| 140_140 | NA | NA | 144_173 | NA | NA | 220_220 | 152_152 | NA | NA |
| 140_140 | 98_106 | 139_150 | 144_156 | 231_233 | 104_106 | 214_214 | 152_152 | 167_172 | 210_215 |
| 140_140 | NA | 139_139 | 156_171 | 213_223 | 104_108 | 220_220 | 152_152 | NA | 213_215 |
| 140_140 | 98_100 | 150_150 | 148_160 | 213_229 | 104_106 | 214_235 | 150_152 | 172_175 | 215_223 |
| 140_142 | 102_104 | 142_142 | 168_175 | 229_235 | 106_106 | 214_216 | 152_152 | 172_175 | 213_213 |
| 140_140 | 98_108 | 142_150 | 146_160 | 213_231 | 104_106 | 216_235 | 152_152 | 167_172 | 210_215 |
| 140_140 | NA | NA | 140_177 | NA | NA | 218_230 | 152_152 | NA | NA |
| 140_142 | 102_104 | 142_150 | 154_156 | 231_237 | 104_106 | 220_228 | 150_152 | 163_172 | 213_249 |
| 140_140 | 102_102 | 142_149 | 146_164 | 207_213 | 104_106 | 218_224 | 150_152 | 167_175 | 210_215 |
| 140_140 | NA | NA | 150_168 | NA | NA | 220_236 | 152_152 | NA | NA |
| 140_142 | NA | NA | 142_171 | NA | NA | 218_220 | 152_152 | NA | NA |
| 140_140 | 104_108 | 139_142 | 145_163 | 213_213 | 104_104 | 220_230 | 152_152 | 163_172 | 210_213 |
| 125_140 | 102_106 | 142_142 | 171_177 | 213_229 | 104_104 | 218_226 | 152_152 | 167_175 | 215_217 |
| 140_140 | 98_98 | 139_142 | 144_144 | 227_229 | 106_106 | 220_222 | 152_152 | 167_175 | 213_215 |
| 140_140 | 102_108 | 150_150 | 144_163 | 227_233 | 104_106 | 224_235 | 150_152 | 163_175 | 215_233 |
| 140_142 | NA | NA | 142_146 | NA | NA | 228_230 | 148_152 | NA | NA |
| 139_140 | 106_108 | 139_150 | 140_168 | 225_227 | 96_106 | 214_218 | 150_152 | 163_167 | 210_217 |
| 140_142 | 98_102 | 139_150 | 159_168 | 213_213 | 106_106 | 210_235 | 152_155 | 172_172 | 210_233 |
| 140_140 | NA | NA | 168_169 | NA | NA | 220_220 | 152_152 | NA | NA |
| 140_140 | NA | NA | 144_168 | NA | NA | 218_220 | 152_152 | NA | NA |
| 140_140 | NA | NA | 165_179 | NA | NA | 218_220 | 152_152 | NA | NA |
| 140_142 | NA | NA | 164_177 | NA | NA | 220_220 | 152_152 | NA | NA |
| 125_140 | 98_108 | 139_139 | 162_181 | 213_225 | 106_106 | 210_220 | 150_152 | 163_163 | 213_245 |
| 140_140 | NA | NA | 140_177 | NA | NA | 222_226 | 152_152 | NA | NA |
| 140_140 | 102_110 | 150_150 | 144_156 | 213_213 | 104_106 | 214_220 | 152_152 | 163_172 | 210_213 |
| 140_140 | 102_108 | 142_150 | 144_158 | 213_227 | 104_104 | 218_220 | 152_152 | 163_163 | 213_213 |
| 138_140 | NA | NA | 175_177 | NA | NA | 222_226 | 152_152 | NA | NA |
| 140_140 | 98_106 | 139_150 | 148_169 | 213_223 | 96_106 | 220_220 | 152_152 | 170_172 | 213_217 |
| 140_140 | 98_98 | 139_150 | 144_156 | 213_213 | 106_108 | 216_235 | 150_152 | 167_175 | 213_213 |
| 140_142 | NA | 142_150 | 140_156 | 225_227 | 106_106 | 214_225 | 150_152 | NA | 210_213 |
| 140_140 | NA | NA | 144_148 | NA | NA | 220_220 | 152_152 | NA | NA |
| 140_140 | 98_102 | 139_152 | 144_171 | 219_231 | 106_106 | 218_220 | 152_152 | 175_175 | 213_213 |
| 140_140 | NA | NA | 146_179 | NA | NA | 218_230 | 152_152 | NA | NA |
| 125_140 | 102_106 | 150_150 | 164_168 | 213_225 | 106_106 | 218_220 | 150_150 | 163_163 | 213_215 |
| 140_140 | 98_102 | 150_150 | 140_170 | 213_229 | 106_108 | 214_222 | 150_150 | 163_172 | 215_231 |
| 140_140 | 102_104 | 142_142 | 160_168 | 213_225 | 106_106 | 220_233 | 150_152 | 175_175 | 213_213 |
| 140_140 | NA | NA | 144_173 | NA | NA | 210_226 | 152_152 | NA | NA |
| 140_142 | 98_98 | 139_150 | 144_164 | 213_229 | 106_106 | 220_220 | 152_152 | 163_172 | 210_215 |
| 140_140 | 98_106 | 142_150 | 156_166 | 213_221 | 106_106 | 216_220 | 150_150 | 167_175 | 213_217 |
| 140_140 | 106_114 | 139_150 | 156_179 | 213_213 | 106_106 | 210_220 | 150_152 | 163_163 | 213_213 |
| 140_140 | 98_102 | 139_142 | 168_170 | 213_213 | 104_106 | 216_236 | 150_152 | 163_163 | 213_250 |
| 140_140 | 98_104 | 150_150 | 156_179 | 213_213 | 104_104 | 216_236 | 148_152 | 167_170 | 256_256 |
| 140_140 | 98_98 | 139_149 | 156_158 | 213_231 | 104_106 | 210_220 | 152_152 | 163_172 | 213_213 |
| 140_140 | 98_108 | 145_150 | 146_164 | 213_213 | 106_106 | 218_218 | 152_155 | 167_167 | 210_231 |
| 140_140 | 98_104 | 139_150 | 171_181 | 213_233 | 106_108 | 220_224 | 150_152 | 163_167 | 231_246 |

| | | | | | | | | | |
|---|---|---|---|---|---|---|---|---|---|
| 140_140 | 104_104 | 139_150 | 150_162 | 213_225 | 106_106 | 212_214 | 150_152 | 172_172 | 215_215 |
| 140_140 | 98_98 | 139_150 | NA | 213_227 | 106_106 | 212_228 | NA | 163_163 | 215_231 |
| 140_142 | 98_110 | 142_150 | 156_171 | 213_227 | 96_106 | 220_228 | 150_152 | 163_177 | 215_215 |
| 140_140 | 98_102 | 139_150 | 156_173 | 225_233 | 104_108 | 218_220 | 152_152 | 163_184 | 213_215 |
| 140_140 | 106_114 | 142_149 | 142_181 | 213_223 | 104_108 | 214_220 | 152_152 | 163_172 | 213_213 |
| 140_140 | 98_108 | 150_150 | 156_166 | 213_227 | 104_104 | 220_228 | 152_152 | 163_172 | 245_271 |
| 140_140 | 98_108 | 139_150 | 156_173 | 221_231 | 104_104 | 210_228 | 150_152 | 172_172 | 210_210 |
| 140_140 | NA | NA | 140_146 | NA | NA | 220_220 | 152_152 | NA | NA |
| 140_144 | NA | NA | 142_144 | NA | NA | 220_220 | 152_152 | NA | NA |
| 140_140 | 98_106 | 139_142 | 160_162 | 213_213 | 104_104 | 218_218 | 152_152 | 163_172 | 217_217 |
| 125_140 | 104_108 | 149_150 | 156_168 | 213_229 | 104_106 | 216_220 | 150_150 | 163_163 | 210_213 |
| 140_140 | NA | NA | 141_173 | NA | NA | 230_230 | 152_152 | NA | NA |
| 140_140 | 104_104 | 139_142 | 144_154 | 213_223 | 106_106 | 224_235 | 152_152 | 172_172 | 213_213 |
| 140_140 | 106_106 | 142_150 | 164_170 | 213_213 | 106_108 | 220_233 | 152_152 | 163_167 | 213_253 |
| 125_140 | 110_114 | 150_150 | 166_169 | 213_213 | 104_106 | 214_220 | 150_152 | 163_172 | 219_243 |
| 140_140 | 104_110 | 150_150 | 144_164 | 223_229 | 104_106 | 210_236 | 150_155 | 163_172 | 215_215 |
| 125_140 | 98_102 | 142_149 | 144_160 | 219_233 | 106_106 | 220_220 | 150_152 | 170_172 | 215_215 |
| 140_140 | 98_98 | 149_150 | 140_144 | 213_213 | 106_106 | 212_218 | 150_152 | 163_172 | 210_215 |
| 140_140 | 98_102 | 150_154 | 156_158 | 213_231 | 104_106 | 210_218 | 150_150 | 167_172 | 213_213 |
| 140_144 | 98_106 | 139_142 | 144_158 | 213_227 | 104_106 | 220_233 | 152_152 | 167_175 | 213_215 |
| 140_140 | 98_102 | 142_149 | 155_173 | 213_213 | 104_106 | 218_220 | 152_155 | 175_175 | 215_251 |
| 125_140 | 98_102 | 150_150 | 162_162 | 227_233 | 104_104 | 214_220 | 150_152 | 172_172 | 229_252 |
| 140_142 | NA | NA | 144_144 | NA | NA | 220_222 | 152_152 | NA | NA |
| 140_140 | 98_104 | 139_142 | 146_156 | 225_227 | 104_106 | 220_222 | 150_152 | 163_172 | 213_233 |
| 140_140 | 104_106 | 139_139 | 150_156 | 213_221 | 106_106 | 210_220 | 150_150 | 163_163 | 210_210 |
| 140_140 | 98_108 | 142_149 | 146_175 | 225_227 | 108_108 | 220_230 | 150_152 | 163_163 | 210_217 |
| 140_140 | 104_108 | 139_139 | 156_158 | 213_225 | 106_106 | 216_222 | 150_152 | 167_172 | 213_249 |
| 125_140 | 102_108 | 142_149 | 156_181 | 213_213 | 104_104 | 218_220 | 150_152 | 167_172 | 210_210 |
| 127_140 | 98_104 | 142_142 | 146_156 | 213_223 | 104_106 | 216_236 | 150_152 | 163_167 | 210_219 |
| 138_140 | NA | NA | 146_175 | NA | NA | 220_228 | 152_152 | NA | NA |
| 140_140 | 98_104 | 149_150 | 168_171 | 213_227 | 104_106 | 220_228 | 152_158 | 163_172 | 210_213 |
| 140_140 | NA | NA | 146_154 | NA | NA | 220_224 | 152_152 | NA | NA |
| 140_140 | 106_110 | 139_149 | 140_164 | 229_239 | 104_106 | 214_220 | 152_152 | 170_172 | 213_255 |
| 140_140 | 102_106 | 139_150 | 144_156 | 213_225 | 104_106 | 216_222 | 152_152 | 172_175 | 213_215 |
| 140_140 | NA | NA | 163_173 | NA | NA | 222_233 | 152_152 | NA | NA |
| 125_140 | 98_104 | 142_149 | 152_164 | 213_225 | 106_106 | 220_239 | 152_152 | 163_167 | 213_233 |
| 138_140 | 98_104 | 139_139 | 171_173 | 213_213 | 104_104 | 218_228 | 152_152 | 167_167 | 215_233 |
| 140_140 | NA | NA | 144_144 | NA | NA | 220_228 | 137_152 | NA | NA |
| 140_140 | 98_98 | 139_139 | 148_165 | 223_227 | 104_106 | 210_222 | 152_152 | 167_175 | 215_259 |
| 125_140 | 98_98 | 139_150 | 139_162 | 213_227 | 106_106 | 220_230 | 150_152 | 172_175 | 213_215 |
| 139_140 | NA | NA | 169_171 | NA | NA | 210_224 | 152_152 | NA | NA |
| 140_140 | 98_106 | 139_142 | 158_173 | 223_225 | 106_106 | 220_220 | 150_152 | 167_179 | 215_239 |
| 140_140 | 98_102 | 142_150 | 156_173 | 229_233 | 104_106 | 216_220 | 152_152 | 163_172 | 213_215 |
| 140_142 | NA | NA | 144_154 | NA | NA | 218_233 | 152_152 | NA | NA |
| 142_144 | NA | NA | 144_158 | NA | NA | 220_224 | 152_152 | NA | NA |
| 140_140 | 104_106 | 139_142 | 146_158 | 211_213 | 104_104 | 218_222 | 152_152 | 167_172 | 215_215 |
| 140_142 | 98_102 | 139_142 | 144_161 | 229_231 | 104_106 | 220_228 | 152_152 | 167_167 | 215_237 |
| 140_140 | 102_102 | 149_150 | 144_171 | 213_225 | 106_106 | 222_230 | 150_152 | 167_170 | 215_257 |

| | | | | | | | | | |
|---|---|---|---|---|---|---|---|---|---|
| 142_146 | 102_114 | 142_150 | 144_181 | 227_229 | 96_106 | 220_220 | 150_152 | 167_172 | 210_249 |
| 140_142 | 102_104 | 142_149 | 156_164 | 213_225 | 106_106 | 212_241 | 150_152 | 163_172 | 233_245 |
| 140_140 | 98_98 | 139_142 | 165_169 | 209_223 | 102_106 | 210_235 | 152_152 | 172_175 | 213_215 |
| 140_140 | 102_102 | 142_142 | 144_160 | 225_231 | 106_106 | 218_220 | 152_152 | 167_175 | 213_215 |
| 142_142 | 98_104 | 139_142 | NA | 221_229 | 104_106 | 220_224 | 152_152 | 167_172 | 213_233 |
| 140_144 | NA | NA | 144_175 | NA | NA | 220_220 | 152_152 | NA | NA |
| 140_140 | NA | NA | 144_171 | NA | NA | 216_220 | 152_152 | NA | NA |
| 140_142 | NA | NA | 144_162 | NA | NA | 218_220 | 152_152 | NA | NA |
| 140_140 | NA | NA | 165_173 | NA | NA | 212_220 | 148_152 | NA | NA |
| 139_140 | NA | NA | 144_154 | NA | NA | 220_220 | 152_152 | NA | NA |
| 125_140 | 98_104 | 142_150 | 139_166 | 215_225 | 104_106 | 214_226 | 152_152 | 163_172 | 213_213 |
| 139_140 | NA | NA | 144_179 | NA | NA | 220_233 | 152_152 | NA | NA |
| 140_140 | NA | NA | 158_159 | NA | NA | 220_230 | 152_152 | NA | NA |
| 140_140 | NA | NA | 168_169 | NA | NA | 220_228 | 152_152 | NA | NA |
| 140_140 | NA | NA | 144_169 | NA | NA | 220_228 | 152_152 | NA | NA |
| 140_141 | NA | NA | 162_177 | NA | NA | 214_220 | 152_152 | NA | NA |
| 140_144 | NA | NA | 144_144 | NA | NA | 214_224 | 152_152 | NA | NA |
| 140_142 | NA | NA | 173_175 | NA | NA | 220_228 | 148_152 | NA | NA |
| 140_140 | NA | NA | 144_163 | NA | NA | 218_235 | 152_152 | NA | NA |
| 140_144 | 98_104 | 139_139 | 162_168 | 223_229 | 106_106 | 218_220 | 152_152 | 172_175 | 233_245 |
| 140_140 | NA | NA | 140_144 | NA | NA | 218_220 | 152_152 | NA | NA |
| 140_144 | 104_106 | 145_150 | 142_173 | 227_233 | 106_106 | 220_226 | 152_152 | 167_172 | 210_237 |
| 140_140 | NA | NA | 144_171 | NA | NA | 220_233 | 152_152 | NA | NA |
| 140_140 | NA | NA | 144_144 | NA | NA | 220_222 | 152_152 | NA | NA |
| 142_142 | NA | NA | 162_175 | NA | NA | 222_228 | 152_152 | NA | NA |
| 140_140 | NA | NA | 142_168 | NA | NA | 220_220 | 152_152 | NA | NA |
| 140_140 | NA | NA | 144_164 | NA | NA | 222_233 | 152_152 | NA | NA |
| 140_140 | NA | NA | 168_175 | NA | NA | 218_220 | 152_152 | NA | NA |
| 140_140 | 98_104 | 139_139 | 158_171 | 207_229 | 106_106 | 218_220 | 152_152 | 175_175 | 213_215 |
| 140_140 | 98_102 | 139_142 | 144_160 | 229_231 | 104_108 | 220_228 | 152_152 | 172_172 | 213_215 |
| 140_140 | 98_106 | 142_142 | 144_169 | 221_227 | 104_106 | 220_222 | 152_155 | 172_175 | 215_257 |
| 140_140 | NA | NA | 160_177 | NA | NA | 220_236 | 152_152 | NA | NA |
| 140_140 | NA | NA | 144_173 | NA | NA | 218_228 | 152_152 | NA | NA |
| 140_140 | NA | NA | 146_160 | NA | NA | 220_220 | 152_152 | NA | NA |
| 140_140 | NA | NA | 146_177 | NA | NA | 220_220 | 152_152 | NA | NA |
| 140_140 | 98_98 | 139_142 | 144_158 | 209_231 | 104_106 | 218_220 | 152_152 | 172_175 | 215_249 |
| 140_140 | 106_106 | 139_139 | 144_169 | 223_229 | 104_106 | 220_220 | 152_152 | 167_167 | 249_263 |
| 140_140 | NA | NA | 144_179 | NA | NA | 233_233 | 152_152 | NA | NA |
| 140_140 | 98_104 | 142_142 | 146_179 | 225_231 | 104_106 | 220_222 | 152_152 | 167_175 | 213_259 |
| 140_140 | 98_98 | 142_142 | 158_171 | 227_227 | 96_106 | 220_226 | 152_152 | 172_172 | 215_215 |
| 140_140 | 98_102 | 142_142 | 142_144 | 227_229 | 104_106 | 220_220 | 152_152 | 172_175 | 213_259 |
| 140_144 | 102_108 | 139_142 | 144_169 | 223_225 | 96_106 | 218_220 | 152_152 | 175_175 | 210_213 |
| 140_140 | 98_98 | 139_142 | 162_169 | 227_231 | 106_106 | 218_220 | 152_152 | 172_172 | 215_215 |
| 140_140 | 98_104 | 139_142 | 144_169 | 229_244 | 106_106 | 220_220 | 152_152 | 167_167 | 215_249 |
| 140_140 | 102_106 | 139_142 | 171_171 | 227_229 | 104_104 | 210_224 | 152_152 | 172_175 | 213_213 |
| 140_142 | NA | NA | 144_144 | NA | NA | 214_226 | 152_152 | NA | NA |
| 139_140 | NA | NA | 168_168 | NA | NA | 220_222 | 152_152 | NA | NA |
| 140_140 | NA | NA | 169_176 | NA | NA | 226_226 | 152_152 | NA | NA |

| | | | | | | | | | |
|---|---|---|---|---|---|---|---|---|---|
| 140_140 | NA | NA | 144_160 | NA | NA | 210_214 | 152_152 | NA | NA |
| 140_142 | NA | NA | 150_171 | NA | NA | 218_222 | 152_152 | NA | NA |
| 140_140 | 104_106 | 139_142 | 144_144 | 229_231 | 104_106 | 228_230 | 152_152 | 172_175 | 213_215 |
| 140_140 | NA | NA | 166_169 | NA | NA | 222_224 | 150_152 | NA | NA |
| 140_140 | 98_98 | 139_139 | 144_154 | 225_227 | 104_104 | 220_222 | 152_152 | 172_175 | 213_213 |
| 140_140 | 98_104 | 139_150 | 160_169 | 221_229 | 96_106 | 220_220 | 152_152 | 172_175 | 213_215 |
| 140_142 | NA | NA | 144_169 | NA | NA | 220_220 | 152_152 | NA | NA |
| 140_140 | NA | NA | 144_169 | NA | NA | 218_224 | 146_152 | NA | NA |
| 140_140 | NA | NA | 144_175 | NA | NA | 220_224 | 152_152 | NA | NA |
| 140_142 | NA | NA | 148_173 | NA | NA | 220_228 | 152_152 | NA | NA |
| 140_140 | NA | NA | 144_158 | NA | NA | 220_220 | 152_152 | NA | NA |
| 140_140 | NA | NA | 156_179 | NA | NA | 220_220 | 152_152 | NA | NA |
| 140_142 | NA | NA | 144_173 | NA | NA | 214_218 | 152_152 | NA | NA |
| 140_140 | NA | NA | 168_171 | NA | NA | 218_220 | 152_152 | NA | NA |
| 140_140 | NA | NA | 150_158 | NA | NA | 220_224 | 150_152 | NA | NA |
| 140_140 | NA | NA | 142_173 | NA | NA | 218_220 | 152_152 | NA | NA |
| 140_140 | 98_106 | 139_142 | 173_175 | 223_231 | 106_106 | 222_222 | 152_152 | 167_172 | 213_215 |
| 140_140 | NA | NA | 171_177 | NA | NA | 220_220 | 152_152 | NA | NA |
| 140_140 | NA | NA | 173_175 | NA | NA | 220_220 | 150_152 | NA | NA |
| 140_140 | NA | NA | 144_155 | NA | NA | 216_220 | 152_152 | NA | NA |
| 140_142 | NA | NA | 169_179 | NA | NA | 218_220 | 152_152 | NA | NA |
| 140_140 | NA | NA | 144_162 | NA | NA | 224_228 | 150_152 | NA | NA |
| 140_142 | NA | NA | 144_169 | NA | NA | 224_226 | 152_152 | NA | NA |
| 140_142 | NA | NA | 162_168 | NA | NA | 220_230 | 152_152 | NA | NA |
| 140_140 | NA | NA | 146_154 | NA | NA | 218_220 | 152_152 | NA | NA |
| 140_142 | NA | NA | 144_168 | NA | NA | 220_235 | 152_152 | NA | NA |
| 140_144 | NA | NA | 144_163 | NA | NA | 220_220 | 152_152 | NA | NA |
| 140_142 | NA | NA | 144_144 | NA | NA | 220_220 | 152_152 | NA | NA |
| 140_140 | NA | NA | 160_181 | NA | NA | 222_226 | 152_152 | NA | NA |
| 140_140 | NA | NA | 148_181 | NA | NA | 220_235 | 152_152 | NA | NA |
| 140_140 | NA | NA | 144_144 | NA | NA | 220_224 | 150_152 | NA | NA |
| 139_140 | NA | NA | 156_175 | NA | NA | 220_236 | 152_152 | NA | NA |
| 140_140 | NA | NA | 158_165 | NA | NA | 218_220 | 152_152 | NA | NA |
| 140_142 | 102_102 | 139_139 | 144_175 | 223_223 | 106_106 | 218_222 | 152_152 | 175_175 | 213_213 |
| 140_140 | 98_102 | 142_142 | 144_168 | 227_227 | 96_104 | 218_220 | 152_152 | 172_175 | 213_215 |
| 140_142 | 98_104 | 139_142 | 146_148 | 221_223 | 104_106 | 220_224 | 152_152 | 172_172 | 213_239 |
| 140_142 | 98_106 | 139_139 | 144_144 | 213_231 | 94_106 | 220_220 | 150_152 | 172_172 | 213_213 |
| 140_140 | 98_98 | 139_142 | 144_154 | 229_231 | 104_106 | 224_224 | 152_152 | 167_175 | 213_215 |
| 140_144 | 98_104 | 139_142 | 146_173 | 225_231 | 96_106 | 220_226 | 152_152 | 167_175 | 213_213 |
| 140_140 | 98_104 | 139_139 | 146_168 | 221_221 | 104_106 | 220_220 | 152_152 | 172_175 | 213_215 |
| 140_140 | 98_98 | 142_142 | 154_160 | 221_227 | 104_106 | 220_220 | 152_152 | 167_167 | 213_229 |
| 140_142 | NA | NA | 173_175 | NA | NA | 218_220 | 125_147 | NA | NA |
| 140_140 | NA | NA | 169_175 | NA | NA | 220_228 | 152_152 | NA | NA |
| 140_142 | NA | NA | 144_144 | NA | NA | 218_220 | 152_152 | NA | NA |
| 140_140 | NA | NA | 146_154 | NA | NA | 220_222 | 152_152 | NA | NA |
| 140_140 | NA | NA | 171_172 | NA | NA | 218_218 | 152_152 | NA | NA |
| 142_142 | 102_104 | 139_139 | 142_175 | 223_227 | 104_106 | 222_228 | 152_152 | 167_172 | 215_215 |
| 140_140 | 98_104 | 142_142 | 144_173 | 223_227 | 96_106 | 214_220 | 152_152 | 167_172 | 210_213 |

| | | | | | | | | | |
|---|---|---|---|---|---|---|---|---|---|
| 140_142 | 98_102 | 139_139 | 164_166 | 225_229 | 106_106 | 220_222 | 152_152 | 172_175 | 213_245 |
| 140_146 | 98_98 | 142_150 | 144_156 | 221_227 | 96_104 | 214_218 | 152_152 | 175_175 | 213_225 |
| 142_142 | 98_104 | 139_142 | 144_162 | 209_225 | 106_106 | 222_226 | 152_152 | 172_175 | 213_215 |
| 140_142 | NA | NA | 144_169 | NA | NA | 218_220 | 152_152 | NA | NA |
| 146_146 | 102_104 | 139_142 | 162_179 | 227_227 | 106_106 | 220_224 | 152_152 | 172_172 | 215_253 |
| 140_140 | 104_104 | 139_142 | 162_171 | 223_227 | 98_104 | 220_224 | 152_152 | 167_172 | 213_215 |
| 140_140 | 98_98 | 142_142 | 144_168 | 227_229 | 104_106 | 220_220 | 152_152 | 167_172 | 213_249 |
| 140_140 | 98_106 | 139_142 | 144_168 | 227_237 | 104_104 | 220_220 | 152_152 | 172_175 | 213_217 |
| 140_142 | 98_106 | 139_142 | 169_175 | 233_235 | 104_106 | 220_220 | 150_152 | 167_175 | 213_213 |
| 142_146 | 98_98 | 139_142 | 162_168 | 227_231 | 104_106 | 220_220 | 152_152 | 175_175 | 210_215 |
| 140_140 | 98_102 | 142_142 | 147_181 | 233_235 | 106_106 | 220_224 | 150_152 | 167_172 | 213_213 |
| 140_140 | 98_98 | 139_142 | 144_168 | 221_225 | 104_106 | 218_220 | 152_152 | 172_172 | 213_215 |
| 140_140 | 102_104 | 142_142 | NA | 223_227 | 104_106 | 218_220 | NA | 167_175 | 210_215 |
| 140_140 | 98_104 | 142_150 | 144_173 | 227_229 | 104_106 | 220_220 | 150_150 | 172_172 | 215_217 |
| 140_140 | NA | NA | 144_171 | NA | NA | 220_224 | 152_152 | NA | NA |
| 140_144 | NA | NA | 144_169 | NA | NA | 220_224 | 150_152 | NA | NA |
| 140_142 | NA | NA | 142_150 | NA | NA | 220_220 | 152_152 | NA | NA |
| 139_139 | NA | NA | 148_175 | NA | NA | 218_218 | 152_152 | NA | NA |
| 140_140 | 98_98 | 139_139 | 144_177 | 223_235 | 106_106 | 216_220 | 152_152 | 167_172 | 213_213 |
| 140_140 | 104_104 | 139_142 | 144_173 | 209_219 | 104_104 | 216_220 | 152_152 | 167_172 | 213_215 |
| 125_146 | 104_106 | 139_149 | 168_175 | 225_227 | 104_106 | 220_220 | 152_152 | 175_175 | 213_237 |
| 142_142 | 98_102 | 139_142 | 144_168 | 223_235 | 106_106 | 218_220 | 152_152 | 172_175 | 213_231 |
| 142_142 | NA | NA | 165_179 | NA | NA | 220_220 | 152_152 | NA | NA |
| 140_142 | NA | NA | 173_177 | NA | NA | 212_228 | 150_152 | NA | NA |
| 140_142 | 98_104 | 149_149 | 168_177 | 225_225 | 104_104 | 220_230 | 152_152 | 172_175 | 213_215 |
| 140_140 | NA | NA | 148_173 | NA | NA | 218_220 | 152_152 | NA | NA |
| 140_142 | 102_104 | 139_142 | 142_144 | 221_225 | 96_106 | 218_220 | 152_152 | 167_172 | 213_213 |
| 140_140 | 102_102 | 139_142 | 162_181 | 213_223 | 102_106 | 218_220 | 152_152 | 167_175 | 213_239 |
| 140_140 | 102_106 | 142_142 | 154_169 | 225_231 | 104_106 | 220_224 | 152_152 | 167_170 | 213_213 |
| 140_142 | 98_98 | 142_142 | 144_158 | 225_227 | 104_106 | 220_220 | 152_152 | 172_172 | 235_253 |
| 140_142 | 106_114 | 142_142 | 158_164 | 225_225 | 106_106 | 218_220 | 152_152 | 172_175 | 213_213 |
| 140_140 | 98_98 | 139_142 | 144_144 | 223_223 | 104_104 | 220_220 | 152_152 | 167_175 | 213_213 |
| 138_142 | 102_104 | 142_142 | 144_175 | 223_225 | 96_106 | 220_224 | 152_152 | 172_175 | 213_213 |
| 140_142 | 102_104 | 139_139 | 171_177 | 221_221 | 104_106 | 220_224 | 152_152 | 175_175 | 215_231 |
| 140_144 | NA | NA | 144_162 | NA | NA | 222_222 | 152_152 | NA | NA |
| 140_142 | NA | NA | 142_177 | NA | NA | 220_222 | 152_152 | NA | NA |
| 140_142 | NA | NA | 154_181 | NA | NA | 220_224 | 152_152 | NA | NA |
| 140_140 | NA | NA | 147_154 | NA | NA | 220_220 | 152_152 | NA | NA |
| 140_142 | NA | NA | 158_169 | NA | NA | 220_222 | 152_152 | NA | NA |
| 140_140 | NA | NA | 146_146 | NA | NA | 220_222 | 152_152 | NA | NA |
| 140_140 | NA | NA | 144_150 | NA | NA | 220_220 | 152_152 | NA | NA |
| 140_142 | NA | NA | 165_173 | NA | NA | 220_222 | 152_152 | NA | NA |
| 140_142 | NA | NA | 169_179 | NA | NA | 220_220 | 152_152 | NA | NA |
| 140_142 | NA | NA | 156_162 | NA | NA | 220_220 | 152_152 | NA | NA |
| 140_140 | 98_106 | 134_139 | 153_155 | 225_229 | 106_106 | 220_224 | 152_152 | 172_172 | 215_217 |
| 140_140 | NA | NA | 144_164 | NA | NA | 226_233 | 150_152 | NA | NA |
| 140_140 | NA | NA | 156_156 | NA | NA | 214_220 | 152_152 | NA | NA |
| 140_140 | NA | NA | 158_161 | NA | NA | 218_220 | 152_152 | NA | NA |

| | | | | | | | | | |
|---|---|---|---|---|---|---|---|---|---|
| 142_142 | NA | NA | 146_156 | NA | NA | 218_220 | 152_152 | NA | NA |
| 140_142 | 102_106 | 139_142 | 162_181 | 227_233 | 104_104 | 220_224 | 152_152 | 172_175 | 213_217 |
| 140_140 | 104_104 | 135_142 | 161_177 | 221_231 | 102_104 | 224_226 | 152_152 | 172_172 | 215_215 |
| 140_140 | 102_104 | 139_139 | 158_169 | 227_229 | 106_106 | 222_228 | 152_152 | 169_175 | 213_259 |
| 142_142 | 102_104 | 142_142 | 144_175 | 225_231 | 104_106 | 220_222 | 152_152 | 167_172 | 215_239 |
| 140_140 | NA | NA | 142_148 | NA | NA | 220_224 | 152_152 | NA | NA |
| 139_140 | NA | NA | 144_156 | NA | NA | 220_222 | 152_152 | NA | NA |
| 140_140 | NA | NA | 144_150 | NA | NA | 218_220 | 152_152 | NA | NA |
| 140_140 | NA | NA | 156_171 | NA | NA | 220_228 | 152_152 | NA | NA |
| 140_142 | NA | NA | 160_160 | NA | NA | 218_220 | 152_152 | NA | NA |
| 140_142 | NA | NA | 161_175 | NA | NA | 220_220 | 149_152 | NA | NA |
| 142_144 | NA | NA | 144_158 | NA | NA | 220_222 | 152_155 | NA | NA |
| 140_140 | NA | NA | 144_148 | NA | NA | 220_230 | 152_152 | NA | NA |
| 140_140 | NA | NA | 142_175 | NA | NA | 218_222 | 152_152 | NA | NA |
| 140_140 | NA | NA | 161_161 | NA | NA | 218_228 | 150_150 | NA | NA |
| 140_144 | NA | NA | 169_179 | NA | NA | 222_233 | 152_152 | NA | NA |
| 144_144 | NA | NA | 144_173 | NA | NA | 222_226 | 152_152 | NA | NA |
| 140_140 | NA | NA | 144_171 | NA | NA | 220_222 | 152_152 | NA | NA |
| 140_142 | NA | NA | 144_156 | NA | NA | 218_220 | 152_155 | NA | NA |
| 142_142 | NA | NA | 144_158 | NA | NA | 218_220 | 152_152 | NA | NA |
| 140_140 | NA | NA | 158_181 | NA | NA | 220_220 | 152_152 | NA | NA |
| 140_140 | NA | NA | 150_160 | NA | NA | 218_222 | 152_152 | NA | NA |
| NA | NA | NA | 158_164 | NA | NA | NA | 152_155 | NA | NA |
| 140_142 | NA | NA | 158_162 | NA | NA | 214_220 | 150_152 | NA | NA |
| 140_146 | NA | NA | 177_181 | NA | NA | 222_222 | 152_152 | NA | NA |
| 125_146 | NA | NA | 176_176 | NA | NA | 220_224 | 152_152 | NA | NA |
| 138_140 | NA | NA | 146_169 | NA | NA | 220_224 | 152_152 | NA | NA |
| 140_144 | NA | NA | 158_168 | NA | NA | 218_218 | 152_152 | NA | NA |
| 142_144 | NA | NA | 154_154 | NA | NA | 220_220 | 152_152 | NA | NA |
| 140_142 | NA | NA | 140_179 | NA | NA | 220_233 | 152_152 | NA | NA |
| 140_140 | NA | NA | 154_168 | NA | NA | 220_222 | 152_152 | NA | NA |
| 140_140 | NA | NA | 177_179 | NA | NA | 218_236 | 152_152 | NA | NA |
| 136_140 | NA | NA | 158_176 | NA | NA | 220_222 | 152_152 | NA | NA |
| 140_140 | NA | NA | 140_158 | NA | NA | 218_222 | 152_152 | NA | NA |
| 142_142 | NA | NA | 142_169 | NA | NA | 218_220 | 152_155 | NA | NA |
| 140_140 | NA | NA | 173_175 | NA | NA | 220_220 | 150_152 | NA | NA |
| 140_140 | NA | NA | 144_148 | NA | NA | 220_224 | 150_152 | NA | NA |
| 140_140 | NA | NA | 144_144 | NA | NA | 216_220 | 152_152 | NA | NA |
| 140_142 | NA | NA | 158_171 | NA | NA | 222_230 | 152_152 | NA | NA |
| 140_140 | NA | NA | 144_146 | NA | NA | 220_220 | 152_152 | NA | NA |
| 140_142 | NA | NA | 144_154 | NA | NA | 220_220 | 152_152 | NA | NA |
| 140_140 | NA | NA | 159_171 | NA | NA | 218_220 | 152_152 | NA | NA |
| 142_142 | NA | NA | 142_171 | NA | NA | 222_222 | 152_152 | NA | NA |
| 142_144 | NA | NA | 142_144 | NA | NA | 218_224 | 152_152 | NA | NA |
| 140_144 | NA | NA | 146_161 | NA | NA | 220_220 | 152_152 | NA | NA |
| 142_142 | NA | NA | 140_144 | NA | NA | 218_235 | 152_152 | NA | NA |
| 142_142 | NA | NA | 154_158 | NA | NA | 218_220 | 152_152 | NA | NA |
| 142_142 | NA | NA | 152_169 | NA | NA | 220_222 | 152_152 | NA | NA |

| | | | | | | | | | |
|---|---|---|---|---|---|---|---|---|---|
| 140_144 | NA | NA | 158_176 | NA | NA | 218_226 | 152_152 | NA | NA |
| 140_140 | NA | NA | 158_162 | NA | NA | 218_218 | 152_152 | NA | NA |
| 140_144 | NA | NA | 144_156 | NA | NA | 218_224 | 152_152 | NA | NA |
| 140_140 | NA | NA | 179_185 | NA | NA | 220_222 | 152_152 | NA | NA |
| 140_140 | NA | NA | 144_169 | NA | NA | 220_220 | 152_152 | NA | NA |
| 140_144 | NA | NA | 148_154 | NA | NA | 222_224 | 152_152 | NA | NA |
| 140_140 | NA | NA | 156_164 | NA | NA | 218_220 | 152_152 | NA | NA |
| 142_142 | NA | NA | 154_154 | NA | NA | 218_233 | 152_152 | NA | NA |
| 140_144 | NA | NA | 154_164 | NA | NA | 218_220 | 152_152 | NA | NA |
| 140_140 | NA | NA | 154_176 | NA | NA | 220_220 | 152_152 | NA | NA |
| 140_140 | NA | NA | 160_175 | NA | NA | 220_228 | 152_152 | NA | NA |
| 140_140 | NA | NA | 142_144 | NA | NA | 218_220 | 152_152 | NA | NA |
| 140_142 | NA | NA | 158_163 | NA | NA | 218_222 | 152_152 | NA | NA |
| 140_144 | NA | NA | 158_165 | NA | NA | 218_220 | 152_152 | NA | NA |
| 142_142 | 102_102 | 139_139 | 154_169 | 225_227 | 96_98 | 218_218 | 152_152 | 172_175 | 213_215 |
| 140_144 | NA | NA | 144_177 | NA | NA | 218_220 | 152_152 | NA | NA |
| 140_140 | 102_104 | 139_145 | 152_177 | 215_223 | 104_106 | 222_224 | 152_152 | 167_172 | 215_235 |
| 140_144 | 102_102 | 139_150 | 156_161 | 223_227 | 96_96 | 220_220 | 152_152 | 167_172 | 213_235 |
| 144_144 | 98_104 | 139_142 | 156_179 | 223_225 | 106_106 | 218_220 | 152_152 | 172_175 | 237_245 |
| 142_144 | 98_106 | 139_142 | 152_171 | 209_211 | 104_106 | 220_224 | 152_152 | 172_175 | 213_237 |
| 140_140 | NA | NA | 142_154 | NA | NA | 218_218 | 152_152 | NA | NA |
| 140_144 | NA | NA | 144_154 | NA | NA | 218_228 | 152_152 | NA | NA |
| 140_140 | NA | NA | 154_171 | NA | NA | 215_220 | 152_152 | NA | NA |
| 142_144 | 98_104 | 139_142 | 161_169 | 225_225 | 106_106 | 218_220 | 152_152 | 172_175 | 215_253 |
| 140_140 | 102_110 | 139_139 | 154_161 | 221_221 | 106_106 | 218_218 | 152_155 | 172_172 | 213_243 |
| 144_144 | 110_110 | 139_139 | 152_161 | 215_227 | 106_106 | 218_220 | 152_152 | 175_175 | 213_261 |
| 140_142 | NA | NA | 154_162 | NA | NA | 220_226 | 152_152 | NA | NA |
| 141_142 | 98_104 | 139_142 | 142_175 | 221_223 | 96_96 | 218_222 | 152_152 | 172_175 | 237_241 |
| 141_144 | 98_98 | 139_142 | 154_157 | 211_219 | 96_106 | 220_220 | 152_152 | 172_175 | 213_243 |
| 140_148 | 98_102 | 139_139 | 154_161 | 211_223 | 104_104 | 218_220 | 152_152 | 175_175 | 235_237 |
| 142_144 | 98_98 | 139_142 | 144_158 | 223_225 | 106_106 | 218_218 | 152_152 | 172_175 | 237_255 |
| 118_140 | NA | NA | 152_173 | NA | NA | 218_220 | 152_152 | NA | NA |
| 142_144 | 98_98 | 139_139 | 169_171 | 211_225 | 98_104 | 218_222 | 152_152 | 167_172 | 217_243 |
| 144_144 | 98_110 | 139_142 | 142_150 | 209_223 | 106_106 | 222_222 | 152_152 | 175_175 | 213_235 |
| 144_146 | 98_98 | 139_139 | 158_158 | 211_225 | 98_106 | 218_220 | 152_152 | 175_175 | 227_261 |
| 140_146 | 98_104 | 139_139 | 161_171 | 211_231 | 106_106 | 218_222 | 152_152 | 175_175 | 213_237 |
| 144_144 | 98_98 | 139_142 | 154_158 | 209_223 | 106_106 | 218_218 | 152_152 | 167_172 | 215_239 |
| 140_144 | NA | NA | 154_171 | NA | NA | 218_218 | 152_152 | NA | NA |
| 142_144 | 98_104 | 141_145 | 157_171 | 225_233 | 104_106 | 218_220 | 152_152 | 172_175 | 237_243 |
| 140_144 | 102_102 | 142_142 | 142_179 | 219_225 | 106_106 | 218_218 | 152_152 | 172_175 | 213_213 |
| 140_144 | 98_102 | 139_139 | 154_156 | 223_225 | 106_106 | 218_220 | 152_152 | 167_175 | 213_255 |
| 140_144 | NA | NA | 154_169 | NA | NA | 220_228 | 152_152 | NA | NA |
| 144_144 | NA | NA | 160_164 | NA | NA | 218_218 | 152_152 | NA | NA |
| 140_144 | 98_104 | 139_142 | 152_154 | 223_223 | 96_106 | 220_222 | 152_152 | 167_172 | 213_247 |
| 140_144 | NA | NA | 154_158 | NA | NA | 218_220 | 152_152 | NA | NA |
| 140_140 | NA | NA | 140_144 | NA | NA | 220_220 | 152_152 | NA | NA |
| 140_140 | NA | NA | 144_160 | NA | NA | 218_218 | 152_152 | NA | NA |
| 140_148 | NA | NA | 154_156 | NA | NA | 218_222 | 152_152 | NA | NA |

| | | | | | | | | | |
|---|---|---|---|---|---|---|---|---|---|
| 140_144 | NA | NA | 148_165 | NA | NA | 218_222 | 152_152 | NA | NA |
| 140_140 | NA | NA | 154_168 | NA | NA | 218_218 | 152_152 | NA | NA |
| 144_144 | NA | NA | 157_173 | NA | NA | 218_220 | 152_152 | NA | NA |
| 140_140 | NA | NA | 171_171 | NA | NA | 220_226 | 152_152 | NA | NA |
| 142_144 | NA | NA | 163_179 | NA | NA | 218_218 | 152_152 | NA | NA |
| 140_140 | NA | NA | 169_176 | NA | NA | 210_222 | 152_152 | NA | NA |
| 140_140 | NA | NA | 142_158 | NA | NA | 220_220 | 150_152 | NA | NA |
| 142_144 | NA | NA | 144_177 | NA | NA | 220_224 | 152_152 | NA | NA |
| 140_142 | NA | NA | 147_179 | NA | NA | 218_226 | 152_155 | NA | NA |
| 142_144 | NA | NA | 144_169 | NA | NA | 224_228 | 152_152 | NA | NA |
| 140_142 | NA | NA | 158_169 | NA | NA | 222_222 | 152_152 | NA | NA |
| 140_144 | NA | NA | 156_165 | NA | NA | 220_224 | 152_152 | NA | NA |
| 140_144 | NA | NA | 156_161 | NA | NA | 218_220 | 152_152 | NA | NA |
| 140_142 | NA | NA | 144_144 | NA | NA | 226_228 | 152_152 | NA | NA |
| 142_144 | NA | NA | 162_168 | NA | NA | 218_218 | 152_152 | NA | NA |
| 140_140 | NA | NA | 142_156 | NA | NA | 220_220 | 152_152 | NA | NA |
| 140_144 | NA | NA | 152_158 | NA | NA | 218_218 | 152_152 | NA | NA |
| 133_142 | NA | NA | 144_168 | NA | NA | 220_220 | 152_152 | NA | NA |
| 142_144 | NA | NA | 156_158 | NA | NA | 218_222 | 152_152 | NA | NA |
| 140_142 | NA | NA | 171_171 | NA | NA | 220_220 | 152_152 | NA | NA |
| 140_140 | NA | NA | 158_161 | NA | NA | 218_218 | 152_152 | NA | NA |
| 140_140 | NA | NA | 172_177 | NA | NA | 216_224 | 152_158 | NA | NA |
| 144_144 | NA | NA | 158_168 | NA | NA | 222_228 | 152_158 | NA | NA |
| 140_144 | NA | NA | 152_156 | NA | NA | 218_220 | 152_152 | NA | NA |
| 140_140 | NA | NA | 142_169 | NA | NA | NA | 152_152 | NA | NA |
| 140_144 | NA | NA | 150_160 | NA | NA | 218_220 | 152_152 | NA | NA |
| 144_144 | NA | NA | 152_161 | NA | NA | 218_218 | 152_152 | NA | NA |
| 140_147 | NA | NA | 162_164 | NA | NA | 218_220 | 152_152 | NA | NA |
| 140_142 | NA | NA | 144_158 | NA | NA | 220_220 | 152_152 | NA | NA |
| 140_144 | NA | NA | 142_147 | NA | NA | 218_222 | 152_152 | NA | NA |
| 140_142 | NA | NA | 147_158 | NA | NA | 218_224 | 152_152 | NA | NA |
| 140_142 | NA | NA | 156_171 | NA | NA | 218_220 | 152_152 | NA | NA |
| 140_142 | NA | NA | 154_177 | NA | NA | 220_224 | 152_152 | NA | NA |
| 140_140 | NA | NA | 171_179 | NA | NA | 220_228 | 152_152 | NA | NA |
| 140_140 | NA | NA | 160_177 | NA | NA | 224_226 | 152_152 | NA | NA |
| 140_140 | NA | NA | 142_177 | NA | NA | 214_218 | 152_152 | NA | NA |
| 140_144 | NA | NA | 156_158 | NA | NA | 218_218 | 152_152 | NA | NA |
| 140_142 | NA | NA | 154_166 | NA | NA | 218_220 | 152_152 | NA | NA |
| 140_140 | NA | NA | 144_158 | NA | NA | 218_222 | 152_152 | NA | NA |
| 142_142 | NA | NA | 144_179 | NA | NA | 218_220 | 152_152 | NA | NA |
| 140_144 | NA | NA | 179_183 | NA | NA | 218_218 | 152_152 | NA | NA |
| 144_144 | NA | NA | 142_154 | NA | NA | 218_218 | 152_152 | NA | NA |
| 140_140 | NA | NA | 148_168 | NA | NA | 212_222 | 152_152 | NA | NA |
| 142_144 | NA | NA | 146_171 | NA | NA | 220_220 | 152_152 | NA | NA |
| 140_140 | NA | NA | 142_142 | NA | NA | 220_220 | 152_152 | NA | NA |
| 140_142 | NA | NA | 171_172 | NA | NA | 220_226 | 152_152 | NA | NA |
| 144_144 | NA | NA | 150_158 | NA | NA | 218_218 | 152_152 | NA | NA |
| 144_144 | NA | NA | 144_163 | NA | NA | 218_218 | 152_152 | NA | NA |

| | | | | | | | | | |
|---|---|---|---|---|---|---|---|---|---|
| 142_142 | NA | NA | 144_148 | NA | NA | 220_220 | 152_152 | NA | NA |
| 138_144 | NA | NA | 144_159 | NA | NA | 218_220 | 152_152 | NA | NA |
| 140_140 | NA | NA | 154_158 | NA | NA | 220_220 | 152_158 | NA | NA |
| 140_142 | NA | NA | 168_181 | NA | NA | 214_228 | 152_152 | NA | NA |
| 140_144 | NA | NA | 144_160 | NA | NA | 224_224 | 152_152 | NA | NA |
| 142_142 | 98_104 | 139_142 | 144_171 | 213_221 | 104_106 | 220_220 | 152_152 | 167_175 | 213_215 |
| 140_140 | 98_106 | 139_142 | 157_157 | 229_231 | 98_106 | 218_220 | 152_152 | 172_175 | 227_229 |
| 142_144 | NA | NA | 154_173 | NA | NA | 220_224 | 152_152 | NA | NA |
| 140_142 | NA | NA | 160_166 | NA | NA | 220_224 | 152_152 | NA | NA |
| 140_142 | NA | NA | 146_154 | NA | NA | 220_220 | 152_152 | NA | NA |
| 140_144 | NA | NA | 144_164 | NA | NA | 218_220 | 152_152 | NA | NA |
| 140_144 | NA | NA | 150_156 | NA | NA | 220_220 | 152_152 | NA | NA |
| 140_140 | NA | NA | 144_144 | NA | NA | 220_220 | 152_152 | NA | NA |
| 140_144 | NA | NA | 154_172 | NA | NA | 218_224 | 152_152 | NA | NA |
| 140_140 | 102_110 | 142_149 | 158_169 | 209_213 | 106_106 | 216_220 | 150_152 | 163_172 | 210_241 |
| 140_140 | 98_106 | 139_142 | 144_154 | 213_227 | 106_106 | 218_220 | 152_152 | 175_175 | 219_263 |
| 127_144 | 98_108 | 139_150 | 163_169 | 219_223 | 104_106 | 224_224 | 152_152 | 163_172 | 213_215 |
| 142_142 | 102_102 | 142_142 | 154_177 | 227_233 | 104_104 | 220_224 | 152_155 | 167_175 | 213_213 |
| 144_144 | 98_100 | 135_139 | 144_154 | 227_229 | 100_104 | 218_220 | 152_152 | 167_172 | 213_215 |
| 140_140 | NA | NA | 154_166 | NA | NA | 220_220 | 152_152 | NA | NA |
| 144_144 | NA | NA | 142_162 | NA | NA | 218_220 | 150_152 | NA | NA |
| 140_140 | 98_104 | 135_139 | 158_177 | 225_229 | 102_104 | 218_228 | 148_152 | 172_175 | 213_237 |
| 140_142 | 98_104 | 142_142 | 144_164 | 221_227 | 106_106 | 220_230 | 152_152 | 172_172 | 213_213 |
| 140_140 | 104_108 | 139_139 | 144_158 | 217_223 | 104_106 | 220_228 | 152_152 | 175_175 | 213_219 |
| 142_142 | NA | NA | 142_175 | NA | NA | 218_226 | 152_152 | NA | NA |
| 140_142 | 98_100 | 139_150 | 175_181 | 221_229 | 96_106 | 216_235 | 152_152 | 172_172 | 210_215 |
| NA | 98_104 | 139_150 | 166_166 | 223_223 | 104_104 | NA | 148_152 | 172_175 | 215_231 |
| 140_142 | 98_104 | 139_139 | 154_181 | 207_223 | 104_106 | 220_220 | 152_152 | 167_175 | 237_239 |
| 140_146 | NA | NA | 162_168 | NA | NA | 220_235 | 152_152 | NA | NA |
| 140_144 | 98_106 | 139_139 | 156_158 | 229_233 | 104_106 | 220_224 | 152_152 | 172_175 | 215_215 |
| 140_144 | 98_102 | 139_145 | 155_172 | 209_227 | 104_104 | 218_224 | 152_152 | 167_172 | 213_249 |
| 144_146 | NA | NA | 161_162 | NA | NA | 218_220 | 152_152 | NA | NA |
| 140_144 | NA | NA | 168_169 | NA | NA | 218_235 | 152_152 | NA | NA |
| 142_144 | NA | NA | 176_179 | NA | NA | 220_226 | 152_152 | NA | NA |
| 140_142 | NA | NA | 144_158 | NA | NA | 220_228 | 152_152 | NA | NA |
| 140_142 | NA | NA | 144_154 | NA | NA | 222_224 | 152_152 | NA | NA |
| 140_144 | 98_98 | 142_142 | 147_181 | 229_229 | 96_106 | 220_220 | 152_152 | 167_172 | 215_257 |
| 140_140 | 104_104 | 139_142 | 169_169 | 225_231 | 96_106 | 218_220 | 152_152 | 172_172 | 213_215 |
| 140_140 | 102_104 | 142_142 | 169_181 | 223_227 | 106_106 | 218_220 | 152_152 | NA | 213_213 |
| 140_142 | 98_102 | 139_142 | 144_169 | 211_225 | 104_104 | 220_220 | 152_152 | 163_172 | 213_231 |
| 140_140 | 104_104 | 142_142 | 171_177 | 225_229 | 96_104 | 218_220 | 152_152 | 167_167 | 213_215 |
| NA | 98_106 | 142_142 | 156_164 | 227_229 | 104_106 | NA | 152_152 | NA | 215_215 |
| 140_140 | NA | NA | 144_146 | NA | NA | 218_230 | 152_152 | NA | NA |
| 142_142 | NA | NA | 144_181 | NA | NA | 220_220 | 152_152 | NA | NA |
| 142_142 | NA | NA | 154_161 | NA | NA | 218_218 | 152_152 | NA | NA |
| 140_144 | NA | NA | 162_171 | NA | NA | 218_222 | 152_152 | NA | NA |
| 140_142 | 102_106 | 139_139 | 144_154 | 221_231 | 106_106 | 220_226 | 152_152 | 172_172 | 213_215 |
| 140_140 | 98_98 | 142_142 | 144_166 | 223_223 | 96_106 | 218_226 | 152_152 | 167_172 | 215_233 |

| | | | | | | | | | |
|---|---|---|---|---|---|---|---|---|---|
| 140_140 | 102_104 | 139_142 | 150_154 | 223_223 | 106_106 | 218_220 | 152_152 | 172_175 | 215_237 |
| 142_144 | 98_98 | 135_142 | 173_179 | 221_223 | 104_106 | 222_226 | 152_152 | 167_175 | 213_215 |
| 140_144 | NA | NA | 164_168 | NA | NA | 222_228 | 152_152 | NA | NA |
| 140_140 | NA | NA | 144_154 | NA | NA | 218_226 | 152_152 | NA | NA |
| 140_142 | NA | NA | 169_171 | NA | NA | 220_220 | 152_152 | NA | NA |
| 140_140 | NA | NA | 154_158 | NA | NA | 218_226 | 152_152 | NA | NA |
| 134_142 | NA | NA | 175_176 | NA | NA | 218_220 | 152_152 | NA | NA |
| 140_144 | NA | NA | 154_181 | NA | NA | 220_220 | 150_152 | NA | NA |
| 140_140 | NA | NA | 152_175 | NA | NA | 224_230 | 152_152 | NA | NA |
| 140_140 | NA | NA | 158_160 | NA | NA | 222_233 | 152_152 | NA | NA |
| 140_140 | NA | NA | 155_169 | NA | NA | 220_224 | 152_152 | NA | NA |
| 138_140 | NA | NA | 144_181 | NA | NA | 218_220 | 152_152 | NA | NA |
| 140_140 | NA | NA | 148_154 | NA | NA | 220_230 | 152_152 | NA | NA |
| 140_140 | NA | NA | 148_150 | NA | NA | 218_220 | 152_152 | NA | NA |
| 140_144 | NA | NA | 154_168 | NA | NA | 222_233 | 152_152 | NA | NA |
| 134_140 | NA | NA | 146_175 | NA | NA | 220_224 | 152_152 | NA | NA |
| 144_147 | NA | NA | 154_162 | NA | NA | 218_226 | 152_152 | NA | NA |
| 140_140 | NA | NA | 169_175 | NA | NA | 218_220 | 152_152 | NA | NA |
| 142_144 | NA | NA | 144_146 | NA | NA | 218_218 | 148_152 | NA | NA |
| 140_142 | NA | NA | 154_181 | NA | NA | 218_228 | 152_152 | NA | NA |
| 142_142 | NA | NA | 144_183 | NA | NA | 220_222 | 152_152 | NA | NA |
| 134_140 | NA | NA | 144_169 | NA | NA | 220_220 | 132_152 | NA | NA |
| 140_140 | NA | NA | 169_177 | NA | NA | 222_228 | 152_152 | NA | NA |
| 140_142 | NA | NA | 144_173 | NA | NA | 220_224 | 152_152 | NA | NA |
| 140_140 | 98_104 | 142_142 | 144_171 | 221_225 | 96_106 | 216_224 | 148_152 | 172_175 | 213_225 |
| 140_142 | 98_98 | 139_139 | 150_154 | 223_225 | 104_104 | 220_235 | 152_152 | 172_175 | 215_245 |
| 140_142 | NA | NA | 180_182 | NA | NA | 218_224 | 148_152 | NA | NA |
| 140_140 | NA | NA | 168_175 | NA | NA | 218_224 | 152_152 | NA | NA |
| 140_140 | NA | NA | 168_171 | NA | NA | 224_228 | 152_152 | NA | NA |
| 140_141 | 104_106 | 142_142 | 144_183 | 227_229 | 104_104 | 218_220 | 152_152 | 167_172 | 212_212 |
| 140_140 | 98_102 | 139_139 | 173_180 | 215_221 | 102_104 | 220_220 | 152_152 | 167_172 | 213_229 |
| 140_144 | NA | NA | 144_158 | NA | NA | 218_220 | 152_152 | NA | NA |
| 140_140 | NA | NA | 154_177 | NA | NA | 218_220 | 152_152 | NA | NA |
| 140_142 | 98_106 | 139_139 | 144_177 | 221_229 | 104_106 | 220_224 | 152_152 | 174_175 | 213_213 |
| 140_140 | 98_98 | 142_145 | 165_178 | 225_227 | 102_104 | 218_220 | 148_152 | 172_175 | 215_229 |
| 140_140 | 98_104 | 142_142 | 163_177 | 225_227 | 96_106 | 218_220 | 152_152 | 172_175 | 215_229 |
| 140_140 | 98_104 | 139_139 | 144_173 | 221_227 | 106_106 | 224_228 | 152_152 | 172_172 | 215_239 |
| 140_140 | 98_106 | 139_141 | 152_152 | 227_227 | 104_108 | 220_224 | 152_152 | 172_174 | 217_233 |
| 140_146 | NA | NA | 144_144 | NA | NA | 220_220 | 152_152 | NA | NA |
| 142_142 | 102_104 | 139_139 | 169_173 | 221_221 | 104_106 | 224_228 | 152_152 | 172_175 | 213_215 |
| 140_142 | 102_104 | 139_139 | 148_168 | 227_229 | 104_106 | 222_224 | 152_152 | 172_175 | 213_233 |
| 140_144 | 102_104 | 139_142 | 144_179 | 221_221 | 104_106 | 226_228 | 152_152 | 175_175 | 213_215 |
| 140_140 | 102_102 | 139_139 | 146_177 | 231_233 | 104_104 | 224_226 | 152_152 | 167_172 | 215_233 |
| 140_140 | 104_104 | 142_142 | 146_177 | 227_235 | 104_104 | 224_226 | 152_152 | 175_175 | 213_235 |
| 140_140 | 102_102 | 139_139 | 183_185 | 221_229 | 104_106 | 222_224 | 152_152 | 167_172 | 213_213 |
| 144_146 | NA | NA | 169_179 | NA | NA | 210_218 | 152_152 | NA | NA |
| 140_140 | 102_105 | 139_142 | 169_177 | 221_227 | 104_104 | 228_228 | 152_152 | 172_175 | 213_215 |
| 140_140 | 102_106 | 139_139 | 146_175 | 221_225 | 106_106 | 224_224 | 152_152 | 175_175 | 213_213 |

| | | | | | | | | | |
|---|---|---|---|---|---|---|---|---|---|
| 140_142 | 106_128 | 139_142 | 169_177 | 229_231 | 104_104 | 224_226 | 152_152 | 172_175 | 213_213 |
| 142_142 | 102_102 | 139_139 | 146_181 | 227_229 | 106_106 | 220_222 | 152_152 | 167_175 | 213_213 |
| 140_140 | NA | NA | 144_146 | NA | NA | 218_220 | 152_152 | NA | NA |
| 140_142 | NA | NA | 144_169 | NA | NA | 220_224 | 152_152 | NA | NA |
| 140_142 | 102_104 | 142_142 | 168_169 | 221_227 | 104_106 | 220_224 | 152_152 | 172_175 | 213_213 |
| 140_142 | NA | NA | 169_175 | NA | NA | 220_220 | 152_152 | NA | NA |
| 140_140 | 98_102 | NA | 179_183 | NA | 104_106 | 218_230 | 152_152 | 172_172 | NA |
| 140_142 | 102_102 | 139_142 | 144_176 | 221_221 | 104_106 | 226_226 | 152_152 | 175_175 | 213_233 |
| 142_144 | 102_102 | 139_139 | 144_144 | 221_229 | 104_106 | 224_226 | 152_152 | 175_175 | 213_233 |
| 140_142 | 102_102 | 139_139 | 144_185 | 221_221 | 104_106 | 220_220 | 152_152 | 175_175 | 213_215 |
| 140_142 | 102_104 | 139_139 | 179_181 | 227_229 | 104_106 | 224_224 | 152_152 | 175_175 | 213_213 |
| 140_140 | 102_102 | 139_142 | 144_176 | 221_221 | 104_106 | 220_226 | 155_155 | 175_175 | 213_233 |
| 140_142 | 102_102 | 139_142 | 144_176 | 221_221 | 104_106 | 220_226 | 152_152 | 175_175 | 213_213 |
| 140_140 | NA | NA | 144_158 | NA | NA | 220_226 | 152_152 | NA | NA |
| 140_142 | 102_102 | 139_142 | 144_176 | 221_221 | 104_106 | 226_226 | 152_152 | 175_175 | 213_233 |
| 140_142 | 98_102 | 139_142 | 146_185 | 221_221 | 104_106 | 220_224 | 152_152 | 172_175 | 213_213 |
| 140_140 | 102_104 | 139_139 | 181_183 | 221_229 | 104_106 | 224_226 | 152_152 | 172_172 | 213_213 |
| 140_142 | NA | 139_139 | 144_144 | 221_229 | 106_106 | 224_226 | 152_152 | NA | 213_213 |
| 140_140 | 102_102 | 139_142 | 144_176 | 221_221 | 104_106 | 226_226 | 152_152 | 175_175 | 213_213 |
| 140_142 | 102_102 | 139_139 | 146_186 | 221_227 | 106_106 | 220_226 | 152_152 | 175_175 | 213_213 |
| 140_140 | 102_104 | 139_142 | 147_169 | 227_231 | 104_104 | 224_228 | 152_152 | 175_175 | 212_212 |
| 142_144 | NA | 138_139 | 177_177 | 221_233 | 106_106 | 220_233 | 152_152 | NA | 213_253 |
| 140_140 | NA | NA | 144_144 | NA | NA | 226_228 | 152_152 | NA | NA |
| 140_144 | 102_106 | 139_142 | 144_146 | 221_233 | 104_104 | 224_224 | 152_152 | 175_177 | 213_213 |
| 140_142 | 102_102 | 139_142 | 152_177 | 223_237 | 104_104 | 220_235 | 152_152 | 167_174 | 213_213 |
| 140_140 | NA | NA | 169_181 | NA | NA | 220_222 | 152_152 | NA | NA |
| 140_140 | 98_102 | 139_139 | 158_179 | 211_231 | 106_106 | 220_224 | 152_152 | 172_175 | 213_215 |
| 140_140 | NA | NA | 155_155 | NA | NA | 218_220 | 152_152 | NA | NA |
| NA | 98_106 | 139_142 | 144_154 | 221_227 | 106_106 | NA | 152_152 | 175_175 | 213_215 |
| NA | 98_104 | 139_142 | 144_160 | 233_235 | 104_106 | NA | 152_152 | 167_175 | 213_239 |
| 140_140 | 102_104 | 142_142 | 152_175 | 221_227 | 96_104 | 218_224 | 152_152 | 172_175 | 213_227 |
| 140_144 | 98_102 | 139_139 | 156_171 | 227_231 | 96_106 | 224_226 | 152_152 | 172_175 | 213_233 |
| 140_140 | 104_108 | 139_142 | 144_177 | 215_229 | 106_106 | 220_224 | 152_152 | 167_175 | 213_239 |
| 140_140 | 98_106 | 139_142 | 171_181 | 221_233 | 104_106 | 218_224 | 152_152 | 167_175 | 213_239 |
| 140_140 | 102_104 | 139_142 | 146_177 | 227_231 | 106_106 | 218_218 | 152_152 | 172_172 | 213_213 |
| 140_140 | 102_106 | 139_142 | 155_176 | 223_227 | 104_106 | 218_224 | 152_152 | 172_172 | 213_213 |
| 140_144 | 102_106 | 139_142 | 144_144 | 229_231 | 104_106 | 224_226 | 152_152 | 175_175 | 213_215 |
| 140_140 | 102_102 | 139_139 | 144_177 | 227_229 | 104_106 | 220_226 | 152_152 | 175_175 | 213_231 |
| 140_140 | 102_102 | 139_139 | 169_181 | 215_215 | 106_106 | 226_226 | 152_152 | 172_177 | 213_213 |
| 142_142 | 102_104 | 139_142 | 144_175 | 229_229 | 104_104 | 222_230 | 152_155 | 175_175 | 213_213 |
| 142_144 | 98_104 | 142_142 | 144_175 | 221_221 | 104_106 | 224_224 | 152_152 | 167_174 | NA |
| 140_140 | 98_98 | 142_142 | 144_155 | 227_229 | 96_106 | 224_230 | 152_152 | 172_175 | 213_213 |
| 140_142 | 98_102 | 142_142 | 144_158 | 227_227 | 104_106 | 220_235 | 152_152 | 172_172 | 213_213 |
| 140_142 | 102_102 | 139_139 | 146_155 | 227_229 | 104_106 | 224_226 | 152_152 | 172_175 | 213_213 |
| 140_142 | 104_106 | 139_139 | 144_169 | 221_223 | 106_106 | 224_224 | 152_152 | 172_175 | 213_213 |
| 144_146 | 98_98 | 142_142 | 144_175 | 227_227 | 104_104 | 220_224 | 152_152 | 172_175 | 213_213 |
| 140_144 | 98_98 | 139_139 | 142_144 | 225_227 | 98_106 | 220_233 | 152_152 | 167_175 | 213_213 |
| 140_140 | 98_98 | 139_142 | 169_171 | 213_225 | 104_106 | 220_233 | 152_152 | 172_175 | 227_264 |

| | | | | | | | | | |
|---|---|---|---|---|---|---|---|---|---|
| 140_142 | 106_106 | 139_142 | 144_171 | 215_231 | 104_104 | 228_228 | 152_152 | 172_175 | 213_215 |
| 140_142 | 102_104 | 139_139 | 144_179 | 229_231 | 104_104 | 224_226 | 152_152 | 175_175 | 213_213 |
| 140_140 | 106_108 | 139_142 | 176_181 | 221_229 | 106_106 | 224_224 | 152_152 | 175_175 | 213_213 |
| 140_140 | 98_98 | 142_142 | 144_176 | 221_231 | 104_106 | 224_226 | 152_152 | 172_175 | 213_213 |
| 140_140 | 98_106 | 139_139 | 146_168 | 211_225 | 96_106 | 216_224 | 152_152 | 172_175 | 213_217 |
| 140_140 | 98_104 | 139_142 | 173_177 | 227_227 | 106_106 | 220_226 | 152_152 | 172_175 | 213_215 |
| 140_144 | 102_106 | 139_142 | 181_181 | 221_229 | 104_104 | 220_224 | 152_152 | 175_175 | 213_227 |
| NA | 102_108 | 142_152 | 140_179 | 221_229 | 106_106 | NA | 150_152 | NA | 210_213 |
| 140_140 | 98_104 | 142_142 | 145_148 | 231_231 | 104_106 | 226_236 | 148_148 | 167_167 | 210_213 |
| 125_142 | 98_98 | 142_149 | 142_177 | 229_229 | 104_106 | 220_220 | 152_152 | 172_172 | 215_245 |
| 140_140 | NA | NA | 169_177 | NA | NA | 218_224 | 152_152 | NA | NA |
| 140_140 | 102_104 | 139_142 | 146_171 | 221_231 | 104_106 | 218_226 | 152_152 | 172_175 | 210_213 |
| 142_142 | 102_106 | 139_142 | 144_183 | 221_227 | 104_104 | 224_224 | 152_152 | 172_175 | 213_213 |
| 140_142 | 102_102 | 139_139 | 144_176 | 227_231 | 104_106 | 224_224 | 152_152 | 175_175 | 213_215 |
| 140_142 | 102_106 | 139_139 | 169_173 | 221_231 | 106_106 | 222_224 | 152_152 | 175_175 | 213_239 |
| 140_142 | 98_102 | 139_142 | 168_181 | 221_221 | 106_106 | 222_235 | 152_152 | 174_175 | 213_215 |
| 140_140 | 102_106 | 139_142 | 146_171 | 221_231 | 98_106 | 220_230 | 152_152 | 172_175 | 213_213 |
| 140_140 | 102_102 | 139_142 | 144_173 | 223_223 | 104_106 | 218_220 | 152_152 | 175_175 | 213_213 |
| 140_140 | 102_102 | 139_142 | 171_181 | 221_229 | 104_106 | 224_224 | 152_152 | 175_175 | 213_233 |
| 140_140 | 102_104 | 139_142 | 173_179 | 219_227 | 104_104 | 220_224 | 152_152 | 167_175 | 213_213 |
| 140_140 | 102_106 | 138_142 | 168_177 | 215_225 | 106_106 | 218_220 | 152_152 | 172_175 | 215_239 |
| 140_142 | 102_104 | 139_150 | 144_152 | 223_231 | 106_108 | 220_222 | 152_152 | 172_175 | 215_215 |
| 142_144 | 98_102 | 139_142 | 150_177 | 211_223 | 106_106 | 220_220 | 152_152 | 164_164 | 213_231 |
| 140_140 | 98_102 | 139_142 | 173_181 | 223_227 | 104_106 | 218_220 | 152_152 | 172_172 | 210_241 |
| 140_144 | 102_104 | 139_139 | 171_177 | 229_231 | 104_106 | 220_224 | 149_152 | 172_172 | 213_245 |
| 140_142 | 102_102 | 139_139 | 154_181 | 227_227 | 104_106 | 220_226 | 152_152 | 175_175 | 213_213 |
| 140_140 | NA | NA | 144_154 | NA | NA | 220_220 | 152_152 | NA | NA |
| 140_142 | 102_104 | 139_142 | 175_176 | 221_225 | 104_104 | 220_226 | 152_152 | 170_172 | 215_233 |
| 140_140 | 102_104 | 142_142 | 155_181 | 229_233 | 104_106 | 220_224 | 152_152 | 167_175 | 213_215 |
| 140_142 | 100_130 | 139_139 | 164_183 | 215_221 | 104_104 | 224_226 | 152_152 | 175_175 | 231_233 |
| 144_144 | 104_104 | 139_145 | 144_160 | 223_225 | 106_106 | 218_224 | 152_152 | 175_175 | 215_249 |
| 140_144 | 98_102 | 142_142 | 144_181 | 221_227 | 96_106 | 220_224 | 152_152 | 172_175 | 213_213 |
| 140_140 | 98_102 | 139_139 | 144_156 | 215_221 | 96_106 | 218_220 | 152_152 | NA | 213_215 |
| 140_140 | 102_104 | 139_142 | 146_175 | 227_229 | 104_106 | 222_226 | 152_152 | 172_175 | 213_213 |
| NA | 98_102 | 139_150 | 166_168 | 213_229 | 96_106 | NA | 150_152 | 167_172 | 213_231 |
| NA | 98_102 | 139_139 | 171_182 | 223_225 | 106_106 | NA | 152_152 | 163_175 | 213_229 |
| 140_144 | NA | NA | 158_158 | NA | NA | 218_218 | 152_152 | NA | NA |
| 140_144 | NA | NA | 154_169 | NA | NA | 218_226 | 152_152 | NA | NA |
| 140_140 | 102_104 | 142_142 | 175_179 | 227_231 | 104_106 | 222_226 | 152_152 | 167_172 | 213_213 |
| 140_140 | 98_98 | 139_142 | 146_177 | 211_231 | 106_106 | 220_226 | 148_152 | 167_175 | 215_217 |
| 140_140 | NA | NA | 175_185 | NA | NA | 220_220 | 152_152 | NA | NA |
| 140_140 | 98_102 | 139_142 | 155_171 | 219_231 | 96_104 | 224_226 | 152_152 | 175_175 | 213_213 |
| 140_140 | 100_104 | 139_139 | 171_181 | 227_227 | 106_106 | 218_224 | 152_152 | 172_175 | 215_249 |
| 140_144 | 98_98 | 139_142 | 142_171 | 223_223 | 96_106 | 220_220 | 152_152 | 175_177 | 219_235 |
| 140_147 | 98_104 | 139_142 | 144_171 | 223_227 | 96_106 | 218_220 | 152_152 | 167_172 | 213_217 |
| 140_142 | 102_102 | 142_142 | 144_171 | 221_227 | 106_106 | 220_226 | 152_152 | 172_172 | 213_215 |
| 140_140 | 98_98 | 135_139 | 150_169 | 213_227 | 102_106 | 210_228 | 152_152 | 167_175 | 215_235 |
| 140_144 | 102_106 | 139_150 | 160_175 | 213_213 | 106_108 | 226_235 | 152_152 | 167_172 | 213_251 |

| | | | | | | | | | |
|---|---|---|---|---|---|---|---|---|---|
| NA | 98_104 | 139_145 | 144_164 | 223_229 | 96_106 | NA | 152_152 | 164_172 | 215_217 |
| NA | 98_106 | 139_142 | 144_158 | 225_229 | 96_106 | NA | 152_152 | 164_172 | 213_217 |
| 140_140 | 98_102 | 139_139 | 163_171 | 225_237 | 106_106 | 218_220 | 152_152 | 175_175 | 213_253 |
| 140_140 | 98_104 | 139_142 | 154_168 | 221_233 | 96_96 | 218_226 | 150_152 | 167_175 | 213_215 |
| 140_140 | NA | NA | 169_179 | NA | NA | 218_220 | 152_152 | NA | NA |
| NA | 102_106 | 142_142 | 142_160 | 221_225 | 106_106 | NA | 152_152 | 167_175 | 213_229 |
| 140_142 | 102_116 | 139_142 | 144_179 | 225_229 | 104_104 | 226_226 | 152_152 | NA | 213_227 |
| 140_142 | NA | NA | 156_156 | NA | NA | 220_222 | 152_152 | NA | NA |
| 140_140 | 98_106 | 139_142 | 146_183 | 223_227 | 106_106 | 220_228 | 152_152 | 175_175 | 213_213 |
| NA | 98_102 | 139_142 | 144_150 | 223_225 | 106_106 | NA | 152_155 | 167_172 | 213_215 |
| 142_144 | 98_104 | 142_142 | 156_164 | 221_239 | 106_106 | 220_230 | 152_152 | 167_172 | 213_215 |
| NA | 104_106 | 142_142 | 146_181 | 221_223 | 106_106 | NA | 152_152 | 167_172 | 215_235 |
| 140_142 | 98_100 | 139_142 | 156_158 | 225_231 | 106_106 | 218_220 | 152_152 | 172_175 | 210_213 |
| 140_140 | NA | NA | 144_177 | NA | NA | 218_220 | 152_152 | NA | NA |
| 140_140 | 98_104 | 139_142 | 171_175 | 221_223 | 106_106 | 218_218 | 152_152 | NA | 215_217 |
| 140_140 | NA | NA | 158_174 | NA | NA | 218_230 | 152_152 | NA | NA |
| 140_140 | NA | NA | 152_177 | NA | NA | 218_220 | 152_152 | NA | NA |
| 140_146 | 100_102 | 139_142 | 171_179 | 209_233 | 104_104 | 218_224 | 152_152 | 175_175 | 215_217 |
| 140_140 | 98_98 | 142_142 | 142_169 | 209_209 | 106_108 | 220_220 | 152_152 | 167_175 | 213_215 |
| 127_140 | NA | NA | 168_177 | NA | NA | 214_220 | 150_152 | NA | NA |
| NA | 98_104 | 139_139 | 164_177 | 223_223 | 96_106 | NA | 147_150 | 172_175 | 217_217 |
| 140_142 | 102_106 | 139_139 | 169_171 | 223_227 | 106_106 | 218_220 | 152_152 | 172_175 | 213_213 |
| NA | 106_106 | 139_139 | 140_142 | 223_223 | 104_104 | NA | 152_152 | 175_175 | 213_241 |
| 140_140 | 98_102 | 139_142 | 168_179 | 211_221 | 106_106 | 218_222 | 152_152 | 167_172 | 213_233 |
| 140_146 | 102_104 | 142_145 | 140_179 | 223_223 | 104_104 | 220_224 | 152_152 | 167_175 | 213_231 |
| 140_144 | 102_102 | 139_139 | 146_156 | 227_229 | 106_106 | 218_220 | 152_152 | 172_175 | 217_235 |
| 142_142 | 100_104 | 142_142 | 155_177 | 211_225 | 106_106 | 218_224 | 152_152 | 167_175 | 213_213 |
| 144_146 | 98_104 | 139_139 | 154_175 | 211_221 | 104_106 | 220_220 | 152_152 | NA | 213_213 |
| 142_147 | 98_116 | 139_142 | 172_180 | 225_237 | 96_106 | 218_220 | 152_152 | 172_175 | 217_217 |
| 140_142 | 98_104 | 142_142 | 168_169 | 211_223 | 96_106 | 218_218 | 152_152 | 172_175 | 213_215 |
| 140_140 | 98_104 | 139_139 | 169_177 | 227_229 | 96_106 | 218_222 | 152_152 | 167_175 | 217_231 |
| NA | 98_98 | 139_142 | 146_173 | 225_227 | 106_106 | NA | 147_152 | 172_174 | 213_247 |
| 140_148 | 102_102 | 139_142 | 144_144 | 223_227 | 104_104 | 220_224 | 152_152 | 163_175 | 213_213 |
| 140_144 | NA | NA | 169_175 | NA | NA | 226_226 | 152_152 | NA | NA |
| 140_144 | 98_104 | 139_142 | 150_158 | 225_225 | 96_98 | 216_218 | 152_152 | 172_175 | 215_235 |
| 140_140 | 104_104 | 139_142 | 144_174 | 223_233 | 104_106 | 220_220 | 152_152 | 167_175 | 213_231 |
| 140_141 | NA | NA | 142_173 | NA | NA | 224_230 | 152_152 | NA | NA |
| 138_142 | 98_104 | 139_142 | 142_143 | 223_227 | 96_104 | 220_220 | 152_152 | 167_175 | 213_213 |
| 140_140 | 98_102 | 139_142 | 150_168 | 223_227 | 104_106 | 218_220 | 152_152 | 172_175 | 213_215 |
| 142_144 | 98_106 | 139_139 | 150_162 | 223_225 | 96_106 | 220_226 | 152_152 | NA | 213_251 |
| 142_142 | NA | NA | 144_152 | NA | NA | 218_218 | 152_152 | NA | NA |
| 140_144 | 104_106 | 139_139 | 162_172 | 221_223 | 96_106 | 210_220 | 152_152 | 174_175 | 213_215 |
| 140_142 | NA | 139_145 | 146_169 | 223_223 | 106_106 | 220_220 | 152_155 | NA | 213_213 |
| 140_144 | 98_104 | 139_142 | 144_175 | 221_233 | 96_106 | 218_228 | 152_152 | 167_167 | 213_213 |
| 140_144 | 104_104 | 142_142 | 168_169 | 209_225 | 106_106 | 218_220 | 152_152 | 167_172 | 213_213 |
| 140_140 | 102_104 | 139_142 | 169_174 | 221_223 | 106_106 | 220_220 | 152_152 | 172_175 | 215_215 |
| 140_142 | 98_98 | 142_142 | 150_154 | 211_225 | 104_106 | 220_220 | 152_152 | 167_175 | 213_247 |
| 142_142 | 98_104 | 139_139 | 157_171 | 223_231 | 104_108 | 220_222 | 152_152 | 175_175 | 213_251 |

| AP019 | AP043 | AP049 | AP068 | AP218 | AP223 | AP226 | AP238 | AP243 | AP249 |
|---|---|---|---|---|---|---|---|---|---|
| 137_144 | 141_144 | 141_141 | 155_157 | 114_114 | 170_179 | 238_238 | 264_264 | 254_254 | 210_217 |
| 137_144 | 141_144 | 141_141 | 157_163 | 114_118 | 179_179 | 238_238 | 264_264 | 254_254 | 217_217 |
| 137_141 | 144_146 | 129_141 | 157_157 | 114_114 | 179_179 | 238_238 | 264_264 | 254_254 | 210_217 |
| 137_141 | 144_144 | 141_141 | 157_157 | 114_114 | 179_179 | 238_238 | 264_264 | 254_254 | 210_217 |
| 137_137 | 144_146 | 129_141 | 157_157 | 114_114 | 179_179 | 238_238 | 264_264 | 254_254 | 210_210 |
| 137_137 | 144_146 | 129_141 | 157_157 | 114_114 | 179_179 | 238_238 | 264_264 | 254_254 | 210_210 |
| 137_137 | 144_144 | 141_141 | 157_157 | 114_120 | 179_179 | 238_238 | 264_264 | 254_254 | 210_210 |
| 137_137 | 141_144 | 141_141 | 155_157 | 114_114 | 179_179 | 238_238 | 264_264 | 254_254 | 217_217 |
| 137_137 | 144_144 | 141_141 | 157_157 | 114_114 | 170_179 | 238_238 | 264_264 | 254_261 | 210_217 |
| 137_140 | 141_144 | 141_141 | 155_157 | 114_120 | 179_179 | 232_238 | 264_272 | 254_254 | 210_210 |
| 137_140 | 141_144 | 141_141 | 157_163 | 114_114 | 170_179 | 236_245 | 264_264 | 254_261 | 210_217 |
| 137_140 | 144_144 | 141_141 | 157_157 | 114_114 | 179_179 | 238_238 | 264_264 | 254_254 | 210_210 |
| 137_140 | 144_144 | 123_141 | 153_155 | 114_114 | 170_179 | 236_236 | 264_264 | 254_254 | 210_210 |
| 137_145 | 136_136 | 141_141 | 151_155 | 114_114 | 170_179 | 238_238 | 264_272 | 254_254 | 210_210 |
| 136_145 | 136_136 | 129_141 | 153_157 | 114_114 | 170_179 | 238_238 | 264_272 | 254_254 | 210_210 |
| 137_140 | 144_144 | 141_141 | 155_155 | 114_114 | 179_179 | 232_238 | 264_272 | 254_254 | 210_210 |
| 137_140 | 144_144 | 141_141 | 155_157 | 114_114 | 179_179 | 232_238 | 264_264 | 254_254 | 210_210 |
| 140_146 | 136_136 | 141_141 | 155_157 | 114_114 | 170_170 | 238_245 | 264_264 | 254_254 | 210_217 |
| 136_137 | 136_144 | 153_153 | 157_165 | 114_120 | 170_176 | 232_238 | 264_266 | 254_254 | 210_210 |
| 137_142 | 136_141 | 141_141 | 155_157 | 114_114 | 170_179 | 232_236 | 264_264 | 254_261 | 210_217 |
| 137_137 | 136_141 | 141_141 | 155_155 | 114_114 | 179_179 | 232_232 | 264_266 | 254_267 | 210_210 |
| 137_142 | 136_150 | 141_141 | 157_157 | 114_114 | 170_179 | 236_238 | 264_264 | 254_261 | 210_210 |
| 137_137 | 136_144 | 141_141 | 157_165 | 114_120 | 179_179 | 236_236 | 264_266 | 254_254 | 210_210 |
| 136_137 | 141_144 | 141_141 | 151_155 | 114_114 | 170_179 | 238_245 | 264_264 | 254_254 | 210_210 |
| 137_144 | 136_144 | 141_141 | 155_157 | 114_114 | 170_176 | 236_238 | 264_264 | 254_254 | 210_217 |
| 136_144 | 138_141 | 123_123 | 151_157 | 114_120 | 170_179 | 236_238 | 264_264 | 254_270 | 210_210 |
| 137_137 | 136_144 | 141_144 | 151_155 | 114_120 | 176_179 | 232_238 | 264_264 | 254_254 | 210_217 |
| 137_142 | 141_144 | 141_144 | 151_157 | 114_120 | 179_179 | 234_236 | 264_264 | 254_254 | 217_217 |
| 140_145 | 141_144 | 141_141 | 151_155 | 114_120 | 179_179 | 234_238 | 264_264 | 254_254 | 217_217 |
| 137_137 | 134_144 | 141_141 | 159_165 | 114_114 | 179_179 | 232_238 | 264_264 | 254_254 | 210_210 |
| 137_137 | 134_141 | 141_141 | 155_157 | 114_114 | 179_179 | 232_238 | 264_264 | 254_254 | 210_210 |
| 136_137 | 134_144 | 129_141 | 155_157 | 114_114 | 179_179 | 238_238 | 264_264 | 254_254 | 210_210 |
| 137_137 | 134_144 | 129_141 | 157_163 | 114_114 | 179_179 | 238_238 | 264_264 | 254_254 | 210_219 |
| 137_137 | 141_144 | 141_141 | 151_157 | 114_114 | 170_179 | 232_238 | 264_264 | 254_254 | 217_217 |
| 137_141 | 144_144 | 141_141 | 151_157 | 114_114 | 179_184 | 232_236 | 264_264 | 254_254 | 210_217 |
| 137_144 | 141_146 | 141_141 | 151_157 | 114_114 | 179_179 | 232_238 | 264_264 | 254_254 | 210_217 |
| 137_141 | 141_144 | 141_141 | 151_157 | 114_114 | 179_179 | 232_236 | 264_264 | 254_254 | 210_217 |
| 137_144 | 144_146 | NA | 151_163 | 114_114 | 179_179 | 232_238 | 264_264 | 255_255 | 210_210 |
| 135_137 | 136_171 | 132_132 | 155_157 | 114_114 | 179_184 | 238_245 | 266_266 | 254_254 | 217_219 |
| 137_137 | 141_141 | 132_132 | 157_165 | 114_114 | 170_184 | 238_245 | 264_266 | 254_254 | 220_220 |
| 137_137 | NA | 132_141 | NA | NA | NA | NA | 266_266 | 254_257 | NA |
| 137_142 | NA | 132_132 | NA | NA | NA | NA | 264_266 | 254_254 | NA |
| NA | NA | 132_141 | NA | NA | NA | NA | 264_264 | 254_254 | NA |
| 137_142 | 138_141 | 132_132 | 153_155 | 114_114 | 179_184 | 245_245 | 266_266 | 254_254 | 210_220 |
| 137_137 | NA | 132_132 | NA | NA | NA | NA | 264_266 | 254_254 | NA |
| 137_137 | 168_171 | 132_132 | 155_157 | 114_122 | 179_184 | 245_248 | 264_264 | 254_254 | 218_225 |
| 137_140 | 146_171 | 132_132 | 157_157 | 114_122 | 179_184 | 245_245 | 264_266 | 254_254 | 212_220 |
| 137_137 | NA | 132_132 | NA | NA | NA | NA | 266_266 | 254_254 | NA |

| | | | | | | | | | |
|---|---|---|---|---|---|---|---|---|---|
| 137_140 | 134_170 | 141_141 | 155_157 | 114_114 | 179_184 | 236_245 | 264_264 | 254_254 | 210_220 |
| 140_140 | 136_141 | 132_141 | 155_161 | 114_114 | 179_184 | 236_245 | 264_264 | 254_254 | 210_225 |
| 137_137 | NA | 132_132 | NA | NA | NA | NA | 264_266 | 254_270 | NA |
| 137_137 | NA | NA | NA | NA | NA | NA | 264_266 | 254_254 | NA |
| 137_137 | 144_171 | 141_141 | 165_165 | 114_122 | 179_182 | 236_236 | 264_266 | 254_254 | 210_225 |
| 137_137 | 141_166 | 132_141 | NA | 114_122 | NA | NA | 266_272 | 254_254 | 209_220 |
| 137_137 | 141_170 | 123_141 | 153_155 | 114_122 | 170_186 | 245_245 | 264_266 | 254_254 | 210_220 |
| 137_140 | 144_173 | 132_141 | 157_157 | 114_122 | 179_184 | 245_245 | 266_266 | 254_254 | 220_220 |
| 137_137 | 141_141 | 132_132 | 155_157 | 114_122 | 184_184 | 245_245 | 266_266 | 254_254 | 220_225 |
| 137_137 | NA | 132_132 | NA | NA | NA | NA | 264_266 | 254_254 | NA |
| 137_137 | 141_141 | 132_141 | 155_157 | 114_122 | 179_184 | 245_245 | 264_266 | 254_254 | 220_225 |
| 137_137 | 141_168 | 132_141 | 157_157 | 114_122 | 179_184 | 245_245 | 264_264 | 254_254 | 220_220 |
| 137_137 | NA | 123_132 | NA | NA | NA | NA | 264_266 | 254_254 | NA |
| 137_137 | NA | 132_141 | NA | NA | NA | NA | 266_266 | 254_254 | NA |
| 137_137 | 134_141 | 132_141 | 157_157 | 114_122 | 184_184 | 238_238 | 264_264 | 254_254 | 220_225 |
| 140_142 | 144_170 | 132_141 | 155_157 | 114_114 | 170_179 | 236_245 | 264_266 | 254_254 | 218_220 |
| 137_137 | 136_171 | 141_141 | 155_157 | 114_122 | 170_184 | 245_245 | 266_266 | 245_254 | 220_220 |
| 140_140 | 141_144 | 132_132 | 153_157 | 114_122 | 170_179 | 245_245 | 264_266 | 254_254 | 210_220 |
| 137_137 | NA | 135_138 | NA | NA | NA | NA | 264_264 | 254_254 | NA |
| 137_140 | 144_144 | 123_141 | 155_157 | 114_122 | 179_179 | 238_238 | 264_266 | 254_254 | 210_218 |
| 137_137 | 144_144 | 132_141 | 157_157 | 114_114 | 179_184 | 236_245 | 264_264 | NA | 210_210 |
| 137_137 | NA | 132_132 | NA | NA | NA | NA | 266_266 | 254_254 | NA |
| 137_137 | NA | 132_132 | NA | NA | NA | NA | 266_266 | 254_254 | NA |
| 137_137 | NA | 132_132 | NA | NA | NA | NA | 266_266 | 254_269 | NA |
| 137_137 | NA | 132_132 | NA | NA | NA | NA | 266_266 | 254_254 | NA |
| 136_142 | 138_148 | 141_141 | 155_161 | 114_122 | 184_184 | 238_245 | 264_266 | 254_254 | 218_220 |
| 137_137 | NA | 132_132 | NA | NA | NA | NA | 266_266 | 254_269 | NA |
| 137_140 | 141_144 | 132_141 | 157_157 | 114_122 | 184_184 | 245_245 | 264_264 | 254_254 | 210_217 |
| 140_140 | 141_144 | 132_166 | 155_157 | 114_114 | 179_179 | 238_245 | 264_266 | 254_254 | 210_220 |
| 137_137 | NA | 132_141 | NA | NA | NA | NA | 266_266 | 254_254 | NA |
| 137_140 | 134_141 | 132_132 | 155_155 | 114_114 | 179_184 | 245_245 | 264_266 | 254_254 | 220_225 |
| 137_140 | 141_141 | 123_123 | 157_165 | 114_114 | 184_184 | 245_245 | 264_266 | 254_254 | 210_222 |
| 137_137 | 144_144 | 141_141 | NA | 114_114 | NA | NA | 264_266 | 254_254 | 220_225 |
| 137_137 | NA | 132_132 | NA | NA | NA | NA | 266_266 | 254_270 | NA |
| 137_142 | 138_144 | 132_141 | 157_161 | 114_114 | 184_186 | 245_245 | 266_266 | 254_254 | 220_220 |
| 137_137 | NA | 132_141 | NA | NA | NA | NA | 264_266 | 254_254 | NA |
| 137_137 | 144_144 | 123_141 | 155_157 | 114_122 | 179_179 | 238_245 | 264_266 | 254_254 | 210_222 |
| 137_137 | 144_144 | 141_153 | 157_157 | 114_114 | 179_184 | 238_245 | 264_264 | 254_254 | 210_218 |
| 137_140 | 141_144 | 132_141 | 157_157 | 114_114 | 184_184 | 245_245 | 266_266 | 254_254 | 210_220 |
| 137_137 | NA | 132_141 | NA | NA | NA | NA | 266_266 | 254_254 | NA |
| 137_142 | 141_141 | 132_141 | 157_161 | 114_122 | 179_184 | 236_238 | 266_266 | 254_254 | 220_225 |
| 137_140 | 134_146 | 141_141 | 151_155 | 114_114 | 179_179 | 236_245 | 264_264 | 254_254 | 210_210 |
| 140_142 | 141_146 | 132_132 | 155_161 | 114_122 | 179_184 | 236_245 | 264_264 | 254_254 | 210_220 |
| 140_140 | 138_141 | 141_141 | 149_149 | 122_122 | 170_179 | 238_245 | 264_266 | 254_254 | 220_222 |
| 137_140 | 141_141 | 132_132 | 157_159 | 114_122 | 170_179 | 238_245 | 264_266 | 254_254 | 210_218 |
| 137_140 | 138_146 | 132_141 | 157_157 | 114_114 | 179_179 | 245_245 | 264_266 | 254_266 | 210_225 |
| 136_142 | 141_144 | 132_141 | 155_155 | 114_122 | 179_179 | 236_238 | 264_266 | 254_254 | 220_225 |
| 140_140 | 136_136 | 132_132 | 155_157 | 114_114 | 179_184 | 236_245 | 264_266 | 254_254 | 210_217 |

| | | | | | | | | | |
|---|---|---|---|---|---|---|---|---|---|
| 137_137 | NA | 132_141 | 157_157 | 114_114 | 179_184 | 245_245 | 264_266 | 254_254 | 210_218 |
| 137_140 | 138_138 | NA | 161_165 | 114_114 | 179_179 | 245_245 | NA | NA | 220_225 |
| 137_137 | 141_141 | 132_132 | 155_157 | 114_114 | 184_184 | 238_245 | 264_266 | 254_254 | 210_210 |
| 137_140 | 136_144 | 132_132 | 155_157 | 114_122 | 179_184 | 238_245 | 264_266 | 254_254 | 218_220 |
| 137_140 | 138_138 | 132_141 | 157_159 | 114_114 | 170_179 | 245_248 | 264_266 | 254_254 | 218_220 |
| 136_137 | 144_187 | 141_141 | 153_157 | 114_122 | 179_184 | 238_245 | 264_266 | 254_254 | 210_220 |
| 136_140 | 134_144 | 132_132 | 155_155 | 114_114 | 179_179 | 238_238 | 264_264 | 254_254 | 210_220 |
| 140_142 | NA | 132_135 | NA | NA | NA | NA | 266_272 | 254_254 | NA |
| 137_137 | NA | 132_132 | NA | NA | NA | NA | 264_266 | 254_254 | NA |
| 137_137 | 143_181 | 141_141 | 157_157 | 114_114 | 179_184 | 245_245 | 264_266 | 254_254 | 210_218 |
| 137_140 | 144_144 | 133_141 | 157_159 | 114_114 | 179_186 | 236_236 | 264_266 | 254_254 | 220_220 |
| 137_137 | NA | 132_132 | NA | NA | NA | NA | 266_272 | 254_254 | NA |
| 137_140 | 141_144 | 132_132 | 153_157 | 114_114 | 179_184 | 236_245 | 264_266 | 254_254 | 218_220 |
| 136_137 | 144_144 | 153_153 | 157_157 | 114_114 | 170_179 | 238_245 | 264_264 | 254_254 | 210_220 |
| 140_140 | 141_144 | 132_141 | 155_157 | 114_114 | 179_184 | 236_245 | 266_266 | 254_254 | 210_218 |
| 136_137 | 134_144 | 141_141 | 157_157 | 114_114 | 179_184 | 238_245 | 264_264 | 254_254 | 218_225 |
| 137_137 | 141_141 | 132_141 | 157_159 | 114_114 | 179_184 | 238_245 | 266_266 | 254_254 | 210_218 |
| 137_140 | 144_144 | 132_141 | 153_157 | 114_114 | 179_184 | 236_245 | 264_264 | 254_254 | 217_220 |
| 137_140 | 144_144 | 132_159 | 155_157 | 114_114 | 170_184 | 238_245 | 266_266 | 254_270 | 218_220 |
| 137_137 | 141_144 | 132_132 | 155_157 | 114_114 | 179_184 | 245_245 | 266_266 | 254_254 | 210_220 |
| 137_140 | 146_185 | 132_141 | 157_159 | 114_114 | 179_179 | 238_245 | 264_266 | 254_254 | 220_225 |
| 137_140 | 136_141 | 132_132 | 157_159 | 114_114 | 184_184 | 238_238 | 265_266 | 254_254 | 210_220 |
| 137_137 | NA | 132_132 | NA | NA | NA | NA | 266_266 | 254_254 | NA |
| 137_140 | 138_146 | 132_132 | 155_157 | 114_122 | 184_184 | 238_245 | 266_266 | 254_254 | 220_225 |
| 136_137 | 144_144 | 132_132 | 155_155 | 114_114 | 179_179 | 236_245 | 264_266 | 254_254 | 210_220 |
| 140_142 | 136_181 | 132_141 | 157_159 | 122_122 | 179_184 | 238_245 | 266_266 | 254_254 | 218_225 |
| 137_137 | 134_171 | 132_132 | 155_157 | 114_114 | 184_184 | 245_245 | 264_264 | 254_254 | 220_220 |
| 137_140 | 138_144 | 132_132 | 155_157 | 114_122 | 179_179 | 238_245 | 266_266 | 254_254 | 210_210 |
| 140_142 | 134_144 | 132_132 | 157_157 | 114_114 | 179_184 | 245_245 | 266_266 | 254_254 | 220_220 |
| 137_137 | NA | 132_141 | NA | NA | NA | NA | 264_266 | 254_270 | NA |
| 137_137 | 141_144 | 132_141 | 157_157 | 114_122 | 179_184 | 236_245 | 264_266 | 254_254 | 210_218 |
| 137_137 | NA | 132_141 | NA | NA | NA | NA | 264_266 | 254_254 | NA |
| 137_142 | NA | 132_141 | 157_157 | 122_122 | 179_184 | 236_245 | 264_264 | 254_254 | 220_225 |
| 137_137 | 141_141 | 123_141 | 155_157 | 114_114 | 184_184 | 236_245 | 264_264 | 254_254 | 210_218 |
| 133_137 | NA | 132_141 | NA | NA | NA | NA | 266_266 | 254_254 | NA |
| 137_137 | 144_170 | 132_141 | 157_157 | 114_122 | 170_170 | 245_245 | 266_266 | 254_254 | 210_225 |
| 137_137 | 141_141 | 132_132 | 155_157 | 114_122 | 179_184 | 245_245 | 266_272 | 254_254 | 218_220 |
| 137_137 | NA | 116_132 | NA | NA | NA | NA | 264_266 | 254_254 | NA |
| 137_142 | 138_141 | 132_132 | 157_157 | 114_122 | 179_184 | 245_245 | 266_266 | 254_254 | 218_220 |
| 137_140 | 134_141 | 132_141 | 155_157 | 114_122 | 179_179 | 236_236 | 264_266 | 254_270 | 210_220 |
| 137_142 | NA | 132_141 | NA | NA | NA | NA | 264_266 | 254_254 | NA |
| 137_137 | 141_200 | 132_141 | 157_161 | 114_122 | 170_184 | 245_245 | 264_264 | 254_254 | 218_218 |
| 137_137 | 171_195 | 132_132 | 155_161 | 114_114 | 179_184 | 238_238 | 264_264 | 254_270 | 220_227 |
| 137_137 | NA | 132_132 | NA | NA | NA | NA | 266_266 | 254_254 | NA |
| 137_142 | NA | 123_132 | NA | NA | NA | NA | 266_266 | 254_254 | NA |
| 137_137 | 136_141 | 132_132 | 157_157 | 114_114 | 179_184 | 245_245 | 264_266 | 254_254 | 220_220 |
| 137_137 | 141_171 | 132_132 | 155_157 | 114_114 | 170_184 | 245_245 | 266_266 | 254_254 | 220_220 |
| 137_137 | 141_183 | 123_132 | 157_159 | 114_122 | 179_184 | 245_245 | 266_266 | 254_254 | 210_220 |

| | | | | | | | | | |
|---|---|---|---|---|---|---|---|---|---|
| 137_137 | 141_144 | 141_156 | 157_159 | 114_114 | 184_184 | 245_245 | 264_266 | 254_254 | 210_218 |
| 136_137 | 144_146 | 132_132 | 155_157 | 114_114 | 170_179 | 245_245 | 264_266 | 254_254 | 220_225 |
| 137_137 | 141_173 | 132_132 | 155_155 | 114_122 | 179_184 | 245_245 | 264_266 | 254_270 | 218_218 |
| 137_137 | 138_171 | 132_132 | 157_163 | 114_114 | 170_184 | 245_245 | 266_266 | 254_254 | 220_220 |
| 137_137 | 141_173 | 132_132 | 157_157 | 114_114 | 170_184 | 245_245 | NA | 254_254 | 220_225 |
| 137_137 | NA | 132_132 | NA | NA | NA | NA | 266_266 | 254_254 | NA |
| 137_137 | NA | 132_132 | NA | NA | NA | NA | 266_266 | 254_254 | NA |
| 137_137 | NA | 132_132 | NA | NA | NA | NA | 264_266 | 254_254 | NA |
| 137_137 | NA | 126_132 | NA | NA | NA | NA | 264_266 | 254_254 | NA |
| 137_137 | NA | 132_132 | NA | NA | NA | NA | 264_266 | 254_254 | NA |
| 137_140 | 138_185 | 132_141 | 157_159 | 114_114 | 184_184 | 245_245 | 266_272 | 254_254 | 220_220 |
| 137_141 | NA | 132_132 | NA | NA | NA | NA | 266_266 | 254_266 | NA |
| 137_137 | NA | 132_141 | NA | NA | NA | NA | 266_266 | 254_254 | NA |
| 137_140 | NA | 132_132 | NA | NA | NA | NA | 264_266 | 254_270 | NA |
| 137_137 | NA | 123_132 | NA | NA | NA | NA | 266_266 | 254_266 | NA |
| 137_137 | NA | 132_132 | NA | NA | NA | NA | 264_266 | 254_254 | NA |
| 137_137 | NA | NA | NA | NA | NA | NA | 264_264 | 254_254 | NA |
| 137_137 | NA | 126_132 | NA | NA | NA | NA | 266_266 | 254_254 | NA |
| 137_137 | NA | 132_132 | NA | NA | NA | NA | 266_266 | 254_254 | NA |
| 137_140 | 173_173 | 123_132 | 156_156 | 114_114 | 170_186 | 245_245 | 264_266 | 254_254 | 220_225 |
| 137_142 | NA | 123_123 | NA | NA | NA | NA | 266_272 | 254_254 | NA |
| 137_142 | 141_175 | 132_141 | 157_157 | 114_114 | 184_184 | 236_245 | 266_266 | 254_254 | 218_225 |
| 137_137 | NA | 132_132 | NA | NA | NA | NA | 264_266 | 254_254 | NA |
| 137_137 | NA | 132_135 | NA | NA | NA | NA | 264_266 | 254_254 | NA |
| 137_137 | NA | 132_132 | NA | NA | NA | NA | 264_266 | 254_254 | NA |
| 137_137 | NA | 132_132 | NA | NA | NA | NA | 266_266 | 254_254 | NA |
| 137_140 | NA | 132_141 | NA | NA | NA | NA | 264_266 | 254_254 | NA |
| 137_140 | NA | 123_132 | NA | NA | NA | NA | 266_266 | 254_254 | NA |
| 137_142 | 136_136 | 132_132 | 157_161 | 114_122 | 170_184 | 245_245 | 264_266 | 254_254 | 220_220 |
| 137_137 | 144_195 | 132_132 | 157_157 | 114_122 | 179_184 | 245_245 | 264_266 | 254_270 | 220_225 |
| 137_137 | 138_141 | 132_132 | 159_163 | 114_114 | 170_184 | 245_245 | 266_266 | 254_254 | 218_218 |
| 137_137 | NA | 132_141 | NA | NA | NA | NA | 266_266 | 254_254 | NA |
| 137_140 | NA | 132_141 | NA | NA | NA | NA | 266_266 | 254_254 | NA |
| 137_137 | NA | 132_132 | NA | NA | NA | NA | 266_266 | 254_254 | NA |
| 137_137 | NA | 132_132 | NA | NA | NA | NA | 266_266 | 254_254 | NA |
| 137_137 | 171_188 | 123_141 | 155_161 | 114_122 | 170_170 | 245_245 | 264_266 | 254_254 | 219_225 |
| 137_137 | 136_144 | 132_132 | 157_157 | 122_122 | 179_186 | 245_245 | 266_266 | 254_266 | 218_225 |
| 140_142 | NA | 132_135 | NA | NA | NA | NA | 266_266 | 254_254 | NA |
| 137_137 | 177_181 | 132_132 | 157_159 | 114_114 | 184_184 | 245_245 | 266_266 | 254_254 | 218_218 |
| 137_137 | 141_141 | 132_141 | 155_157 | 114_114 | 184_184 | 245_245 | 266_266 | 254_254 | 214_225 |
| 137_142 | 143_144 | 132_132 | 157_159 | 114_122 | 170_179 | 245_245 | 266_266 | 254_254 | 218_220 |
| 137_137 | 141_144 | 132_132 | 157_157 | 114_122 | 179_184 | 245_245 | 266_266 | 254_254 | 220_220 |
| 137_137 | 134_138 | 132_132 | 157_163 | 122_122 | 184_184 | 245_245 | 264_266 | 254_254 | 220_225 |
| 137_137 | 136_141 | 132_132 | 157_159 | 114_122 | 179_186 | 245_245 | 264_266 | 254_266 | 218_220 |
| 137_137 | 141_175 | 123_132 | 156_156 | 114_122 | 184_184 | 245_245 | 264_264 | 254_254 | 218_220 |
| 137_137 | NA | 132_132 | NA | NA | NA | NA | 266_266 | 254_254 | NA |
| 137_137 | NA | 132_132 | NA | NA | NA | NA | 264_266 | 254_254 | NA |
| 137_140 | NA | 132_132 | NA | NA | NA | NA | 264_266 | 254_254 | NA |

| | | | | | | | | | |
|---|---|---|---|---|---|---|---|---|---|
| 137_141 | NA | 132_132 | NA | NA | NA | NA | 266_266 | 254_254 | NA |
| 137_140 | NA | 132_141 | NA | NA | NA | NA | 264_266 | 254_254 | NA |
| 137_142 | 138_141 | 132_132 | 157_157 | 114_122 | 174_184 | 245_245 | 264_266 | 254_254 | 218_220 |
| 137_137 | NA | 132_141 | NA | NA | NA | NA | 264_266 | 254_254 | NA |
| 137_137 | 187_188 | 131_132 | 157_161 | 114_114 | 170_184 | 245_248 | 264_266 | 254_254 | 218_220 |
| 137_140 | 141_171 | 132_132 | 155_157 | 114_122 | 179_184 | 238_248 | 266_266 | 254_254 | 220_225 |
| 137_137 | NA | 132_132 | NA | NA | NA | NA | 266_266 | 254_254 | NA |
| 137_137 | NA | 132_132 | NA | NA | NA | NA | 264_266 | 254_254 | NA |
| 137_137 | NA | 132_141 | NA | NA | NA | NA | 264_266 | 254_254 | NA |
| 137_137 | NA | 132_132 | NA | NA | NA | NA | 266_266 | 254_270 | NA |
| 137_137 | NA | 132_132 | NA | NA | NA | NA | 266_266 | 254_270 | NA |
| 137_137 | NA | 132_135 | NA | NA | NA | NA | 264_266 | 254_254 | NA |
| 137_137 | NA | 132_132 | NA | NA | NA | NA | 264_266 | 254_270 | NA |
| 137_137 | NA | 132_132 | NA | NA | NA | NA | 264_266 | 254_254 | NA |
| 137_140 | NA | 132_132 | NA | NA | NA | NA | 264_266 | 254_254 | NA |
| 137_137 | NA | 123_132 | NA | NA | NA | NA | 264_272 | 254_254 | NA |
| 137_142 | 134_181 | 132_141 | 157_161 | 114_122 | 179_186 | 245_245 | 266_266 | 254_266 | 220_220 |
| 137_137 | NA | 141_141 | NA | NA | NA | NA | 264_266 | 254_254 | NA |
| 137_137 | NA | 132_132 | NA | NA | NA | NA | 264_266 | 254_254 | NA |
| 137_137 | NA | 132_141 | NA | NA | NA | NA | 266_266 | 254_254 | NA |
| 137_137 | NA | 123_132 | NA | NA | NA | NA | 266_266 | 254_254 | NA |
| 137_137 | NA | 132_141 | NA | NA | NA | NA | 264_266 | 254_254 | NA |
| 137_137 | NA | 132_135 | NA | NA | NA | NA | 266_266 | 254_254 | NA |
| 137_137 | NA | 123_132 | NA | NA | NA | NA | 266_266 | 254_266 | NA |
| 137_137 | NA | 132_132 | NA | NA | NA | NA | 266_266 | 254_254 | NA |
| 137_137 | NA | 132_132 | NA | NA | NA | NA | 264_266 | 254_254 | NA |
| 137_137 | NA | 132_132 | NA | NA | NA | NA | 264_272 | 254_254 | NA |
| 137_137 | NA | 132_132 | NA | NA | NA | NA | 264_264 | 254_254 | NA |
| 136_137 | NA | 132_132 | NA | NA | NA | NA | 266_266 | 254_270 | NA |
| 137_142 | NA | 135_141 | NA | NA | NA | NA | 266_266 | 254_270 | NA |
| 137_137 | NA | 132_132 | NA | NA | NA | NA | 266_266 | 254_254 | NA |
| 137_137 | NA | 132_132 | NA | NA | NA | NA | 266_266 | 254_254 | NA |
| 137_137 | NA | 132_132 | NA | NA | NA | NA | 266_266 | 254_254 | NA |
| 137_140 | NA | 132_132 | 157_157 | 114_114 | 179_179 | 245_245 | 266_266 | 254_254 | 209_209 |
| 133_137 | 170_171 | 132_132 | 157_157 | 114_114 | 184_184 | 245_245 | 264_266 | 254_254 | 220_220 |
| 137_137 | 136_141 | 132_132 | 155_157 | 114_114 | 184_184 | 245_245 | 266_266 | 254_254 | 220_220 |
| 133_137 | 141_171 | 132_141 | 157_163 | 114_114 | 170_184 | 245_245 | 266_266 | 254_270 | 218_220 |
| 137_137 | 150_171 | 132_132 | 155_157 | 114_122 | 174_179 | 245_245 | 266_266 | 254_270 | 210_225 |
| 137_140 | 141_141 | 132_159 | 157_157 | 114_114 | 179_179 | 245_245 | 266_272 | 254_254 | 218_218 |
| 137_137 | 197_199 | 132_132 | 155_155 | 114_122 | 170_184 | 245_245 | 264_264 | 254_254 | 209_220 |
| 137_140 | 170_171 | 132_159 | 157_159 | 114_114 | 184_184 | 245_245 | 264_266 | 254_254 | 218_220 |
| 137_140 | NA | 132_132 | NA | NA | NA | NA | 264_266 | 254_254 | NA |
| 137_137 | NA | 132_132 | NA | NA | NA | NA | 266_266 | 254_254 | NA |
| 137_137 | NA | 132_141 | NA | NA | NA | NA | 266_266 | 255_266 | NA |
| 137_137 | NA | 132_132 | NA | NA | NA | NA | 266_266 | 254_254 | NA |
| 137_137 | NA | 132_135 | NA | NA | NA | NA | 266_272 | 254_254 | NA |
| 133_137 | 177_183 | 132_141 | 159_163 | 114_114 | 179_184 | 245_245 | 264_266 | 254_254 | 209_218 |
| 137_137 | 143_143 | 132_141 | 157_157 | 114_122 | 184_186 | 245_245 | 266_266 | 254_254 | 220_220 |

| | | | | | | | | | |
|---|---|---|---|---|---|---|---|---|---|
| 137_137 | 144_187 | 132_141 | 157_161 | 114_122 | 184_184 | 238_245 | 266_266 | 254_254 | 220_220 |
| 137_137 | 143_185 | 132_135 | 161_163 | 114_114 | 170_179 | 245_245 | 266_266 | 254_254 | 220_220 |
| 137_142 | 134_181 | 123_135 | 155_159 | 114_122 | 170_179 | 245_245 | 264_264 | 254_254 | 218_225 |
| 137_142 | NA | 132_132 | NA | NA | NA | NA | 266_266 | 254_254 | NA |
| 136_137 | 141_171 | 132_132 | 155_157 | 114_114 | 170_179 | 245_245 | 266_266 | 254_254 | 220_225 |
| 136_137 | 141_141 | 132_132 | 157_161 | 114_122 | 179_184 | 245_245 | 264_266 | 270_270 | 209_220 |
| 136_137 | 144_170 | 132_132 | 157_161 | 114_114 | 179_184 | 245_245 | 264_266 | 254_254 | 220_225 |
| 137_140 | 144_170 | 132_132 | 155_157 | 122_122 | 170_184 | 245_245 | 266_266 | 254_270 | 220_220 |
| 137_137 | 141_141 | 132_141 | 157_161 | 114_122 | 184_184 | 245_245 | 266_266 | 254_270 | 220_220 |
| 137_137 | 179_181 | 132_132 | 157_157 | 114_114 | 179_184 | 245_245 | 266_266 | 254_254 | 218_220 |
| 137_137 | 170_175 | 132_141 | 157_157 | 114_114 | 179_184 | 245_245 | 264_266 | 254_254 | 218_220 |
| 137_137 | 141_183 | 141_141 | 155_157 | 114_114 | 170_184 | 245_245 | 266_266 | 254_270 | 218_225 |
| 137_137 | 141_141 | NA | 155_157 | 114_114 | 179_184 | 245_245 | NA | NA | 220_220 |
| 137_137 | 170_171 | 132_132 | 157_159 | 114_114 | 184_184 | 245_245 | 264_266 | 254_270 | 220_220 |
| 137_137 | NA | 132_132 | NA | NA | NA | NA | 264_266 | 254_254 | NA |
| 136_137 | NA | 132_132 | NA | NA | NA | NA | 266_266 | 254_254 | NA |
| 136_137 | NA | 132_132 | NA | NA | NA | NA | 264_266 | 254_254 | NA |
| 137_137 | NA | 132_132 | NA | NA | NA | NA | 266_266 | 254_254 | NA |
| 137_137 | 164_166 | 135_135 | 157_157 | 114_122 | 179_184 | 238_245 | 264_266 | 254_254 | 220_220 |
| 137_137 | 141_173 | 132_132 | 157_157 | 114_114 | 170_186 | 245_245 | 264_266 | 254_254 | 220_220 |
| 137_137 | 141_171 | 141_141 | 157_161 | 114_122 | 184_184 | 236_245 | 264_264 | 251_254 | 218_220 |
| 137_137 | 141_171 | 132_141 | 157_157 | 114_122 | 179_184 | 245_245 | 264_266 | 254_254 | 210_220 |
| 137_137 | NA | 132_132 | NA | NA | NA | NA | 266_266 | 254_254 | NA |
| 137_137 | NA | 132_141 | NA | NA | NA | NA | 266_266 | 254_254 | NA |
| 136_137 | 144_177 | 132_141 | 155_157 | 114_114 | 170_179 | 245_245 | 266_266 | 254_254 | 220_220 |
| 137_137 | NA | 132_138 | NA | NA | NA | NA | 266_266 | 254_254 | NA |
| 137_137 | 141_144 | 132_141 | 157_163 | 114_114 | 184_184 | 245_245 | 264_266 | 254_254 | 220_220 |
| 137_137 | 141_166 | 123_132 | 157_163 | 114_122 | 170_184 | NA | 264_266 | 254_254 | 218_220 |
| 137_140 | 141_141 | 132_132 | 155_157 | 122_122 | 184_184 | 245_245 | 266_266 | 254_254 | 218_218 |
| 137_137 | 170_173 | 132_135 | 157_163 | 114_122 | 170_184 | 245_245 | 264_266 | 254_254 | 217_220 |
| 137_137 | 170_171 | 132_135 | 157_157 | 114_114 | 184_184 | 245_245 | 264_266 | 254_254 | 220_220 |
| 137_140 | 170_177 | 132_135 | 159_163 | 114_114 | 170_184 | 238_245 | 264_266 | 254_254 | 220_220 |
| 133_137 | 173_177 | 132_135 | 157_163 | 122_122 | 184_184 | 245_245 | 266_266 | 254_254 | 220_225 |
| 137_137 | 141_141 | 132_132 | 157_157 | 114_114 | 184_186 | 245_245 | 266_266 | 254_254 | 220_225 |
| 137_137 | NA | 132_141 | NA | NA | NA | NA | 266_266 | 254_254 | NA |
| 137_140 | NA | 123_141 | NA | NA | NA | NA | 266_266 | 254_254 | NA |
| 137_137 | NA | 132_141 | NA | NA | NA | NA | 264_264 | 254_254 | NA |
| 137_140 | NA | 132_132 | NA | NA | NA | NA | 264_266 | 254_266 | NA |
| 137_137 | NA | 132_135 | NA | NA | NA | NA | 266_266 | 254_270 | NA |
| 137_137 | NA | 132_132 | NA | NA | NA | NA | 266_266 | 254_254 | NA |
| 136_137 | NA | 132_132 | NA | NA | NA | NA | 266_266 | 254_254 | NA |
| 140_140 | NA | 132_132 | NA | NA | NA | NA | 266_266 | 254_254 | NA |
| 136_137 | NA | 132_132 | NA | NA | NA | NA | 266_266 | 254_266 | NA |
| 137_137 | NA | 132_132 | NA | NA | NA | NA | 266_266 | 254_254 | NA |
| 137_137 | 144_170 | 132_135 | 157_163 | 114_114 | 179_184 | 245_245 | 266_266 | 254_254 | 220_220 |
| 136_137 | NA | 132_135 | NA | NA | NA | NA | 264_264 | 254_254 | NA |
| 137_140 | NA | 132_141 | NA | NA | NA | NA | 266_268 | 254_255 | NA |
| 137_137 | NA | 132_132 | NA | NA | NA | NA | 264_266 | 254_254 | NA |

| | | | | | | | | | |
|---|---|---|---|---|---|---|---|---|---|
| 137_137 | NA | 132_132 | NA | NA | NA | NA | 266_266 | 254_254 | NA |
| 137_137 | 134_134 | 132_132 | 157_157 | 114_114 | 170_184 | 245_245 | 266_266 | 254_254 | 214_225 |
| 137_137 | 141_170 | 132_132 | 157_157 | 114_114 | 170_184 | 245_245 | 266_266 | 254_254 | 220_220 |
| 137_137 | 141_141 | 132_132 | 155_157 | 122_122 | 179_184 | 245_245 | 266_266 | 254_266 | 218_220 |
| 137_137 | 134_144 | 132_132 | 157_157 | 114_114 | 170_184 | 245_245 | 266_266 | 254_254 | 220_220 |
| 137_137 | NA | 132_132 | NA | NA | NA | NA | 264_266 | 254_254 | NA |
| 137_137 | NA | 132_132 | NA | NA | NA | NA | 266_266 | 254_266 | NA |
| 137_137 | NA | 132_132 | NA | NA | NA | NA | 266_266 | 254_254 | NA |
| 137_137 | NA | 132_132 | NA | NA | NA | NA | 264_266 | 254_266 | NA |
| 137_137 | NA | 132_141 | NA | NA | NA | NA | 266_266 | 254_254 | NA |
| 137_137 | NA | 132_132 | NA | NA | NA | NA | 266_266 | 254_254 | NA |
| 137_137 | NA | 123_123 | NA | NA | NA | NA | 266_266 | 254_254 | NA |
| 137_137 | NA | 132_132 | NA | NA | NA | NA | 266_266 | 254_254 | NA |
| 137_137 | NA | 132_132 | NA | NA | NA | NA | 266_266 | 254_254 | NA |
| 137_137 | NA | 132_141 | NA | NA | NA | NA | 264_266 | 254_267 | NA |
| 137_137 | NA | 132_132 | NA | NA | NA | NA | 266_266 | 254_254 | NA |
| 137_140 | NA | 132_132 | NA | NA | NA | NA | 266_266 | 254_254 | NA |
| 137_137 | NA | 132_132 | NA | NA | NA | NA | 264_264 | 254_254 | NA |
| 137_140 | NA | 132_141 | NA | NA | NA | NA | 266_266 | 254_254 | NA |
| 137_137 | NA | 123_132 | NA | NA | NA | NA | 264_266 | 254_254 | NA |
| 133_137 | NA | 132_132 | NA | NA | NA | NA | 266_266 | 254_270 | NA |
| 137_140 | NA | 132_135 | NA | NA | NA | NA | 264_264 | 254_254 | NA |
| NA | NA | 132_132 | NA | NA | NA | NA | 266_266 | 254_254 | NA |
| 137_140 | NA | 141_141 | NA | NA | NA | NA | 264_266 | 261_267 | NA |
| 137_137 | NA | 132_132 | NA | NA | NA | NA | 264_266 | 254_254 | NA |
| 137_137 | NA | 132_132 | NA | NA | NA | NA | 264_266 | 254_270 | NA |
| 137_137 | NA | 132_132 | NA | NA | NA | NA | 266_266 | 254_254 | NA |
| 137_137 | NA | 132_132 | NA | NA | NA | NA | 264_266 | 254_254 | NA |
| 136_137 | NA | 132_132 | NA | NA | NA | NA | 264_264 | 254_254 | NA |
| 137_140 | NA | 132_132 | NA | NA | NA | NA | 266_266 | 254_266 | NA |
| 137_140 | NA | 132_132 | NA | NA | NA | NA | 266_266 | 254_254 | NA |
| 137_137 | NA | NA | NA | NA | NA | NA | 266_266 | 254_254 | NA |
| 137_137 | NA | 132_132 | NA | NA | NA | NA | 266_272 | 254_254 | NA |
| 137_140 | NA | 132_132 | NA | NA | NA | NA | 266_266 | 254_254 | NA |
| 137_137 | NA | 132_132 | NA | NA | NA | NA | 266_266 | 254_254 | NA |
| 137_137 | NA | 132_132 | NA | NA | NA | NA | 264_266 | 254_254 | NA |
| 135_137 | NA | 132_132 | NA | NA | NA | NA | 264_266 | 254_254 | NA |
| 137_137 | NA | 132_132 | NA | NA | NA | NA | 266_266 | 254_254 | NA |
| 137_137 | NA | 132_132 | NA | NA | NA | NA | 264_266 | 254_254 | NA |
| 137_137 | NA | 132_132 | NA | NA | NA | NA | 266_266 | 254_254 | NA |
| 137_137 | NA | 132_132 | NA | NA | NA | NA | 266_266 | 254_254 | NA |
| 137_137 | NA | 135_135 | NA | NA | NA | NA | 264_266 | 254_270 | NA |
| 137_137 | NA | 132_132 | NA | NA | NA | NA | 264_266 | 254_254 | NA |
| 136_137 | NA | 132_132 | NA | NA | NA | NA | 264_266 | 254_254 | NA |
| 137_140 | NA | 132_132 | NA | NA | NA | NA | 264_266 | 254_254 | NA |
| 136_137 | NA | 132_141 | NA | NA | NA | NA | 266_266 | 254_254 | NA |
| 137_137 | NA | 132_132 | NA | NA | NA | NA | 266_266 | 254_254 | NA |
| 137_137 | NA | 132_141 | NA | NA | NA | NA | 264_266 | 254_254 | NA |

| | | | | | | | | | |
|---|---|---|---|---|---|---|---|---|---|
| 137_137 | NA | 132_132 | NA | NA | NA | NA | 264_266 | 254_254 | NA |
| 137_137 | NA | 132_141 | NA | NA | NA | NA | 266_266 | 254_254 | NA |
| 137_137 | NA | 123_132 | NA | NA | NA | NA | 266_272 | 254_254 | NA |
| 137_137 | NA | 132_132 | NA | NA | NA | NA | 264_266 | 254_254 | NA |
| 137_137 | NA | NA | NA | NA | NA | NA | 264_266 | 254_254 | NA |
| 137_137 | NA | 132_132 | NA | NA | NA | NA | 264_266 | 254_254 | NA |
| 137_140 | NA | 132_132 | NA | NA | NA | NA | 264_266 | 254_254 | NA |
| 137_137 | NA | 132_135 | NA | NA | NA | NA | 264_264 | 254_254 | NA |
| 137_137 | NA | 132_132 | NA | NA | NA | NA | 264_266 | 254_254 | NA |
| 137_137 | NA | 132_141 | NA | NA | NA | NA | 264_266 | 254_254 | NA |
| 137_140 | NA | 123_159 | NA | NA | NA | NA | 266_266 | 254_270 | NA |
| 137_140 | NA | 132_141 | NA | NA | NA | NA | 264_266 | 254_254 | NA |
| 137_137 | NA | 132_132 | NA | NA | NA | NA | 266_266 | 254_254 | NA |
| 137_137 | NA | 132_132 | NA | NA | NA | NA | 266_266 | 254_254 | NA |
| 137_140 | 141_141 | 132_132 | 157_159 | 114_114 | 184_184 | 245_245 | 266_266 | 254_254 | 218_218 |
| 137_137 | NA | 132_141 | NA | NA | NA | NA | 266_266 | 254_254 | NA |
| 137_142 | 141_177 | 132_132 | 155_157 | 114_114 | 179_184 | 245_245 | 266_266 | 254_254 | 214_225 |
| 137_140 | 136_141 | 132_159 | 155_155 | 114_114 | 170_184 | 245_245 | 266_266 | 254_254 | 218_225 |
| 137_137 | 141_185 | 132_159 | 157_157 | 114_114 | 170_170 | 245_245 | 266_266 | 254_254 | 218_218 |
| 137_137 | 141_177 | 132_132 | 157_159 | 114_114 | 179_179 | 245_245 | 264_266 | 254_254 | 218_225 |
| 137_142 | NA | 132_132 | NA | NA | NA | NA | 266_266 | 254_254 | NA |
| 137_137 | NA | 132_141 | NA | NA | NA | NA | 266_266 | 254_254 | NA |
| 137_137 | NA | 123_132 | NA | NA | NA | NA | 264_266 | 254_254 | NA |
| 137_137 | 141_171 | 123_132 | 155_163 | 114_114 | 174_184 | 245_245 | 266_266 | 262_267 | 218_218 |
| 137_137 | 141_144 | 132_159 | 157_157 | 114_114 | 174_184 | 245_245 | 264_266 | 254_254 | 217_218 |
| 137_137 | 141_141 | 132_132 | 155_159 | 114_114 | 174_174 | 245_245 | 264_266 | 254_254 | 214_220 |
| 137_140 | NA | 132_132 | NA | NA | NA | NA | 264_266 | 254_254 | NA |
| 137_137 | 141_141 | 132_132 | 157_159 | 114_114 | 170_170 | 245_245 | 266_266 | 254_270 | 218_218 |
| 137_137 | 121_141 | 132_159 | 157_157 | 114_114 | 170_179 | 245_245 | 266_266 | 254_266 | 217_218 |
| 140_147 | 144_173 | 132_132 | 157_159 | 114_114 | 179_184 | 245_245 | 266_266 | 254_266 | 214_217 |
| 137_137 | 141_141 | 132_132 | 155_157 | 114_114 | 170_184 | 245_245 | 266_266 | 254_266 | 214_220 |
| 137_147 | NA | 132_132 | NA | NA | NA | NA | 264_272 | 254_267 | NA |
| 137_137 | 141_183 | 132_132 | 157_157 | 114_114 | 170_174 | 245_245 | 266_266 | 254_254 | 218_222 |
| 137_140 | 141_197 | 135_159 | 155_157 | 114_114 | 170_184 | 245_245 | 264_266 | 254_254 | 217_220 |
| 136_137 | 164_170 | 123_132 | 157_157 | 114_114 | 174_174 | 245_245 | 266_266 | 254_267 | 217_218 |
| 137_137 | 121_195 | 132_132 | 157_157 | 114_114 | 170_184 | 245_245 | 266_266 | 254_254 | 217_218 |
| 137_137 | 121_141 | 123_132 | 157_157 | 114_114 | 170_184 | 245_245 | 264_266 | 266_266 | 214_218 |
| 137_137 | NA | 132_132 | NA | NA | NA | NA | 264_266 | 254_267 | NA |
| 137_140 | 141_144 | 141_141 | 157_159 | 114_114 | 170_174 | 245_245 | 266_266 | 254_254 | 225_225 |
| 137_137 | 141_175 | 132_132 | 157_157 | 114_114 | NA | 245_245 | 264_266 | 254_254 | 217_218 |
| 137_137 | 136_176 | 129_129 | 157_163 | 114_114 | 174_174 | 232_245 | 264_266 | 254_254 | 214_218 |
| 137_137 | NA | 132_132 | NA | NA | NA | NA | 266_266 | 254_254 | NA |
| 137_137 | NA | 132_141 | NA | NA | NA | NA | 264_266 | 254_254 | NA |
| 137_137 | 141_141 | 132_159 | 157_159 | 114_114 | 170_184 | 245_245 | 264_266 | 254_254 | 218_218 |
| 137_140 | NA | 123_132 | NA | NA | NA | NA | 264_266 | 254_254 | NA |
| 137_137 | NA | 132_132 | NA | NA | NA | NA | 266_266 | 254_254 | NA |
| 140_140 | NA | 132_159 | NA | NA | NA | NA | 266_266 | 254_254 | NA |
| 137_137 | NA | 123_132 | NA | NA | NA | NA | 264_266 | 254_254 | NA |

| | | | | | | | | | |
|---|---|---|---|---|---|---|---|---|---|
| 137_140 | NA | 132_141 | NA | NA | NA | NA | 266_266 | 254_254 | NA |
| NA | NA | 132_141 | NA | NA | NA | NA | 266_266 | 254_254 | NA |
| 137_137 | NA | 132_132 | NA | NA | NA | NA | 266_266 | 254_254 | NA |
| 137_137 | NA | 132_132 | NA | NA | NA | NA | 264_272 | 254_254 | NA |
| 140_147 | NA | 132_159 | NA | NA | NA | NA | 264_264 | 254_254 | NA |
| 137_137 | NA | 132_132 | NA | NA | NA | NA | 264_266 | 254_254 | NA |
| 137_140 | NA | 132_132 | NA | NA | NA | NA | 266_266 | 254_266 | NA |
| 137_137 | NA | 132_132 | NA | NA | NA | NA | 266_266 | 254_254 | NA |
| 137_137 | NA | 132_132 | NA | NA | NA | NA | 264_266 | 254_254 | NA |
| 137_137 | NA | 132_132 | NA | NA | NA | NA | 266_266 | 254_254 | NA |
| 137_137 | NA | 132_132 | NA | NA | NA | NA | 266_266 | 254_254 | NA |
| 137_137 | NA | 132_132 | NA | NA | NA | NA | 266_266 | 254_254 | NA |
| 137_137 | NA | 132_159 | NA | NA | NA | NA | 264_272 | 266_266 | NA |
| 137_137 | NA | 132_132 | NA | NA | NA | NA | 266_266 | 254_254 | NA |
| 137_137 | NA | 132_159 | NA | NA | NA | NA | 264_266 | 254_254 | NA |
| 137_140 | NA | 132_132 | NA | NA | NA | NA | 266_266 | 254_254 | NA |
| 137_140 | NA | 132_159 | NA | NA | NA | NA | 264_266 | 254_254 | NA |
| 137_137 | NA | 132_132 | NA | NA | NA | NA | 266_266 | 254_254 | NA |
| 137_137 | NA | 132_132 | NA | NA | NA | NA | 264_266 | 254_254 | NA |
| 137_142 | NA | 132_132 | NA | NA | NA | NA | 266_266 | 254_254 | NA |
| 137_137 | NA | 132_141 | NA | NA | NA | NA | 266_266 | 254_254 | NA |
| 137_140 | NA | 132_141 | NA | NA | NA | NA | 264_266 | 254_254 | NA |
| 137_137 | NA | 132_132 | NA | NA | NA | NA | 264_266 | 254_266 | NA |
| 137_137 | NA | 132_132 | NA | NA | NA | NA | 264_266 | 254_254 | NA |
| 137_140 | NA | 132_132 | NA | NA | NA | NA | 264_272 | 254_254 | NA |
| 137_137 | NA | 132_132 | NA | NA | NA | NA | 264_266 | 254_254 | NA |
| 137_140 | NA | 132_132 | NA | NA | NA | NA | 266_266 | 254_254 | NA |
| 137_147 | NA | 141_141 | NA | NA | NA | NA | 266_266 | 254_254 | NA |
| 137_137 | NA | 132_132 | NA | NA | NA | NA | 266_266 | 254_270 | NA |
| 137_140 | NA | 132_132 | NA | NA | NA | NA | 266_266 | 254_254 | NA |
| 137_137 | NA | 132_132 | NA | NA | NA | NA | 266_266 | 254_254 | NA |
| 137_137 | NA | 132_163 | NA | NA | NA | NA | 264_266 | 254_254 | NA |
| 137_137 | NA | 132_132 | NA | NA | NA | NA | 266_266 | 254_254 | NA |
| 137_142 | NA | 132_132 | NA | NA | NA | NA | 266_266 | 254_254 | NA |
| 137_137 | NA | 132_132 | NA | NA | NA | NA | 264_266 | 254_254 | NA |
| 137_140 | NA | 132_159 | NA | NA | NA | NA | 266_266 | 254_254 | NA |
| 137_140 | NA | 132_159 | NA | NA | NA | NA | 264_266 | 254_254 | NA |
| 137_137 | NA | 132_132 | NA | NA | NA | NA | 264_266 | 254_254 | NA |
| 137_137 | NA | 132_132 | NA | NA | NA | NA | 264_266 | 254_254 | NA |
| 137_137 | NA | 132_132 | NA | NA | NA | NA | 264_264 | 254_254 | NA |
| 137_137 | NA | 132_132 | NA | NA | NA | NA | 266_266 | 254_266 | NA |
| 137_137 | NA | 132_135 | NA | NA | NA | NA | 264_266 | 254_254 | NA |
| 137_140 | NA | 132_141 | NA | NA | NA | NA | 264_266 | 254_254 | NA |
| 137_137 | NA | 132_132 | NA | NA | NA | NA | 266_266 | 254_254 | NA |
| 137_137 | NA | 132_132 | NA | NA | NA | NA | 266_266 | 254_254 | NA |
| 137_137 | NA | 123_135 | NA | NA | NA | NA | 264_266 | 254_254 | NA |
| 137_137 | NA | 132_163 | NA | NA | NA | NA | 264_264 | 254_266 | NA |
| 137_137 | NA | 132_132 | NA | NA | NA | NA | 266_266 | 254_270 | NA |

| | | | | | | | | | |
|---|---|---|---|---|---|---|---|---|---|
| 137_137 | NA | 132_132 | NA | NA | NA | NA | 266_266 | 254_254 | NA |
| 137_137 | NA | 132_132 | NA | NA | NA | NA | 266_266 | 254_254 | NA |
| 137_137 | NA | 132_141 | NA | NA | NA | NA | 264_266 | 254_254 | NA |
| 137_137 | NA | 132_132 | NA | NA | NA | NA | 264_266 | 254_254 | NA |
| 137_137 | NA | 132_132 | NA | NA | NA | NA | 264_266 | 254_254 | NA |
| 137_137 | 141_173 | 123_141 | 155_157 | 114_122 | 170_179 | 241_245 | 264_266 | 254_254 | 217_217 |
| 136_137 | 141_141 | 132_132 | 155_163 | 114_114 | 184_192 | 245_245 | 266_266 | 254_270 | 218_218 |
| 137_137 | NA | 132_132 | NA | NA | NA | NA | 264_266 | 254_254 | NA |
| 137_141 | NA | 132_132 | NA | NA | NA | NA | 266_266 | 254_254 | NA |
| 137_137 | NA | 132_132 | NA | NA | NA | NA | 264_266 | 254_254 | NA |
| 137_140 | NA | 123_132 | NA | NA | NA | NA | 264_266 | 254_266 | NA |
| 137_137 | NA | 132_159 | NA | NA | NA | NA | 266_266 | 254_254 | NA |
| 137_137 | NA | 123_131 | NA | NA | NA | NA | 264_266 | 254_254 | NA |
| 137_140 | NA | 132_132 | NA | NA | NA | NA | 266_266 | 254_254 | NA |
| 137_142 | 141_183 | 132_153 | 163_165 | 114_114 | 170_179 | 238_245 | 264_266 | 254_254 | 217_225 |
| 137_137 | 141_188 | 132_132 | 157_163 | 114_114 | 184_184 | 245_245 | 266_266 | 254_270 | 220_225 |
| 137_137 | 136_141 | 141_141 | 157_165 | 114_114 | 170_184 | 238_245 | 266_266 | 254_254 | 210_220 |
| 137_137 | 138_138 | 132_132 | 157_159 | 114_114 | 179_184 | 245_245 | 264_266 | 254_254 | 220_225 |
| 137_137 | 141_141 | 132_132 | 157_159 | 114_114 | 184_184 | 245_245 | 264_266 | 254_254 | 217_220 |
| 137_140 | NA | 132_132 | NA | NA | NA | NA | 264_266 | 254_254 | NA |
| 137_137 | NA | 132_159 | NA | NA | NA | NA | 266_266 | 254_270 | NA |
| 137_137 | 171_177 | NA | 157_157 | 114_114 | 179_184 | 245_245 | 264_266 | 254_254 | 220_220 |
| 137_137 | 136_136 | 132_132 | 157_157 | 114_114 | 170_170 | 245_245 | 264_266 | 254_254 | 218_220 |
| 137_137 | 134_144 | 132_132 | 157_161 | 114_114 | 179_184 | 245_245 | 264_266 | 254_254 | 214_218 |
| 137_140 | NA | 132_132 | NA | NA | NA | NA | 264_264 | 254_254 | NA |
| 137_140 | 134_173 | 132_132 | 151_157 | 114_114 | 179_179 | 245_245 | 266_272 | 254_254 | 217_220 |
| NA | 141_170 | NA | 157_157 | 114_114 | 170_184 | 245_245 | 266_266 | 254_266 | 220_220 |
| 137_140 | 141_183 | 132_141 | 157_161 | 114_114 | 170_170 | 245_245 | 264_266 | 254_254 | 217_220 |
| 137_140 | NA | 132_141 | NA | NA | NA | NA | 266_266 | 254_254 | NA |
| 137_137 | 173_188 | 132_132 | 156_163 | 114_122 | 174_184 | 245_245 | 264_266 | 254_254 | 218_218 |
| 137_137 | 136_141 | NA | 155_157 | 114_114 | 191_192 | 245_245 | 264_266 | 254_254 | 218_220 |
| 137_137 | NA | 132_132 | NA | NA | NA | NA | 266_266 | 254_254 | NA |
| 137_137 | NA | 132_132 | NA | NA | NA | NA | 266_266 | 254_254 | NA |
| 137_137 | NA | 132_132 | NA | NA | NA | NA | 264_264 | 254_254 | NA |
| 137_140 | NA | 132_132 | NA | NA | NA | NA | 266_266 | 254_254 | NA |
| 137_137 | NA | 132_141 | NA | NA | NA | NA | 266_266 | 254_254 | NA |
| 137_137 | 134_188 | 132_132 | 155_157 | 114_122 | 179_184 | 245_245 | 264_266 | 254_254 | 218_220 |
| 137_137 | 141_144 | 132_132 | 155_157 | 114_114 | 184_186 | 245_245 | 264_264 | 254_254 | 214_225 |
| 137_137 | 141_141 | 132_135 | 157_157 | 114_114 | 174_184 | 245_245 | 264_266 | 254_254 | 218_225 |
| 137_137 | 144_173 | 132_132 | 163_163 | 114_114 | 184_184 | 245_245 | 264_266 | 254_254 | 214_220 |
| 137_140 | 136_144 | 132_132 | 155_157 | 114_114 | 184_184 | 245_245 | 266_272 | 254_254 | 214_220 |
| NA | 144_185 | 141_141 | 155_155 | 114_114 | 184_184 | 245_245 | 264_266 | 254_254 | 218_218 |
| 137_137 | NA | 116_132 | NA | NA | NA | NA | 266_266 | 254_254 | NA |
| 137_142 | NA | 132_132 | NA | NA | NA | NA | 266_266 | 254_254 | NA |
| 137_137 | NA | 132_159 | NA | NA | NA | NA | 266_266 | 254_254 | NA |
| 137_140 | NA | 132_132 | NA | NA | NA | NA | 266_266 | 254_270 | NA |
| 137_137 | 141_170 | 132_132 | 157_161 | 114_114 | 179_184 | 245_245 | 266_272 | 254_266 | 220_222 |
| 137_140 | 170_181 | 132_132 | 157_163 | 114_114 | 174_184 | 245_245 | 266_266 | 254_254 | 220_225 |

| | | | | | | | | | |
|---|---|---|---|---|---|---|---|---|---|
| 137_137 | 134_144 | 132_132 | 155_157 | 114_114 | 184_184 | 245_245 | 266_266 | 266_270 | 218_220 |
| 137_142 | 134_141 | 132_132 | 157_157 | 114_114 | 170_184 | 245_245 | 266_266 | 254_254 | 218_225 |
| 137_137 | NA | 132_141 | NA | NA | NA | NA | 264_266 | 254_254 | NA |
| 137_140 | NA | 132_132 | NA | NA | NA | NA | 266_266 | 254_254 | NA |
| 137_137 | NA | 132_132 | NA | NA | NA | NA | 264_266 | 254_254 | NA |
| 137_140 | NA | 132_141 | NA | NA | NA | NA | 266_266 | 254_254 | NA |
| 137_137 | NA | 132_132 | NA | NA | NA | NA | 264_266 | 254_254 | NA |
| 137_140 | NA | 132_141 | NA | NA | NA | NA | 266_266 | 254_254 | NA |
| 137_137 | NA | 132_132 | NA | NA | NA | NA | 266_266 | 254_254 | NA |
| 137_140 | NA | 132_132 | NA | NA | NA | NA | 266_266 | 254_254 | NA |
| 137_137 | NA | 132_132 | NA | NA | NA | NA | 266_266 | 254_254 | NA |
| 137_140 | NA | 132_132 | NA | NA | NA | NA | 264_266 | 254_254 | NA |
| 137_137 | NA | 132_132 | NA | NA | NA | NA | 266_266 | 254_254 | NA |
| 137_137 | NA | 132_132 | NA | NA | NA | NA | 264_266 | 254_254 | NA |
| 137_137 | NA | 132_132 | NA | NA | NA | NA | 266_266 | 254_254 | NA |
| 137_140 | NA | 132_132 | NA | NA | NA | NA | 264_266 | 254_254 | NA |
| 137_137 | NA | 132_132 | NA | NA | NA | NA | 264_266 | 254_270 | NA |
| 137_137 | NA | 132_132 | NA | NA | NA | NA | 264_264 | 254_254 | NA |
| 137_140 | NA | NA | NA | NA | NA | NA | 266_266 | 254_254 | NA |
| 137_137 | NA | 132_132 | NA | NA | NA | NA | 264_266 | 254_254 | NA |
| 137_137 | NA | 132_132 | NA | NA | NA | NA | 266_266 | 254_254 | NA |
| 137_137 | NA | 132_141 | NA | NA | NA | NA | 264_264 | 254_254 | NA |
| 137_140 | NA | 132_132 | NA | NA | NA | NA | 264_266 | 254_270 | NA |
| 137_137 | NA | 132_132 | NA | NA | NA | NA | 266_266 | 254_254 | NA |
| 137_140 | 141_168 | NA | 161_163 | 114_114 | 184_184 | 245_245 | 266_266 | 254_254 | 218_220 |
| 137_137 | 138_141 | 132_132 | 163_165 | 114_122 | 170_184 | 245_245 | 266_266 | 254_254 | 220_225 |
| 137_142 | NA | NA | NA | NA | NA | NA | 266_266 | 254_254 | NA |
| 137_137 | NA | 132_132 | NA | NA | NA | NA | 266_266 | 254_254 | NA |
| 137_142 | NA | 132_132 | NA | NA | NA | NA | 266_266 | 254_254 | NA |
| 137_137 | 134_141 | 132_132 | 157_157 | 114_122 | 170_170 | 245_245 | 266_266 | 254_254 | 219_225 |
| 137_140 | 141_199 | 132_132 | 155_157 | 114_114 | 184_184 | 245_245 | 266_266 | 254_254 | 220_225 |
| 137_137 | NA | 132_132 | NA | NA | NA | NA | 266_266 | 254_254 | NA |
| 137_140 | NA | 132_132 | NA | NA | NA | NA | 266_266 | 254_254 | NA |
| 137_137 | 173_179 | 116_132 | 157_159 | 114_114 | 179_184 | 245_245 | 266_266 | 254_254 | 217_220 |
| 137_137 | 170_170 | 132_142 | 157_163 | 114_114 | 170_184 | 245_245 | 266_272 | 254_254 | 210_218 |
| 137_137 | 141_170 | 132_146 | 157_157 | 114_122 | 170_184 | 245_245 | 266_272 | 254_254 | 210_218 |
| 137_137 | 173_187 | 132_141 | 157_157 | 114_114 | 170_179 | 245_245 | 266_266 | 254_254 | 218_220 |
| 137_137 | 134_141 | 132_132 | NA | 114_114 | NA | 245_245 | 264_266 | 254_254 | 220_225 |
| 137_137 | NA | 132_132 | NA | NA | NA | NA | 266_266 | 254_254 | NA |
| 137_137 | 134_173 | 132_132 | 155_157 | 114_114 | 179_179 | 245_245 | 264_266 | 254_254 | 220_225 |
| 137_137 | 162_173 | 132_132 | 155_157 | 114_114 | 170_170 | 245_245 | 264_266 | 254_254 | 220_220 |
| 137_137 | 170_170 | 123_132 | 157_159 | 114_114 | 179_184 | 245_245 | 264_266 | 254_254 | 220_227 |
| 137_137 | 170_183 | 116_116 | 155_157 | 114_114 | 184_184 | 245_245 | 266_266 | 254_254 | 217_220 |
| 137_137 | 144_171 | 132_132 | 155_159 | 114_114 | 179_184 | 245_245 | 266_266 | 254_254 | 220_225 |
| 137_137 | 141_179 | 132_132 | 157_157 | 114_114 | 179_184 | 245_245 | 266_266 | 254_254 | 220_220 |
| 137_137 | NA | 132_132 | NA | NA | NA | NA | 266_266 | 254_254 | NA |
| 137_137 | 173_173 | 132_132 | 157_159 | 114_114 | 170_170 | 245_245 | 264_266 | 254_254 | 220_225 |
| 137_140 | 163_170 | 132_132 | 155_157 | 114_114 | 184_184 | 245_245 | 266_266 | 254_254 | 220_225 |

| | | | | | | | | | |
|---|---|---|---|---|---|---|---|---|---|
| 137_140 | 162_173 | 132_132 | 155_157 | 114_114 | 170_179 | 245_245 | 266_266 | 254_254 | 220_227 |
| 137_137 | 171_173 | 132_132 | 155_157 | 114_114 | 170_179 | 245_245 | 264_266 | 254_254 | 220_225 |
| 137_137 | NA | 132_132 | NA | NA | NA | NA | 264_266 | 254_266 | NA |
| 137_140 | NA | 132_132 | NA | NA | NA | NA | 264_266 | 254_254 | NA |
| 137_137 | 134_166 | 132_132 | 155_157 | 114_114 | 184_184 | 245_245 | 266_266 | 254_254 | 220_220 |
| 137_140 | NA | 132_132 | NA | NA | NA | NA | 266_266 | 254_254 | NA |
| 137_137 | NA | 132_132 | 155_155 | 114_114 | 179_179 | 245_245 | 264_266 | 254_254 | NA |
| 137_137 | 175_183 | 132_132 | 157_158 | 114_120 | 184_184 | 245_245 | 266_266 | 254_254 | 220_220 |
| 137_137 | 170_175 | 132_132 | 155_157 | 114_120 | 184_184 | 245_245 | 264_266 | 254_254 | 220_220 |
| 137_137 | 141_183 | 132_132 | 157_157 | 114_120 | 170_184 | 245_245 | 266_266 | 254_254 | 220_220 |
| 137_137 | 175_179 | 132_132 | 157_157 | 114_114 | 179_184 | 245_245 | 266_266 | 254_254 | 220_220 |
| 137_137 | 175_183 | 132_132 | 157_158 | 114_114 | 179_184 | 245_245 | 266_266 | 254_254 | 220_220 |
| 137_137 | 175_183 | 132_132 | 158_159 | 114_114 | 179_184 | 245_245 | 266_266 | 254_254 | 220_220 |
| 137_137 | NA | 123_123 | NA | NA | NA | NA | 264_266 | 254_254 | NA |
| 137_137 | 175_175 | 132_132 | 158_159 | 114_120 | 184_184 | 245_245 | 266_266 | 254_254 | 220_220 |
| 137_137 | 175_175 | 132_132 | 157_157 | 114_114 | 179_184 | 245_245 | 266_272 | 254_254 | 220_220 |
| 137_137 | 141_173 | 132_132 | 155_157 | 114_114 | 179_184 | 245_245 | 266_266 | 254_254 | 220_220 |
| 137_137 | 134_183 | 132_132 | NA | 114_120 | NA | NA | 264_266 | 254_254 | 220_220 |
| 137_137 | 175_175 | 132_132 | 158_159 | 114_120 | 184_184 | 245_245 | 266_266 | 254_254 | 220_220 |
| 137_137 | 175_183 | 132_133 | 157_159 | 114_120 | 179_179 | 245_245 | 266_266 | 254_254 | 220_220 |
| 137_137 | 134_170 | 132_132 | 155_157 | 114_114 | 179_184 | 245_245 | 264_264 | 254_254 | 225_225 |
| 137_137 | 134_173 | 132_132 | NA | 114_114 | NA | NA | 266_266 | 254_254 | 220_220 |
| 137_140 | NA | 132_132 | NA | NA | NA | NA | 264_266 | 254_254 | NA |
| 137_137 | NA | 132_132 | 155_157 | 114_114 | 179_184 | 245_245 | 266_266 | 254_254 | 217_225 |
| 137_137 | 134_170 | NA | 157_159 | 114_114 | 170_184 | 245_245 | 266_266 | 254_254 | 225_227 |
| 137_137 | NA | 132_132 | NA | NA | NA | NA | 266_266 | 254_254 | NA |
| 137_140 | 134_191 | 132_132 | 157_163 | 114_122 | 179_184 | 245_245 | 264_266 | 254_254 | 220_220 |
| 137_137 | NA | 135_141 | NA | NA | NA | NA | 264_266 | 254_254 | NA |
| NA | 166_170 | 132_132 | 158_163 | 114_114 | 170_170 | 245_245 | 264_264 | 254_270 | 218_218 |
| NA | 134_136 | 132_132 | 157_159 | 114_114 | 179_184 | 245_245 | 266_266 | 254_254 | 218_220 |
| 137_137 | 175_177 | 132_132 | 157_159 | 114_114 | 170_179 | 245_245 | 264_266 | 254_254 | 219_220 |
| 137_137 | 134_134 | 132_132 | 157_157 | 114_114 | 170_179 | 245_245 | 266_266 | 254_254 | 217_220 |
| 137_137 | 171_173 | 132_132 | 155_157 | 114_114 | 170_179 | 245_245 | 266_266 | 254_254 | 209_225 |
| 137_137 | 162_168 | 132_132 | 155_157 | 114_114 | 184_184 | 245_245 | 266_266 | 254_254 | 220_225 |
| 137_137 | 141_141 | 132_132 | 155_157 | 114_114 | 179_184 | 245_245 | 266_266 | 254_254 | 220_227 |
| 137_137 | 168_170 | 132_132 | 157_159 | 114_114 | 170_179 | 245_245 | 266_266 | 254_254 | 220_225 |
| 137_137 | 134_179 | 132_132 | 157_157 | 114_114 | 179_184 | 245_245 | 264_266 | 254_254 | 220_220 |
| 137_137 | 146_175 | 132_132 | 155_157 | 114_122 | 179_184 | 245_245 | 266_266 | 254_254 | 220_225 |
| 137_137 | 171_171 | 132_132 | 155_159 | 114_122 | 179_184 | 245_245 | 264_266 | 254_254 | 220_220 |
| 137_137 | 173_179 | 132_132 | 159_161 | 114_114 | 179_184 | 245_245 | 266_266 | 254_254 | 217_220 |
| 137_137 | NA | 132_132 | 157_159 | 114_114 | 179_179 | 245_245 | 266_266 | 254_254 | 220_220 |
| 137_137 | 141_141 | 132_132 | 155_155 | 114_114 | 184_184 | 245_245 | 266_266 | 254_254 | 220_220 |
| 137_137 | 141_141 | 132_132 | 157_157 | 114_114 | 179_184 | 245_245 | 266_266 | 254_254 | 220_220 |
| 137_137 | 144_200 | 132_132 | 157_157 | 114_114 | 179_179 | 245_245 | 266_266 | 254_254 | 220_220 |
| 137_137 | 141_141 | 132_132 | 157_157 | 114_114 | 179_184 | 245_245 | 264_266 | 254_254 | 220_220 |
| 136_137 | 141_141 | 132_132 | 157_157 | 114_114 | 184_184 | 245_245 | 266_266 | 254_254 | 220_220 |
| 137_137 | 136_141 | 132_132 | 159_159 | 114_114 | 184_184 | 245_245 | 266_266 | 254_254 | 225_225 |
| 136_137 | 136_200 | 132_132 | 157_157 | 114_114 | 179_184 | 245_245 | 266_266 | 254_270 | 225_225 |

| | | | | | | | | | |
|---|---|---|---|---|---|---|---|---|---|
| 137_137 | 141_166 | 132_132 | 155_157 | 114_114 | 184_186 | 245_245 | 266_266 | 254_254 | 220_227 |
| 137_137 | 168_179 | 132_132 | 155_157 | 114_114 | 184_184 | 245_245 | 266_266 | 254_261 | 220_220 |
| 137_137 | 141_171 | 132_132 | 157_159 | 114_114 | 179_179 | 245_245 | 266_266 | 254_254 | 220_225 |
| 137_137 | 141_177 | 132_132 | 155_155 | 114_114 | 170_179 | 245_245 | 266_266 | 254_254 | 218_220 |
| 137_137 | 138_193 | 132_132 | 159_159 | 114_122 | 170_184 | 245_245 | 264_266 | 254_254 | 218_218 |
| 137_137 | 141_179 | 132_132 | 157_157 | 114_114 | 179_179 | 245_245 | 264_266 | 254_254 | 220_220 |
| 137_137 | 173_177 | 132_132 | 155_157 | 114_114 | 179_184 | 245_245 | 266_266 | 254_254 | 220_220 |
| NA | 141_170 | 132_132 | 155_157 | 114_114 | 179_179 | 245_245 | 264_266 | 254_254 | 210_220 |
| 137_137 | 134_179 | NA | 157_167 | 114_114 | 170_184 | 244_245 | 264_264 | 254_254 | 218_220 |
| 137_137 | 141_173 | 132_132 | 157_163 | 114_114 | 170_170 | 245_245 | 264_266 | 254_254 | 218_220 |
| 137_137 | NA | 132_132 | NA | NA | NA | NA | 266_266 | 254_254 | NA |
| 137_137 | 170_170 | 132_132 | 157_157 | 114_114 | 179_179 | 245_245 | 264_266 | 254_254 | 220_220 |
| 137_140 | 144_168 | 116_132 | 157_157 | 114_114 | 179_184 | 245_245 | 264_266 | 254_254 | 220_225 |
| 137_137 | 141_177 | 132_132 | 157_157 | 114_114 | 170_184 | 245_245 | 266_266 | 254_254 | 217_225 |
| 137_137 | 141_170 | 132_132 | 155_155 | 114_114 | 179_179 | 245_245 | 264_266 | 254_254 | 220_220 |
| 137_137 | 134_134 | 132_132 | 155_157 | 114_114 | 179_184 | 245_245 | 266_266 | 254_254 | 220_220 |
| 137_137 | 141_173 | 132_132 | 155_157 | 114_114 | 179_184 | 245_245 | 266_266 | 254_254 | 220_220 |
| 137_137 | 144_144 | 132_132 | 157_157 | 114_114 | 179_179 | 245_245 | 266_266 | 254_254 | 220_220 |
| 137_137 | 141_171 | 132_132 | 155_157 | 114_114 | 184_184 | 245_245 | 266_266 | 254_254 | 218_225 |
| 137_137 | 171_173 | 132_132 | 155_155 | 114_114 | 184_184 | 245_245 | 266_266 | 254_254 | 218_225 |
| 137_137 | 136_141 | 132_132 | 157_159 | 114_114 | 184_184 | 245_245 | 266_266 | 254_254 | 217_220 |
| 137_137 | 166_173 | 132_132 | 155_157 | 114_114 | 184_184 | 245_245 | 264_266 | 254_254 | 218_225 |
| 137_137 | 136_181 | 132_132 | 155_157 | 114_122 | 179_184 | 245_245 | 266_266 | 254_266 | 218_218 |
| 137_137 | 141_170 | 132_132 | 155_155 | 114_114 | 179_184 | 245_245 | 266_266 | 254_254 | 218_220 |
| 137_137 | 141_141 | 132_159 | 157_157 | 114_114 | 170_179 | 245_245 | 264_266 | 254_254 | 220_220 |
| 137_137 | 141_141 | 132_159 | 155_157 | 114_114 | 184_184 | 245_245 | 266_266 | 254_254 | 220_220 |
| 137_137 | NA | 132_132 | NA | NA | NA | NA | 264_266 | 254_254 | NA |
| 137_137 | 141_203 | 132_132 | 157_159 | 114_114 | 179_184 | 245_245 | 266_266 | 254_254 | 219_225 |
| 137_137 | 136_185 | 132_132 | 155_157 | 114_114 | 179_184 | 245_245 | 266_266 | 254_254 | 219_225 |
| 137_137 | 141_175 | 116_132 | 155_157 | 114_114 | 184_184 | 245_245 | 266_266 | 254_254 | 220_220 |
| 137_137 | 164_173 | 132_132 | 155_157 | 114_122 | 179_184 | 245_245 | 266_266 | 254_254 | 220_225 |
| 137_137 | 134_141 | 132_132 | 157_161 | 114_114 | 170_184 | 245_245 | 266_266 | 254_254 | 220_220 |
| 137_137 | 134_144 | 132_132 | 157_157 | 114_114 | 184_184 | 238_245 | 266_266 | 254_254 | 218_218 |
| 137_137 | 141_177 | 132_132 | 157_159 | 114_114 | 179_179 | 245_245 | 266_266 | 254_254 | 220_220 |
| NA | 134_144 | 132_141 | 157_159 | 114_122 | 170_179 | 245_245 | 264_266 | 254_254 | 210_225 |
| NA | 141_187 | 132_132 | 157_157 | 114_114 | 170_170 | 245_245 | 264_266 | 254_254 | 225_227 |
| 137_137 | NA | 132_132 | NA | NA | NA | NA | 266_266 | 254_266 | NA |
| 137_137 | NA | 132_132 | NA | NA | NA | NA | 266_266 | 254_254 | NA |
| 137_137 | 134_179 | 132_132 | 155_157 | 114_114 | 184_184 | 245_245 | 266_266 | 254_254 | 225_225 |
| 137_137 | 141_191 | NA | 157_161 | 114_114 | 184_184 | 245_245 | 266_272 | 254_270 | 220_220 |
| 137_140 | NA | 132_132 | NA | NA | NA | NA | 266_266 | 254_254 | NA |
| 137_137 | 183_185 | 123_132 | 155_155 | 114_114 | 170_184 | 245_245 | 266_266 | 254_270 | 220_225 |
| 137_137 | 141_146 | 132_132 | 155_157 | 114_114 | 170_170 | 245_245 | 266_266 | 254_254 | 217_220 |
| 137_137 | 141_141 | 132_132 | 155_157 | 114_114 | 179_184 | 245_245 | 266_266 | 254_254 | 218_220 |
| 137_137 | 141_166 | 132_132 | 157_157 | 114_114 | 184_184 | 245_245 | 266_266 | 254_254 | 218_220 |
| 137_137 | 141_141 | 132_132 | 157_157 | 114_114 | 179_184 | 245_245 | 266_266 | 254_254 | 220_220 |
| 137_140 | 141_204 | 132_132 | 161_163 | 114_114 | 170_184 | 245_245 | 266_266 | 254_266 | 218_220 |
| 137_140 | 134_170 | 132_132 | 157_157 | 114_114 | 184_184 | 245_245 | 264_266 | 254_266 | 218_220 |

| | | | | | | | | | |
|---|---|---|---|---|---|---|---|---|---|
| NA | 144_144 | 132_132 | 157_157 | 114_122 | 170_170 | 245_245 | 266_266 | 254_254 | 220_220 |
| NA | 134_134 | 123_132 | 155_157 | 114_122 | 170_184 | 245_245 | 266_266 | 254_254 | 218_220 |
| 137_140 | 144_164 | 132_132 | 157_157 | 114_122 | 184_184 | 245_245 | 266_266 | 266_270 | 209_220 |
| 137_137 | 134_168 | 132_132 | 155_163 | 114_114 | 170_184 | 245_245 | 266_266 | 254_270 | 218_218 |
| 140_142 | NA | 132_132 | NA | NA | NA | NA | 266_266 | 254_254 | NA |
| NA | 134_141 | 116_132 | 155_157 | 114_114 | 170_184 | 245_245 | 266_266 | 254_254 | 220_220 |
| 137_137 | 141_144 | 132_132 | 155_157 | 114_122 | 184_184 | 245_245 | 266_266 | 254_266 | 220_220 |
| 137_140 | NA | 132_141 | NA | NA | NA | NA | 266_266 | 254_254 | NA |
| 137_137 | 134_144 | 132_132 | 155_157 | 114_114 | 184_184 | 245_245 | 266_266 | 254_254 | 209_220 |
| NA | 134_134 | 132_132 | 157_157 | 114_114 | 170_184 | 245_245 | 266_266 | 254_254 | 214_218 |
| 137_137 | 140_179 | 132_132 | 157_157 | 114_114 | 184_184 | 245_245 | 264_266 | 254_254 | 220_220 |
| NA | 141_170 | 132_132 | 157_163 | 114_114 | 170_184 | 245_245 | 266_266 | 254_254 | 218_220 |
| 140_140 | 179_188 | 123_132 | 157_157 | 114_114 | 184_184 | 245_245 | 264_266 | 254_254 | 220_225 |
| 137_137 | NA | 132_132 | NA | NA | NA | NA | 266_266 | 254_254 | NA |
| 137_137 | 141_170 | 132_132 | 155_157 | 114_114 | 170_184 | 245_245 | 266_266 | 254_270 | 217_218 |
| 137_137 | NA | 123_123 | NA | NA | NA | NA | 264_266 | 254_266 | NA |
| 137_137 | NA | 132_132 | NA | NA | NA | NA | 266_266 | 254_254 | NA |
| 137_137 | 134_141 | 132_132 | 155_157 | 114_114 | 179_184 | 245_245 | 266_266 | 254_254 | 225_225 |
| 137_140 | 144_168 | 132_132 | 155_159 | 114_114 | 184_184 | 245_245 | 266_266 | 254_270 | 218_218 |
| 137_142 | NA | 132_141 | NA | NA | NA | NA | 264_266 | 254_254 | NA |
| NA | 141_183 | NA | 159_161 | 114_114 | 179_184 | 245_245 | 266_266 | 254_254 | 209_220 |
| 137_140 | 134_199 | 123_132 | 155_157 | 114_122 | 179_184 | 245_245 | 266_266 | 254_266 | 209_209 |
| NA | 175_183 | 132_132 | 157_157 | 114_114 | 179_179 | 238_245 | 266_266 | 254_266 | 220_220 |
| 137_142 | 141_179 | 132_132 | 157_157 | 114_114 | 170_174 | 238_245 | 266_266 | 254_254 | 217_218 |
| 137_137 | 136_138 | 132_132 | 157_157 | 114_122 | 184_184 | 245_245 | 264_266 | 254_270 | 220_220 |
| 137_140 | 166_183 | 159_159 | 157_165 | 114_114 | 170_179 | 245_245 | 266_266 | 254_254 | 210_220 |
| 137_140 | 134_185 | 132_132 | 163_163 | 114_114 | 170_184 | 245_245 | 266_266 | 254_254 | 214_225 |
| 137_137 | 141_141 | 132_132 | 156_157 | 114_122 | 170_184 | 245_245 | 264_264 | 254_254 | 225_225 |
| 137_142 | 134_138 | NA | 157_163 | 114_114 | 179_184 | 245_245 | 264_266 | 254_254 | 220_225 |
| 137_137 | 134_141 | NA | 157_157 | 114_114 | 184_184 | 245_245 | 264_264 | 254_254 | 220_227 |
| 137_137 | 134_173 | 132_132 | 155_163 | 114_114 | 184_184 | 245_245 | 266_266 | 254_266 | 225_225 |
| NA | 134_170 | 132_132 | 157_159 | 114_114 | 174_184 | 245_245 | 264_266 | 254_254 | 225_225 |
| 137_137 | 134_144 | 132_132 | 157_157 | 114_114 | 179_179 | 245_245 | 264_266 | 254_254 | 220_220 |
| 140_140 | NA | 132_132 | NA | NA | NA | NA | 266_266 | 254_254 | NA |
| 133_140 | 141_168 | 132_132 | 157_159 | 114_114 | 179_186 | 245_245 | 266_266 | 254_259 | 220_220 |
| 137_137 | 175_177 | 132_132 | 155_157 | 114_114 | 170_170 | 245_245 | 264_266 | 254_270 | 220_220 |
| 137_137 | NA | 132_141 | NA | NA | NA | NA | 266_266 | 254_254 | NA |
| 137_137 | 141_144 | 132_132 | 157_163 | 114_114 | 184_186 | 245_245 | 266_266 | 266_270 | 220_220 |
| 137_140 | 144_144 | 129_132 | 155_157 | 114_114 | 184_184 | 238_245 | 264_266 | 254_270 | 220_220 |
| 137_137 | 134_168 | 132_132 | 157_157 | 114_114 | 179_184 | 245_245 | 266_266 | 254_254 | 217_220 |
| 140_144 | NA | 132_132 | NA | NA | NA | NA | 266_266 | 254_254 | NA |
| 137_137 | 134_141 | 132_132 | 155_163 | 114_114 | 170_179 | 245_245 | 266_268 | 254_270 | 217_220 |
| 137_137 | 141_141 | 132_132 | NA | 114_114 | NA | NA | 266_266 | 254_254 | 220_225 |
| 137_137 | 134_141 | 132_132 | 157_165 | 114_114 | 170_184 | 245_245 | 264_264 | 254_270 | 209_220 |
| 137_137 | 195_197 | 159_159 | 157_157 | 114_114 | 184_184 | 245_245 | 266_266 | 254_254 | 220_225 |
| 137_140 | 171_175 | 132_132 | 155_157 | 114_114 | 170_184 | 245_245 | 264_266 | 254_254 | 218_220 |
| 137_137 | 134_141 | 132_132 | 157_161 | 114_114 | 170_184 | 245_245 | 266_266 | 254_254 | 220_225 |
| 137_137 | 141_144 | 132_132 | 157_163 | 114_114 | 184_184 | 245_245 | 266_266 | 254_254 | 220_220 |

| AP273 | AP288 | AP289 | HB_C16_01 | HB_C16_02 | HB_C16_05 |
|---|---|---|---|---|---|
| 109_109 | 130_136 | 184_184 | 253_274 | 243_269 | 98_102 |
| 109_109 | 130_130 | 183_184 | 253_298 | 243_253 | 77_102 |
| 109_109 | 130_130 | 184_184 | 253_302 | 243_284 | 77_102 |
| 106_109 | 130_136 | 184_199 | 302_302 | 284_284 | 77_77 |
| 109_109 | 130_136 | 184_199 | 253_302 | 243_284 | 77_104 |
| 109_109 | 130_130 | 184_184 | 253_278 | 243_243 | 77_102 |
| 109_109 | 130_136 | 184_199 | 253_302 | 243_243 | 77_102 |
| 109_109 | 130_130 | 184_184 | 253_298 | 243_253 | 77_102 |
| 109_109 | 130_136 | 184_199 | 253_292 | 236_240 | 86_106 |
| 106_109 | 130_130 | 175_199 | 253_296 | 240_240 | 77_106 |
| 109_109 | 130_130 | 175_199 | 276_298 | 255_255 | 79_79 |
| 107_109 | 130_130 | 184_199 | 276_286 | 253_253 | 77_79 |
| 106_109 | 130_130 | 184_199 | 276_278 | 253_257 | 77_79 |
| 109_109 | 130_136 | 184_184 | 259_282 | 271_273 | 94_106 |
| 106_109 | 130_130 | 184_184 | 286_302 | 283_283 | 77_77 |
| 106_109 | 130_130 | 184_199 | 253_259 | 240_240 | 106_106 |
| 106_109 | 130_130 | 184_199 | 253_259 | 240_273 | 106_106 |
| 109_109 | 130_140 | 184_184 | 253_286 | 243_283 | 77_104 |
| 106_106 | 130_130 | 184_184 | 286_298 | 249_283 | 77_77 |
| 109_109 | 130_130 | 184_184 | 288_298 | 249_249 | 77_77 |
| 106_109 | 130_130 | 184_184 | 253_253 | 240_240 | 97_106 |
| 109_109 | 130_130 | 184_203 | 253_259 | 240_264 | 96_106 |
| 106_109 | 130_130 | 184_184 | 253_290 | 240_255 | 79_79 |
| 106_109 | 130_136 | 184_199 | 253_290 | 241_251 | 77_105 |
| 106_109 | 130_130 | 184_184 | 259_290 | 240_251 | 77_98 |
| 109_109 | 130_130 | 184_199 | 253_259 | 240_251 | 77_99 |
| 109_109 | 130_130 | 184_199 | 282_282 | 267_270 | 77_77 |
| 109_109 | 130_130 | 184_184 | 276_298 | 267_288 | 77_77 |
| 107_109 | 130_130 | 184_199 | 276_282 | 267_273 | 77_79 |
| 109_112 | 130_130 | 184_184 | 278_284 | 245_252 | 77_77 |
| 109_109 | 130_130 | 184_199 | 259_278 | 243_245 | 77_106 |
| 109_109 | 130_130 | 184_184 | 278_290 | 245_253 | 77_77 |
| 109_109 | 130_130 | 184_184 | 278_290 | 245_253 | 77_77 |
| 109_109 | 130_130 | 184_217 | 253_286 | 276_284 | 77_77 |
| 109_109 | 130_140 | 184_184 | 253_282 | 257_276 | 77_77 |
| 109_109 | 130_147 | 184_184 | 253_286 | 257_278 | 77_79 |
| 109_109 | 130_140 | 183_184 | 253_282 | 255_278 | 77_77 |
| 109_109 | 130_130 | 183_184 | 286_286 | 257_257 | 77_79 |
| 107_109 | 136_146 | 184_185 | 261_292 | 245_245 | 71_81 |
| 107_109 | 130_136 | 187_241 | 282_284 | 254_278 | 71_79 |
| 93_109 | 136_136 | 214_220 | NA | 265_273 | 79_79 |
| 109_109 | 136_136 | 185_185 | NA | 241_245 | 71_77 |
| 93_93 | 136_136 | NA | NA | 245_254 | 71_71 |
| 109_109 | 130_136 | 184_202 | 280_282 | 241_270 | 77_79 |
| 107_109 | 136_136 | 179_202 | NA | 245_245 | 71_79 |
| 107_109 | 130_136 | 185_231 | 263_282 | 245_257 | 71_79 |
| 109_109 | 136_136 | 198_221 | 261_282 | 245_249 | 71_77 |
| 107_109 | 146_146 | 184_189 | NA | 245_269 | 77_79 |

| | | | | | |
|---|---|---|---|---|---|
| 107_109 | 130_136 | 184_184 | 286_335 | 248_253 | 77_79 |
| 107_109 | 130_136 | 184_237 | 282_284 | 253_255 | 79_79 |
| 109_109 | 136_136 | 216_223 | NA | 244_245 | 71_79 |
| 107_109 | 136_136 | 187_241 | NA | 245_245 | 71_79 |
| 109_109 | 130_136 | 184_187 | 263_284 | 245_245 | 79_79 |
| 109_109 | 130_136 | 202_204 | NA | 251_267 | 79_79 |
| 107_109 | 130_136 | 184_184 | 276_286 | 293_299 | 79_79 |
| 107_109 | 136_136 | 187_198 | 282_286 | 241_245 | 79_81 |
| 107_109 | 136_136 | 198_231 | 261_280 | 245_269 | 71_79 |
| 107_109 | 136_146 | 239_239 | NA | 241_245 | 79_79 |
| 109_109 | 130_130 | 184_187 | 280_280 | 235_244 | 71_79 |
| 107_109 | 133_136 | 192_225 | 280_290 | 247_254 | 71_79 |
| 107_109 | 130_136 | 239_241 | NA | 241_278 | 71_71 |
| 109_109 | 136_146 | 231_252 | NA | 245_254 | 79_79 |
| 107_109 | 136_136 | 199_237 | 263_319 | 247_254 | 71_79 |
| 107_109 | 136_136 | 221_225 | 286_288 | 245_254 | 71_79 |
| 109_109 | 130_136 | 221_239 | 261_298 | 245_269 | 71_77 |
| 109_109 | 130_130 | 184_202 | 280_282 | 254_254 | 71_79 |
| 109_109 | 136_136 | 231_252 | NA | 245_245 | 71_79 |
| 109_109 | 130_136 | 185_227 | 261_286 | 269_275 | 71_77 |
| 109_109 | 130_130 | 185_231 | 282_300 | NA | 77_79 |
| 109_109 | 136_136 | 189_217 | NA | 245_247 | 79_81 |
| 109_109 | 136_136 | 214_241 | NA | 245_245 | 77_79 |
| 107_109 | 130_136 | 189_223 | NA | 254_261 | 77_79 |
| 107_109 | 136_136 | 187_217 | NA | 245_245 | 79_79 |
| 109_109 | 130_146 | 184_204 | 290_360 | 245_251 | 77_77 |
| 109_109 | 136_136 | 214_221 | NA | 265_267 | 71_79 |
| 109_109 | 130_136 | 184_239 | 284_321 | 241_275 | 79_81 |
| 109_109 | 130_133 | 227_231 | 286_300 | 236_236 | 77_79 |
| 109_109 | 136_136 | 187_223 | NA | 245_267 | 77_79 |
| 109_109 | 136_136 | 185_192 | 261_321 | 245_254 | 71_79 |
| 109_109 | 130_136 | 184_184 | 288_294 | 255_265 | 77_79 |
| 109_109 | 130_130 | 184_196 | NA | 251_263 | 77_79 |
| 109_109 | 136_136 | 202_237 | NA | 248_265 | 79_79 |
| 107_109 | 136_136 | 184_223 | 284_284 | 241_269 | 71_77 |
| 107_109 | 136_136 | 184_248 | NA | 245_275 | 71_79 |
| 109_109 | 130_130 | 184_187 | 282_290 | 240_253 | 71_79 |
| 109_109 | 130_136 | 184_210 | 280_296 | 265_265 | 77_77 |
| 109_109 | 130_136 | 187_189 | 317_333 | 244_253 | 71_79 |
| 109_109 | 136_136 | 183_192 | NA | 245_254 | 71_71 |
| 109_109 | 130_136 | 184_239 | 278_282 | 245_245 | 71_79 |
| 109_109 | 130_130 | 184_208 | 261_288 | 251_255 | 77_79 |
| 107_109 | 133_136 | 189_241 | 282_284 | 275_275 | 79_79 |
| 109_109 | 130_136 | 187_196 | 282_282 | 251_251 | 79_79 |
| 109_109 | 130_136 | 223_227 | 282_286 | 245_245 | 77_79 |
| 107_109 | 130_136 | 184_187 | 263_278 | 245_247 | 71_79 |
| 109_109 | 130_136 | 184_187 | 263_284 | 245_248 | 71_77 |
| 109_109 | 136_136 | 184_185 | 276_329 | 256_256 | 71_77 |

| | | | | | |
|---|---|---|---|---|---|
| 109_109 | 130_136 | 184_206 | 263_305 | 267_271 | 71_77 |
| 109_109 | NA | 184_202 | 282_290 | NA | 77_79 |
| 109_109 | 130_130 | 184_184 | 276_286 | 244_253 | 79_79 |
| 107_109 | 130_136 | 184_187 | 263_286 | 245_269 | 71_79 |
| 109_109 | 136_136 | 184_202 | 280_309 | 235_257 | 79_79 |
| 109_109 | 130_136 | 184_187 | 288_331 | 245_245 | 77_79 |
| 109_109 | 130_136 | 184_187 | 263_290 | 245_276 | 71_77 |
| 109_109 | 136_136 | 210_217 | NA | 245_245 | 71_79 |
| 107_109 | 130_136 | 187_189 | NA | 254_254 | 71_71 |
| 109_109 | 130_130 | 184_204 | 290_319 | 255_257 | 71_81 |
| 109_109 | 136_146 | 184_185 | 261_296 | 244_267 | 77_77 |
| 109_109 | 136_136 | 198_241 | NA | 246_271 | 77_79 |
| 109_109 | 136_136 | 185_216 | 263_282 | 245_254 | 77_79 |
| 109_109 | 130_136 | 184_187 | 282_282 | 245_248 | 71_77 |
| 109_109 | 130_136 | 184_187 | 261_288 | 265_281 | 77_77 |
| 109_109 | 130_136 | 187_237 | 305_311 | 235_275 | 71_77 |
| 109_109 | 130_130 | 184_239 | 282_286 | 257_263 | 77_77 |
| 107_109 | 130_130 | 184_239 | 276_286 | 271_275 | 77_79 |
| 107_109 | 130_136 | 185_239 | 282_290 | 253_275 | 77_79 |
| 107_109 | 130_136 | 189_189 | 319_331 | 245_247 | 71_79 |
| 107_109 | 130_136 | 184_187 | 288_288 | 241_276 | 71_79 |
| 109_109 | 130_130 | 184_190 | 288_290 | 257_261 | 77_79 |
| 107_107 | 136_136 | 199_227 | NA | 245_246 | 71_77 |
| 107_109 | 136_136 | 185_185 | 286_290 | 247_267 | 77_79 |
| 109_109 | 130_136 | 184_184 | 263_292 | 265_278 | 77_77 |
| 109_109 | 130_140 | 184_184 | 276_288 | 241_257 | 77_79 |
| 107_109 | 130_136 | 187_187 | 286_290 | 245_247 | 71_79 |
| 109_109 | 136_136 | 184_202 | 261_284 | 247_265 | 77_79 |
| 107_109 | 136_136 | 185_187 | 276_282 | 235_244 | 77_79 |
| 107_109 | 136_136 | 185_210 | NA | 241_254 | 77_77 |
| 109_109 | 130_130 | 184_184 | 288_290 | 248_271 | 77_79 |
| 93_109 | 136_136 | 187_190 | NA | 252_254 | 71_77 |
| 109_109 | 130_130 | 184_198 | 276_292 | 245_245 | 79_79 |
| 109_109 | 136_136 | 184_184 | 261_290 | 245_259 | 77_79 |
| 109_109 | 136_146 | 185_223 | NA | 244_244 | 77_79 |
| 109_109 | 130_130 | 187_235 | 263_282 | 247_257 | 77_77 |
| 107_109 | 136_136 | 185_223 | 280_305 | 241_245 | 71_79 |
| 107_109 | 136_136 | 184_184 | NA | 256_271 | 71_71 |
| 109_109 | 136_136 | 187_225 | 263_278 | 245_248 | 79_79 |
| 109_109 | 130_130 | 227_227 | 278_296 | 248_267 | 77_79 |
| 109_109 | 136_136 | 184_187 | NA | 254_254 | 71_79 |
| 109_109 | 130_136 | 184_184 | 261_284 | 251_291 | 79_79 |
| 109_109 | 130_136 | 184_187 | 282_286 | 254_261 | 71_79 |
| 107_109 | 133_136 | 198_227 | NA | 244_245 | 71_79 |
| 107_109 | 146_146 | 187_229 | NA | 245_254 | 71_77 |
| 107_109 | 136_145 | 221_227 | 261_261 | 245_248 | 77_79 |
| 109_109 | 136_136 | 233_237 | 282_338 | 245_245 | 71_71 |
| 109_109 | 130_136 | NA | 284_290 | 248_265 | 71_79 |

| | | | | | |
|---|---|---|---|---|---|
| 107_109 | 136_136 | 198_245 | 282_282 | 241_265 | 71_74 |
| 109_109 | 136_136 | 184_237 | 284_286 | 245_245 | 77_79 |
| 109_109 | 136_146 | 185_251 | 331_331 | 248_291 | 74_79 |
| 109_109 | 136_136 | 231_241 | 280_313 | 245_248 | 77_77 |
| 107_109 | NA | 187_241 | 300_311 | NA | 77_79 |
| 109_109 | 130_136 | 185_199 | NA | 245_254 | 79_79 |
| 107_109 | 136_136 | 185_231 | NA | 245_245 | 79_79 |
| 107_109 | 136_136 | 187_198 | NA | 245_248 | 71_77 |
| 107_109 | 136_136 | 189_233 | NA | 245_254 | 71_79 |
| 107_109 | 136_136 | 183_184 | NA | 245_291 | 79_79 |
| 109_109 | 130_136 | 185_221 | 278_338 | 244_257 | 71_77 |
| 107_107 | 136_136 | 184_227 | NA | 245_245 | 71_79 |
| 109_109 | 130_136 | 185_185 | NA | 245_254 | 71_79 |
| 107_109 | 136_136 | 208_210 | NA | 245_245 | 77_77 |
| 109_109 | 136_136 | 204_223 | NA | 244_246 | 71_71 |
| 109_109 | 130_130 | 185_189 | NA | 246_246 | 77_79 |
| 109_109 | 130_136 | 187_231 | NA | 245_245 | 71_79 |
| 109_109 | 136_136 | 198_202 | NA | 244_245 | 79_79 |
| 107_109 | 136_136 | 206_216 | NA | 241_247 | 71_71 |
| 107_109 | 136_136 | 229_229 | 284_298 | 245_245 | 71_79 |
| NA | 136_136 | 190_231 | NA | 241_244 | 71_79 |
| 109_109 | 130_136 | 185_187 | 261_313 | 276_293 | 71_77 |
| 109_109 | 136_136 | 216_235 | NA | 247_254 | 71_77 |
| 109_109 | 136_136 | 183_184 | NA | 244_245 | 71_79 |
| 107_109 | 133_136 | 184_185 | NA | 245_247 | 71_71 |
| 109_109 | 136_136 | 184_184 | NA | 241_245 | 71_77 |
| 107_109 | 130_136 | 184_192 | NA | 254_276 | 71_76 |
| 107_109 | 136_136 | 208_231 | NA | 244_250 | 77_81 |
| 109_109 | 136_136 | 185_216 | 280_302 | 245_252 | 71_71 |
| 107_109 | 136_136 | 185_206 | 338_340 | 245_254 | 77_79 |
| 107_109 | 136_136 | 187_239 | 280_300 | 245_245 | 71_71 |
| 107_109 | 136_146 | 239_239 | NA | 244_245 | 79_79 |
| 107_109 | 136_136 | 187_239 | NA | 245_254 | 71_71 |
| 107_109 | 136_136 | 235_239 | NA | 254_254 | 71_71 |
| 107_109 | 136_136 | 235_239 | NA | 245_245 | 71_71 |
| 107_107 | 136_136 | 185_233 | 278_292 | 245_246 | 71_71 |
| 107_109 | 136_136 | 198_231 | 261_284 | 241_245 | 77_79 |
| 107_109 | 136_136 | 223_229 | NA | 245_245 | 71_79 |
| 107_109 | 136_136 | 185_225 | 282_284 | 235_245 | 71_71 |
| 109_109 | 136_136 | 185_220 | 261_309 | 245_248 | 71_79 |
| 107_109 | 136_136 | 189_233 | 313_323 | 245_245 | 79_79 |
| 107_109 | 136_136 | 231_233 | 309_323 | 245_245 | 71_79 |
| 107_109 | 136_136 | 187_235 | 261_284 | 244_245 | 71_77 |
| 107_109 | 130_136 | 199_199 | 296_311 | 245_254 | 71_77 |
| 107_107 | 130_146 | 187_231 | 298_333 | 245_248 | 71_71 |
| 107_109 | 130_136 | 185_185 | NA | 254_254 | 79_79 |
| 107_109 | 136_136 | 187_206 | NA | 241_256 | 77_77 |
| 107_109 | 130_132 | NA | NA | 245_245 | 71_79 |

| | | | | | |
|---|---|---|---|---|---|
| 109_109 | 130_136 | 212_227 | NA | 245_254 | 71_71 |
| 107_109 | 136_136 | 189_225 | NA | 245_265 | 77_79 |
| 107_109 | 136_136 | 184_185 | 263_282 | 245_254 | 71_71 |
| 107_109 | 130_136 | 187_190 | NA | 245_251 | 77_79 |
| 109_109 | 136_136 | 185_198 | 261_282 | 241_256 | 77_77 |
| 107_107 | 136_136 | 184_196 | 263_263 | 244_245 | 71_79 |
| 107_109 | 136_136 | 184_235 | NA | 254_254 | 71_71 |
| 109_109 | 133_136 | 189_221 | NA | 244_250 | 71_77 |
| 109_109 | 136_136 | 184_185 | NA | 245_299 | 71_79 |
| 107_107 | 130_146 | 221_231 | NA | 245_245 | 71_77 |
| 109_109 | 136_136 | 185_217 | NA | 244_245 | 79_79 |
| 109_109 | 136_140 | 216_225 | NA | 245_253 | 79_79 |
| 107_109 | 136_136 | 187_216 | NA | 244_245 | 71_71 |
| 109_109 | 136_136 | 260_260 | NA | 244_254 | 71_79 |
| 109_109 | 136_136 | 187_223 | NA | 236_245 | 77_79 |
| 107_109 | 136_146 | 187_216 | NA | 245_254 | 77_79 |
| 109_109 | 130_136 | 189_229 | 280_292 | 244_254 | 71_79 |
| 107_107 | 136_136 | 184_187 | NA | 241_245 | 71_79 |
| 107_109 | 130_136 | 182_229 | NA | 236_245 | 71_79 |
| 107_109 | 136_136 | 187_231 | NA | 235_245 | 71_79 |
| 107_109 | 136_136 | 185_192 | NA | 245_245 | 71_77 |
| 109_109 | 130_136 | 185_189 | NA | 241_245 | 71_79 |
| 107_109 | 136_136 | 185_190 | NA | 245_245 | 77_79 |
| 107_109 | 136_136 | 187_225 | NA | 245_254 | 71_79 |
| 109_109 | 130_136 | 227_229 | NA | 244_244 | 71_79 |
| 107_109 | 133_136 | 184_198 | NA | 245_245 | 71_79 |
| 107_109 | 136_146 | 217_235 | NA | 241_245 | 79_79 |
| 109_109 | 136_136 | 189_212 | NA | 247_254 | 71_71 |
| 107_109 | 136_136 | 185_202 | NA | 245_245 | 71_77 |
| 107_109 | 136_136 | 187_231 | NA | 254_286 | 71_79 |
| 107_109 | 136_136 | 190_199 | NA | 245_254 | 71_77 |
| 109_109 | 136_146 | 184_187 | NA | 245_255 | 79_79 |
| 109_109 | 136_136 | 185_223 | NA | 245_246 | 77_79 |
| 109_109 | 136_136 | 187_187 | 280_282 | 245_245 | 71_77 |
| 107_107 | 130_136 | 198_235 | 280_282 | 245_245 | 77_77 |
| 109_109 | 136_136 | 190_221 | 261_282 | 246_256 | 71_79 |
| 109_109 | 136_136 | 187_189 | 282_282 | 245_252 | 77_79 |
| 109_109 | 136_136 | 185_185 | 282_282 | 245_254 | 77_79 |
| 107_109 | 133_136 | 185_189 | 282_284 | 241_241 | 71_71 |
| 109_109 | 130_136 | 189_225 | 280_282 | 245_252 | 77_79 |
| 109_109 | 130_136 | 184_185 | 282_282 | 245_245 | 71_71 |
| 107_109 | 136_136 | 185_185 | NA | 241_254 | 71_71 |
| 109_109 | 130_130 | 194_235 | NA | 254_255 | 71_77 |
| 109_109 | 133_136 | 223_227 | NA | 244_254 | 71_77 |
| 109_109 | 136_136 | 185_229 | NA | 245_256 | 79_79 |
| 107_109 | 136_136 | 187_233 | NA | 254_275 | 71_79 |
| 109_109 | 136_146 | 185_220 | 286_337 | 248_284 | 71_77 |
| 107_107 | 136_136 | 189_237 | 282_302 | 244_245 | 71_71 |

| | | | | | |
|---|---|---|---|---|---|
| 107_109 | 130_136 | 189_208 | 263_288 | 245_245 | 71_79 |
| 107_107 | 136_136 | 187_223 | 282_282 | 244_245 | 71_71 |
| 107_109 | 136_136 | 189_189 | 282_284 | 254_254 | 71_71 |
| 107_109 | 136_136 | 189_223 | NA | 245_254 | 71_71 |
| 107_109 | 136_136 | 185_187 | 282_300 | 244_244 | 79_79 |
| 107_107 | 130_136 | 187_202 | 276_282 | 245_248 | 79_79 |
| 107_107 | 136_136 | 185_187 | 282_327 | 245_254 | 77_79 |
| 107_107 | 136_140 | 187_239 | 261_282 | 245_245 | 77_79 |
| 109_109 | 136_136 | 187_187 | 280_282 | 236_244 | 77_79 |
| 109_109 | 130_136 | 235_237 | 261_276 | 245_269 | 77_79 |
| 107_109 | 136_136 | 189_245 | 282_321 | 236_245 | 71_79 |
| 107_107 | 136_136 | 202_239 | 261_276 | 236_244 | 77_79 |
| 109_109 | NA | 187_187 | 261_321 | NA | 77_79 |
| 107_109 | 136_140 | 185_187 | 282_282 | 251_267 | 71_71 |
| 109_109 | 136_136 | 189_223 | NA | 254_254 | 71_79 |
| 107_109 | 136_136 | 198_199 | NA | 241_245 | 71_79 |
| 109_109 | 136_136 | 185_189 | NA | 246_256 | 71_79 |
| 109_109 | 130_136 | 190_190 | NA | 245_254 | 71_71 |
| 107_107 | 130_140 | 185_192 | 282_304 | 245_245 | 71_79 |
| 107_109 | 133_136 | 187_198 | 282_284 | 236_244 | 71_77 |
| 107_109 | 133_140 | 189_189 | 261_282 | 245_254 | 71_77 |
| 107_109 | 136_136 | 210_237 | 282_282 | 244_245 | 77_79 |
| 107_109 | 136_136 | 198_247 | NA | 235_273 | 71_71 |
| 109_109 | 136_136 | 184_210 | NA | 248_254 | 71_79 |
| 107_109 | 136_146 | 214_251 | 284_304 | 255_269 | 77_79 |
| 107_109 | 136_136 | 184_220 | NA | 245_245 | 71_79 |
| 107_109 | 136_136 | 208_214 | 284_335 | 245_269 | 77_79 |
| 107_109 | 136_136 | 187_190 | NA | NA | 71_71 |
| 107_109 | 136_136 | 241_242 | 284_340 | 244_271 | 71_71 |
| 107_109 | 136_136 | 187_216 | 280_309 | 244_247 | 71_71 |
| 109_109 | 136_136 | 212_247 | 305_346 | 244_256 | 71_79 |
| 109_109 | 136_136 | 220_221 | 282_286 | 244_250 | 71_77 |
| 109_109 | 136_140 | 185_187 | 280_280 | 254_267 | 71_77 |
| 109_109 | 136_136 | 187_187 | 305_325 | NA | 71_77 |
| 109_109 | 136_136 | 187_198 | NA | 253_283 | 71_79 |
| 109_109 | 136_140 | 183_184 | NA | 244_250 | 71_79 |
| 109_109 | 136_136 | 210_248 | NA | 244_250 | 71_79 |
| 109_109 | 136_136 | 185_225 | NA | 271_275 | 79_79 |
| 107_109 | 133_136 | 185_239 | NA | NA | 77_79 |
| 109_109 | 130_136 | 189_202 | NA | 245_254 | 71_71 |
| 109_109 | 136_140 | 189_206 | NA | 236_265 | 77_79 |
| 107_109 | 136_136 | 184_185 | NA | 244_267 | 77_79 |
| 109_109 | 136_136 | 199_239 | NA | 245_246 | 71_71 |
| 109_109 | 136_136 | 208_245 | NA | 245_248 | 71_79 |
| 107_109 | 136_136 | 187_223 | 280_284 | 245_248 | 71_77 |
| 109_109 | 136_136 | 187_231 | NA | 245_265 | 71_79 |
| 109_109 | 136_146 | 229_229 | NA | 241_261 | 77_77 |
| 107_109 | 136_136 | 184_204 | NA | 244_244 | 77_79 |

| | | | | | |
|---|---|---|---|---|---|
| 107_109 | 133_136 | 204_237 | NA | 265_265 | 71_77 |
| 109_109 | 136_136 | NA | 307_309 | 245_247 | 71_71 |
| 109_109 | 136_146 | 185_185 | 282_296 | 245_254 | 71_71 |
| 109_109 | 140_146 | 210_210 | 286_352 | 235_241 | 71_77 |
| 109_109 | 136_136 | 217_223 | 261_284 | 254_256 | 71_79 |
| 107_107 | 136_136 | 190_190 | NA | 244_245 | 71_77 |
| 107_109 | 136_136 | 225_233 | NA | 236_244 | 79_79 |
| 107_109 | 136_136 | 212_241 | NA | 244_250 | 77_77 |
| 109_109 | 136_136 | 187_187 | NA | 245_254 | 71_77 |
| 107_107 | 133_136 | 231_245 | NA | 245_245 | 71_79 |
| 107_109 | 136_146 | 184_184 | NA | 241_248 | 71_77 |
| 109_109 | 136_136 | 187_198 | NA | 245_267 | 79_79 |
| 109_109 | 136_136 | 189_235 | NA | 241_245 | 77_79 |
| 109_109 | 130_140 | 189_189 | NA | 250_267 | 79_79 |
| 109_109 | 136_136 | 190_199 | NA | 259_271 | 77_77 |
| 109_109 | 136_136 | 221_242 | NA | 245_245 | 79_79 |
| 109_109 | 136_136 | 223_239 | NA | 245_245 | 71_74 |
| 107_109 | 136_136 | 187_208 | NA | 245_245 | 71_71 |
| 107_107 | 136_146 | 185_235 | NA | 248_248 | 71_79 |
| 107_109 | 136_136 | 184_198 | NA | 245_245 | 79_79 |
| 109_109 | 136_146 | 190_225 | NA | 241_256 | 79_79 |
| 107_109 | 136_136 | 185_208 | NA | 241_245 | 79_79 |
| 122_122 | 136_136 | NA | NA | 248_256 | NA |
| 107_109 | 130_136 | 199_203 | NA | 244_255 | 71_79 |
| 109_109 | 136_146 | 184_187 | NA | 245_256 | 71_79 |
| 107_109 | 136_136 | 185_185 | NA | 245_273 | 71_77 |
| 109_109 | 136_140 | 223_223 | NA | 241_254 | 79_79 |
| 109_109 | 136_136 | 185_247 | NA | 248_248 | 71_71 |
| 109_109 | 136_140 | 185_185 | NA | 245_256 | 71_77 |
| 107_109 | 136_136 | 184_185 | NA | 254_267 | 77_79 |
| 107_109 | 136_136 | 184_187 | NA | 256_296 | 79_79 |
| 107_109 | 136_136 | 185_187 | NA | 245_245 | 77_77 |
| 109_109 | 136_146 | 184_235 | NA | 245_288 | 71_79 |
| 107_109 | 136_136 | 185_185 | NA | 245_256 | 79_79 |
| 107_109 | 136_136 | 185_190 | NA | 245_255 | 71_77 |
| 109_109 | 136_136 | 187_229 | NA | 241_245 | 71_79 |
| 109_109 | 130_136 | 185_220 | NA | 245_245 | 77_79 |
| 107_109 | 136_136 | 192_220 | NA | 245_245 | 71_71 |
| 107_109 | 136_136 | 185_185 | NA | 245_254 | 71_77 |
| 107_109 | 136_136 | 187_187 | NA | 245_248 | 74_79 |
| 107_109 | 136_140 | 187_192 | NA | 245_254 | 71_79 |
| 107_109 | 136_146 | 185_187 | NA | 245_249 | 71_79 |
| 109_109 | 136_146 | 245_245 | NA | 245_245 | 71_77 |
| 107_109 | 136_136 | 185_206 | NA | 245_254 | 71_71 |
| 107_109 | 136_136 | 183_185 | NA | 245_255 | 71_77 |
| 109_109 | 136_140 | 187_189 | NA | 249_276 | 71_71 |
| 107_107 | 136_136 | 185_190 | NA | 245_284 | 71_71 |
| 107_109 | 136_136 | 225_225 | NA | 245_245 | 71_71 |

| | | | | | |
|---|---|---|---|---|---|
| 107_109 | 136_136 | 189_237 | NA | 245_283 | 79_79 |
| 107_109 | 136_136 | 189_233 | NA | 244_267 | 71_71 |
| 107_109 | 136_136 | 185_185 | NA | 245_256 | 71_79 |
| 107_109 | 136_136 | 187_187 | NA | 244_270 | 71_71 |
| 109_109 | 136_136 | 229_242 | NA | 241_245 | 71_77 |
| 109_109 | 136_136 | 187_235 | NA | 245_286 | 79_79 |
| 107_109 | 136_136 | 184_223 | NA | 245_248 | 71_71 |
| 109_109 | 136_136 | 198_237 | NA | 244_245 | 71_71 |
| 107_109 | 130_136 | 185_229 | NA | 241_245 | 77_79 |
| 107_107 | 136_136 | 183_199 | NA | 245_256 | 71_79 |
| 109_109 | 136_136 | 185_187 | NA | 278_278 | 71_71 |
| 107_109 | 136_136 | 187_194 | NA | 248_256 | 71_71 |
| 107_109 | 136_140 | 185_187 | NA | 254_276 | 71_71 |
| 107_109 | 136_136 | 185_189 | NA | 244_248 | 71_71 |
| 107_109 | 136_136 | 229_233 | 315_317 | 249_249 | 77_77 |
| 107_109 | 136_136 | 185_185 | NA | 245_259 | 71_77 |
| 109_112 | 136_140 | 185_245 | 280_315 | 245_248 | 71_81 |
| 107_109 | 136_136 | 185_185 | 261_309 | 245_273 | 77_79 |
| 107_109 | 136_136 | 184_185 | 282_315 | 245_245 | 71_79 |
| 107_107 | 136_136 | 185_212 | 261_300 | 241_250 | 71_79 |
| 107_107 | 136_136 | 185_199 | NA | 248_284 | 71_71 |
| 107_107 | 136_140 | 187_187 | NA | 241_248 | 77_79 |
| 107_107 | 130_136 | 187_208 | NA | 245_245 | 71_79 |
| 107_107 | 136_146 | 184_225 | 307_319 | 247_247 | 71_79 |
| 107_109 | 136_136 | 185_187 | 286_315 | 245_247 | 71_77 |
| 107_109 | 136_136 | 184_221 | 290_315 | 245_245 | 71_71 |
| 109_109 | 136_140 | 190_190 | NA | 246_265 | 77_79 |
| 107_107 | 136_136 | 183_185 | 325_333 | 245_276 | 71_79 |
| 107_107 | 136_136 | 184_184 | 292_315 | 288_288 | 79_79 |
| 106_109 | 136_136 | 189_221 | 305_319 | 261_299 | 71_79 |
| 107_109 | 136_136 | 185_231 | 313_333 | 248_256 | 71_79 |
| 107_116 | 136_136 | 185_185 | NA | 245_245 | 71_71 |
| 107_109 | 136_136 | 185_185 | 298_307 | 247_267 | 79_79 |
| 107_109 | 136_136 | 185_185 | 313_329 | 273_273 | 77_79 |
| 107_107 | 136_136 | 185_187 | 309_309 | 248_248 | 71_71 |
| 106_109 | 136_136 | 185_185 | 294_348 | 248_261 | 79_79 |
| 107_107 | 136_136 | 217_223 | 282_315 | 248_283 | 71_79 |
| 107_107 | 136_136 | 184_185 | NA | 288_288 | 71_79 |
| 109_109 | 136_136 | 185_220 | 302_348 | 248_248 | 71_79 |
| 107_109 | 136_140 | 212_235 | NA | 245_245 | 77_79 |
| 107_107 | 136_136 | 185_185 | 290_325 | 248_248 | 71_79 |
| 107_107 | 136_136 | 221_242 | NA | 246_248 | 71_71 |
| 107_107 | 136_136 | NA | NA | 244_248 | 71_71 |
| 107_109 | 136_136 | 185_185 | 302_325 | 245_276 | 71_71 |
| 109_109 | 136_136 | 185_233 | NA | 249_249 | 77_79 |
| 107_109 | 130_136 | 184_185 | NA | 244_245 | 71_71 |
| 107_109 | 136_136 | 185_185 | NA | 248_248 | 71_79 |
| 109_109 | 136_136 | 185_220 | NA | 256_256 | 79_79 |

| | | | | | |
|---|---|---|---|---|---|
| 107_109 | 136_136 | 185_187 | NA | 244_249 | 71_71 |
| 107_107 | 136_136 | 202_202 | NA | 241_245 | 71_79 |
| 109_109 | 136_136 | 185_187 | NA | 284_284 | 71_79 |
| 109_109 | 136_136 | 185_187 | NA | 244_254 | 71_77 |
| 107_107 | 136_136 | 184_223 | NA | 248_249 | 71_71 |
| 107_109 | 136_136 | 185_187 | NA | 244_247 | 77_77 |
| 107_109 | 130_136 | 184_190 | NA | 251_254 | 71_77 |
| 107_109 | 136_136 | 187_187 | NA | 245_245 | 79_79 |
| 107_107 | 130_140 | 187_217 | NA | 248_254 | 79_79 |
| 109_109 | 136_136 | 187_231 | NA | 245_245 | 71_77 |
| 109_109 | 136_136 | 185_185 | NA | 245_254 | 71_71 |
| 106_109 | 133_136 | 185_190 | NA | 248_248 | 71_79 |
| 107_107 | 136_136 | 229_231 | NA | 245_286 | 71_79 |
| 107_109 | 136_136 | 187_190 | NA | 245_254 | 71_79 |
| 107_107 | 136_136 | 185_233 | NA | 247_276 | 71_71 |
| 107_109 | 140_146 | 233_233 | NA | 241_245 | 71_71 |
| 107_107 | 136_136 | 231_237 | NA | 248_248 | 71_71 |
| 107_107 | 130_136 | 185_237 | NA | 235_248 | 79_79 |
| 107_107 | 136_136 | 185_239 | NA | 284_284 | 71_74 |
| 109_109 | 136_136 | 187_187 | NA | 253_288 | 71_79 |
| 109_109 | 136_140 | 185_185 | NA | 245_258 | 71_71 |
| 109_109 | 130_136 | 184_187 | NA | 243_276 | 71_77 |
| 107_109 | 136_136 | 198_198 | NA | 245_281 | 79_79 |
| 107_109 | 136_136 | 184_185 | NA | 248_281 | 71_71 |
| 109_109 | 136_136 | NA | NA | 254_267 | 71_79 |
| 107_107 | 136_136 | 185_185 | NA | 248_281 | 79_79 |
| 106_107 | 136_136 | 185_187 | NA | 244_248 | 71_79 |
| 107_109 | 130_136 | 185_237 | NA | 244_245 | 71_77 |
| 109_109 | 136_140 | 223_223 | NA | 245_266 | 71_71 |
| 107_109 | 133_136 | 185_208 | NA | NA | 71_79 |
| 107_109 | 136_136 | 183_187 | NA | 241_254 | 71_79 |
| 107_107 | 136_136 | 184_192 | NA | 244_264 | 71_79 |
| 109_109 | 136_136 | 187_187 | NA | 244_278 | 79_79 |
| 109_109 | 136_136 | 225_233 | NA | 248_248 | 71_79 |
| 107_109 | 136_136 | 227_227 | NA | 245_245 | 71_79 |
| 109_109 | 136_140 | 187_221 | NA | 248_281 | 79_79 |
| 107_109 | 136_136 | 185_185 | NA | NA | 71_71 |
| 107_107 | 136_136 | 185_235 | NA | 245_248 | 79_81 |
| 109_109 | 130_136 | 185_185 | NA | 246_263 | 71_71 |
| 109_109 | 136_136 | 212_212 | NA | 241_241 | 79_79 |
| 107_109 | 136_136 | 185_194 | NA | 256_256 | 71_77 |
| 107_109 | 136_136 | 185_202 | NA | 245_246 | 79_79 |
| 107_107 | 136_136 | 184_194 | NA | 245_248 | 71_71 |
| 107_109 | 136_136 | 225_225 | NA | 247_247 | 71_71 |
| 109_109 | NA | 206_206 | NA | NA | 71_79 |
| 107_109 | 136_136 | 187_198 | NA | 245_248 | 71_79 |
| 107_109 | 136_136 | 216_220 | NA | 244_257 | 71_71 |
| 107_107 | 136_136 | NA | NA | 244_284 | 71_79 |

| | | | | | |
|---|---|---|---|---|---|
| 107_109 | 136_136 | 233_239 | NA | 245_286 | 71_79 |
| 107_107 | 136_136 | 185_187 | NA | 254_256 | 71_79 |
| 107_109 | 136_140 | 262_264 | NA | 256_256 | 77_77 |
| 109_109 | 136_136 | 185_233 | NA | 245_296 | 71_77 |
| 109_109 | 136_136 | 185_196 | NA | 245_245 | 71_77 |
| 109_109 | 130_136 | 225_225 | 263_337 | 245_254 | 71_71 |
| 107_107 | 130_136 | NA | 280_282 | 245_245 | 79_79 |
| 109_109 | 136_136 | 189_189 | NA | 244_245 | 71_71 |
| 107_109 | 130_136 | 187_187 | NA | 245_256 | 71_71 |
| 109_109 | 136_136 | 241_241 | NA | 245_261 | 71_79 |
| 106_109 | 136_136 | 187_207 | NA | 272_272 | 71_79 |
| 107_107 | 136_136 | 185_235 | NA | 245_261 | 71_79 |
| 107_107 | 136_140 | 185_199 | NA | 245_254 | 71_77 |
| 107_109 | 136_136 | 185_235 | NA | 248_278 | 71_77 |
| 109_109 | 130_136 | 184_184 | 282_290 | 245_245 | 71_77 |
| 107_109 | 136_136 | 189_189 | 300_305 | 244_248 | 71_79 |
| 109_109 | 130_136 | 184_184 | 282_319 | 245_245 | 71_71 |
| 107_109 | 133_133 | 187_187 | 309_340 | 245_254 | 71_71 |
| 109_109 | 136_136 | 223_223 | 282_286 | 245_245 | 71_71 |
| 109_109 | 136_136 | 187_189 | NA | 241_254 | 77_79 |
| 107_109 | 136_136 | 185_185 | NA | 272_272 | 71_77 |
| 107_107 | 130_136 | 185_190 | 298_313 | 254_286 | 71_71 |
| 107_109 | 130_136 | 231_231 | 282_298 | 245_254 | 71_71 |
| 109_109 | 136_136 | 199_242 | 278_288 | 241_254 | 71_77 |
| 109_109 | 136_136 | 187_221 | NA | 245_254 | 71_79 |
| 107_109 | 136_136 | NA | 302_331 | 244_254 | 79_79 |
| NA | 136_136 | NA | 280_282 | 245_256 | NA |
| 109_109 | 136_136 | 184_212 | 300_305 | 244_245 | 71_77 |
| 107_109 | 136_140 | 227_242 | NA | 245_245 | 71_77 |
| 107_107 | 140_140 | 192_192 | 278_280 | 235_248 | 79_79 |
| 107_109 | 136_136 | 187_187 | 261_300 | 244_245 | 71_71 |
| 107_109 | 136_136 | 185_185 | NA | 278_284 | 79_79 |
| 107_109 | 136_136 | 185_239 | NA | 245_267 | 71_71 |
| 107_109 | 136_136 | 185_187 | NA | 245_256 | 71_71 |
| 109_109 | 136_136 | 185_190 | NA | 245_254 | 71_71 |
| 107_109 | 136_140 | 231_255 | NA | 254_254 | 71_77 |
| 107_109 | 136_136 | 183_183 | 280_325 | 244_254 | 71_71 |
| 107_109 | 136_136 | 185_241 | 282_307 | 245_271 | 71_77 |
| 107_109 | 136_136 | 185_221 | 305_315 | 245_245 | 71_71 |
| 107_107 | 136_136 | 185_185 | 305_305 | 254_271 | 71_79 |
| 107_109 | 136_136 | 187_216 | 282_317 | 241_244 | 71_71 |
| NA | 136_136 | NA | 282_282 | 245_269 | NA |
| 109_109 | 130_136 | 241_241 | NA | 254_265 | 77_79 |
| 107_109 | 136_140 | 185_185 | NA | 254_254 | 71_71 |
| 107_109 | 136_136 | 185_221 | NA | 256_256 | 71_71 |
| 107_107 | 136_136 | 185_185 | NA | 245_281 | 71_77 |
| 107_107 | 136_136 | 239_239 | 263_321 | 254_254 | 71_79 |
| 109_109 | 136_136 | 185_185 | 286_315 | 245_248 | 71_71 |

| | | | | | |
|---|---|---|---|---|---|
| 107_107 | 130_136 | 194_221 | 319_331 | 244_245 | 79_79 |
| 107_109 | 136_136 | 184_184 | 309_313 | 256_256 | 71_71 |
| 107_109 | 130_136 | 214_221 | NA | 241_265 | 71_79 |
| 109_109 | 136_136 | 187_229 | NA | 245_254 | 71_79 |
| 109_109 | 136_136 | NA | NA | 245_254 | 71_71 |
| 107_109 | 136_136 | 192_229 | NA | 245_264 | 79_79 |
| 109_109 | 136_136 | 237_237 | NA | 241_245 | 71_79 |
| 107_107 | 136_136 | 185_242 | NA | 245_245 | 77_77 |
| 109_109 | 136_136 | 187_247 | NA | 245_254 | 71_79 |
| 109_109 | 130_136 | 235_235 | NA | 245_288 | 71_79 |
| 107_109 | 136_140 | 185_185 | NA | 245_245 | 71_77 |
| 107_109 | 136_136 | 184_235 | NA | 276_286 | 71_79 |
| 107_109 | 136_136 | 216_216 | NA | 244_244 | 71_79 |
| 107_109 | 136_136 | 184_187 | NA | 236_245 | 71_79 |
| 107_109 | 136_136 | 187_187 | NA | 275_281 | 71_79 |
| 107_109 | 136_136 | 190_235 | NA | 245_284 | 71_79 |
| 109_109 | 130_136 | 187_221 | NA | 245_258 | 79_79 |
| 109_109 | 136_136 | 231_239 | NA | 248_284 | 79_79 |
| 107_109 | 136_136 | 187_248 | NA | 256_256 | 71_79 |
| 107_107 | 136_136 | 185_233 | NA | 249_262 | 71_79 |
| 109_109 | 136_136 | 185_185 | NA | NA | 71_77 |
| 109_109 | 136_136 | 187_199 | NA | 244_278 | 71_79 |
| 107_109 | 136_136 | 235_235 | NA | 254_265 | 79_79 |
| 107_109 | 136_136 | 190_190 | NA | 245_245 | 71_71 |
| 107_109 | 130_136 | 231_231 | 284_305 | 244_245 | 71_77 |
| 109_109 | 136_136 | 189_189 | 284_354 | 254_256 | 71_71 |
| 107_109 | 136_136 | 187_187 | NA | 254_254 | 79_79 |
| 109_109 | 136_140 | 187_189 | NA | 245_254 | 71_79 |
| 109_109 | 136_136 | 237_237 | NA | 254_254 | 71_71 |
| 109_109 | 136_136 | 190_227 | 311_321 | 254_254 | 70_79 |
| 109_109 | 136_136 | 192_245 | 305_311 | 245_254 | 77_79 |
| 107_109 | 136_136 | 185_185 | NA | 241_254 | 71_79 |
| 107_109 | 136_136 | NA | NA | 248_258 | 77_79 |
| 109_109 | 136_140 | 187_187 | 282_305 | 245_254 | 71_79 |
| 107_109 | 130_136 | 187_187 | 278_282 | 245_245 | 71_77 |
| 107_109 | 130_136 | 187_187 | 278_304 | 245_245 | 71_71 |
| 109_109 | 136_140 | 185_196 | 282_337 | 254_254 | 71_79 |
| 109_109 | 136_140 | 227_229 | 282_321 | 245_254 | 71_79 |
| 109_109 | 136_136 | 185_233 | NA | 244_245 | 71_77 |
| 109_112 | 136_140 | 185_185 | 280_309 | 248_254 | 71_71 |
| 109_109 | 136_140 | 196_223 | 286_305 | 245_252 | 71_71 |
| 109_109 | 136_136 | NA | 282_309 | 254_254 | 71_71 |
| 107_109 | 136_140 | NA | 305_317 | 245_245 | 71_71 |
| 109_109 | 136_136 | 237_237 | 305_309 | 245_254 | 71_79 |
| 109_109 | 136_136 | 233_235 | 305_315 | 245_245 | 71_79 |
| 109_112 | 136_136 | 190_190 | NA | 254_258 | 71_71 |
| 109_109 | 136_136 | 241_241 | 309_317 | 254_273 | 79_79 |
| 107_109 | 136_136 | NA | 302_317 | 245_245 | 71_79 |

| | | | | | |
|---|---|---|---|---|---|
| 109_109 | 136_136 | 187_187 | 282_313 | 245_252 | 71_71 |
| 109_109 | 136_136 | 185_185 | 282_305 | 245_245 | 71_79 |
| 107_107 | 136_136 | 216_216 | NA | 247_267 | 77_79 |
| 107_109 | 136_136 | 185_185 | NA | 247_254 | 79_112 |
| 109_109 | 136_136 | 235_251 | 309_315 | 245_245 | 71_71 |
| 107_109 | 136_136 | 233_242 | NA | 245_245 | 71_77 |
| 107_109 | 136_140 | 185_185 | 307_335 | 247_247 | 71_71 |
| 109_109 | 136_136 | NA | 309_309 | 245_245 | 71_71 |
| 109_109 | 136_136 | 227_227 | 282_309 | 245_254 | 71_79 |
| 109_109 | 136_136 | 227_242 | 307_309 | 245_245 | 71_71 |
| 107_109 | 136_136 | NA | 282_309 | 245_245 | 71_71 |
| 109_109 | 136_136 | NA | 309_309 | 245_245 | 71_71 |
| 109_109 | 136_136 | NA | 282_309 | 245_245 | 71_79 |
| 109_109 | 136_136 | 184_184 | NA | 275_288 | 77_79 |
| 109_109 | 136_136 | 227_227 | 309_309 | 245_245 | 71_71 |
| 109_109 | 136_136 | NA | 284_307 | 245_254 | 71_79 |
| 107_109 | 136_143 | NA | 284_307 | 245_245 | 71_79 |
| 109_109 | 136_136 | 227_227 | NA | 245_245 | 71_79 |
| 109_109 | 136_136 | 227_227 | 282_309 | 245_245 | 71_79 |
| 109_109 | 136_136 | 227_227 | 282_307 | 245_245 | 71_79 |
| 109_109 | 130_136 | 225_225 | 286_309 | 245_256 | 71_79 |
| 109_109 | 136_136 | 233_248 | NA | 247_254 | 71_71 |
| 109_109 | 136_136 | 185_185 | NA | 245_275 | 77_79 |
| 107_109 | 136_140 | NA | 298_305 | 245_254 | 71_77 |
| 107_109 | 136_136 | 231_248 | 288_313 | 254_276 | 71_77 |
| 109_109 | 136_136 | 233_233 | NA | 275_288 | 77_79 |
| 107_109 | 136_136 | 185_187 | 304_305 | 245_254 | 71_79 |
| 107_109 | 136_136 | 185_185 | NA | 245_254 | 71_77 |
| NA | 136_136 | NA | 282_350 | 248_254 | NA |
| NA | 136_136 | NA | 309_321 | 247_254 | NA |
| 107_109 | 136_136 | 223_225 | 311_317 | 245_254 | 71_71 |
| 109_109 | 136_136 | 223_223 | 304_315 | 245_245 | 71_79 |
| 109_109 | 136_140 | 187_187 | 309_319 | 276_276 | 71_79 |
| 109_109 | 136_140 | 185_185 | 304_305 | 245_247 | 71_71 |
| 109_109 | 136_140 | 185_185 | 288_309 | 245_245 | 71_77 |
| 109_109 | 136_136 | NA | 302_325 | 244_250 | 71_71 |
| 109_109 | 136_136 | 229_229 | 282_346 | 245_245 | 71_71 |
| 109_109 | 136_140 | NA | 302_319 | 245_245 | 71_79 |
| 109_109 | 136_136 | 229_231 | 288_319 | 245_252 | 71_71 |
| 109_109 | 136_136 | 187_187 | 315_340 | 254_260 | 71_79 |
| 109_109 | 136_136 | NA | 309_319 | 254_254 | 71_79 |
| 109_109 | 136_136 | 197_197 | 307_311 | 245_245 | 71_77 |
| 109_109 | 136_140 | NA | 311_311 | 254_254 | 71_71 |
| 109_109 | 136_140 | 185_185 | 282_309 | 241_245 | 71_79 |
| 109_112 | 136_136 | NA | 282_309 | 241_245 | 71_79 |
| 107_109 | 130_136 | 187_225 | 282_288 | 245_245 | 71_71 |
| 109_109 | 136_140 | 235_245 | 309_331 | 245_267 | 71_79 |
| 107_109 | 136_136 | 183_245 | 309_368 | 245_245 | 71_71 |

| | | | | | |
|---|---|---|---|---|---|
| 109_109 | 136_136 | 187_187 | 282_286 | 245_245 | 71_71 |
| 109_109 | 136_140 | 189_189 | 284_302 | 245_254 | 71_71 |
| 107_109 | 136_136 | 241_241 | 307_309 | 245_254 | 71_71 |
| 107_109 | 136_136 | 185_187 | 278_309 | 245_254 | 71_79 |
| 107_109 | 130_140 | 221_221 | 278_346 | 248_256 | 71_71 |
| 107_109 | 136_140 | 185_202 | 305_307 | 254_254 | 71_77 |
| 109_109 | 130_140 | NA | 298_304 | NA | 71_71 |
| NA | 130_146 | NA | 280_286 | 245_251 | NA |
| 109_109 | 136_143 | 184_233 | 286_288 | 245_254 | 71_79 |
| 109_109 | 136_136 | 185_187 | 282_284 | 245_259 | 71_79 |
| 107_109 | 136_136 | 202_202 | NA | 244_245 | 71_79 |
| 109_109 | 136_136 | 185_185 | 286_302 | 245_252 | 71_71 |
| 109_109 | 136_136 | NA | 307_315 | 245_271 | 71_77 |
| 109_109 | 136_140 | 225_225 | 282_286 | 245_254 | 71_71 |
| 109_109 | 136_136 | 242_245 | 307_309 | 245_254 | 71_71 |
| 107_109 | 136_136 | 235_235 | 327_352 | 245_288 | 77_77 |
| 109_109 | 136_140 | 229_239 | 282_346 | 251_254 | 71_79 |
| 109_109 | 136_140 | NA | 282_286 | 245_245 | 71_71 |
| 107_109 | 136_136 | NA | 307_317 | 244_254 | 77_79 |
| 109_109 | 136_136 | 227_227 | 313_315 | 244_254 | 71_79 |
| 109_109 | 136_140 | 189_189 | 286_302 | 245_245 | 71_71 |
| 107_109 | 136_136 | 185_225 | 286_331 | 244_254 | 77_79 |
| 107_109 | 136_146 | 184_192 | 307_325 | 280_280 | 71_71 |
| 107_109 | 136_136 | 189_189 | 282_323 | 245_254 | 71_79 |
| 107_109 | 136_140 | 187_225 | 307_307 | 248_248 | 71_71 |
| 107_109 | 136_136 | NA | 282_282 | 245_245 | 71_71 |
| NA | 136_136 | NA | NA | 241_271 | NA |
| 107_109 | 136_136 | NA | 282_309 | 245_245 | 71_71 |
| 109_109 | 136_140 | 185_233 | 282_286 | 245_254 | 71_71 |
| 109_109 | 131_136 | 185_185 | 282_309 | 245_245 | 71_79 |
| 109_109 | 136_146 | 189_202 | 263_284 | 244_250 | 71_77 |
| 109_112 | 136_136 | NA | 284_284 | 245_254 | 71_71 |
| 107_109 | 136_136 | 184_235 | 282_305 | 250_250 | 71_77 |
| 109_109 | 130_140 | 187_187 | 286_305 | 245_245 | 71_79 |
| NA | 130_136 | NA | 286_292 | 278_278 | NA |
| NA | 136_146 | NA | 307_344 | 244_245 | NA |
| 107_109 | 127_136 | NA | NA | 246_254 | 71_71 |
| 107_109 | 136_140 | 245_245 | NA | 245_254 | 71_71 |
| 109_109 | 136_136 | 189_189 | 282_307 | 245_245 | 71_79 |
| 109_109 | 136_136 | 227_227 | 282_304 | 245_245 | 71_77 |
| 107_109 | 136_140 | 248_248 | NA | 254_254 | 71_77 |
| 107_109 | 136_136 | NA | 284_300 | 254_254 | 71_79 |
| 107_109 | 136_140 | 184_217 | 286_302 | 245_273 | 71_79 |
| 107_107 | 136_136 | 185_191 | 282_313 | 247_269 | 71_79 |
| 107_109 | 130_136 | 184_185 | 284_313 | 245_256 | 71_79 |
| 109_109 | 136_136 | 235_235 | 321_323 | 245_254 | 77_79 |
| 107_109 | 136_136 | 187_235 | 282_337 | 245_248 | 71_76 |
| 109_109 | 136_136 | 185_217 | 261_307 | 245_254 | 71_79 |

| | | | | | |
|---|---|---|---|---|---|
| NA | 136_136 | NA | 278_282 | 241_254 | NA |
| NA | 136_136 | NA | 300_311 | 245_248 | NA |
| 107_109 | 136_140 | 185_233 | 305_311 | 248_276 | 71_79 |
| 107_107 | 136_136 | 185_185 | 253_304 | 245_257 | 79_79 |
| 107_109 | 136_136 | 185_223 | NA | 259_265 | 71_79 |
| NA | 136_136 | NA | 300_327 | 245_245 | NA |
| 109_109 | 136_140 | 185_185 | 261_309 | 245_245 | 71_71 |
| 107_109 | 136_136 | 185_233 | NA | 243_254 | 71_79 |
| 109_109 | 136_140 | 220_220 | 305_342 | 254_254 | 71_71 |
| NA | 136_136 | NA | 307_311 | 245_286 | NA |
| 107_109 | 136_136 | 187_229 | 261_302 | 245_248 | 71_79 |
| NA | 136_136 | NA | 315_323 | 254_267 | NA |
| 107_109 | 136_136 | 187_194 | 282_305 | 248_248 | 71_71 |
| 109_109 | 136_136 | 187_233 | NA | 248_248 | 71_77 |
| 107_109 | 136_136 | 185_185 | 331_344 | 253_254 | 79_79 |
| 107_109 | 136_136 | 185_225 | NA | 245_254 | 71_79 |
| 107_109 | 136_136 | 233_233 | NA | 245_256 | 71_71 |
| 107_107 | 136_136 | 229_229 | 317_344 | 248_248 | 71_79 |
| 107_109 | 136_136 | 185_185 | 288_309 | 245_269 | 71_79 |
| 107_109 | 136_140 | 184_239 | NA | 245_251 | 77_77 |
| NA | 136_136 | NA | 288_315 | 244_249 | NA |
| 109_109 | 136_140 | 233_233 | 282_284 | 245_256 | 71_77 |
| NA | 136_136 | NA | 282_311 | 254_256 | NA |
| 109_109 | 136_136 | 184_185 | 317_319 | 244_244 | 71_79 |
| 109_109 | 136_136 | 216_216 | 282_284 | 244_269 | 71_77 |
| 107_109 | 136_140 | 185_185 | 313_338 | 244_244 | 71_79 |
| 109_109 | 140_146 | 185_185 | 323_335 | 245_254 | 71_79 |
| 109_109 | 136_140 | 185_242 | 284_323 | 245_248 | 79_79 |
| 107_107 | 136_136 | 185_185 | 282_300 | 256_276 | 71_71 |
| 107_109 | 136_136 | 185_185 | 282_282 | 245_265 | 71_71 |
| 107_107 | 130_136 | 185_227 | 315_323 | 245_254 | 71_81 |
| NA | 136_136 | NA | 261_329 | 245_265 | NA |
| 107_109 | 136_136 | 187_187 | 304_346 | 247_254 | 71_79 |
| 109_109 | 136_136 | 185_185 | NA | 245_275 | 71_71 |
| 109_109 | 136_136 | 187_214 | 284_311 | 248_269 | 77_77 |
| 107_107 | 136_136 | 206_208 | 261_282 | 254_271 | 71_79 |
| 107_109 | 136_140 | 187_187 | NA | 245_273 | 71_79 |
| 107_109 | 130_136 | 184_187 | 290_317 | 245_267 | 71_79 |
| 109_109 | 136_136 | 210_220 | 311_323 | 241_248 | 71_79 |
| 109_109 | 130_136 | 185_235 | 305_311 | 254_254 | 71_79 |
| 107_109 | 136_136 | 187_192 | NA | 245_254 | 71_71 |
| 107_109 | 136_140 | 184_187 | 253_307 | 245_248 | 71_71 |
| 107_107 | 136_136 | 185_185 | NA | 245_245 | 71_77 |
| 107_107 | 136_140 | 185_237 | 284_331 | 245_245 | 79_79 |
| 107_109 | 136_136 | 185_210 | 261_286 | 245_245 | 79_79 |
| 107_109 | 136_136 | 185_185 | 282_315 | 245_248 | 71_79 |
| 107_107 | 136_136 | 187_194 | 282_331 | 245_262 | 71_71 |
| 109_109 | 136_136 | 192_220 | 282_296 | 254_261 | 71_71 |

```
R version 4.2.2 (2022-10-31 ucrt)
Platform: x86_64-w64-mingw32/x64 (64-bit)
Running under: Windows 10 x64 (build 19045)

Matrix products: default

locale:
[1] LC_COLLATE=Turkish_Turkey.utf8  LC_CTYPE=Turkish_Turkey.utf8    LC_MONETARY=Turkish_Turkey.utf8 L

attached base packages:
[1] parallel  stats     graphics  grDevices utils     datasets  methods   base

other attached packages:
 [1] gradientForest_0.1-32 extendedForest_1.6.1  tidyr_1.3.0          envirem_2.3          palinsol_0.93        gsl_2.
 [8] lattice_0.21-8        maps_3.4.1            moments_0.14.1       cluster_2.1.4        rworldmap_1.3-6      RColorBr
[15] spam_2.9-1            gtools_3.9.4          raster_3.6-23        ggplot2_3.4.2        viridis_0.6.3        viridisLite_0.4.
[22] splancs_2.01-43       sp_1.6-0              adespatial_0.3-21    hierfstat_0.5-11     PopGenReport_3.0.7   knitr_
[29] sf_1.0-10             spData_2.2.2          pegas_1.2            ape_5.7-1            magrittr_2.0.3       poppr_2.9.4
[36] ade4_1.7-22           dplyr_1.1.2           pkgbuild_1.4.2       pak_0.4.0            usdm_2.1-6           terra_1.7-39

loaded via a namespace (and not attached):
  [1] uuid_1.1-0          plyr_1.8.8          igraph_1.4.1        splines_4.2.2       gstat_2.1-1         rncl_0.8.7          gap.dat
  [9] foreach_1.5.2       htmltools_0.5.4     gdata_2.18.0.1      fansi_1.0.4         doParallel_1.0.17   R.utils_2.12.2
 [17] jpeg_0.1-10         colorspace_2.1-0    mmod_1.3.3          xfun_0.37           rgdal_1.6-6         callr_3.7.3         cr
 [25] zoo_1.8-11          iterators_1.0.14    glue_1.6.2          gtable_0.3.1        seqinr_4.2-23       polysat_1.7-7       a
 [33] mvtnorm_1.1-3       DBI_1.1.3           GGally_2.1.2        shapefiles_0.7.2    Rcpp_1.0.10         xtable_1.8-4
 [41] foreign_0.8-83      proxy_0.4-27        dotCall64_1.0-2     intervals_0.15.3    dismo_1.3-9         httr_1.4.5
 [49] calibrate_1.7.7     wk_0.7.1            ellipsis_0.3.2      pkgconfig_2.0.3     reshape_0.8.9       XML_3.99-0.13
 [57] utf8_1.2.3          tidyselect_1.2.0    rlang_1.1.0         reshape2_1.4.4      later_1.3.0         munsell_0.5.0       a
 [65] cli_3.6.0           generics_0.1.3      evaluate_0.20       stringr_1.5.0       fastmap_1.1.1       yaml_2.3.7          pr
 [73] s2_1.1.2            RgoogleMaps_1.4.5.3 pbapply_1.7-0       nlme_3.1-160        mime_0.12           R.oo_1.25.0
 [81] compiler_4.2.2      beeswarm_0.4.0      png_0.1-8           e1071_1.7-13        RSAGA_1.4.0         spacetime_1.
 [89] stringi_1.7.12      gdistance_1.6       ps_1.7.2            Matrix_1.5-1        classInt_0.4-9      vegan_2.6-4         pe
 [97] pillar_1.9.0        lifecycle_1.0.3     combinat_0.0-8      maptools_1.1-6      cowplot_1.1.1       httpuv_1.6.9
[105] promises_1.2.0.1    KernSmooth_2.23-20  gridExtra_2.3       vipor_0.4.5         codetools_0.2-18    boot_1.3
[113] mgcv_1.8-41         hms_1.1.2           grid_4.2.2          class_7.3-20        rmarkdown_2.20      shiny_1.7.4
```

C_NUMERIC=C LC_TIME=Turkish_Turkey.utf8

| Locus | Alleles | Private alleles | Genotyping error | Null alleles |
|---|---|---|---|---|
| A007 | 53 | 15 | 1,79 | 0,03 |
| A014 | 10 | 2 | 2,68 | 0,10 |
| A028 | 15 | 4 | 6,26 | 0,08 |
| A043 | 16 | 3 | 3,06 | 0,05 |
| A079 | 14 | 5 | 0,89 | 0,03 |
| A088 | 14 | 5 | 0,49 | 0,07 |
| A107 | 44 | 6 | 4,00 | 0,01 |
| A113 | 18 | 2 | 0,97 | 0,05 |
| AB024 | 8 | 2 | 0,97 | 0,04 |
| AB124 | 19 | 5 | 3,28 | 0,03 |
| AC006 | 11 | 3 | 1,69 | 0,06 |
| AC306 | 12 | 3 | 1,39 | 0,05 |
| AP001 | 34 | 9 | 3,88 | 0,06 |
| AP019 | 11 | 2 | 2,12 | 0,00 |
| AP043 | 36 | 9 | 5,83 | 0,05 |
| AP049 | 18 | 7 | 3,07 | 0,08 |
| AP068 | 12 | 2 | 4,02 | -0,01 |
| AP218 | 4 | 1 | 0,00 | 0,01 |
| AP223 | 9 | 3 | 2,68 | 0,04 |
| AP226 | 8 | 3 | 1,34 | 0,10 |
| AP238 | 5 | 1 | 0,27 | 0,04 |
| AP243 | 12 | 6 | 0,98 | 0,01 |
| AP249 | 11 | 1 | 3,40 | 0,09 |
| AP273 | 7 | 2 | 1,32 | 0,01 |
| AP288 | 11 | 5 | 1,79 | 0,06 |
| AP289 | 49 | 11 | 4,42 | 0,12 |
| HB_C16_01 | 48 | 8 | 1,35 | 0,02 |
| HB_C16_02 | 47 | 3 | 6,61 | 0,08 |
| HB_C16_05 | 18 | 12 | 1,34 | 0,04 |

| Expected H | Observed H | Fis | Fit | Fst |
|---:|---:|---:|---:|---:|
| 0,95 | 0,89 | 0,03 | 0,07 | 0,04 |
| 0,65 | 0,48 | 0,08 | 0,27 | 0,21 |
| 0,39 | 0,28 | 0,25 | 0,29 | 0,06 |
| 0,54 | 0,46 | 0,10 | 0,15 | 0,06 |
| 0,78 | 0,73 | 0,01 | 0,07 | 0,07 |
| 0,68 | 0,57 | 0,04 | 0,17 | 0,14 |
| 0,94 | 0,93 | 0,00 | 0,02 | 0,02 |
| 0,89 | 0,79 | 0,01 | 0,13 | 0,12 |
| 0,60 | 0,54 | 0,09 | 0,10 | 0,02 |
| 0,80 | 0,74 | 0,03 | 0,08 | 0,06 |
| 0,25 | 0,18 | 0,04 | 0,30 | 0,27 |
| 0,75 | 0,67 | 0,00 | 0,12 | 0,12 |
| 0,78 | 0,67 | 0,04 | 0,15 | 0,12 |
| 0,33 | 0,33 | -0,01 | 0,02 | 0,03 |
| 0,87 | 0,77 | 0,08 | 0,12 | 0,05 |
| 0,43 | 0,31 | 0,13 | 0,30 | 0,19 |
| 0,65 | 0,66 | -0,03 | -0,02 | 0,01 |
| 0,26 | 0,25 | -0,03 | 0,05 | 0,08 |
| 0,67 | 0,60 | 0,03 | 0,12 | 0,09 |
| 0,35 | 0,21 | 0,08 | 0,43 | 0,38 |
| 0,45 | 0,39 | 0,01 | 0,14 | 0,14 |
| 0,16 | 0,15 | 0,06 | 0,07 | 0,01 |
| 0,75 | 0,59 | 0,07 | 0,22 | 0,16 |
| 0,46 | 0,44 | -0,02 | 0,05 | 0,07 |
| 0,42 | 0,34 | 0,03 | 0,21 | 0,19 |
| 0,91 | 0,68 | 0,21 | 0,26 | 0,05 |
| 0,94 | 0,91 | 0,00 | 0,04 | 0,04 |
| 0,85 | 0,71 | 0,14 | 0,17 | 0,03 |
| 0,66 | 0,60 | 0,03 | 0,09 | 0,06 |

| Evenness | G'st | Gst | Jost's D |
|---|---|---|---|
| 0,68 | 0,43 | 0,03 | 0,41 |
| 0,70 | 0,31 | 0,13 | 0,20 |
| 0,43 | 0,10 | 0,06 | 0,04 |
| 0,52 | 0,12 | 0,05 | 0,07 |
| 0,76 | 0,33 | 0,09 | 0,26 |
| 0,73 | 0,28 | 0,11 | 0,19 |
| 0,68 | 0,27 | 0,02 | 0,25 |
| 0,82 | 0,48 | 0,08 | 0,43 |
| 0,73 | 0,16 | 0,06 | 0,10 |
| 0,62 | 0,28 | 0,06 | 0,23 |
| 0,43 | 0,38 | 0,29 | 0,11 |
| 0,78 | 0,19 | 0,05 | 0,14 |
| 0,48 | 0,25 | 0,06 | 0,20 |
| 0,45 | 0,04 | 0,03 | 0,01 |
| 0,54 | 0,29 | 0,06 | 0,24 |
| 0,46 | 0,35 | 0,21 | 0,16 |
| 0,56 | 0,01 | 0,00 | 0,01 |
| 0,55 | 0,06 | 0,04 | 0,02 |
| 0,80 | 0,23 | 0,08 | 0,16 |
| 0,49 | 0,40 | 0,29 | 0,12 |
| 0,77 | 0,26 | 0,15 | 0,12 |
| 0,38 | 0,02 | 0,01 | 0,00 |
| 0,68 | 0,56 | 0,21 | 0,44 |
| 0,76 | 0,13 | 0,07 | 0,06 |
| 0,52 | 0,35 | 0,22 | 0,15 |
| 0,54 | 0,45 | 0,06 | 0,42 |
| 0,63 | 0,47 | 0,04 | 0,45 |
| 0,40 | 0,22 | 0,04 | 0,19 |
| 0,78 | 0,20 | 0,07 | 0,13 |

| | Eur | Co.Aeg | EC.Anat |
|---|---|---|---|
| **A107:AC306** | | | 0,0001 |
| **AB124:AP288** | | 0,0001 | |
| **AC006:HB_C16_02** | 0,0001 | | |
| **HB_C16_01:HB_C16_02** | 0,0001 | | |

| Population | N | Private alleles | Mean richness | Total richness | H |
|---|---|---|---|---|---|
| C.BlkS | 34 | 1 | 1,42 | 41,17 | 3,53 |
| Co.Aeg | 55 | 8 | 1,59 | 46,16 | 4,01 |
| E.Med | 80 | 23 | 1,54 | 44,55 | 4,38 |
| EC.Anat | 29 | 6 | 1,46 | 42,36 | 3,37 |
| Erz-Kar | 51 | 6 | 1,60 | 46,34 | 3,93 |
| ES.Marm | 29 | 6 | 1,63 | 47,23 | 3,37 |
| Eur | 38 | 28 | 1,54 | 44,80 | 3,64 |
| L.Cauc | 44 | 8 | 1,58 | 45,76 | 3,78 |
| Thrace | 77 | 25 | 1,63 | 47,38 | 4,34 |
| U.Euph | 74 | 11 | 1,58 | 45,87 | 4,30 |
| W.Anat | 65 | 6 | 1,57 | 45,49 | 4,17 |
| W.BlkS | 53 | 5 | 1,58 | 45,72 | 3,97 |
| Zagros | 43 | 7 | 1,57 | 45,64 | 3,76 |

| Hexp | Hobs | Fis | Fst |
|---|---|---|---|
| 0,51 | 0,50 | 0,05 | 0,15 |
| 0,60 | 0,58 | -0,03 | 0,11 |
| 0,54 | 0,50 | 0,05 | 0,08 |
| 0,47 | 0,43 | 0,06 | 0,20 |
| 0,66 | 0,61 | 0,15 | 0,10 |
| 0,64 | 0,62 | 0,00 | -0,05 |
| 0,55 | 0,54 | 0,02 | 0,03 |
| 0,58 | 0,54 | 0,11 | 0,05 |
| 0,64 | 0,62 | 0,03 | -0,05 |
| 0,59 | 0,55 | 0,02 | 0,11 |
| 0,57 | 0,56 | 0,00 | 0,09 |
| 0,58 | 0,53 | 0,06 | 0,10 |
| 0,58 | 0,53 | 0,05 | 0,04 |

| | Eur | Thrace | ES.Marm | Co.Aeg | W.Anat | W.BlkS | C.BlkS | L.Cauc |
|---|---|---|---|---|---|---|---|---|
| **Eur** | NA | NA | NA | NA | NA | NA | NA | NA |
| **Thrace** | 0,39 | NA | NA | NA | NA | NA | NA | NA |
| **ES.Marm** | 0,46 | 0,00 | NA | NA | NA | NA | NA | NA |
| **Co.Aeg** | 0,65 | 0,11 | 0,06 | NA | NA | NA | NA | NA |
| **W.Anat** | 0,66 | 0,13 | 0,08 | 0,02 | NA | NA | NA | NA |
| **W.BlkS** | 0,67 | 0,15 | 0,09 | 0,02 | 0,02 | NA | NA | NA |
| **C.BlkS** | 0,75 | 0,24 | 0,18 | 0,08 | 0,14 | 0,09 | NA | NA |
| **L.Cauc** | 0,70 | 0,27 | 0,22 | 0,13 | 0,16 | 0,15 | 0,06 | NA |
| **Erz-Kar** | 0,64 | 0,11 | 0,06 | -0,01 | 0,02 | 0,03 | -0,02 | -0,02 |
| **U.Euph** | 0,66 | 0,14 | 0,09 | 0,02 | 0,02 | 0,02 | 0,05 | 0,10 |
| **EC.Anat** | 0,74 | 0,28 | 0,24 | 0,16 | 0,14 | 0,13 | 0,20 | 0,28 |
| **E.Med** | 0,69 | 0,18 | 0,13 | 0,05 | 0,05 | 0,04 | 0,09 | 0,15 |
| **Zagros** | 0,69 | 0,19 | 0,13 | 0,05 | 0,04 | 0,03 | 0,08 | 0,08 |

| | Eur | Thrace | ES.Marm | Co.Aeg | W.Anat | W.BlkS | C.BlkS | L.Cauc |
|---|---|---|---|---|---|---|---|---|
| **Eur** | NA | NA | NA | NA | NA | NA | NA | NA |
| **Thrace** | 0,09 | NA | NA | NA | NA | NA | NA | NA |
| **ES.Marm** | 0,10 | 0,00 | NA | NA | NA | NA | NA | NA |
| **Co.Aeg** | 0,16 | 0,02 | 0,01 | NA | NA | NA | NA | NA |
| **W.Anat** | 0,17 | 0,03 | 0,02 | 0,00 | NA | NA | NA | NA |
| **W.BlkS** | 0,17 | 0,03 | 0,02 | 0,00 | 0,00 | NA | NA | NA |
| **C.BlkS** | 0,21 | 0,05 | 0,04 | 0,02 | 0,03 | 0,02 | NA | NA |
| **L.Cauc** | 0,18 | 0,05 | 0,04 | 0,03 | 0,03 | 0,03 | 0,01 | NA |
| **Erz-Kar** | 0,15 | 0,02 | 0,01 | 0,00 | 0,00 | 0,01 | 0,00 | 0,00 |
| **U.Euph** | 0,17 | 0,03 | 0,02 | 0,00 | 0,01 | 0,00 | 0,01 | 0,02 |
| **EC.Anat** | 0,22 | 0,07 | 0,06 | 0,04 | 0,04 | 0,03 | 0,05 | 0,07 |
| **E.Med** | 0,19 | 0,04 | 0,03 | 0,01 | 0,01 | 0,01 | 0,02 | 0,03 |
| **Zagros** | 0,18 | 0,04 | 0,03 | 0,01 | 0,01 | 0,01 | 0,02 | 0,02 |

| | Eur | Thrace | ES.Marm | Co.Aeg | W.Anat | W.BlkS | C.BlkS | L.Cauc |
|---|---|---|---|---|---|---|---|---|
| **Eur** | NA | NA | NA | NA | NA | NA | NA | NA |
| **Thrace** | 0,28 | NA | NA | NA | NA | NA | NA | NA |
| **ES.Marm** | 0,33 | 0,00 | NA | NA | NA | NA | NA | NA |
| **Co.Aeg** | 0,52 | 0,07 | 0,04 | NA | NA | NA | NA | NA |
| **W.Anat** | 0,52 | 0,08 | 0,05 | 0,01 | NA | NA | NA | NA |
| **W.BlkS** | 0,54 | 0,09 | 0,05 | 0,01 | 0,01 | NA | NA | NA |
| **C.BlkS** | 0,61 | 0,16 | 0,11 | 0,05 | 0,08 | 0,05 | NA | NA |
| **L.Cauc** | 0,57 | 0,18 | 0,15 | 0,08 | 0,10 | 0,09 | 0,03 | NA |
| **Erz-Kar** | 0,51 | 0,07 | 0,04 | -0,01 | 0,01 | 0,02 | -0,01 | -0,01 |
| **U.Euph** | 0,52 | 0,09 | 0,06 | 0,01 | 0,01 | 0,01 | 0,03 | 0,06 |
| **EC.Anat** | 0,59 | 0,18 | 0,15 | 0,09 | 0,08 | 0,07 | 0,11 | 0,17 |
| **E.Med** | 0,55 | 0,11 | 0,08 | 0,03 | 0,03 | 0,02 | 0,05 | 0,09 |
| **Zagros** | 0,56 | 0,12 | 0,09 | 0,03 | 0,03 | 0,02 | 0,05 | 0,05 |

| | Eur | Thrace | ES.Marm | Co.Aeg | W.Anat | W.BlkS | C.BlkS | L.Cauc |
|---|---|---|---|---|---|---|---|---|
| **Eur** | NA | NA | NA | NA | NA | NA | NA | NA |
| **Thrace** | 0,16 | NA | NA | NA | NA | NA | NA | NA |
| **ES.Marm** | 0,19 | 0,00 | NA | NA | NA | NA | NA | NA |
| **Co.Aeg** | 0,28 | 0,04 | 0,02 | NA | NA | NA | NA | NA |

| | | | | | | | | |
|---|---|---|---|---|---|---|---|---|
| **W.Anat** | 0,29 | 0,05 | 0,03 | 0,01 | NA | NA | NA | NA |
| **W.BlkS** | 0,29 | 0,06 | 0,03 | 0,01 | 0,01 | NA | NA | NA |
| **C.BlkS** | 0,35 | 0,09 | 0,06 | 0,02 | 0,05 | 0,01 | NA | NA |
| **L.Cauc** | 0,30 | 0,10 | 0,09 | 0,05 | 0,07 | 0,06 | 0,01 | NA |
| **Erz-Kar** | 0,26 | 0,04 | 0,02 | -0,01 | 0,01 | 0,00 | -0,06 | -0,01 |
| **U.Euph** | 0,28 | 0,05 | 0,04 | 0,01 | 0,01 | 0,01 | 0,01 | 0,04 |
| **EC.Anat** | 0,36 | 0,12 | 0,11 | 0,07 | 0,07 | 0,06 | 0,10 | 0,13 |
| **E.Med** | 0,31 | 0,07 | 0,05 | 0,02 | 0,02 | 0,02 | 0,02 | 0,07 |
| **Zagros** | 0,30 | 0,07 | 0,05 | 0,02 | 0,02 | 0,01 | 0,02 | 0,03 |

| Erz-Kar | U.Euph | EC.Anat | E.Med | Zagros |
|---|---|---|---|---|
| NA | NA | NA | NA | NA |
| NA | NA | NA | NA | NA |
| NA | NA | NA | NA | NA |
| NA | NA | NA | NA | NA |
| NA | NA | NA | NA | NA |
| NA | NA | NA | NA | NA |
| NA | NA | NA | NA | NA |
| NA | NA | NA | NA | NA |
| NA | NA | NA | NA | NA |
| -0,03 | NA | NA | NA | NA |
| 0,19 | 0,14 | NA | NA | NA |
| 0,04 | 0,03 | 0,06 | NA | NA |
| 0,00 | 0,01 | 0,14 | 0,04 | NA |

| Erz-Kar | U.Euph | EC.Anat | E.Med | Zagros |
|---|---|---|---|---|
| NA | NA | NA | NA | NA |
| NA | NA | NA | NA | NA |
| NA | NA | NA | NA | NA |
| NA | NA | NA | NA | NA |
| NA | NA | NA | NA | NA |
| NA | NA | NA | NA | NA |
| NA | NA | NA | NA | NA |
| NA | NA | NA | NA | NA |
| NA | NA | NA | NA | NA |
| -0,01 | NA | NA | NA | NA |
| 0,05 | 0,03 | NA | NA | NA |
| 0,01 | 0,01 | 0,02 | NA | NA |
| 0,00 | 0,00 | 0,03 | 0,01 | NA |

| Erz-Kar | U.Euph | EC.Anat | E.Med | Zagros |
|---|---|---|---|---|
| NA | NA | NA | NA | NA |
| NA | NA | NA | NA | NA |
| NA | NA | NA | NA | NA |
| NA | NA | NA | NA | NA |
| NA | NA | NA | NA | NA |
| NA | NA | NA | NA | NA |
| NA | NA | NA | NA | NA |
| NA | NA | NA | NA | NA |
| NA | NA | NA | NA | NA |
| -0,02 | NA | NA | NA | NA |
| 0,12 | 0,08 | NA | NA | NA |
| 0,02 | 0,02 | 0,03 | NA | NA |
| 0,00 | 0,01 | 0,08 | 0,02 | NA |

| Erz-Kar | U.Euph | EC.Anat | E.Med | Zagros |
|---|---|---|---|---|
| NA | NA | NA | NA | NA |
| NA | NA | NA | NA | NA |
| NA | NA | NA | NA | NA |
| NA | NA | NA | NA | NA |

| | | | | |
|---|---|---|---|---|
| NA | NA | NA | NA | NA |
| NA | NA | NA | NA | NA |
| NA | NA | NA | NA | NA |
| NA | NA | NA | NA | NA |
| NA | NA | NA | NA | NA |
| -0,01 | NA | NA | NA | NA |
| 0,09 | 0,07 | NA | NA | NA |
| 0,01 | 0,01 | 0,03 | NA | NA |
| 0,00 | 0,01 | 0,07 | 0,02 | NA |

| Subset | Test | Obs | Std Obs | Expectation | Variance | Alter | Pvalue |
|---|---|---|---|---|---|---|---|
| all | Variations within samples | 5,13 | -8,19 | 5,70 | 0,07 | less | 0,01 |
| all | Variations between samples | 0,33 | 4,82 | 0,00 | 0,07 | greater | 0,01 |
| all | Variations between Population | 0,32 | 40,01 | 0,00 | 0,01 | greater | 0,01 |
| t-a | Variations within samples | 7,29 | -3,21 | 7,62 | 0,10 | less | 0,01 |
| t-a | Variations between samples | 0,21 | 1,47 | -0,01 | 0,15 | greater | 0,10 |
| t-a | Variations between Population | 0,30 | 17,05 | 0,00 | 0,02 | greater | 0,01 |
| z-a | Variations within samples | 2,95 | -1,75 | 3,07 | 0,07 | less | 0,03 |
| z-a | Variations between samples | 0,09 | 1,39 | 0,00 | 0,07 | greater | 0,13 |
| z-a | Variations between Population | 0,04 | 5,33 | 0,00 | 0,01 | greater | 0,01 |
| z-l | Variations within samples | 2,68 | -2,24 | 2,81 | 0,06 | less | 0,03 |
| z-l | Variations between samples | 0,12 | 2,25 | 0,00 | 0,05 | greater | 0,04 |
| z-l | Variations between Population | 0,02 | 2,26 | 0,00 | 0,01 | greater | 0,03 |
| z-c | Variations within samples | 2,87 | -4,21 | 3,18 | 0,07 | less | 0,01 |
| z-c | Variations between samples | 0,29 | 4,73 | 0,00 | 0,06 | greater | 0,01 |
| z-c | Variations between Population | 0,03 | 2,00 | 0,00 | 0,01 | greater | 0,04 |

| Var | Global | A to T | A to C | A to L to Z | C to Z |
|---|---|---|---|---|---|
| MEM2 | 0,1849 | 0,1937 | 0,0165 | 0,2017 | 0,1925 |
| MEM1 | 0,1164 | 0,2404 | 0,1468 | 0,1249 | 0,0401 |
| PETwettest | 0,0240 | 0,0193 | 0,0053 | 0,0068 | 0,0095 |
| Pwarmest | 0,0220 | 0,0068 | 0,1102 | 0,0029 | 0,0647 |
| isothermality | 0,0177 | 0,0153 | 0,0199 | 0,0129 | 0,0331 |
| Tdriest | 0,0176 | 0,0039 | 0,0479 | 0,0071 | 0,0140 |
| aridity | 0,0153 | 0,0027 | 0,0413 | 0,0055 | 0,0268 |
| Tseasonality | 0,0143 | 0,0034 | 0,0066 | 0,0096 | 0,0096 |
| TUR_alt | 0,0143 | 0,0071 | 0,0169 | 0,0159 | 0,0171 |
| continentality | 0,0135 | 0,0067 | 0,0054 | 0,0108 | 0,0056 |
| Pwettest | 0,0123 | 0,0049 | 0,0248 | 0,0025 | 0,0268 |
| Pdriest | 0,0123 | 0,0238 | 0,0070 | 0,0044 | 0,0124 |
| minTwarm | 0,0114 | 0,0030 | 0,0021 | 0,0157 | 0,0021 |
| PETdriest | 0,0110 | 0,0030 | 0,0221 | 0,0024 | 0,0133 |
| Pcoldest | 0,0105 | 0,0127 | 0,0040 | 0,0025 | 0,0038 |
| minTcold | 0,0097 | 0,0064 | 0,0038 | 0,0096 | 0,0056 |
| Pseasonality | 0,0082 | 0,0046 | 0,0032 | 0,0066 | 0,0111 |
| Pdry | 0,0079 | 0,0122 | 0,0090 | 0,0025 | 0,0094 |
| Twarmest | 0,0076 | 0,0024 | 0,0005 | 0,0099 | 0,0018 |
| Tcoldest | 0,0076 | 0,0058 | 0,0048 | 0,0059 | 0,0064 |
| Pwet | 0,0070 | 0,0051 | 0,0062 | 0,0031 | 0,0065 |
| maxTcold | 0,0067 | 0,0036 | 0,0041 | 0,0076 | 0,0023 |
| diurnalTrange | 0,0064 | 0,0085 | 0,0075 | 0,0061 | 0,0044 |
| PETwarmest | 0,0063 | 0,0057 | 0,0102 | 0,0042 | 0,0112 |
| meanT | 0,0061 | 0,0016 | 0,0027 | 0,0060 | 0,0030 |
| Twettest | 0,0061 | 0,0037 | 0,0024 | 0,0055 | 0,0036 |
| gdd0 | 0,0058 | 0,0025 | 0,0019 | 0,0057 | 0,0020 |
| annualTrange | 0,0058 | 0,0040 | 0,0050 | 0,0072 | 0,0043 |
| annualP | 0,0056 | 0,0102 | 0,0067 | 0,0020 | 0,0049 |
| roughness | 0,0054 | 0,0039 | 0,0044 | 0,0032 | 0,0047 |
| PETseasonality | 0,0052 | 0,0084 | 0,0052 | 0,0038 | 0,0070 |
| embergerQ | 0,0052 | 0,0082 | 0,0047 | 0,0070 | 0,0020 |
| maxTwarm | 0,0049 | 0,0026 | 0,0030 | 0,0054 | 0,0043 |
| annualPET | 0,0046 | 0,0017 | 0,0038 | 0,0041 | 0,0059 |
| PETcoldest | 0,0045 | 0,0043 | 0,0025 | 0,0030 | 0,0030 |
| thermicity | 0,0045 | 0,0013 | 0,0026 | 0,0041 | 0,0021 |
| moisture | 0,0038 | 0,0031 | 0,0027 | 0,0040 | 0,0049 |
| gdd5 | 0,0030 | 0,0014 | 0,0030 | 0,0044 | 0,0022 |
| topoWet | 0,0018 | 0,0030 | 0,0016 | 0,0019 | 0,0035 |
| count10 | 0,0013 | 0,0006 | 0,0006 | 0,0016 | 0,0004 |

| Model | All Variables | Environmental | Important | Selected | Thracian | Anatolian | Levantine |
|---|---|---|---|---|---|---|---|
| Global | 0,64 | 0,34 | 0,50 | 0,20 | 0,73 | 0,64 | 0,74 |
| A to T | 0,66 | 0,23 | 0,53 | 0,09 | 0,75 | 0,57 | NA |
| A to C | 0,58 | 0,42 | 0,46 | 0,29 | NA | 0,52 | NA |
| A to L to Z | 0,55 | 0,22 | 0,38 | 0,06 | NA | 0,58 | 0,76 |
| C to Z | 0,59 | 0,36 | 0,46 | 0,23 | NA | NA | NA |

| Caucasian | Zagrosian |
|---|---|
| 0,67 | 0,41 |
| NA | NA |
| 0,64 | NA |
| NA | 0,31 |
| 0,70 | 0,48 |
